# Mapping of Quorum Sensing Landscape of Commensal and Pathogenic Staphylococci Reveals a Largely Inhibitory Interaction Network

**DOI:** 10.1101/2025.01.30.635662

**Authors:** Bengt H. Gless, Benjamin S. Sereika-Bejder, Iben Jensen, Martin S. Bojer, Katerina Tsiko, Sabrina H. Schmied, Ludovica Vitolo, Bruno Toledo-Silva, Sarne De Vliegher, Hanne Ingmer, Christian A. Olsen

**Affiliations:** Center for Biopharmaceuticals and Department of Drug Design and Pharmacology, Faculty of Health and Medical Sciences, University of Copenhagen, Jagtvej 160, DK-2100, Copenhagen, Denmark; Department of Veterinary and Animal Sciences, Faculty of Health and Medical Sciences, University of Copenhagen, Stigbøjlen 4, DK-1870 Frederiksberg C, Denmark; M-team & Mastitis and Milk Quality Research Unit, Department of Internal Medicine, Reproduction and Population Medicine, Faculty of Veterinary Medicine, Ghent University, Salisburylaan 133, B-9820 Merelbeke, Belgium

**Keywords:** Staphylococci, autoinducing peptides, cyclic peptides, RiPPs, virulence, MRSA

## Abstract

Staphylococci utilize secreted autoinducing peptides (AIPs) to regulate group behaviour through a process called quorum sensing (QS). For staphylococcal pathogens such as *S. aureus*, QS regulates expression of major virulence factors and QS inhibition has been proposed as an alternative to antibiotics for treatment of infections with methicillin resistant *S. aureus* (MRSA). Here, we surveyed the interaction map between QS systems of the pathogens *Staphylococcus aureus, Staphylococcus epidermidis*, and *Staphylococcus lugdunensis* and the 36 currently known AIPs from 22 staphylococcal species. We identified seven of these ribosomally synthesized and post-translationally modified peptides (RiPPs) in this study and all synthetic peptides were assessed for their ability to modulate QS. The mapped interactions of >280 native QS pairings were divided into human- and animal-associated staphylococci showing substantial differences in inhibitory potencies between the groups. In particular, AIPs of the bovine-associated species *S. simulans* displayed potential as QS inhibitors in the strains investigated in this study and were therefore chosen as starting point for a structure–activity relationship study. This study provides insights into the requirements for QS interference, yielding the most potent inhibitors reported to date for *S. epidermidis* and *S. lugdunensis*. Further, we tested an *S. simulans* AIP as anti-virulence agent in an assay to assess risk of acquired suppression of the inhibitory effect, and we established an assay set-up to successfully monitor *agr* deactivation of virulent MRSA by the QS inhibitor. Finally, a peptide was shown to attenuate skin infection caused by MRSA in a mouse model. Our results reveal a complex network of staphylococcal interactions and provide further impetus for the development of therapeutic strategies, based on QS modulation to target antibiotic-resistant pathogens.

**TOC Graphic:** 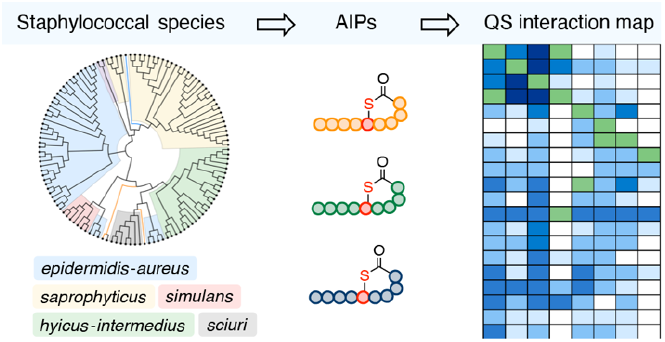

## INTRODUCTION

Staphylococci are Gram-positive bacteria that frequently colonize humans and animals representing some of the most abundant microbes found in the human microbiota.^1^ Among the numerous staphylococcal species, there are harmless commensal species while others, especially *S. aureus*, are pathogenic. All staphylococci have genes encoding a quorum sensing (QS) system that enables changes in group behavior and gene expression in response to cell density.^2, 3^ This cell-to-cell communication plays an important role in the transition from harmless skin colonizer to invasive pathogen and is regulated through the secretion and detection of autoinducing peptides (AIPs), which are 7–12 residue peptides, containing a characteristic thiolactone peptide cycle at the C-terminus (lactone for *S. intermedius* group).^4–6^ The AIP-mediated QS machinery is encoded by a chromosomal locus termed *accessory gene regulator* (*agr*), which controls the expression of genes involved in biofilm formation, surface adhesion and toxin production as well as the Agr proteins involved in the QS process (Supplementary Figure S1).^5, 6^ The *agr* system has been studied in detail for *S. aureus* but *agr* loci are found in all staphylococci, suggesting that each species utilizes a unique AIP as QS signaling molecule. Another key feature of AIP secretion is QS interference with *agr* systems of other staphylococcal species and *agr* specificity groups within the same species.^2, 7^ This phenomenon has been thoroughly studied for *S. aureus* and many non-cognate AIPs act as potent inhibitors to its QS system.^8–15^ QS interference has been less studied in other staphylococci; however, it might be a common occurrence in shared habitats of staphylococci, resulting in altered gene expression levels of co-inhabiting species susceptible to *agr* inhibition by non-cognate AIPs (Figure 1).

**Figure 1.**
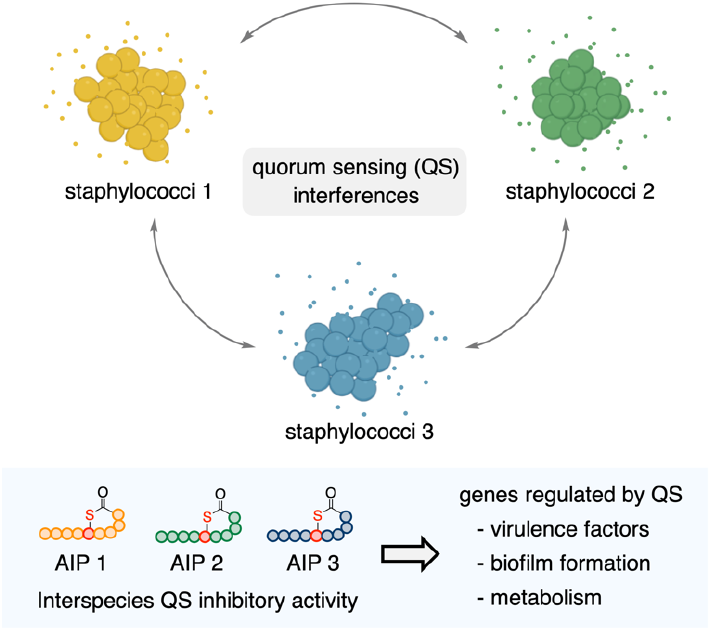
Quorum sensing interference of co-inhabiting staphylococci. The QS interference within habitats of multiple staphylococcal species is complex as each staphylococcal species and *agr* specificity group secretes a unique AIP. Co-inhabiting staphylococci are exposed to non-cognate AIPs, which can interfere with their QS systems depending on their inhibitory potency and thereby alter gene expression and change group behavior.

Determining the influence of QS interference on bacterial multi-species human and animal microbiotas is difficult due to their complex nature and the multitude of non-QS interactions.^3^ Nevertheless, recent advances focused on QS interference with *S. aureus* by commensal staphylococci in the context of atopic dermatitis and therapy.^12, 16, 17^ Early-phase clinical studies have shown a beneficial effect from the secreted inhibitory AIPs of *S. hominis* on the outcome of atopic dermatitis caused by *S. aureus* as a result of QS inhibition.^18^ These are promising results from the perspective of investigating anti-virulence strategies based on QS inhibition as alternatives to traditional treatment with antibiotics or as synergistic options together with antibiotics.^19, 20^

QS inhibitor development had and still has a focus on the AIPs of *S. aureus* and structure–activity relationship studies thereof, which afforded potent inhibitors of *S. aureus* itself.^21–24^ However, recent developments in technologies for the identification of AIP, has led to a significant increase of known non-aureus staphylococcal and mammaliicoccal AIPs.^10–15,25^ Several of these AIPs are potent QS inhibitors of *S. aureus* displaying effective *in vivo* attenuation of infections caused by methicillin-resistant strains of *S. aureus* (MRSA).^10,13–15^ The effects of non-cognate AIPs on *agr* systems different from *S. aureus* have been more scarcely investigated, including that of the common skin colonizer *S. epidermidis*, despite its abundance on the human skin.^26–29^ The roles of *S. epidermidis* as symbiont are manifold^30^ with recent studies even showing a beneficial role for the host,^31, 32^ while at the same time being able to cause medical device infections.^33^ A few studies have investigated modifications to the cognate AIPs, to create activators and inhibitors of its *agr* system.^28, 34, 35^ Similarly, the human skin commensal *S. lugdunensis* has been reported to cause severe endocarditis^36^ and QS interference with its *agr* system has been rarely investigated.^37^

Thus, the substantial increase in recently identified AIPs, combined with the lack of exploration of their interactions, encouraged us to systematically map the QS interactions of all known AIPs (**1**–**36**) with eight *agr* reporter strains of three therapeutically relevant species *S. aureus, S. epidermidis*, and *S. lugdunensis*. Our motivation was two-fold: first, to create a defining data set of QS interactions as a resource to explore trends between bacterial species in the same habitat; and secondly, to discover potent inhibitory interactions, especially against the less studied *agr* systems of *S. epidermidis* and *S. lugdunensis*.

## RESULTS

### Identification of new autoinducing peptides and quorum sensing interaction map

We previously developed a native chemical ligation-based trapping method for the rapid identification of AIPs from bacterial supernatants, by exploiting the chemo-selective reaction between thioesters and N-terminal cysteine residues,^38^ which led to a substantial increase in the number of known AIPs (Supplementary Figure S2).^11^ Here, we report the additional identification of seven previously unidentified AIPs from a collection of human and animal isolates, namely: *S. capitis* AIP-I (**15**), *S. cohnii* AIP-I (**19**), *S. pasteuri* AIP-I (**22**), *S. devriesei* AIP-I (**23**), *S. succinus* AIP-I (**24**), as well as *S. equorum* AIPs I (**25**) and II (**26**) (Supplementary Figure S2–9). This elevates the number of currently known staphylococcal AIPs to 38 [36 unique structures (**1**–**36**)], originating from 22 staphylococcal species from across 5 of the 6 phylogenetic species groups as classified through multi-locus sequence typing (Supplementary Table S1).^39^ In order to create a comparative and reproducible data set of QS interactions of all known AIPs against *S. aureus, S. epidermidis*, and *S. lugdunensis*, we utilized widely-used fluorescent reporter strains, which have a naturally functioning *agr* system that produces GFP/YFP once the *agr* dependent promotor P3 is activated.^40^ As a first step, we compiled our library of 36 unique AIPs by chemical synthesis (Supplementary Schemes S1–3) and established an assay setup in which the peptides were initially screened at 1 μM and at 50 nM concentration. In cases where we observed >95% inhibition at 50 nM AIP concentration, lower concentrations of 2.5 nM and 0.125 nM were tested (Figure 2 and Supplementary Figures S10–17).

**Figure 2.**
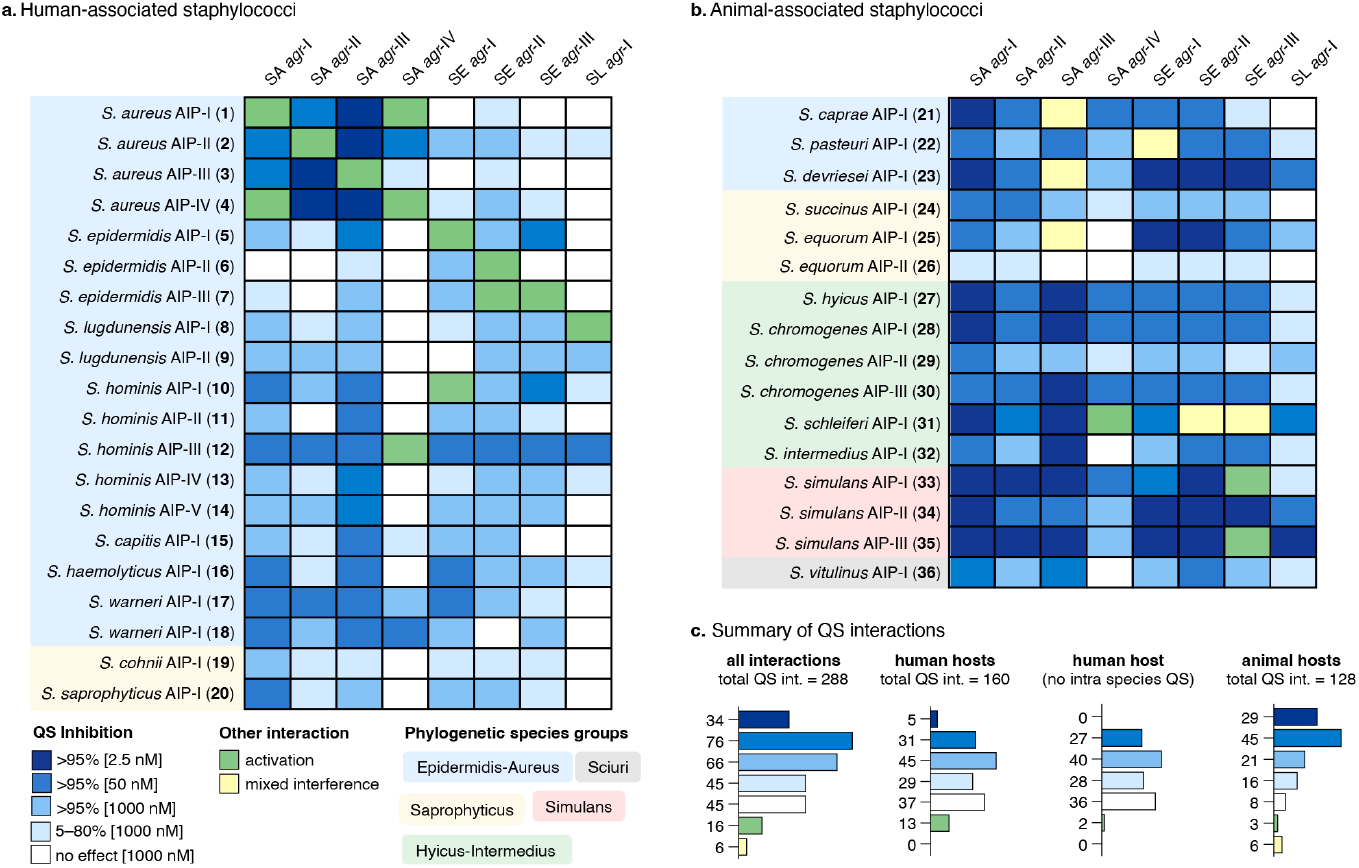
Quorum sensing (QS) interaction map. **a**) Map of human-associated staphylococci. **b**) Map of animal-associated staphylococci. **c**) Summary of QS interactions with human and animal hosts. Synthetic AIPs were tested at several concentrations (1000 nM, 50 nM, 2.5 nM) against fluorescent reporter strains of *S. aureus* (SA) *agr*-I–IV, *S. epidermidis* (SE) *agr*-I–III, and *S. lugdunensis* (SL) *agr*-I to assess their QS modulation abilities. Blue shading of boxes represents different potencies of QS inhibition (>95% at 2.5 nM, 50 nM and 1000 nM or 5–80% at 1000 nM), white boxes represent no interaction at 1000 nM, green boxes represent QS activation and yellow boxes represent “mixed interference” (activation at 1000 nM and inhibition at 50 nM or partial inhibition over a range of concentrations).

To make sure that our systematic survey, using fluorescent reporter strains, correlated with previously reported QS inhibition values, we determined half maximal inhibitory concentration (IC_50_) values for selected AIPs in both fluorescent reporter strain assays (Supplementary Figures S18–20) and *β*-lactamase reporter strain assays for QS inhibition of *S. aureus agr*-I–IV (Supplementary Figures S21– 24),^11^ which showed excellent correlation between single concentration data points and full dose– response experiments as well as with previously reported QS inhibition values collected from the literature (Supplementary Tables S2–S3).

All staphylococcal species were divided into human- and animal-associated species based on their most commonly reported hosts,^1, 41–44^ where *S. aureus*, a known colonizer of both humans and animals, was included in the human group. The resulting maps contain >280 QS interactions of native AIPs, representing the largest resource of its kind (Figure 2). The likelihood of interactions between certain human and animal-associated species to occur might be low; however, these interactions still represent a promising source for the discovery of potent QS inhibitors. Most measured QS interactions were inhibitory (227 of 288, 79%), where a clear difference between AIPs from human- and animal-associated species could be observed (Figure 2). The QS interactions of human-associated AIPs with *S. aureus, S. epidermidis*, and *S. lugdunensis* only exhibited >95% inhibition at 2.5 nM AIP concentration for intra-species interferences between *S. aureus* specificity groups. Further, most of the combinations that produced no effect at AIP concentration of 1 μM were from the human-associated group of AIPs (37 of 45, 82%) (Figure 2a and 2c). In contrast, most QS interactions of animal-associated AIPs (74 of 128, 58%) reached >95% inhibition at 2.5 nM or 50 nM AIP concentration (Figure 2b and 2c). The seven AIPs from species primarily associated with bovine colonization (*S. hyicus, S. chromogenes*, and *S. simulans*), displayed potent interactions, responsible for more than half of the examples of AIPs exhibiting >95% inhibition at 2.5 nM concentration (Figure 2c).

All AIPs that increased the fluorescence readout compared to the control wells were monitored continuously overnight for growth and fluorescence together with all previously known activators (Figure 2a and 2b, shown in green). Often an early increase in fluorescence output accompanied by a delay in growth was observed (Supplementary Figures S25,26). Interestingly, we found *S. epidermidis* AIP-III (**7**) to be an activator of *S. epidermidis agr*-II, in contrast to previous experiments with bacterial supernatant where the AIP had no reported effect.^45^ Further, several cross-species activators were discovered: *S. hominis* AIP-I (**10**) activated *S. epidermidis agr*-I, *S. hominis* AIP-III (**12**) activated *S. aureus agr*-IV, and *S. simulans* AIP-I (**33**) and AIP-III (**35**) activated *S. epidermidis agr*-III. We observed inconsistent inhibition behavior of several AIPs (**21**–**23, 25** and **31**) against some reporter strains (Figure 2b, shown in yellow). This behavior manifested itself either by causing activation at 1 μM and inhibition at lower concentrations or by showing 70% inhibition at multiple concentrations from 2.5–1000 nM, which was also recently observed for analogs of *S. epidermidis* AIPs.^35^

The QS interaction map identified peptides that were inhibitory across all staphylococcal reporter strains, namely *S. hyicus* AIP-I (**27**) and *S. chromogenes* AIP-I (**28**), which potently inhibited all *S. aureus* and *S. epidermidis agr* variants but only weakly inhibited *S. lugdunensis agr*-I as well as *S. simulans* AIP-II (**34**), which inhibited all eight *agr* systems (Figure 2b). Interestingly, all AIPs that acted as strong inhibitors of *S. aureus, S. epidermidis*, and *S. lugdunensis* (**27, 28, 30**–**35**) were 9-mer peptides, differing in length from the cognate AIPs of the reporter species (7, 8, and 12 residues), and they were the only AIPs with positively charged residues at the N-terminus (Supplementary Table S1). We found the AIPs of *S. simulans* interesting, because they displayed strong QS interaction profiles, including the most potent inhibition of *S. simulans* and the activation of *S. epidermidis agr*-III by *S. simulans* AIPs I (**33**) and III (**35**) but not by *S. simulans* AIP-II (**34**). We therefore peformed a structure–activity relationship study to glean further insights about the function of these molecules.

### Structure–activity relationship study of autoinducing peptides of *S. simulans*

*S. simulans* is primarily an animal-associated staphylococcal species, commonly found in bovine livestock,^46^ although human infections have also been documented, particularly involving *agr*-I type strains.^13^ The species has three confirmed AIPs (**33**–**35**), which share structural features.^11, 13^ *S. simulans* AIPs I (**33**) and III (**35**) share an identical exo-tail sequence, KYNP, which is also part of the exo-tail of *S. epidermidis* AIP-III (**7**) and could therefore explain the activating properties towards *S. epidermidis agr*-III (Figure 3a). Further, *S. simulans* AIPs II (**34**) and III (**35**) share an identical macrocycle and are both highly potent inhibitors of *S. lugdunensis* QS, compared to the *S. simulans* AIP-I (**33**) (Figure 3a). As starting point for our structure–activity relationship study, we determined the IC_50_ values of *S. simulans* AIP-I–III (**33**–**35**) giving sub- or low nanomolar potencies against *S. aureus agr*-I–III groups as well as *S. epidermidis agr*-I–II (Supplementary Figures S27–29). For *S. aureus agr*-IV, the IC_50_ value was slightly lower for **33** (10 nM) compared to **34** (32 nM) and **35** (34 nM) and against *S. lugdunensis agr*-I, **34** and **35** displayed sub- or low nanomolar potencies with the IC_50_ value of **35** (0.48 nM) being 400-fold lower compared to the most potent, previously reported inhibitors of *S. lugdunensis*.^37^ *S. simulans* AIP-II (**34**) displayed sub-nanomolar inhibition against *S. epidermidis agr*-III, while **33** and **35** acted as activators.

**Figure 3.**
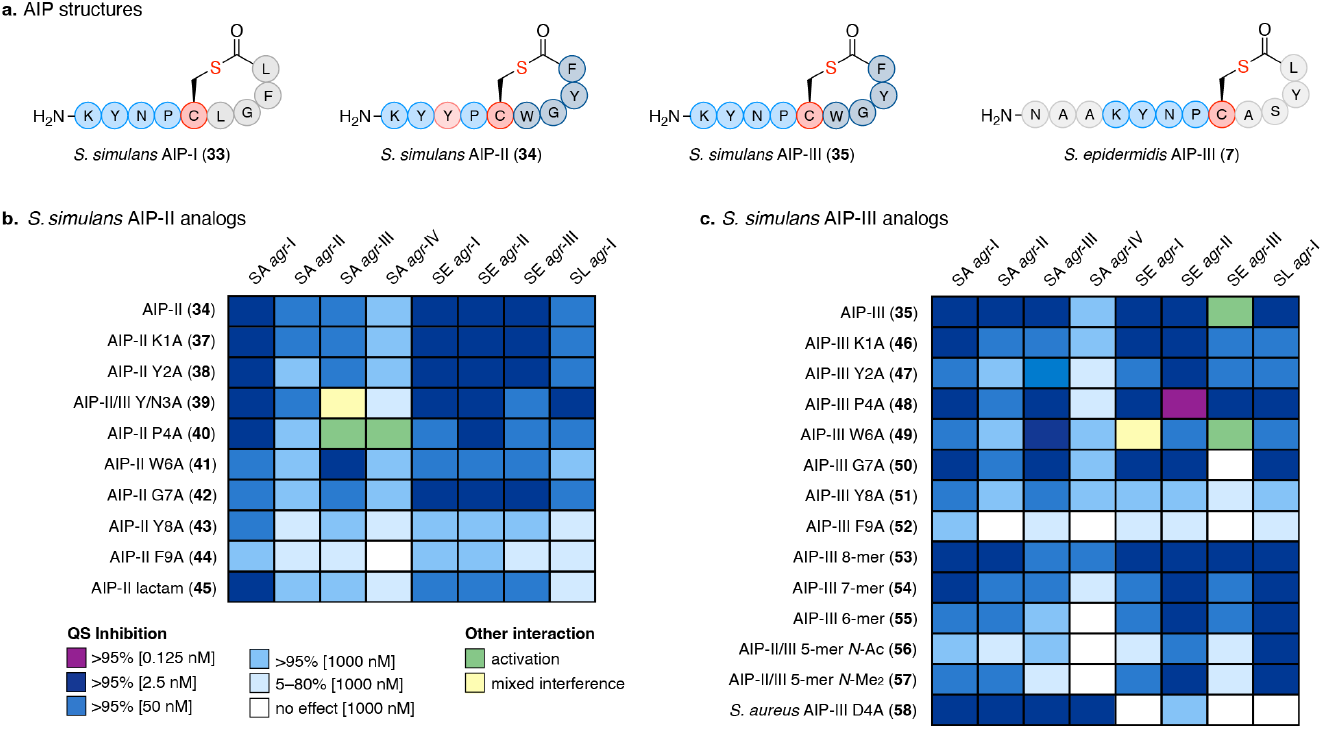
Structure–activity relationship study of *S. simulans* AIP-II (34) and AIP-III (35). **a**) Structures of *S. simulans* AIP-I–III (**33–35**) compared to SE-AIP-III (**7**). **b**) Alanine scan of *S. simulans* AIP-II (**34**). **c**) Alanine and truncation scan of *S. simulans* AIP-III (**35**). Synthetic peptides were tested at several concentration (1000 nM, 50 nM, 2.5 nM, 0.125 nM) against fluorescent reporter strains of *S. aureus* (SA) *agr*-I–IV, *S. epidermidis* (SE) *agr*-I–III, and *S. lugdunensis* (SL) *agr*-I. Shading of boxes represents different potencies of QS inhibition (>95% at 0.125 nM, 2.5 nM, 50 nM, and 1000 nM or 5–80% at 1000 nM), white boxes represent no interaction at 1000 nM, green boxes represent QS activation, and yellow boxes represent “mixed interference” (activation at 1000 nM and inhibition at 50 nM or partial inhibition over a range of concentrations).

Based on these results, we conducted alanine scans on both *S. simulans* AIPs II (**34**) and III (**35**), and tested the peptides to identify important residues for the activation of *S. epidermidis agr*-III and the inhibition of *S. lugdunensis* (Figure 3b and 3c, Supplementary Figures S30–34). The alanine (Ala) mutation of *S. simulans* AIP-II (**34**) starting from the N-terminus, K1A-II (**37**) and Y2A-II (**38**), showed minor effects to the QS interaction profile (Figure 3b). The Y3A-II mutant (**39**) also represents the N3A-III mutation of *S. simulans* AIP-III (**35**) as these peptides differ only in this position, and this common mutant acted as an inhibitor of *S. epidermidis agr*-III. Interestingly, the mutant P4A-II (**40**) became an activator of *S. aureus agr*-III and *agr*-IV, which we confirmed in a continuous assay (Supplementary Figure S25). Changes to the macrocycle in W6A-II (**41**) resulted in decreased inhibitory potency, except against *S. aureus agr*-III, while reducing structural flexibility by substiting glycine in G7A-II (**42**) furnished a decrease against all *S. aureus* strains. In agreement with the previous consensus,^47^ mutations to the C-terminal hydrophobic residues F8A-II (**43**) and L9A-II (**44**) led to significant loss in potency against all reporter strains. A thioester-to-amide analogue of **34** (**45**) (Supplementary Scheme S4), resulted in a decrease in potency against all species apart from *S. aureus agr*-I (Figure 3b). For the Ala-scan of *S. simulans* AIP-III (**35**), the two N-terminal mutants K1A-III (**46**) and Y2A-III (**47**) both lost the ability to activate *S. epidermidis agr*-III and showed reduced overall inhibitory potencies (Figure 3c). The proline mutation P4A-III (**48**) led to an increase in inhibition of *S. epidermidis* and *S. lugdunensis* and was the only tested peptide resulting in >95% inhibition at 0.125 nM AIP concentration. Interestingly, the mutation W6A-III (**49**) in the macrocycle had no effect on *S. epidermidis agr*-III activation, while otherwise leading to a weaker inhibition profile. Like G7A-II (**42**) also G7A-III (**50**) had less effect on inhibition but led to loss of activation of *S. epidermidis agr*-III. Finally, substitution of the two hydrophobic C-terminal residues in F8A-III (**51**) and L9A-III (**52**) led to diminished potency of the peptides as also observed for *S. simulans* AIP-II above. Next, we performed a truncation scan of *S. simulans* AIP-III (**35**), revealing that the inhibition of *S. aureus* was generally reduced by each truncation going from octamer to pentamer length, with the N-terminus of the pentamer being either acetylated or bis-N-methylated,^48^ to circumvent spontaneous rearrangement to the corresponding homodetic pentamer^49^ (**53**–**57**) (Figure 3c). The only exception was an increased inhibition of *S. aureus agr*-IV by **53** and the truncations had minor effects on *S. epidermidis* until the 6-mer (**55**), except for the loss of *S. epidermidis agr*-III activation. As anticipated, the macrocycle represented the key feature for potent inhibition of *S. lugdunensis agr*-I as all truncations including a 5-mer with di-methylated N-terminus (**57**) remained highly potent. Finally, we included a known inhibitor of all *S. aureus agr* groups, *S. aureus* AIP-III D4A (**58**)^22^ and observed potent inhibition of *S. aureus* with the only observed >95% inhibition at 2.5 nM against *S. aureus agr*-IV but weak interference with *S. epidermidis* and *S. lugdunensis* (Figure 3c).

Having established a foundation to design future QS inhibitors based on *S. simulans* AIP, we were interested to examine the potential of such compounds as anti-virulence agents.

### Autoinducing peptide of *S. simulans* as anti-virulence agent

Anti-virulence treatments based on QS inhibition for *S. aureus*, in particular MRSA, have been postulated since the discovery of *agr* cross-inhibition and could become an important addition in fighting resistant infections as it may attenuate their severity.^50^ However, despite a single recent clinical study with a commensal *S. hominis* strain,^18^ QS-based anti-virulence strategies require further development^51^ and certain questions need to be answered: firstly can *S. aureus* develop resistance towards QS inhibition; and secondly, how effective are QS inhibitors against already virulent bacteria. We attempted to address whether bacteria would develop resistance towards QS inhibitors, by assessing the effects of prolonged treatment of a fluorescent *S. aureus agr*-I P3-YFP reporter strain with *S. simulans* AIP-II (**34**) (Figure 4a). The bacteria were passaged daily for 15 days with and without addition of compound **34** and the *agr* activity was measured daily by flow cytometry, showing full repression of *agr* activity at 2 nM dosing of compound **34** over the full period (Figure 4b). After the 15 passages, all cultures were passaged once without addition of compound **34**, followed by assessment of the sensitivity of the strains towards inhibition of QS by compound **34**. Thus, all three cultures were exposed to a dilution series of the inhibitory AIP and no change in the potency was observed (Figure 4c). Despite the simplicity of this experiment, the results represent a first indicator that repeated treatments with AIP-based anti-virulence agents do not cause rapid resistance development.

**Figure 4.**
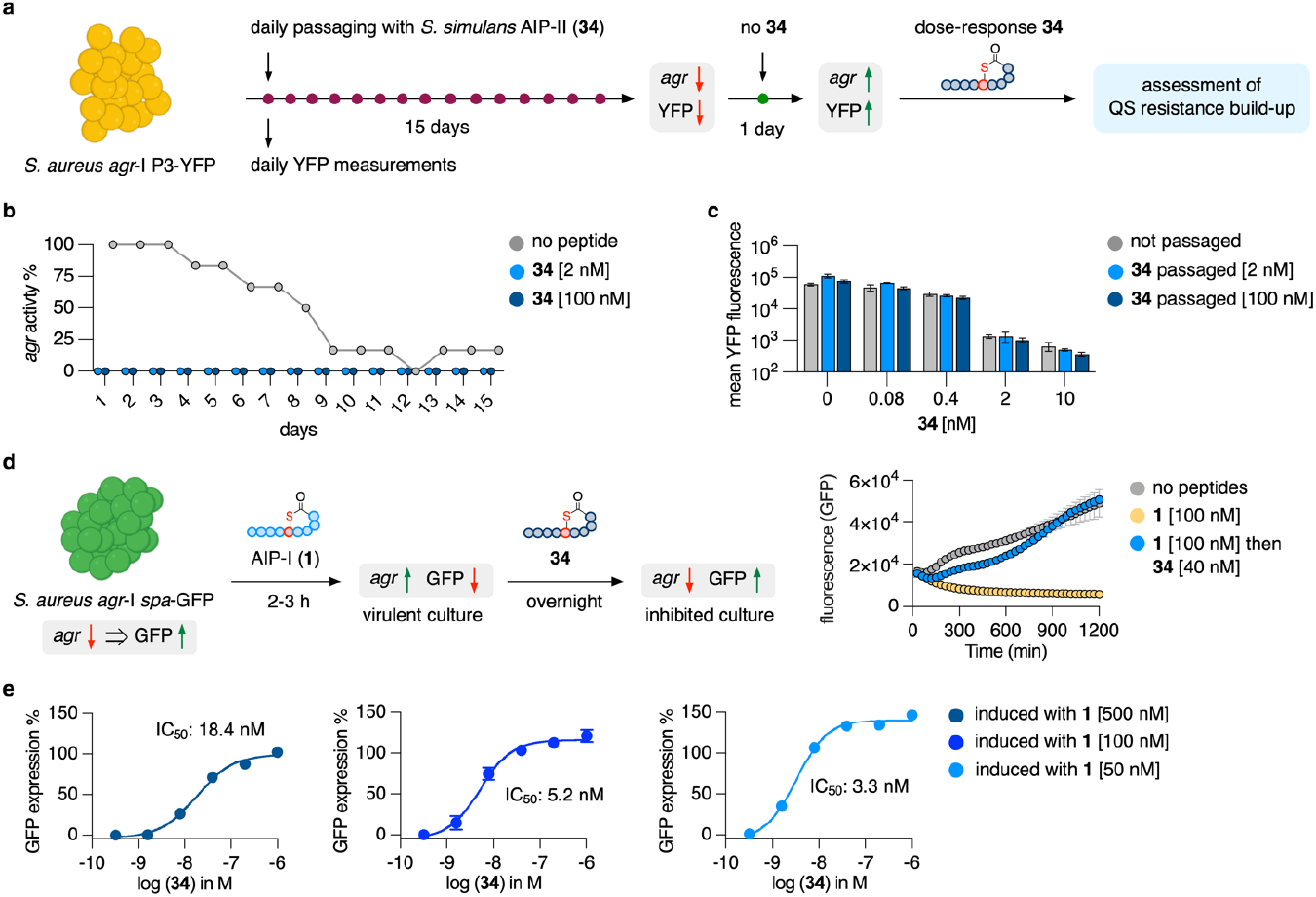
QS resistance development and *agr* deactivation assays. **a**) QS resistance development in a *S. aureus agr*-I P3-YFP reporter strain was examined by passaging cultures daily in the presence of *S. simulans* AIP-II (**34**) for 15 days. On day 16, the cultures were passaged without **34** and dose-response curves against **34** were measured to assess changes in its inhibitory potency. Fluorescence measurements were performed by flow cytometry. **b**) Activity of *agr* over 15 days measured by flow cytometry. The *agr* activity of untreated cultures decreases during treatment while daily treatments with **34** repressed *agr* activity at 2 nM and 100 nM. **c**) Passaged cultures were treated with **34** (10–0.08 nM) and remained equally susceptible to QS inhibition by **34** as not passaged cultures. **d**) The *S. aureus agr*-I *spa*-GFP reporter strain (JE2, MRSA) was treated with *S. aureus* AIP-I (**1**) during early growth to activate *agr* and repress GFP expression. At the start of exponential growth, **34** was added and GFP expression was monitored continuously. Treatment with **34** resulted in immediate increase of fluorescence while induced cultures with **1** remained non-fluorescent. **e**) IC_50_ values were determined for **34** against *S. aureus agr*-I *spa*-GFP induced with different concentrations of **1**. GFP expression was plotted relative to not induced cultures.

The majority of QS inhibition data in the literature is performed by treating cultures with inhibitor before the *agr* system was activated thereby measuring the prevention of *agr* activation. However, treatments would more often require *agr* deactivation of virulent bacteria. To better mimic a potential treatment scenario, we therefore transformed JE2, a highly virulent MRSA USA300 isolate of the *agr*-I type, with a *spa*-GFP reporter plasmid^52^ to create a reporter strain that enables the measurement of *agr* deactivation. The *spa* gene is down regulated when *agr* is activated and will therefore not become fluorescent when the QS system is active (Figure 4d). We induced the *agr* system of reporter cultures with cognate AIP **1** at 100 nM upon inoculation in fresh medium, and once early exponential phase was reached, the inhibitor **34** (40 nM) was added and GFP expression was monitored. Cultures induced with **1**, remained non-fluorescent over the time of the assay, in contrast to cultures treated with **34**, which rapidly started to express GFP and reached fluorescence levels like un-induced cultures because of *agr* deactivation (Figure 4d, Supplementary Figure S35). Next, we induced the *spa*-GFP reporter strain with different concentrations of **1** followed by serial dilutions of **34**, affording IC_50_ values for deactivation of *agr* in the low nanomolar range, which increased when challenged by induction with higher concentrations of **1** (Figure 4e). In comparison, the IC_50_ value for prevention of *agr* activation of *S. aureus agr*-I by compound **34**, measured in the P3-YFP reporter assay (0.45 nM), is 40-fold lower than the highest measured IC_50_ value for *agr* deactivation (18.4 nM at induction with 0.5 μM of **1**). The combined observations of these two *in vitro* experiments highlight that QS-based anti-virulence agents are unlikely to induce resistance and can turn off a fully activated *agr* system in MRSA.

Finally, we assessed the potential of *S. simulans* AIP-II (**34**) as an anti-virulence agent in an *in vivo* MRSA (*agr*-I) mouse skin infection model (Figure 5a–d, Supplementary Table S5). The importance of a functioning *agr* system for *S. aureus* during infection to evade the immune response has been established and it was shown that inhibition of *agr* during early stages of infection can lead to improved disease outcome 48–72 h after its initiation.^50^ Thus, *S. simulans* AIP-II (**34**) (100 μM) was added to the MRSA inoculum (10^7^ CFU) that was applied to the skin and compared to vehicle and daily treatment with the commercial antibacterial product Fucidin^®^ (2% fusidic acid). A significant reduction in the skin lesion size was observed after 48 h and 96 h for mice treated with **34** (*P* = 0.0174 and *P* = 0.0249) as well as fusidic acid (*P* = 0.0108 and *P* = 0.0202) compared to the vehicle control (Figure 5a,c). Further, a significant decrease in bacterial load (~60-fold, *P* = 0.009) was observed for mice treated with **34** compared to vehicle control after 4 days, which was comparable to daily treatment with fusidic acid (~38-fold, *P* = 0.0168) (Figure 5b).

**Figure 5.**
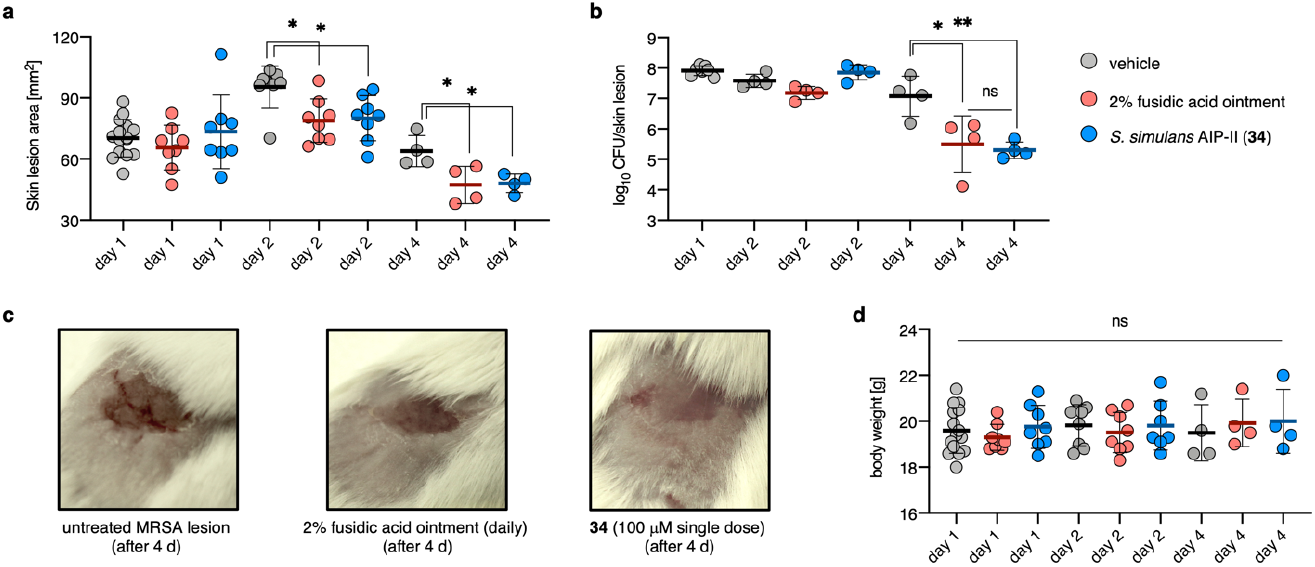
*S. simulans* AIP-II (34) attenuates MRSA infection in a murine skin model. **a**) Murine skin infection model performed with vehicle control group [day 1 (n = 16), day 2 (n = 8), day 4 (n = 4)], fusidic acid (daily application of a 38.7 mM ointment (Fucidin) [day 2 (n = 8) and 4 (n = 4)] and **34** (single treatment at day 0 at 100 μM [day 2 (n = 8) and 4 (n = 4)]. MRSA inoculum: 10^7^ CFU. Skin lesions measured on day 1, 2, and 4 showing significant reduction in lesion size for mice treated with fusidic acid and **34. b**) CFU count determined after day 1, 2, and 4 showing significant reduction in bacterial load per skin lesion. **c**) Representative pictures taken of MRSA lesions at day 4 for untreated and treated mice. **d**) Body weight measured at day 1, 2, and 4 showing no statistically significant differences. Data is presented as mean and error bars represent the standard deviation (SD) of the mean. *P* > 0.05 (ns), *P* < 0.05 (*), *P* < 0.01 (**).

No statistical difference between treatment or vehicle or time of the infection was found for the body weight, which was expected as no systemic infections were observed (Figure 5d). These results are encouraging for the prospects of anti-virulence treatments of staphylococcal infections with non-antibiotic peptides, such as **34**. The high extent of bacterial clearance is most likely a result of a more efficient immune response towards non-virulent MRSA bacteria.

## DISCUSSION

Altering gene expression of co-inhabiting staphylococci through secreted AIPs represents an intriguing ability of the *agr* system. The *agr* loci can be found in the genomes of all staphylococcal species and 38 AIPs from 22 species have now been identified from bacterial supernatants, pointing towards a broad utilization of the *agr* system, including potential QS interference. The colonization of humans and animals by a wide range of staphylococcal species emphasizes that many species share the same environments, providing an arena for biologically relevant inter-staphylococcal interactions, such as QS interference. Here, we report the most comprehensive mapping of QS interference by staphylococcal AIPs performed to date, including seven newly identified AIPs together with all previously reported AIPs. Testing this collection of AIPs against fluorescent reporter strains of *S. aureus agr*-I–IV, *S. epidermidis agr*-I–III, and *S. lugdunensis agr*-I provided a map of >280 QS interactions, which revealed a largely cross-inhibitory network, and lead to the discovery of several potent inhibitors against all the eight tested *agr* systems as well as previously unknown cross-species activators. To further scrutinize the requirements for the potent inhibition and activation profiles of the *S. simulans* AIPs, a structure–activity relationship study was performed based on *S. simulans* AIP-II (**34**) and *S. simulans* AIP-III (**35**), including the evaluation of alanine mutants and truncated peptides. The results of this exercise highlighted the importance of different structural features of the peptides. Truncations of the exo-tail affected the ability to inhibit *S. aureus* the most, while the macrocycle alone was enough to effectively inhibit *S. lugdunensis*. The *S. simulans* AIPs represent the first potent inhibitors of QS in *S. lugdunensis* with a 400-fold increase in potency compared to previously reported inhibitors.^37^ Furthermore, *S. simulans* AIP-II (**34**) was investigated for its potential as an anti-virulence agent, by assessing whether MRSA would develop resistance towards the inhibitor upon dosing over two weeks and assessing whether *agr* could be deactivated in a pathogenic MRSA strain. No detectable resistance development was observed after 15-day long treatment with **34** and we could demonstrate that **34** could shut down the fully active *agr* system of an MRSA isolate. Finally, a significant effect of this inhibitory AIP on the colonization and pathogenesis of MRSA *in viv*o was demonstrated in a mouse model, highlighting the power of gene repression through QS inhibition.^10,12,13,18^

Our results highlight the potential importance of the *agr* system and cross-species interference on the colonization of commensal staphylococci and on the pathogenesis of for example *S. aureus*. Though, the impact of the substantial QS interference among commensal staphylococci on the human microbiota remains to be explored further.

It is our hope that the mapping of cross-species QS interactions initiated in the present work will help provide insights into the roles of *agr* systems in future investigations. Furthermore, our findings highlight the potential utility of natural scaffolds as a promising platform for the development of inhibitors for anti-virulence treatment of *Staphylococcus* infections.

## ASSOCIATED CONTENT

The Supporting Information contains supplementary figures illustrating AIP trapping experiments, dose–response curves and bar graphs for tested compounds, supplementary schemes showing the compounds syntheses, and supplementary tables containing assay data and library compound sequences. Experimental procedures, materials, methods, and compound characterization data is provided as well as copies of HPLC chromatograms and copies ^1^H and ^13^C NMR spectra (PDF).

## AUTHOR INFORMATION

### Author contributions

CRediT: **Bengt H. Gless** conceptualization, formal analysis, investigation, methodology, data curation, supervision, visualization, writing-original draft, writing-review & editing; **Benjamin S. Sereika-Bejder** data curation, formal analysis, investigation, methodology, data curation, supervision, visualization, writing-review & editing; **Iben Jensen** formal analysis, investigation, methodology; **Martin S. Bojer** investigation, methodology, supervision, writing-review & editing; **Katerina Tsiko** investigation, methodology; **Sabrina H. Schmied** investigation; **Ludovica Vitolo** investigation; **Bruno Toledo-Silva** investigation; **Sarne De Vliegher** resources, supervision, writing-review & editing; **Hanne Ingmer** resources, funding acquisition, supervision, writing-review & editing; **Christian A. Olsen** conceptualization, resources, funding acquisition, project administration, supervision, visualization, writing-original draft, writing-review & editing.

### Notes

The authors declare no competing interest.

## ACKNOWLEDGMENTS

We thank Prof. Alexander Horswill (Univerity of Colorado) for donation of fluorescent reporter strains. We thank Carina Vingsbo Lundberg and Karen Juhl from Statens Serum Institut (DK) for performing the mouse studies under contract. We thank Peter Damborg (University of Copenhagen) and Paal S. Andersen (Statens Serum Institut (DK)) for contributing bacterial isolates. This work was supported by the Danish Independent Research Council–Natural Sciences (Grant No. 0135-00427B; C.A.O.), the LEO Foundation Open Competition Grant program (LF-OC-19-000039 and LF-OC-21-000901; CAO), and the Novo Nordisk Foundation–Interdisciplinary Synergy Programme (Grant No. 0077593; H.I.).

## REFERENCES

1. Otto, M. Staphylococci in the human microbiome: the role of host and interbacterial interactions. Current Opinion in Microbiology 2020, 53, 71–77.

2. Williams, P.; Hill, P.; Bonev, B.; Chan, W. C. Quorum-sensing, intra-and inter-species competition in the staphylococci. Microbiology 2023, 169.

3. Parlet, C. P.; Brown, M. M.; Horswill, A. R. Commensal Staphylococci Influence Staphylococcus aureus Skin Colonization and Disease. Trends in Microbiology 2019, 27, 497–507.

4. Ji, G.; Beavis, R. C.; Novick, R. P. Cell density control of staphylococcal virulence mediated by an octapeptide pheromone. Proceedings of the National Academy of Sciences 1995, 92, 12055–12059.

5. Thoendel, M.; Kavanaugh, J. S.; Flack, C. E.; Horswill, A. R. Peptide signaling in the Staphylococci. Chemical Reviews 2011, 111, 117–151.

6. Wang, B.; Muir, T. W. Regulation of virulence in Staphylococcus aureus: molecular mechanisms and remaining puzzles. Cell Chem Biol 2016, 23, 214–224.

7. Ji, G.; Beavis, R.; Novick, R. P. Bacterial Interference Caused by Autoinducing Peptide Variants. Science 1997, 276, 2027–2030.

8. Otto, M.; Süßmuth, R.; Vuong, C.; Jung, G.; Götz, F. Inhibition of virulence factor expression in Staphylococcus aureus by the Staphylococcus epidermidis agr pheromone and derivatives. FEBS Letters 1999, 450, 257–262.

9. Canovas, J.; Baldry, M.; Bojer, M. S.; Andersen, P. S.; Gless, B. H.; Grzeskowiak, P. K.; Stegger, M.; Damborg, P.; Olsen, C. A.; Ingmer, H. Cross-talk between Staphylococcus aureus and other staphylococcal species via the agr quorum sensing system. Frontiers in Microbiology 2016, 7, 1733.

10. Paharik, A. E.; Parlet, C. P.; Chung, N.; Todd, D. A.; Rodriguez, E. I.; Van Dyke, M. J.; Cech, N. B.; Horswill, A. R. Coagulase-negative staphylococcal strain prevents Staphylococcus aureus colonization and skin infection by blocking quorum sensing. Cell Host Microbe 2017, 22, 1–11.

11. Gless, B. H.; Bojer, M. S.; Peng, P.; Baldry, M.; Ingmer, H.; Olsen, C. A. Identification of autoinducing thiodepsipeptides from staphylococci enabled by native chemical ligation. Nature Chemistry 2019, 11, 463–469.

12. Williams, M. R.; Costa, S. K.; Zaramela, L. S.; Khalil, S.; Todd, D. A.; Winter, H. L.; Sanford, J. A.; O’Neill, A. M.; Liggins, M. C.; Nakatsuji, T.; Cech, N. B.; Cheung, A. L.; Zengler, K.; Horswill, A. R.; Gallo, R. L. Quorum sensing between bacterial species on the skin protects against epidermal injury in atopic dermatitis. Science Translational Medicine 2019, 11, eaat8329.

13. Brown, M. M.; Kwiecinski, J. M.; Cruz, L. M.; Shahbandi, A.; Todd, D. A.; Cech, N. B.; Horswill, A. R. Novel Peptide from Commensal Staphylococcus simulans Blocks Methicillin-Resistant Staphylococcus aureus Quorum Sensing and Protects Host Skin from Damage. Antimicrobial Agents and Chemotherapy 2020, 64, e00172–00120.

14. Severn, M. M.; Cho, Y.-S. K.; Manzer, H. S.; Bunch, Z. L.; Shahbandi, A.; Todd, D. A.; Cech, N. B.; Horswill, A. R. The Commensal Staphylococcus warneri Makes Peptide Inhibitors of MRSA Quorum Sensing that Protect Skin from Atopic or Necrotic Damage. Journal of Investigative Dermatology 2022, 142, 3349–3352.e3345.

15. Severn Morgan, M.; Williams Michael, R.; Shahbandi, A.; Bunch Zoie, L.; Lyon Laurie, M.; Nguyen, A.; Zaramela Livia, S.; Todd Daniel, A.; Zengler, K.; Cech Nadja, B.; Gallo Richard, L.; Horswill Alexander, R. The Ubiquitous Human Skin Commensal Staphylococcus hominis Protects against Opportunistic Pathogens. mBio 2022, 13, e00930–00922.

16. Nakamura, Y.; Takahashi, H.; Takaya, A.; Inoue, Y.; Katayama, Y.; Kusuya, Y.; Shoji, T.; Takada, S.; Nakagawa, S.; Oguma, R.; Saito, N.; Ozawa, N.; Nakano, T.; Yamaide, F.; Dissanayake, E.; Suzuki, S.; Villaruz, A.; Varadarajan, S.; Matsumoto, M.; …; Shimojo, N. Staphylococcus Agr virulence is critical for epidermal colonization and associates with atopic dermatitis development. Science Translational Medicine 2020, 12, eaay4068.

17. Tamai, M.; Yamazaki, Y.; Ito, T.; Nakagawa, S.; Nakamura, Y. Pathogenic role of the staphylococcal accessory gene regulator quorum sensing system in atopic dermatitis. Frontiers in Cellular and Infection Microbiology 2023, 13.

18. Nakatsuji, T.; Hata, T. R.; Tong, Y.; Cheng, J. Y.; Shafiq, F.; Butcher, A. M.; Salem, S. S.; Brinton, S. L.; Rudman Spergel, A. K.; Johnson, K.; Jepson, B.; Calatroni, A.; David, G.; Ramirez-Gama, M.; Taylor, P.; Leung, D. Y. M.; Gallo, R. L. Development of a human skin commensal microbe for bacteriotherapy of atopic dermatitis and use in a phase 1 randomized clinical trial. Nature Medicine 2021, 27, 700–709.

19. Maura, D.; Ballok, A. E.; Rahme, L. G. Considerations and caveats in anti-virulence drug development. Curr Opin Microbiol 2016, 33, 41–46.

20. Dickey, S. W.; Cheung, G. Y. C.; Otto, M. Different drugs for bad bugs: antivirulence strategies in the age of antibiotic resistance. Nat Rev Drug Discov 2017, 16, 457–471.

21. Mayville, P.; Ji, G.; Beavis, R.; Yang, H.; Goger, M.; Novick, R. P.; Muir, T. W. Structure-activity analysis of synthetic autoinducing thiolactone peptides from Staphylococcus aureus responsible for virulence. Proceedings of the National Academy of Sciences 1999, 96, 1218–1223.

22. Tal-Gan, Y.; Stacy, D. M.; Foegen, M. K.; Koenig, D. W.; Blackwell, H. E. Highly potent inhibitors of quorum sensing in Staphylococcus aureus revealed through a systematic synthetic study of the group-III autoinducing peptide. J Am Chem Soc 2013, 135, 7869–7882.

23. Vasquez, J. K.; Blackwell, H. E. Simplified Autoinducing Peptide Mimetics with Single-Nanomolar Activity Against the Staphylococcus aureus AgrC Quorum Sensing Receptor. ACS Infectious Diseases 2019, 5, 484–492.

24. West, K. H. J.; Gahan, C. G.; Kierski, P. R.; Calderon, D. F.; Zhao, K.; Czuprynski, C. J.; McAnulty, J. F.; Lynn, D. M.; Blackwell, H. E. Sustained Release of a Synthetic Autoinducing Peptide Mimetic Blocks Bacterial Communication and Virulence In Vivo. Angewandte Chemie International Edition 2022, 61, e202201798.

25. Todd, D. A.; Parlet, C. P.; Crosby, H. A.; Malone, C. L.; Heilmann, K. P.; Horswill, A. R.; Cech, N. B. Signal biosynthesis inhibition with ambuic acid as a strategy to target antibiotic-resistant infections. Antimicrobial Agents and Chemotherapy 2017, 61, e00263–00217.

26. Otto, M.; Echner, H.; Voelter, W.; Götz, F. Pheromone Cross-Inhibition between Staphylococcus aureus and Staphylococcus epidermidis. Infection and Immunity 2001, 69, 1957–1960.

27. Yang, T.; Tal-Gan, Y.; Paharik, A. E.; Horswill, A. R.; Blackwell, H. E. Structure-function analyses of a Staphylococcus epidermidis autoinducing peptide reveals motifs critical for AgrC-type receptor modulation. ACS Chem Biol 2016, 11, 1982–1991.

28. Vasquez, J. K.; West, K. H. J.; Yang, T.; Polaske, T. J.; Cornilescu, G.; Tonelli, M.; Blackwell, H. E. Conformational Switch to a β-Turn in a Staphylococcal Quorum Sensing Signal Peptide Causes a Dramatic Increase in Potency. Journal of the American Chemical Society 2020, 142, 750–761.

29. Severn, M. M.; Horswill, A. R. Staphylococcus epidermidis and its dual lifestyle in skin health and infection. Nature Reviews Microbiology 2023, 21, 97–111.

30. Brown, M. M.; Horswill, A. R. Staphylococcus epidermidis—Skin friend or foe? PLOS Pathogens 2020, 16, e1009026.

31. Liu, Q.; Liu, Q.; Meng, H.; Lv, H.; Liu, Y.; Liu, J.; Wang, H.; He, L.; Qin, J.; Wang, Y.; Dai, Y.; Otto, M.; Li, M. Staphylococcus epidermidis Contributes to Healthy Maturation of the Nasal Microbiome by Stimulating Antimicrobial Peptide Production. Cell Host & Microbe 2020, 27, 68–78.e65.

32. Linehan, J. L.; Harrison, O. J.; Han, S.-J.; Byrd, A. L.; Vujkovic-Cvijin, I.; Villarino, A. V.; Sen, S. K.; Shaik, J.; Smelkinson, M.; Tamoutounour, S.; Collins, N.; Bouladoux, N.; Dzutsev, A.; Rosshart, S. P.; Arbuckle, J. H.; Wang, C.-R.; Kristie, T. M.; Rehermann, B.; Trinchieri, G.; …; Belkaid, Y. Non-classical Immunity Controls Microbiota Impact on Skin Immunity and Tissue Repair. Cell 2018, 172, 784–796.e718.

33. Otto, M. Staphylococcus epidermidis — the ‘accidental’ pathogen. Nature Reviews Microbiology 2009, 7, 555–567.

34. West, K. H. J.; Shen, W.; Eisenbraun, E. L.; Yang, T.; Vasquez, J. K.; Horswill, A. R.; Blackwell, H. E. Non-Native Peptides Capable of Pan-Activating the agr Quorum Sensing System across Multiple Specificity Groups of Staphylococcus epidermidis. ACS Chemical Biology 2021, 16, 1070–1078.

35. Eisenbraun, E. L.; Vulpis, T. D.; Prosser, B. N.; Horswill, A. R.; Blackwell, H. E. Synthetic Peptides Capable of Potent Multigroup Staphylococcal Quorum Sensing Activation and Inhibition in Both Cultures and Biofilm Communities. Journal of the American Chemical Society 2024, 146, 15941–15954.

36. Heilbronner, S.; Foster, T. J. Staphylococcus lugdunensis: a Skin Commensal with Invasive Pathogenic Potential. Clinical Microbiology Reviews 2021, 34, e00205–00220.

37. Gordon, C. P.; Olson, S. D.; Lister, J. L.; Kavanaugh, J. S.; Horswill, A. R. Truncated Autoinducing Peptides as Antagonists of Staphylococcus lugdunensis Quorum Sensing. J Med Chem 2016, 59, 8879–8888.

38. Dawson, P. E.; Muir, T. W.; Clark-Lewis, I.; Kent, S. B. H. Synthesis of proteins by native chemical ligation. Science 1994, 266, 776.

39. Lamers, R. P.; Muthukrishnan, G.; Castoe, T. A.; Tafur, S.; Cole, A. M.; Parkinson, C. L. Phylogenetic relationships among Staphylococcus species and refinement of cluster groups based on multilocus data. BMC Evolutionary Biology 2012, 12, 171.

40. Malone, C. L.; Boles, B. R.; Lauderdale, K. J.; Thoendel, M.; Kavanaugh, J. S.; Horswill, A. R. Fluorescent reporters for Staphylococcus aureus. Journal of Microbiological Methods 2009, 77, 251–260.

41. Nagase, N.; Sasaki, A.; Yamashita, K.; Shimizu, A.; Wakita, Y.; Kitai, S.; Kawano, J. Isolation and Species Distribution of Staphylococci from Animal and Human Skin. Journal of Veterinary Medical Science 2002, 64, 245–250.

42. Bagcigil, F. A.; Moodley, A.; Baptiste, K. E.; Jensen, V. F.; Guardabassi, L. Occurrence, species distribution, antimicrobial resistance and clonality of methicillin- and erythromycin-resistant staphylococci in the nasal cavity of domestic animals. Veterinary Microbiology 2007, 121, 307–315.

43. Pyörälä, S.; Taponen, S. Coagulase-negative staphylococci—emerging mastitis pathogens. Veterinary Microbiology 2009, 134, 3–8.

44. Becker, K.; Heilmann, C.; Peters, G. Coagulase-Negative Staphylococci. Clinical Microbiology Reviews 2014, 27, 870–926.

45. Olson, M. E.; Todd, D. A.; Schaeffer, C. R.; Paharik, A. E.; Van Dyke, M. J.; Büttner, H.; Dunman, P. M.; Rohde, H.; Cech, N. B.; Fey, P. D.; Horswill, A. R. Staphylococcus epidermidis agr quorum-sensing system: signal identification, cross talk, and importance in colonization. J Bacteriol 2014, 196, 3482–3493.

46. Rall, V. L. M.; Miranda, E. S.; Castilho, I. G.; Camargo, C. H.; Langoni, H.; Guimarães, F. F.; Araújo Júnior, J. P.; Fernandes Júnior, A. Diversity of Staphylococcus species and prevalence of enterotoxin genes isolated from milk of healthy cows and cows with subclinical mastitis. Journal of Dairy Science 2014, 97, 829–837.

47. Polaske, T. J.; West, K. H. J.; Zhao, K.; Widner, D. L.; York, J. T.; Blackwell, H. E. Chemical and Biomolecular Insights into the Staphylococcus aureus Agr Quorum Sensing System: Current Progress and Ongoing Challenges. Israel Journal of Chemistry 2023, 63, e202200096.

48. Bejder, B. S.; Monda, F.; Gless, B. H.; Bojer, M. S.; Ingmer, H.; Olsen, C. A. A short-lived peptide signal regulates cell-to-cell communication in Listeria monocytogenes. Commun Biol 2024, 7, 942.

49. Gless, B. H.; Bejder, B. S.; Monda, F.; Bojer, M. S.; Ingmer, H.; Olsen, C. A. Rearrangement of Thiodepsipeptides by S --> N Acyl Shift Delivers Homodetic Autoinducing Peptides. J Am Chem Soc 2021, 143, 10514–10518.

50. Wright, J. S. I.; Jin, R.; Novick, R. P. Transient interference with staphylococcal quorum sensing blocks abscess formation. Proc Natl Acad Sci U S A 2005, 102, 1691–1696.

51. Otto, M. Critical Assessment of the Prospects of Quorum-Quenching Therapy for Staphylococcus aureus Infection. In International Journal of Molecular Sciences, 2023; Vol. 24.

52. Kupferwasser, L. I.; Yeaman, M. R.; Nast, C. C.; Kupferwasser, D.; Xiong, Y.-Q.; Palma, M.; Cheung, A. L.; Bayer, A. S. Salicylic acid attenuates virulence in endovascular infections by targeting global regulatory pathways in Staphylococcus aureus. The Journal of Clinical Investigation 2003, 112, 222–233.

