## Supplementary information for "Mapping of Quorum Sensing Landscape of Commensal and Pathogenic Staphylococci Reveals a Largely Inhibitory Interaction Network"

### 1. Supplementary figures

#### The staphylococcal *accessory gene regulator* (*agr*) system

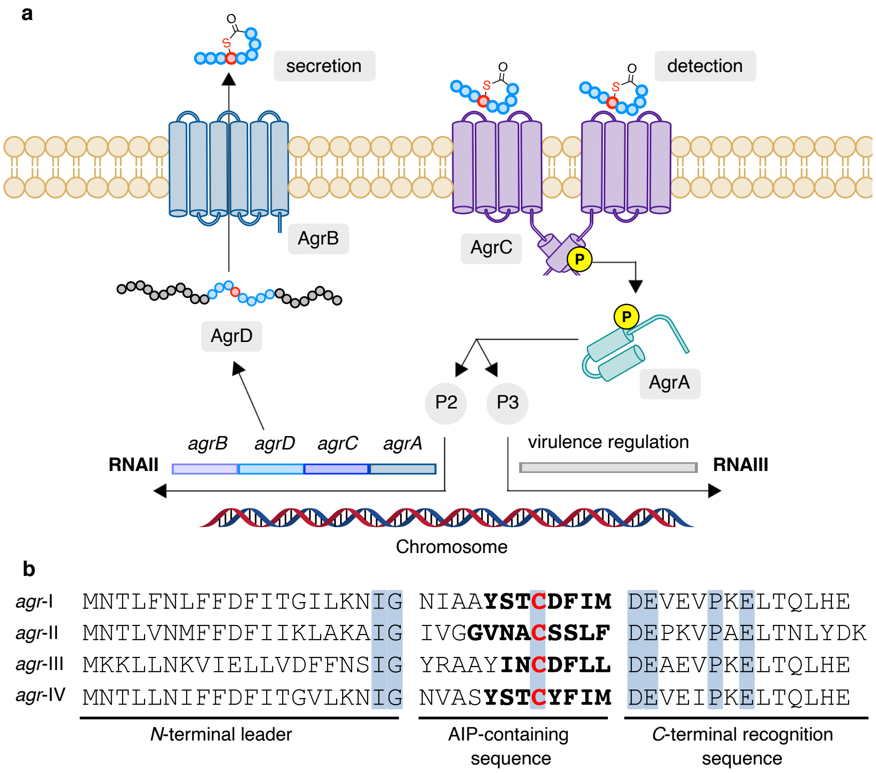

**Supplementary Figure S1. The staphylococcal *agr* system. a**, The chromosomal *agr* locus consists of the two promoter regions P2 and P3, which initiate the transcription of RNAII and RNAIII, respectively. RNAII encodes the four protein components (AgrA, AgrB, AgrC, and AgrD) of the *agr* system, while RNAIII is a regulatory RNA controlling the expression of virulence factors. In the first step of the *agr* circuit, the AIP-precursor peptide AgrD (44–46 amino acids for *S. aureus*) is processed by the membrane-embedded endopeptidase AgrB installing the thiolactone functionality. The peptide is further translocated to the extracellular space and finally cleaved to release the mature AIP. Once a concentration threshold of the AIP molecules is reached due to increased cell density, the AgrC receptor, a homodimeric membrane-bound histidine kinase, is activated. AIP-induced activation is followed by auto-phosphorylation of AgrC and subsequent phosphoryl transfer to AgrA, the response regulator of the *agr* system, making the AgrC–AgrA interaction a classical two-component regulatory system. Phosphorylated AgrA binds to the promotors P2 and P3, resulting in upregulated transcription of RNAII and RNAIII, which lead to a positive feedback loop for AgrBDCA expression as well as upregulated expression of virulence factors. **b**, AgrD peptides consist of three domains, the *C*-terminal recognition sequence, the *N*-terminal leader peptide and in between the 12 amino acid long AIP-containing sequence. Conserved residues are highlighted in blue boxes.

#### Native chemical ligation (NCL) trapping of autoinducing peptides (AIPs)

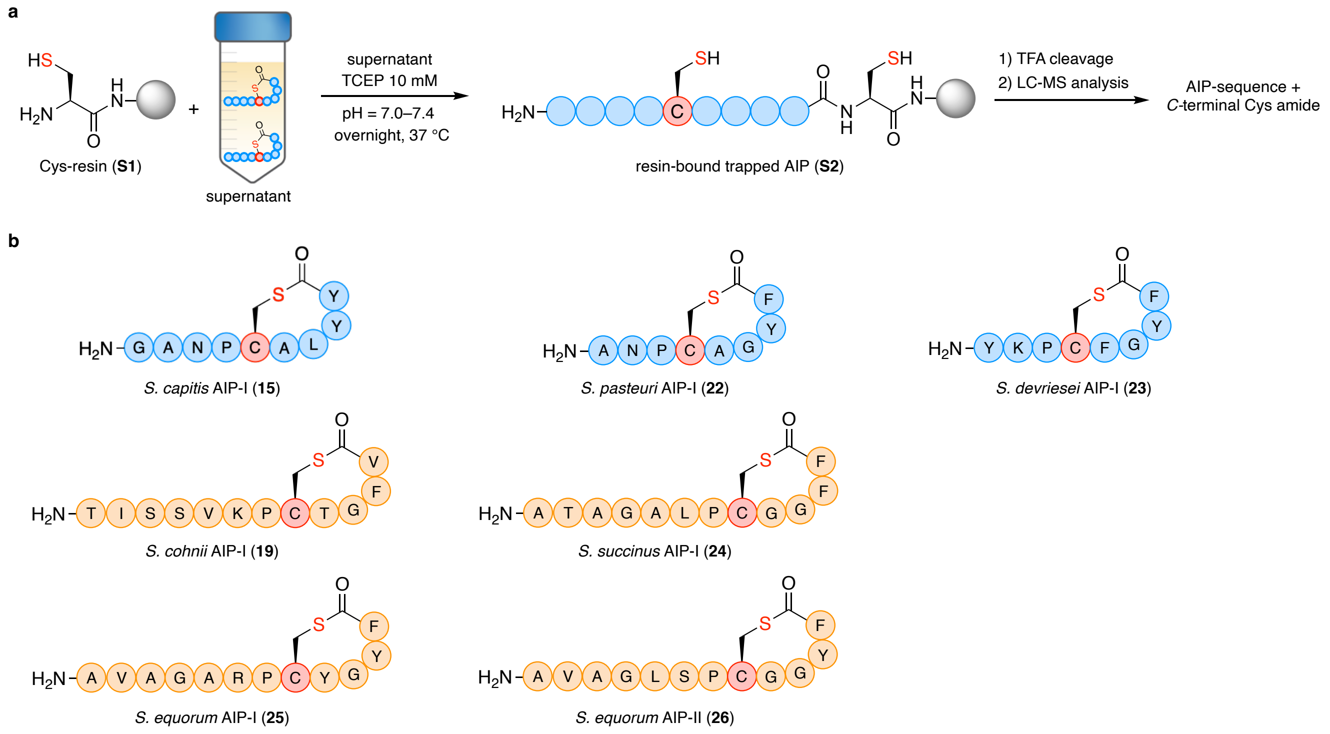

**Supplementary Figure S2. Identification of AIPs through NCL trapping. a**, Cys-resin (**S1**) is incubated overnight in pH-adjusted bacterial supernatant containing tris(2-carboxyethyl)phosphine (TCEP, 10 mM) enabling chemoselective trapping of AIPs. The resin with the trapped AIP (**S2**) is washed extensively and subsequently treated with trifluoroacetic acid (TFA) to release the trapped AIP. The concentrated cleavage solution is analyzed by liquid-chromatography mass-spectrometry (LC-MS) for the 7 possible AIPs sequences with an additional *C*-terminal Cys amide extracted from the AIP-precursor peptide AgrD. **b**, Newly indentified AIPs in this study.

#### LC-MS traces of sequence-guided identifications of AIPs

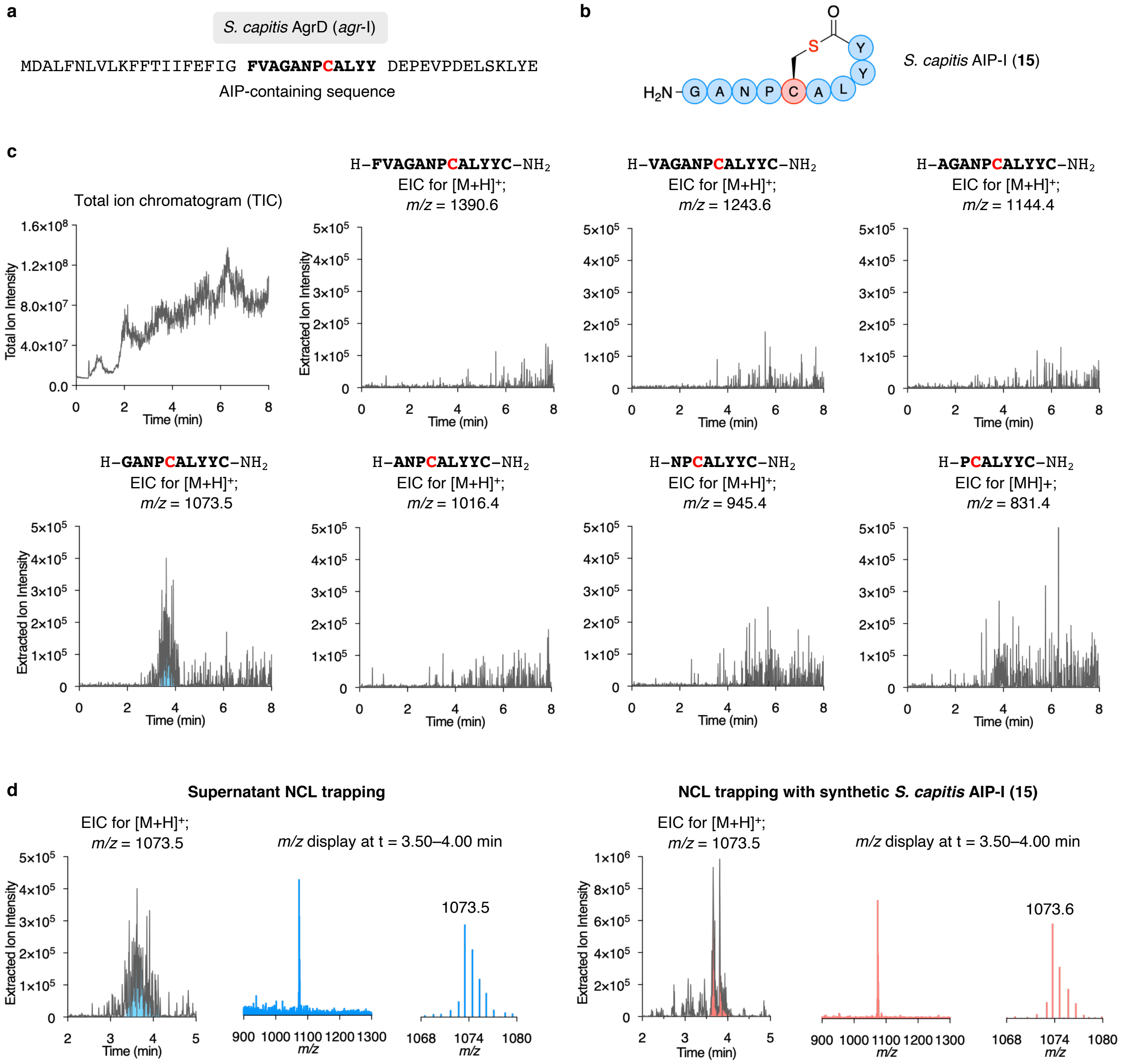

**Supplementary Figure S3. NCL trapping of *S. capitis* AIP-I (15). a**, AgrD sequence of *S. pasteuri agr*-I. **b**, Structure of *S. capitis* AIP-I (**15**). **c**, LC-MS analysis of the TFA cleavage solution: total ion chromatogram (TIC) and extracted ion chromatograms (EIC) of *m/z* = $[M+H]^+$ of possible linear AIP sequences including a C-terminal cysteine amide. **d**, NCL trapping of synthetic *S. capitis* AIP-I (**15**) confirms the identity of trapped AIP **15** from bacterial supernatant.

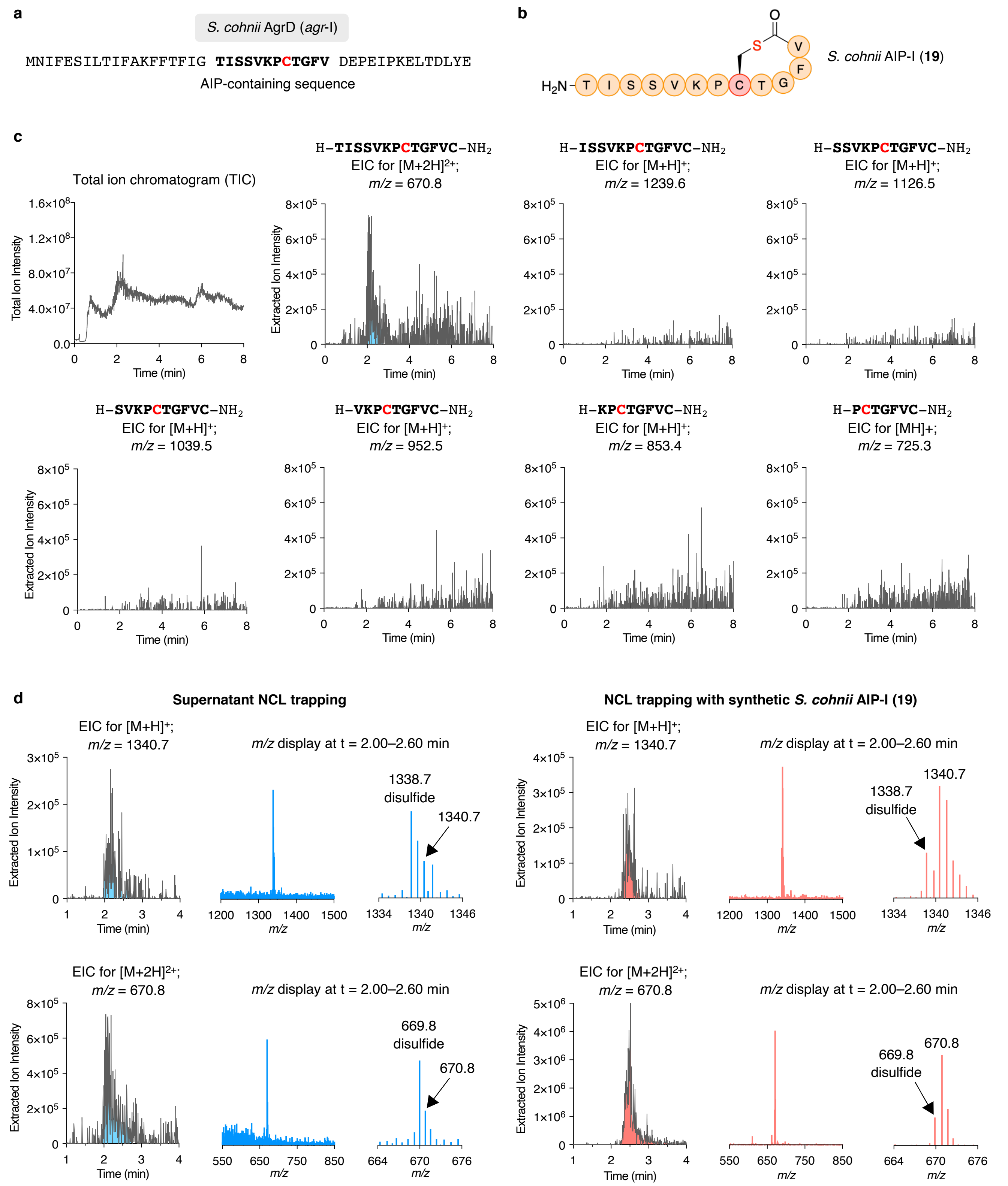

**Supplementary Figure S4. NCL trapping of *S. cohnii* AIP-I (19). a**, AgrD sequence of *S. cohnii agr*-I. **b**, Structure of *S. cohnii* AIP-I (**19**). **c**, LC-MS analysis of the TFA cleavage solution: TIC and EICs of *m/z* = $[M+H]^+$ of possible linear AIP sequences including a C-terminal cysteine amide. **d**, NCL trapping of synthetic *S. cohnii* AIP-I (**19**) confirms the identity of trapped AIP **19** from bacterial supernatant.

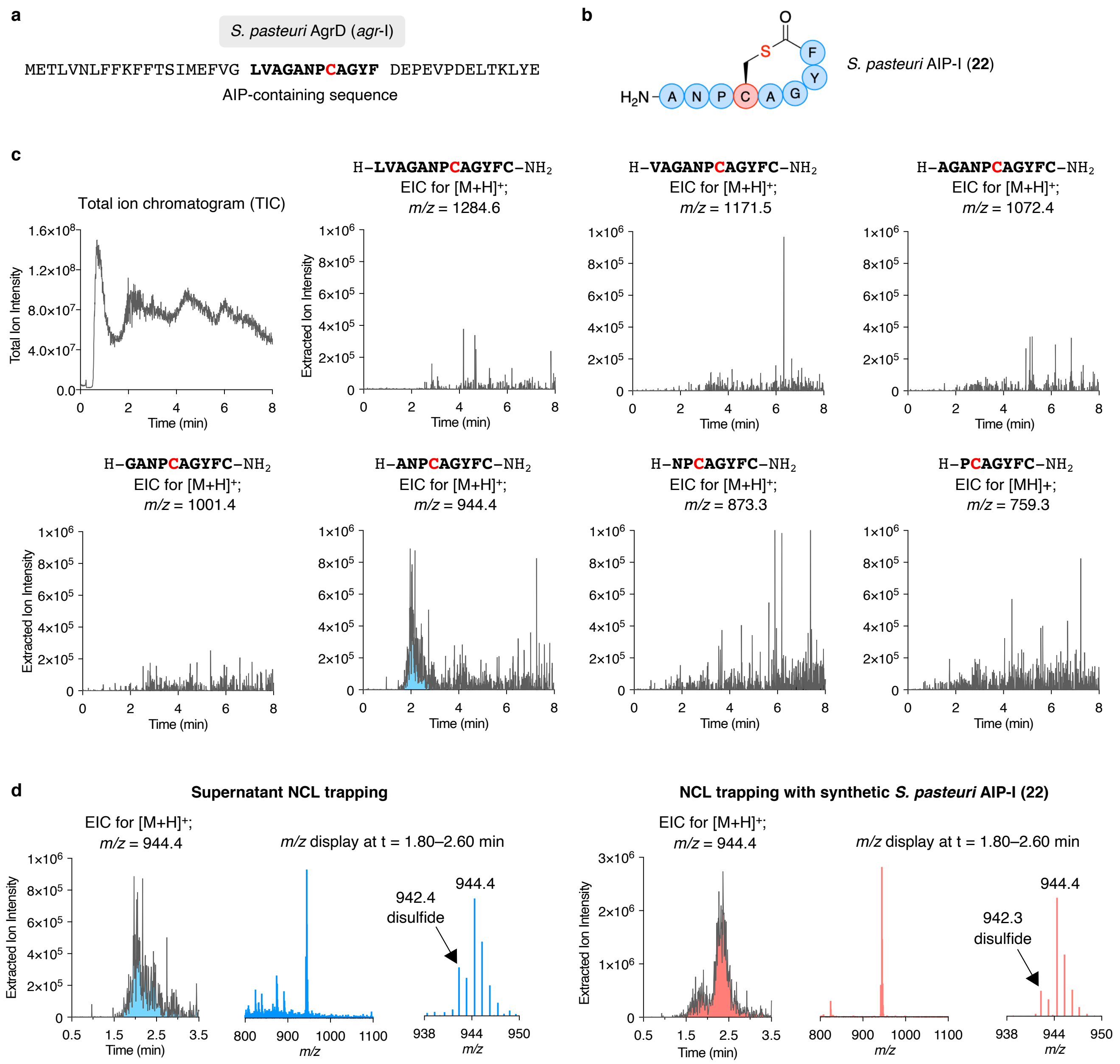

**Supplementary Figure S5. NCL trapping of *S. pasteuri* AIP-I (22).** **a**, AgrD sequence of *S. pasteuri agr*-I. **b**, Structure of *S. pasteuri* AIP-I (**22**). **c**, LC-MS analysis of the TFA cleavage solution: TIC and EICs of *m/z* = $[M+H]^+$ of possible linear AIP sequences including a C-terminal cysteine amide. **d**, NCL trapping of synthetic *S. pasteuri* AIP-I (**22**) confirms the identity of trapped AIP **22** from bacterial supernatant.

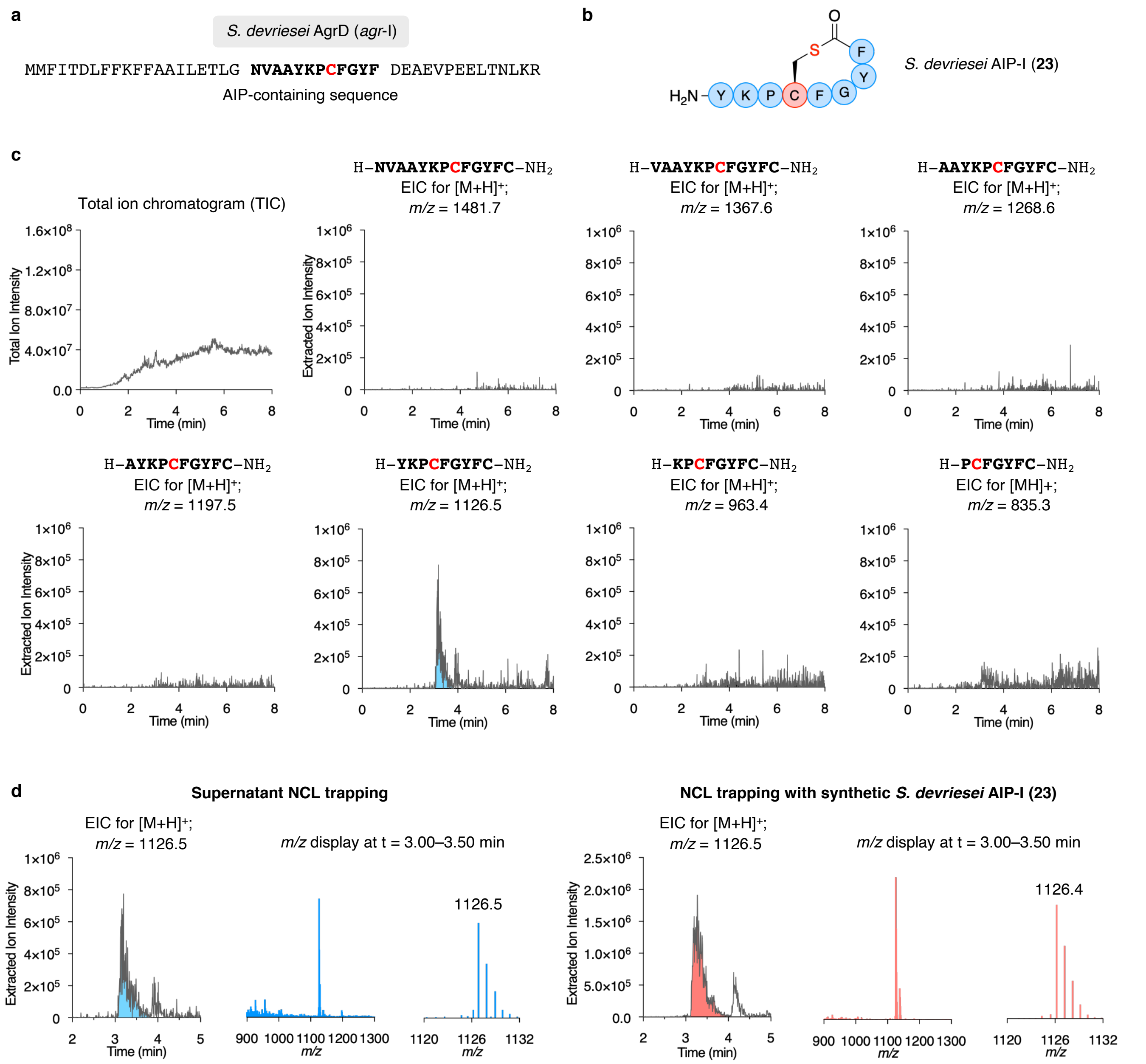

**Supplementary Figure S6. NCL trapping of *S. devriesei* AIP-I (23). a**, AgrD sequence of *S. devriesei agr*-I. **b**, Structure of *S. devriesei* AIP-I (**23**). **c**, LC-MS analysis of the TFA cleavage solution: TIC and EICs of *m/z* = $[M+H]^+$ of possible linear AIP sequences including a C-terminal cysteine amide. **d**, NCL trapping of synthetic *S. devriesei* AIP-I (**23**) confirms the identity of trapped AIP **23** from bacterial supernatant.

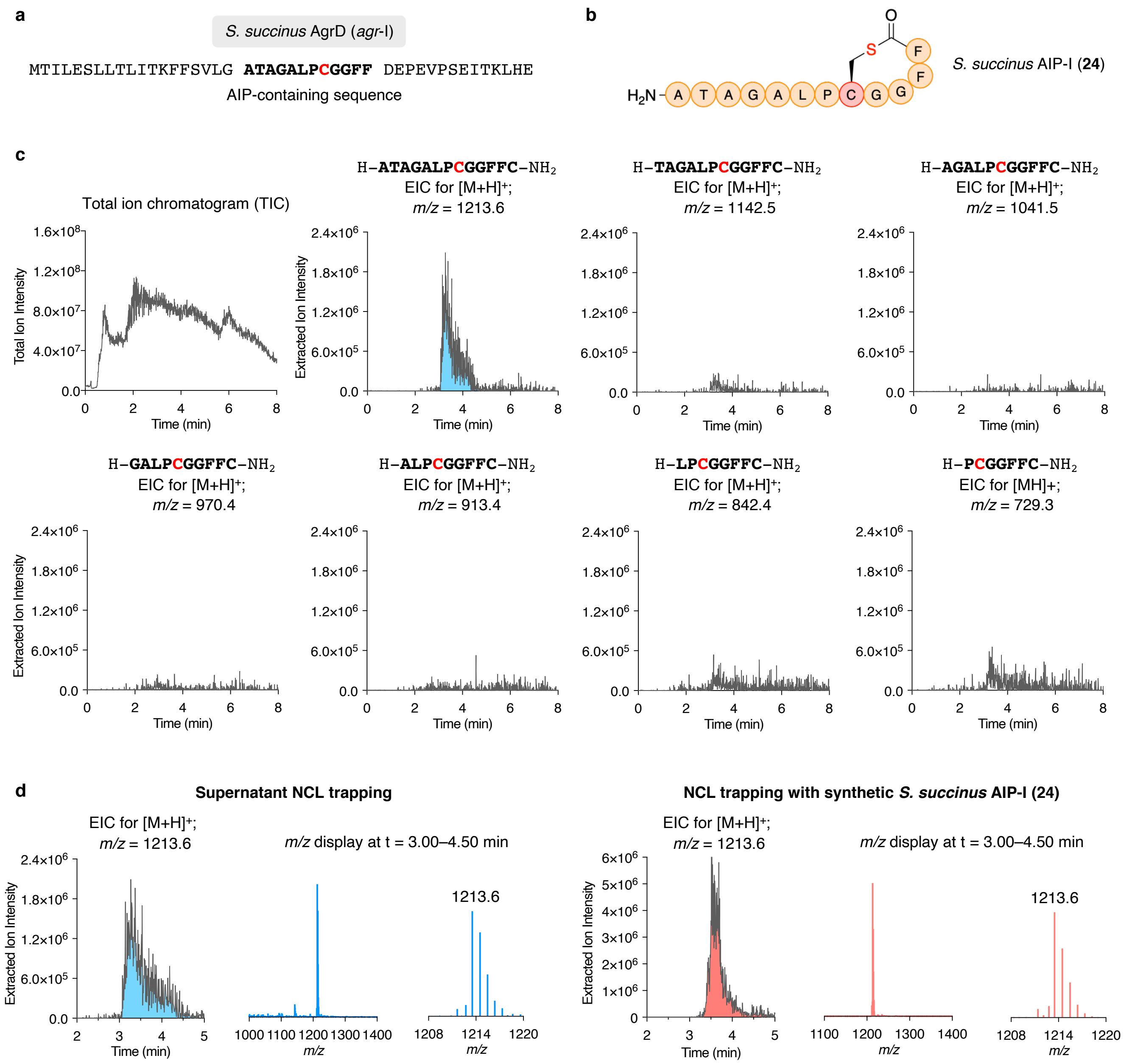

**Supplementary Figure S7. NCL trapping of *S. succinus* AIP-I (24).** **a**, AgrD sequence of *S. succinus agr*-I. **b**, Structure of *S. succinus* AIP-I (**24**). **c**, LC-MS analysis of the TFA cleavage solution: TIC and EICs of $m/z = [M+H]^+$ of possible linear AIP sequences including a C-terminal cysteine amide. **d**, NCL trapping of synthetic *S. succinus* AIP-I (**24**) confirms the identity of trapped AIP **24** from bacterial supernatant.

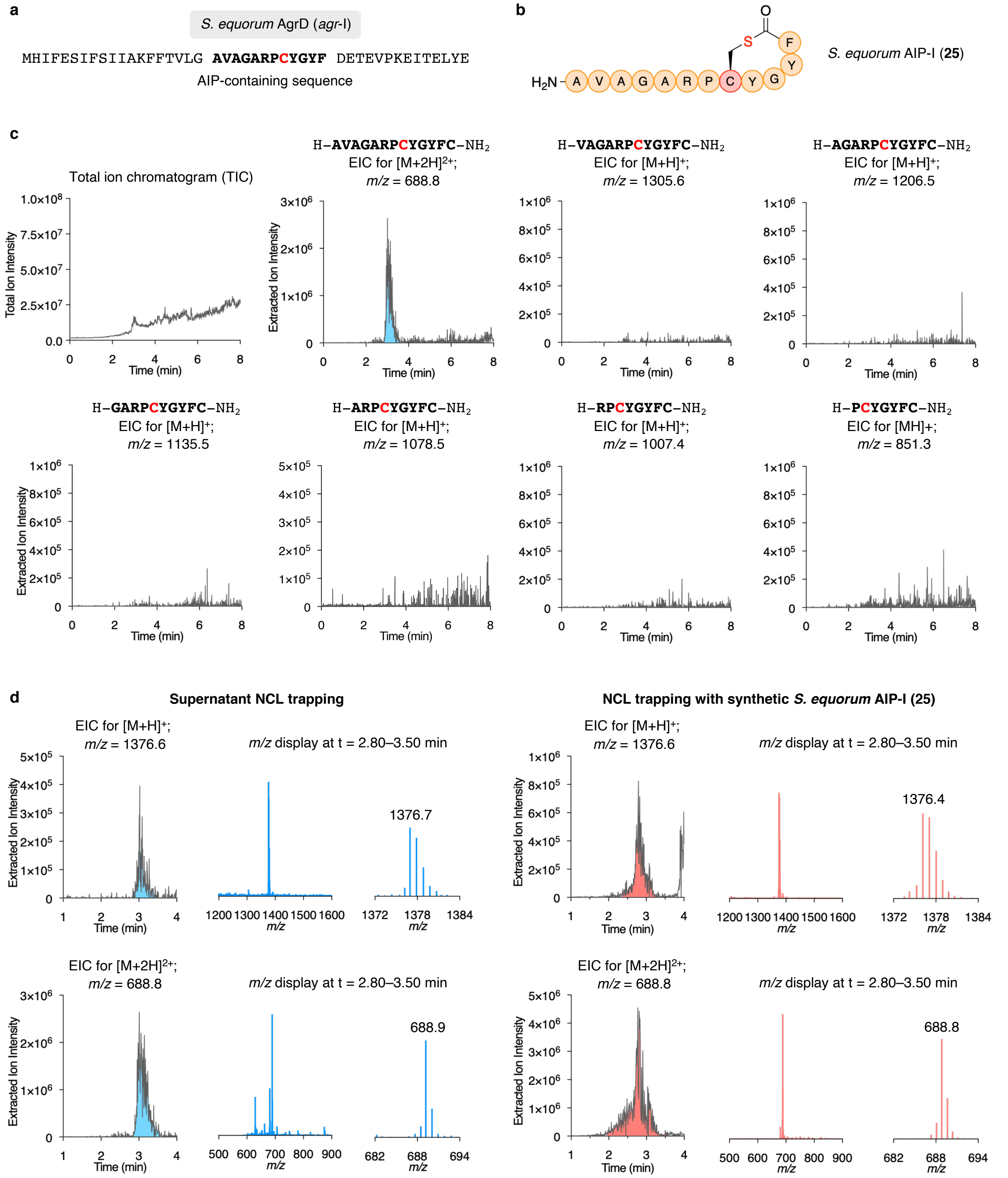

**Supplementary Figure S8. NCL trapping of *S. equorum* AIP-I (25). a**, AgrD sequence of *S. equorum agr*-I. **b**, Structure of *S. equorum* AIP-I (**25**). **c**, LC-MS analysis of the TFA cleavage solution: TIC and EICs of $m/z$ = $[M+H]^+$ of possible linear AIP sequences including a C-terminal cysteine amide. **d**, NCL trapping of synthetic *S. equorum* AIP-I (**25**) confirms the identity of trapped AIP **25** from bacterial supernatant.

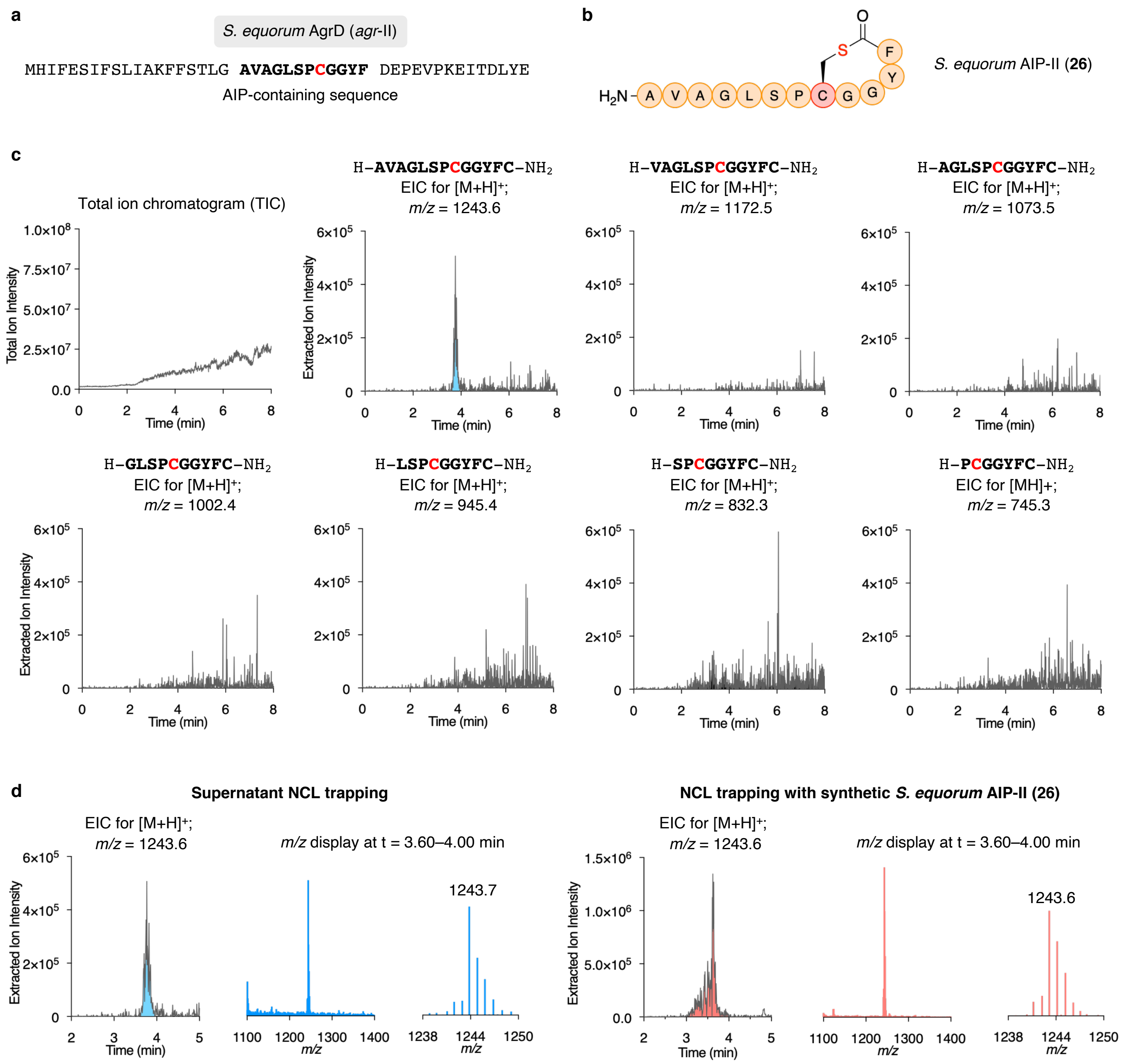

**Supplementary Figure S9. NCL trapping of *S. equorum* AIP-II (26). a**, AgrD sequence of *S. equorum agr*-II. **b**, Structure of *S. equorum* AIP-II (**26**). **c**, LC-MS analysis of the TFA cleavage solution: TIC and EICs of *m/z* = $[M+H]^+$ of possible linear AIP sequences including a C-terminal cysteine amide. **d**, NCL trapping of synthetic *S. equorum* AIP-II (**26**) confirms the identity of trapped AIP **26** from bacterial supernatant.

#### Bar graphs for *agr* interference using fluorescence reporter assays of native AIPs

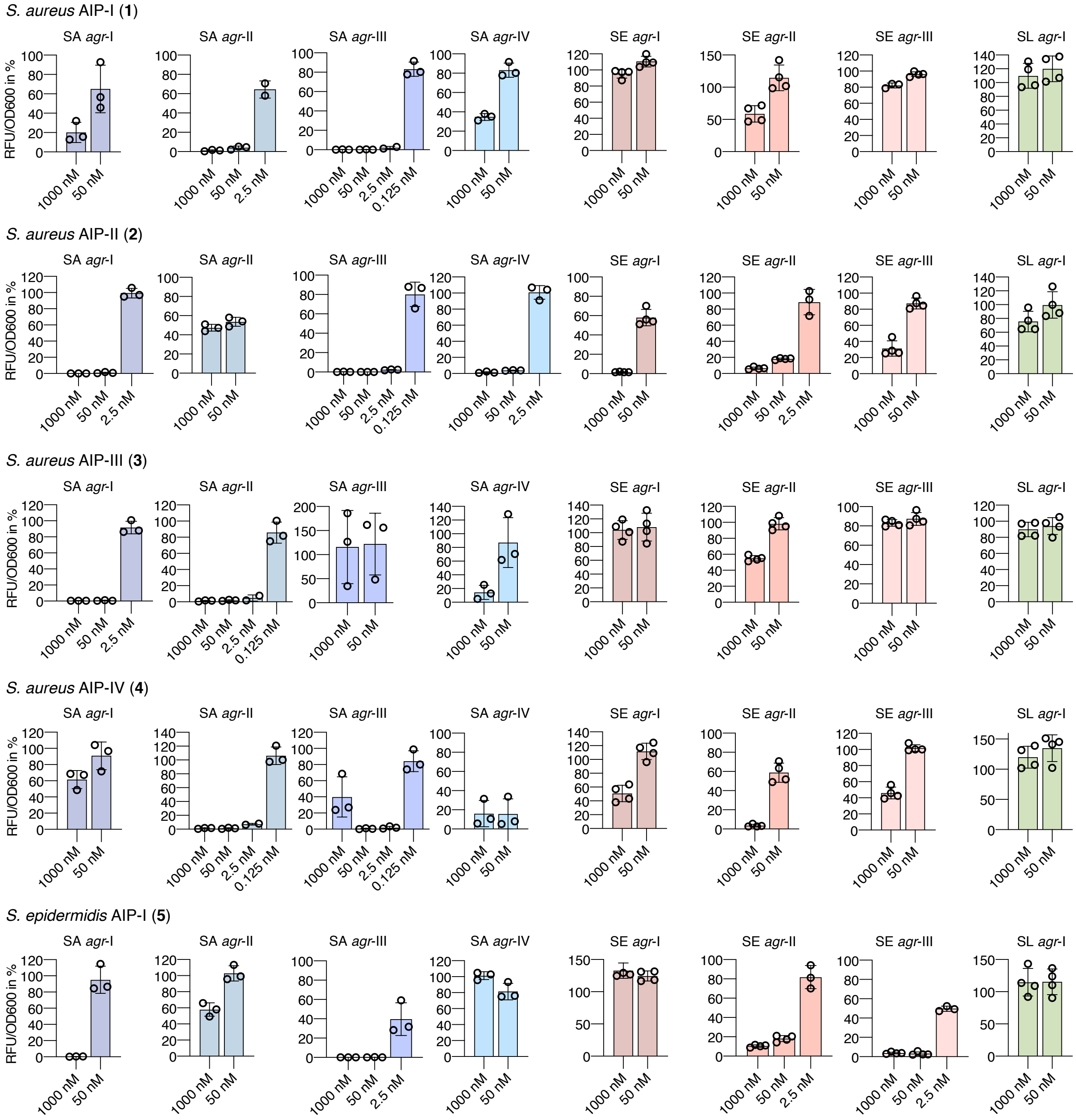

**Supplementary Figure S10. Fluorescence reporter strain assay for *agr* interference with native AIPs.** Fluorescent reporter strains of *S. aureus* (SA), *S. epidermidis* (SE), S. *lugdunensis* (SL) were treated with AIPs at 1000 nM and 50 nM. AIPs were further tested at 2.5 nM and 0.125 nM in case >75% inhibition was observed at higher concentrations. Error bars are the standard error of the mean (SEM) of at least two individual biological assays performed in technical triplicate.

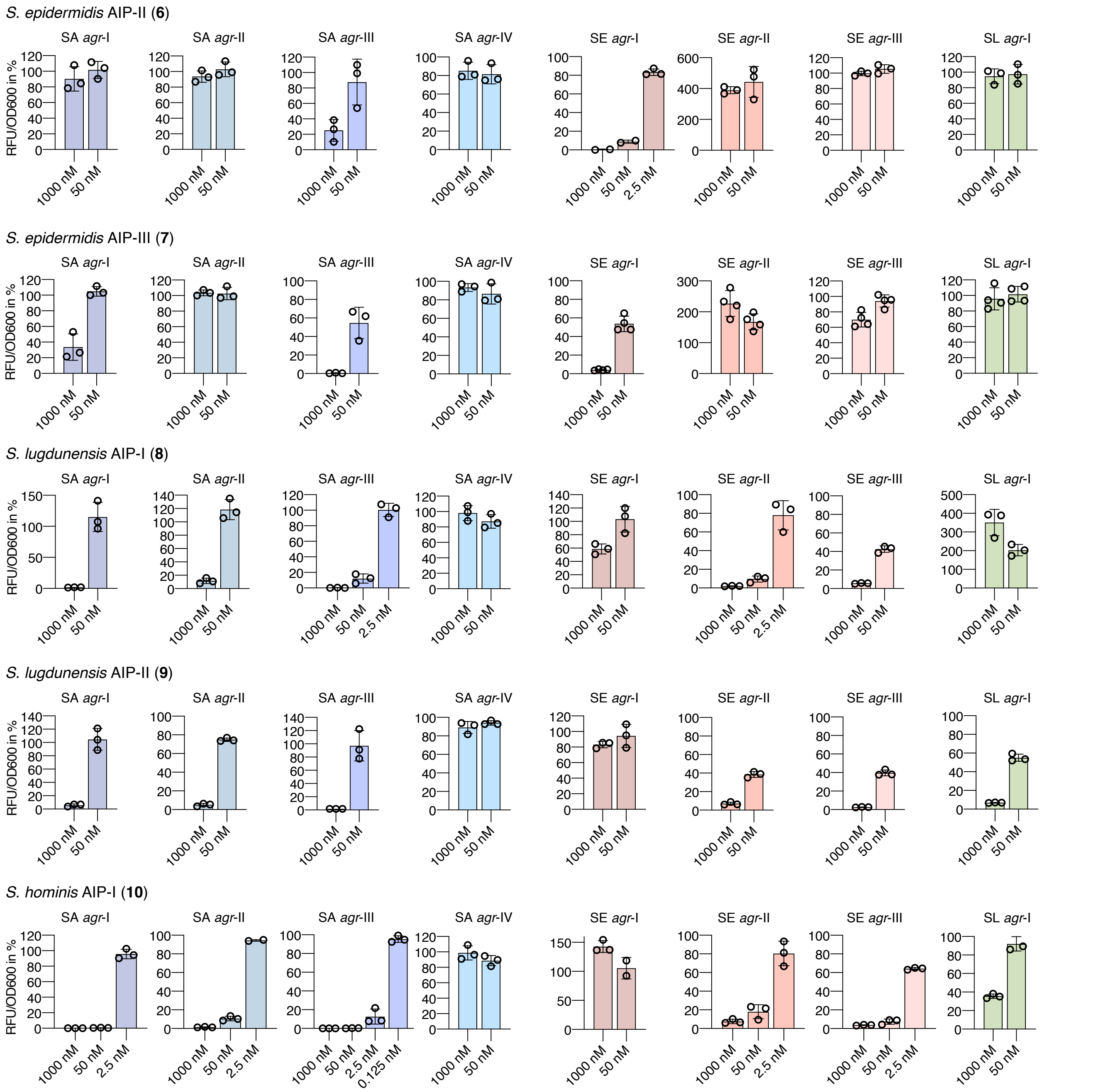

**Supplementary Figure S11. Fluorescence reporter strain assay for *agr* interference with with native AIPs.** Fluorescent reporter strains of of *S. aureus* (SA), *S. epidermidis* (SE), S. *lugdunensis* (SL) were treated with AIPs at 1000 nM and 50 nM. AIPs were further tested at 2.5 nM and 0.125 nM in case >75% inhibition was observed at higher concentrations. Error bars are the SEM of at least two individual biological assays performed in technical triplicate.

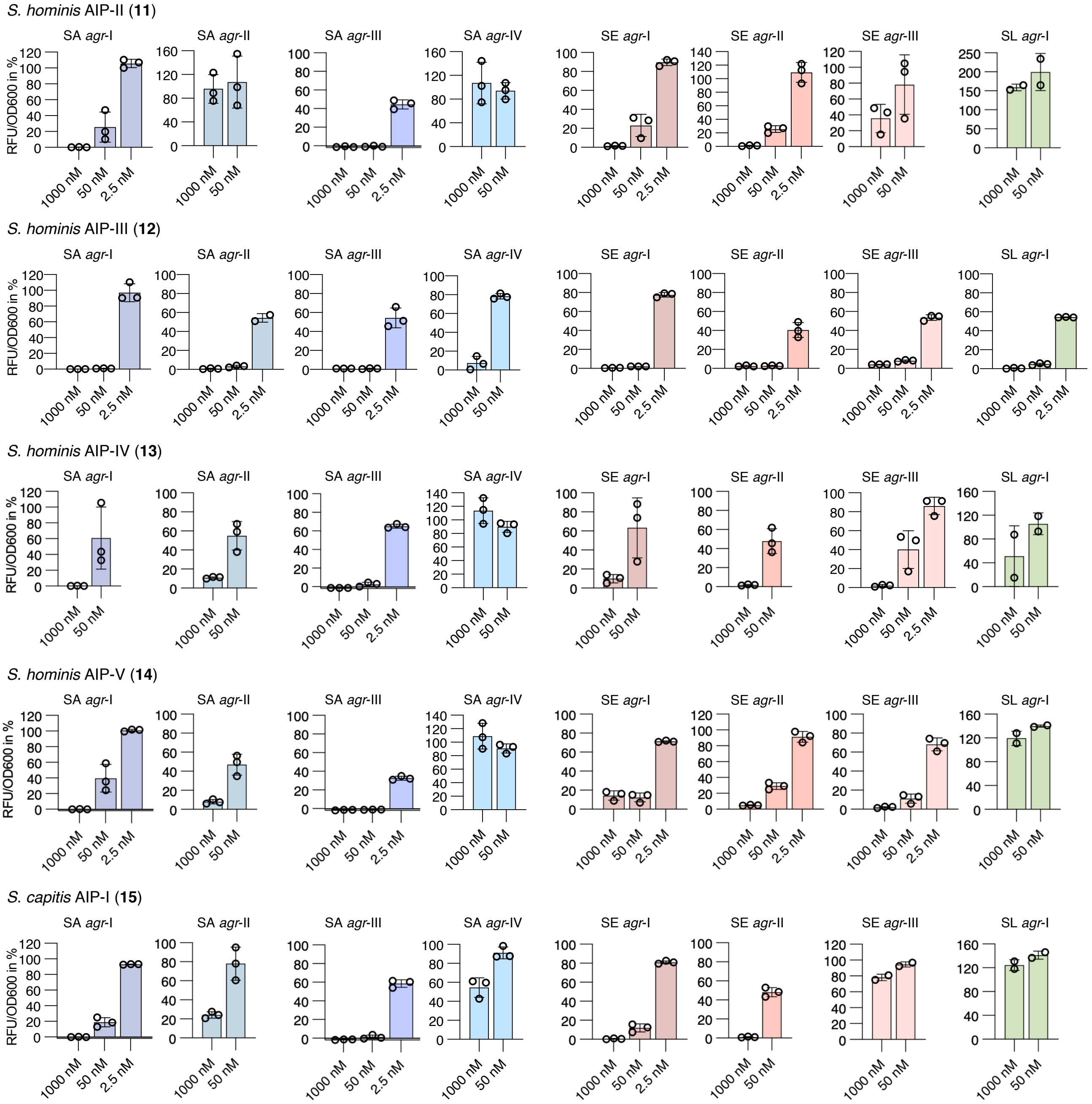

**Supplementary Figure S12. Fluorescence reporter strain assay for *agr* interference with native AIPs.** Fluorescent reporter strains of of *S. aureus* (SA), *S. epidermidis* (SE), S. *lugdunensis* (SL) were treated with AIPs at 1000 nM and 50 nM. AIPs were further tested at 2.5 nM and 0.125 nM in case >75% inhibition was observed at higher concentrations. Error bars are the SEM of at least two individual biological assays performed in technical triplicate.

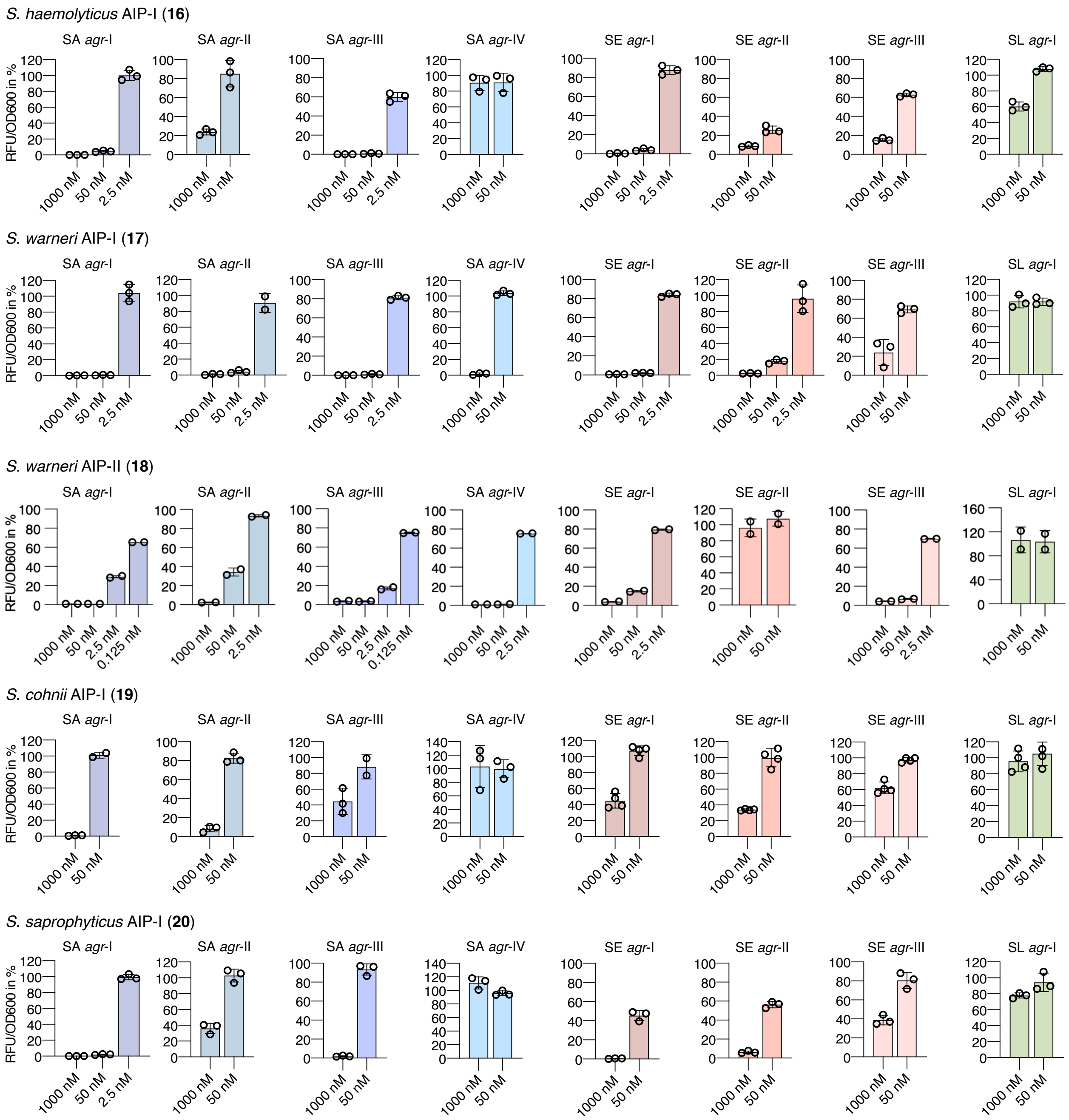

**Supplementary Figure S13. Fluorescence reporter strain assay for *agr* interference with native AIPs.** Fluorescent reporter strains of of *S. aureus* (SA), *S. epidermidis* (SE), S. *lugdunensis* (SL) were treated with AIPs at 1000 nM and 50 nM. AIPs were further tested at 2.5 nM and 0.125 nM in case >75% inhibition was observed at higher concentrations. Error bars are the SEM of at least two individual biological assays performed in technical triplicate.

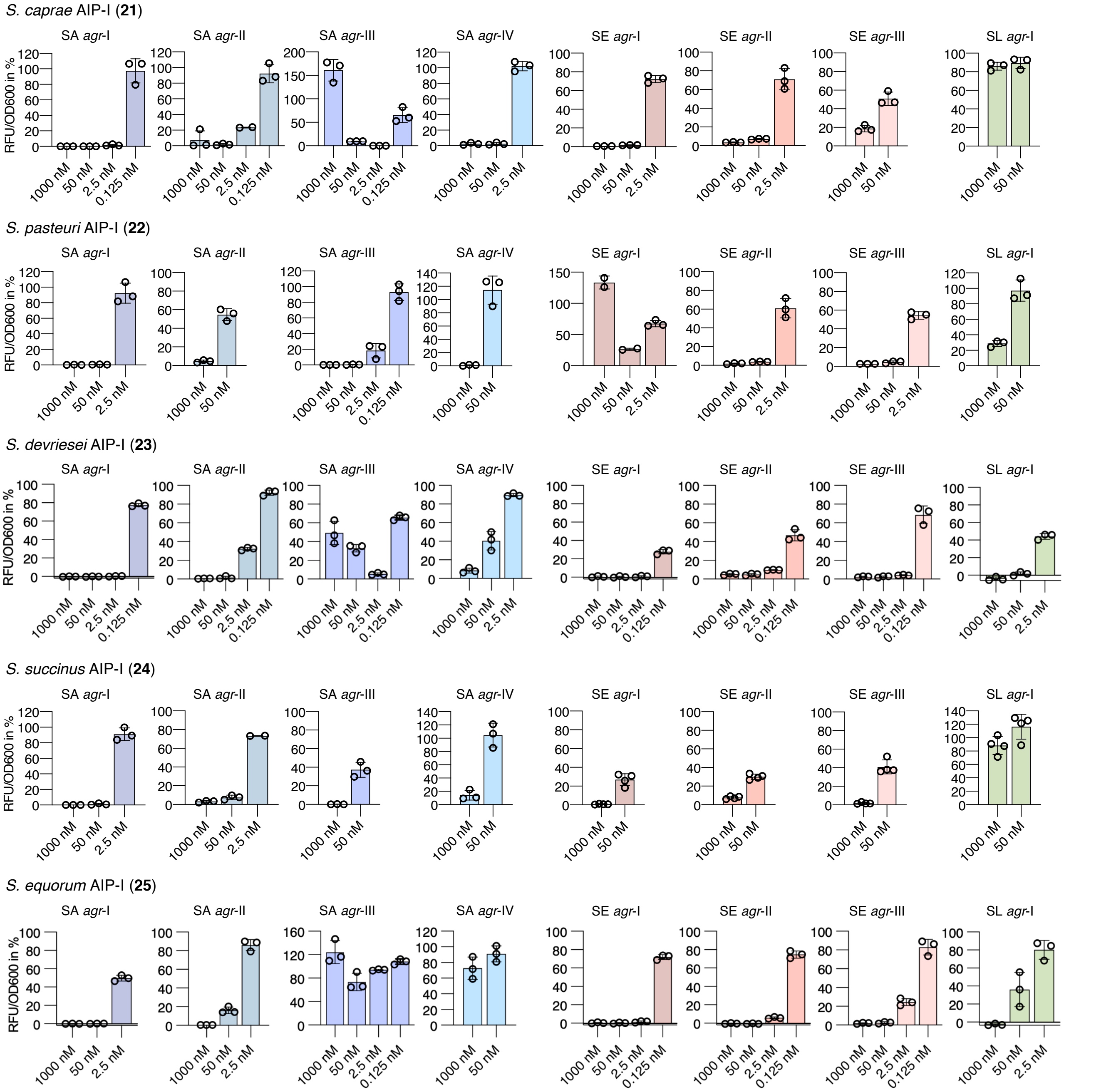

**Supplementary Figure S14. Fluorescence reporter strain assay for *agr* interference with native AIPs.** Fluorescent reporter strains of of *S. aureus* (SA), *S. epidermidis* (SE), S. *lugdunensis* (SL) were treated with AIPs at 1000 nM and 50 nM. AIPs were further tested at 2.5 nM and 0.125 nM in case >75% inhibition was observed at higher concentrations. Error bars are the SEM of at least two individual biological assays performed in technical triplicate.

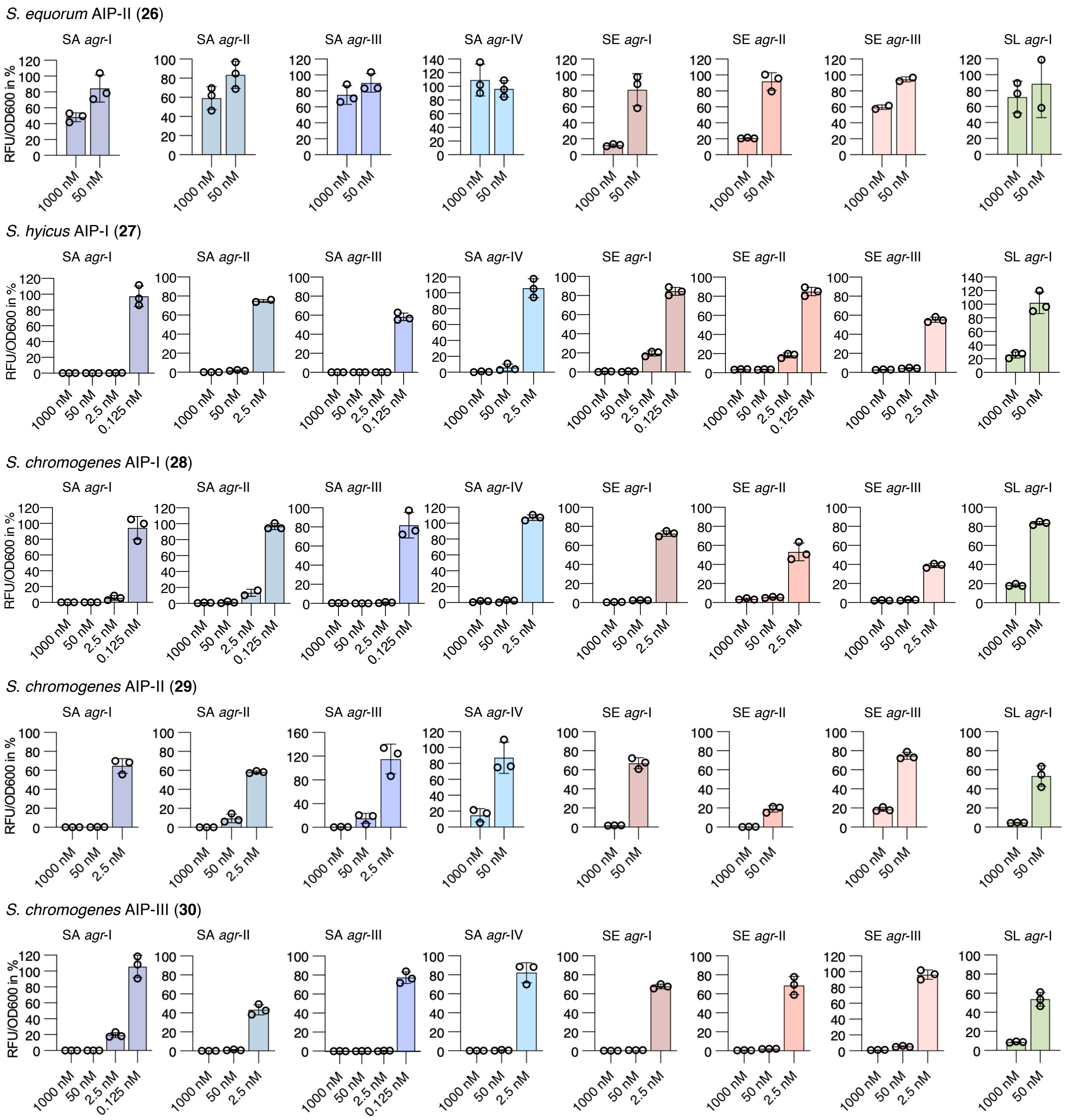

**Supplementary Figure S15. Fluorescence reporter strain assay for *agr* interference with native AIPs.** Fluorescent reporter strains of of *S. aureus* (SA), *S. epidermidis* (SE), S. *lugdunensis* (SL) were treated with AIPs at 1000 nM and 50 nM. AIPs were further tested at 2.5 nM and 0.125 nM in case >75% inhibition was observed at higher concentrations. Error bars are the SEM of at least two individual biological assays performed in technical triplicate.

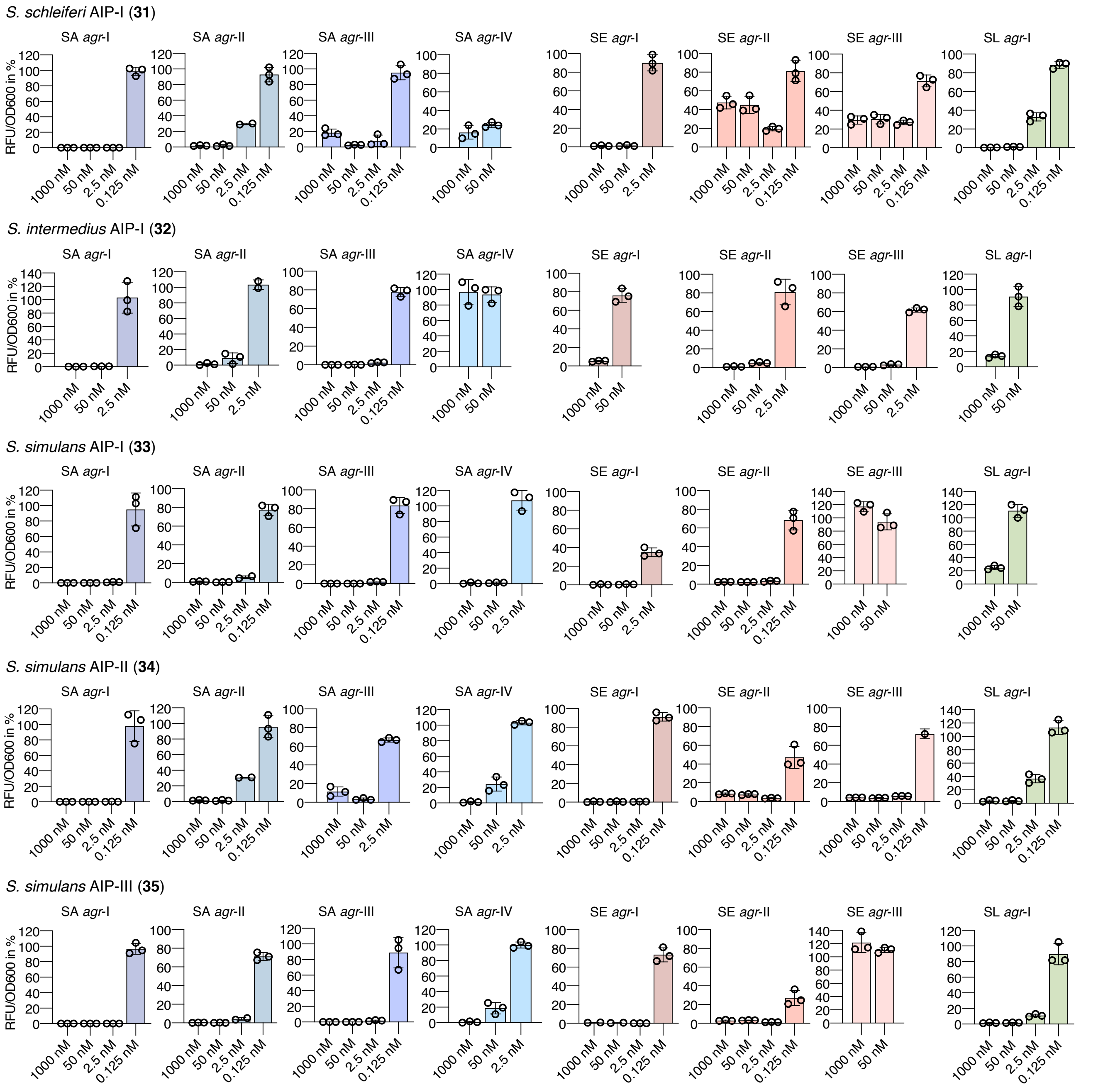

**Supplementary Figure S16. Fluorescence reporter strain assay for *agr* interference with native AIPs.** Fluorescent reporter strains of of *S. aureus* (SA), *S. epidermidis* (SE), S. *lugdunensis* (SL) were treated with AIPs at 1000 nM and 50 nM. AIPs were further tested at 2.5 nM and 0.125 nM in case >75% inhibition was observed at higher concentrations. Error bars are the SEM of at least two individual biological assays performed in technical triplicate.

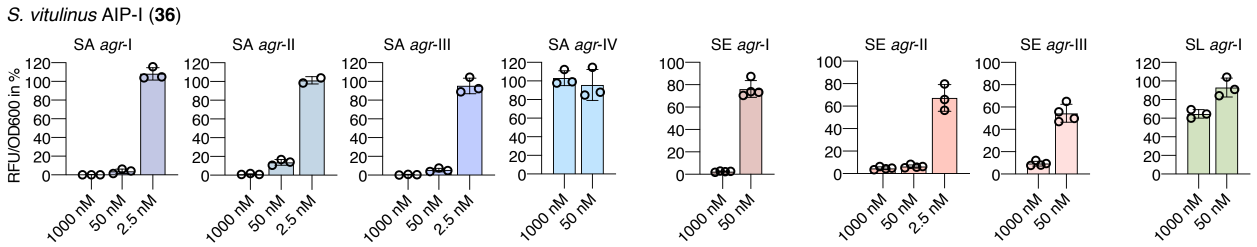

**Supplementary Figure S17. Fluorescence reporter strain assay for *agr* interference with native AIPs.** Fluorescent reporter strains of of *S. aureus* (SA), *S. epidermidis* (SE), S. *lugdunensis* (SL) were treated with AIPs at 1000 nM and 50 nM. AIPs were further tested at 2.5 nM and 0.125 nM in case >75% inhibition was observed at higher concentrations. Error bars are the SEM of at least two individual biological assays performed in technical triplicate.

#### Dose-response curves for *agr* interference for heat map validation

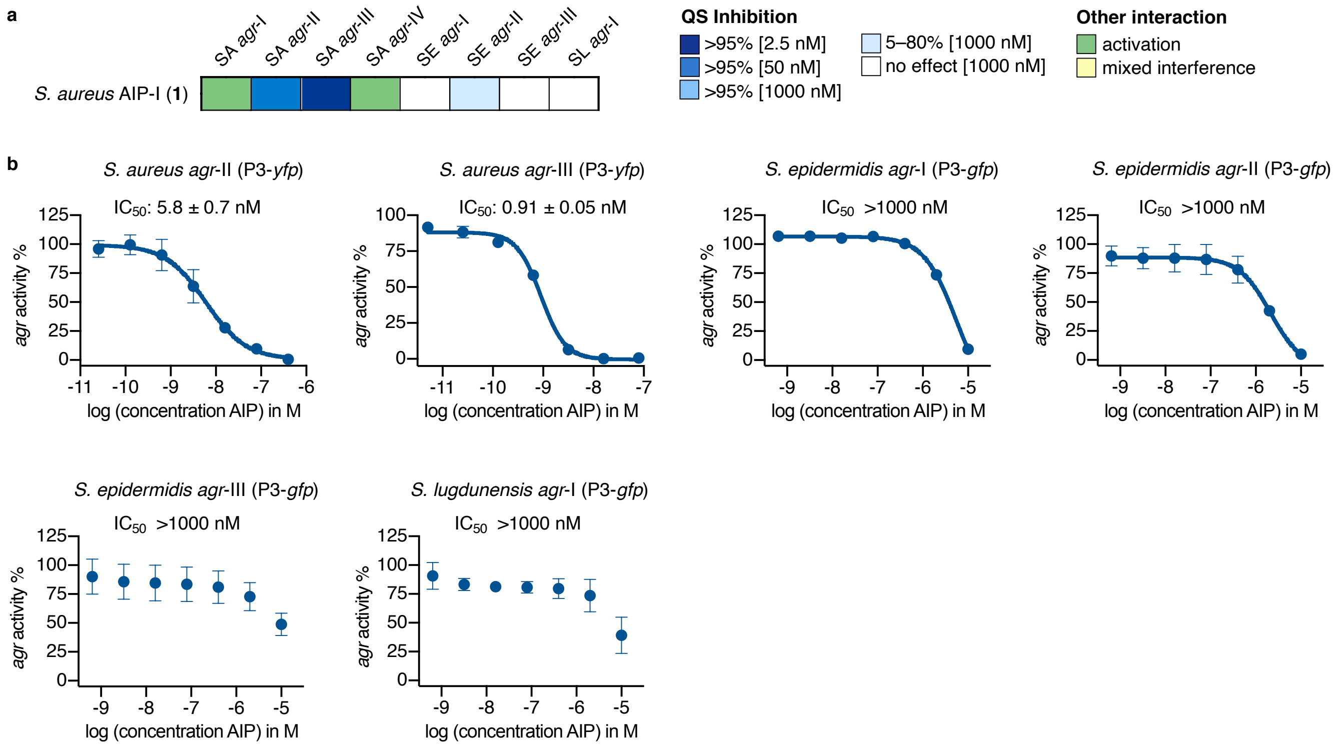

**Supplementary Figure S18. Heat map assay validation for *S. aureus* AIP-I (1) through full dose-response curves. a**, Heat map data of *S. aureus* AIP-I (**1**). **b**, IC50 values of *S. aureus* AIP-I (**1**) against reporter strains of of *S. aureus* (SA), *S. epidermidis* (SE), S. *lugdunensis* (SL) were determined through fluorescence readout as a measure of *agr* activity. The curves were generated from three individual assays performed in technical duplicate and shown error bars shown error bars are the standard deviation of the mean (SD).

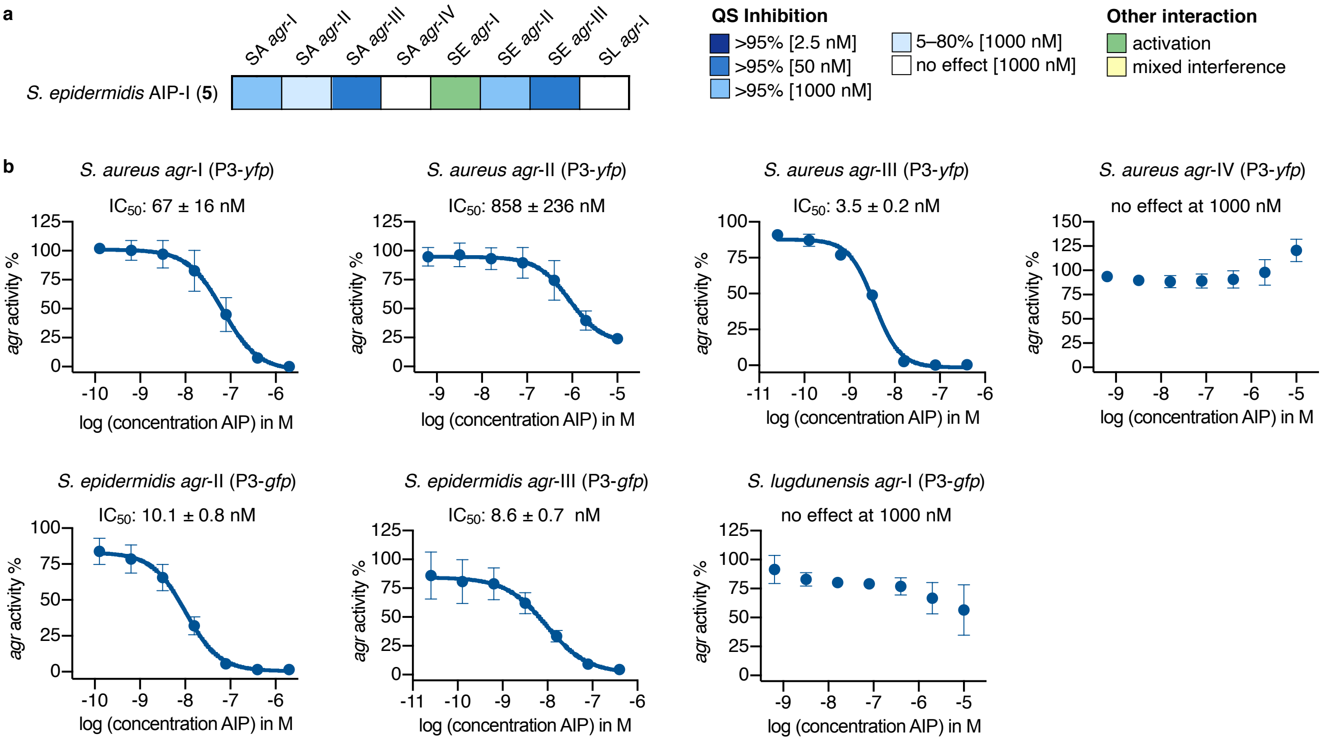

**Supplementary Figure S19. Heat map assay validation for *S. epidermidis* AIP-I (5) through full dose-response curves. a**, Heat map data of *S. epidermidis* AIP-I (**5**). **b**, IC50 values of *S. epidermidis* AIP-I (**5**) against reporter strains of of *S. aureus* (SA), *S. epidermidis* (SE), S. *lugdunensis* (SL) were determined through fluorescence readout as a measure of *agr* activity. The curves were generated from three individual assays performed in technical duplicate and shown error bars shown error bars are the standard deviation of the mean (SD).

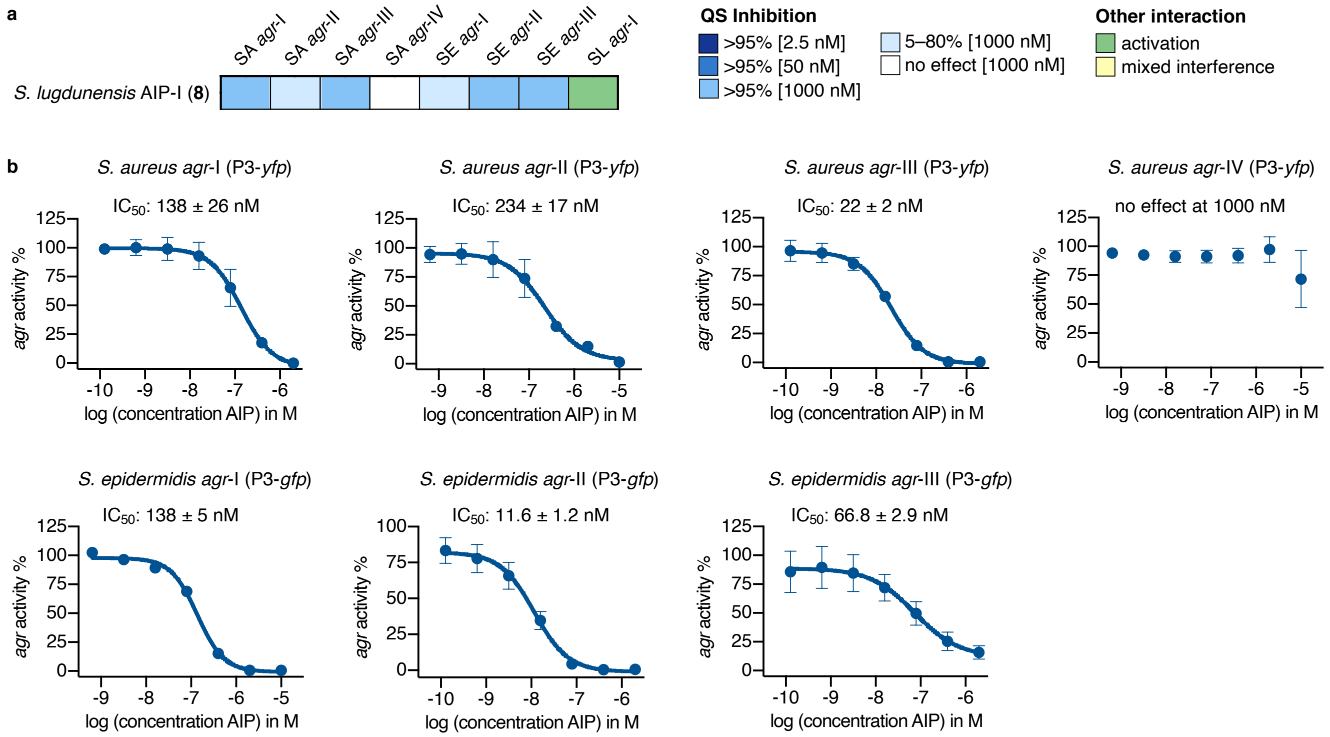

**Supplementary Figure S20. Heat map assay validation for *S. lugdunensis* AIP-I (8) through full dose-response curves. a**, Heat map data of *S. lugdunensis* AIP-I (**8**). **b**, IC50 values of *S. lugdunensis* AIP-I (**8**) against reporter strains of of *S. aureus* (SA), *S. epidermidis* (SE), S. *lugdunensis* (SL) were determined through fluorescence readout as a measure of *agr* activity. The curves were generated from three individual assays performed in technical duplicate and shown error bars shown error bars are the standard deviation of the mean (SD).

#### Dose-response curves for AgrC interference using *β*-lactamase reporter assays

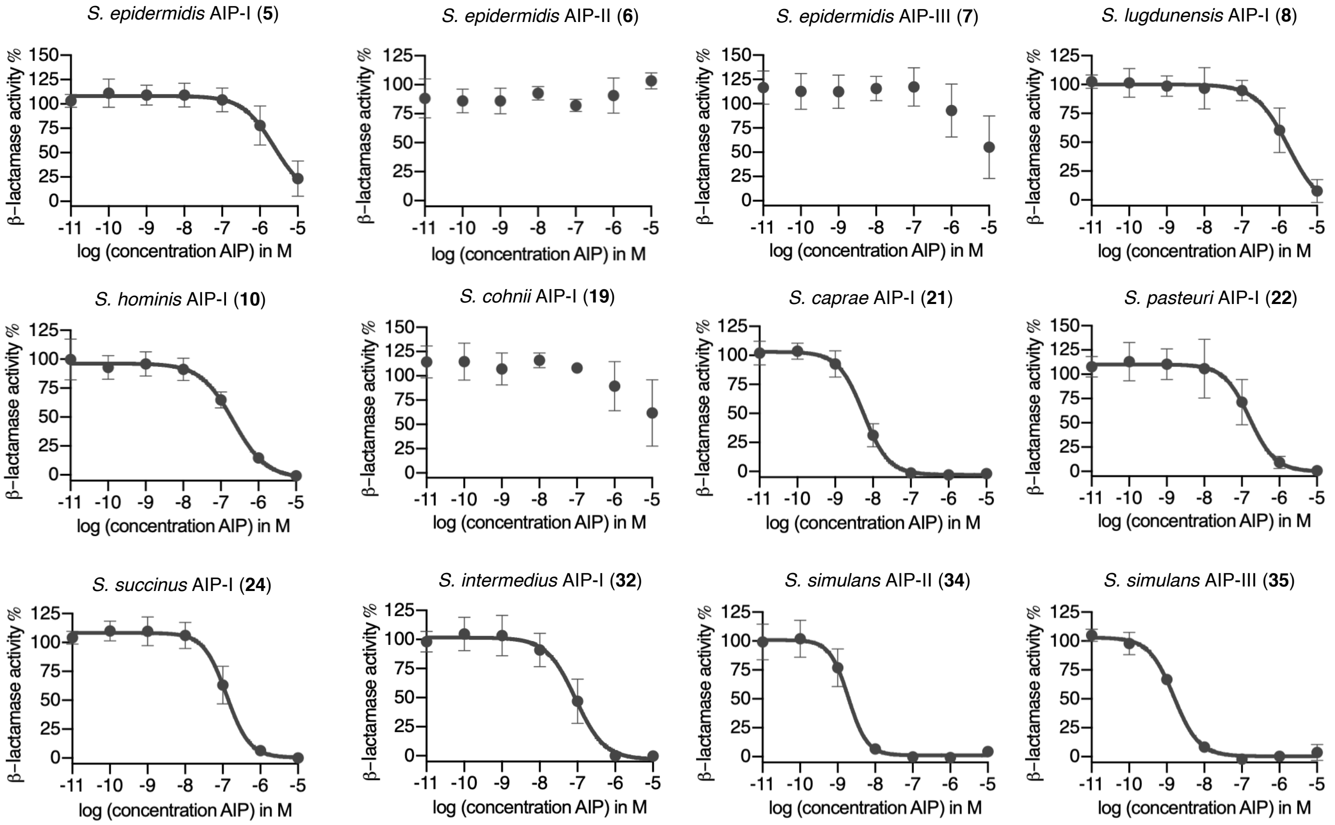

**Supplementary Figure S21. Dose-response curves of AgrC inhibition of *S. aureus agr*-I.** Inhibition properties of synthetic AIPs (10 µM to 10 pM) were determined through *β*-lactamase activity using the *S. aureus agr*-I reporter strain in the presence of 100 nM *S. aureus* AIP-I (**1**). The curves were generated from three individual assays performed in technical duplicate and shown error bars are the standard deviation of the mean (SD).

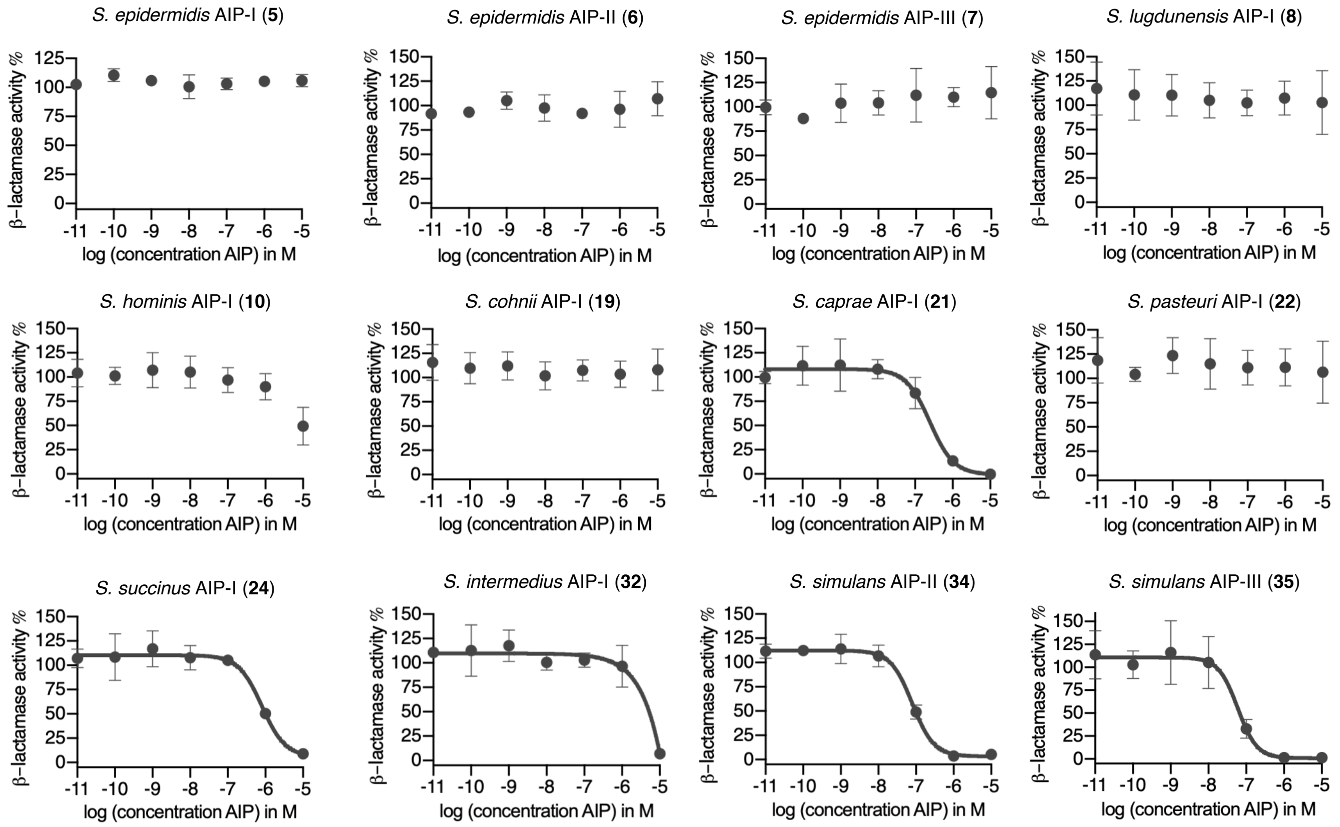

**Supplementary Figure S22. Dose-response curves of AgrC inhibition of *S. aureus agr*-II.** Inhibition properties of synthetic AIPs (10 μM to 10 pM) were determined through *β*-lactamase activity using the *S. aureus agr*-II reporter strain in the presence of 100 nM *S. aureus* AIP-II (**2**). The curves were generated from three individual assays performed in technical duplicate and shown error bars are the standard deviation of the mean (SD).

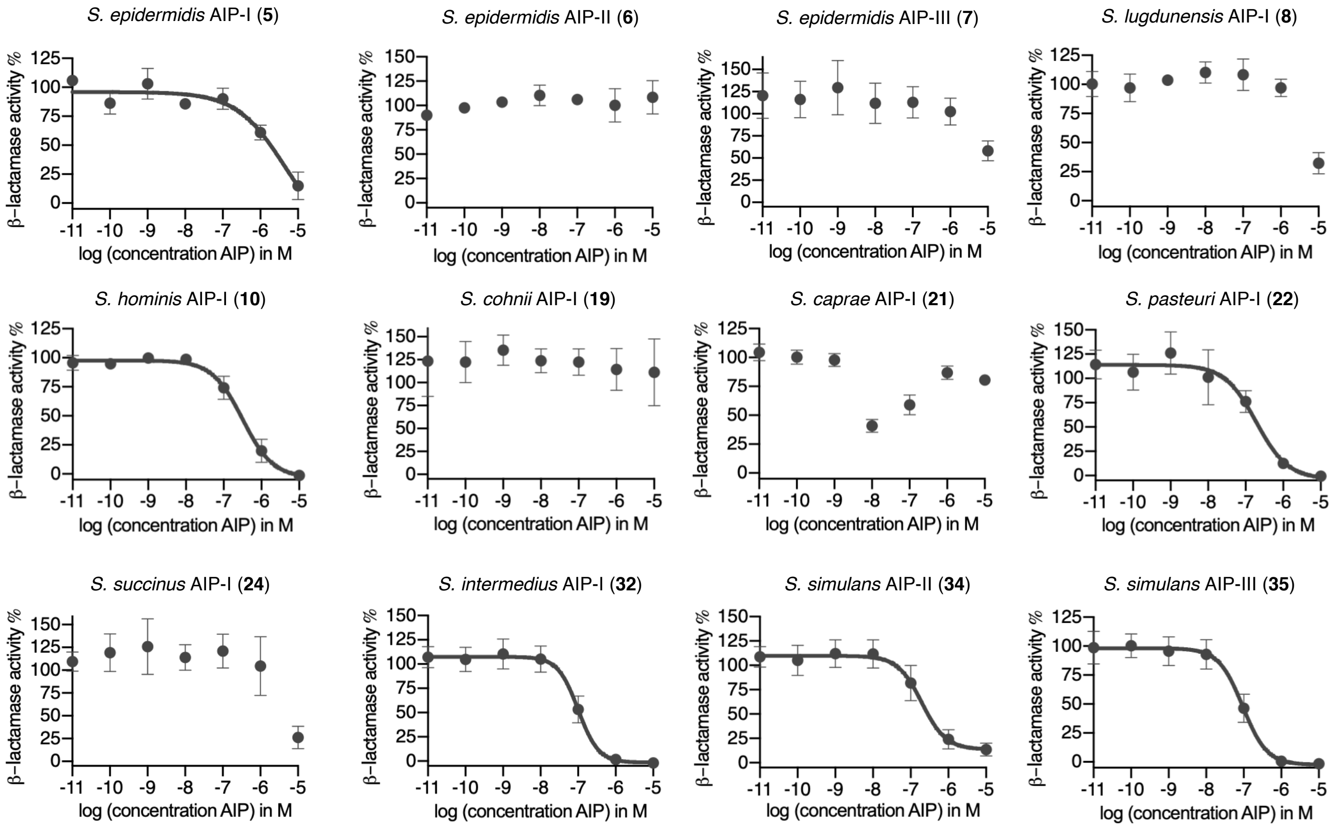

**Supplementary Figure S23. Dose-response curves of AgrC inhibition of *S. aureus agr*-III.** Inhibition properties of synthetic AIPs (10 µM to 10 pM) were determined through *β*-lactamase activity using the *S. aureus agr*-III reporter strain in the presence of 100 nM *S. aureus* AIP-III (**3**). The curves were generated from three individual assays performed in technical duplicate and shown error bars are the standard deviation of the mean (SD).

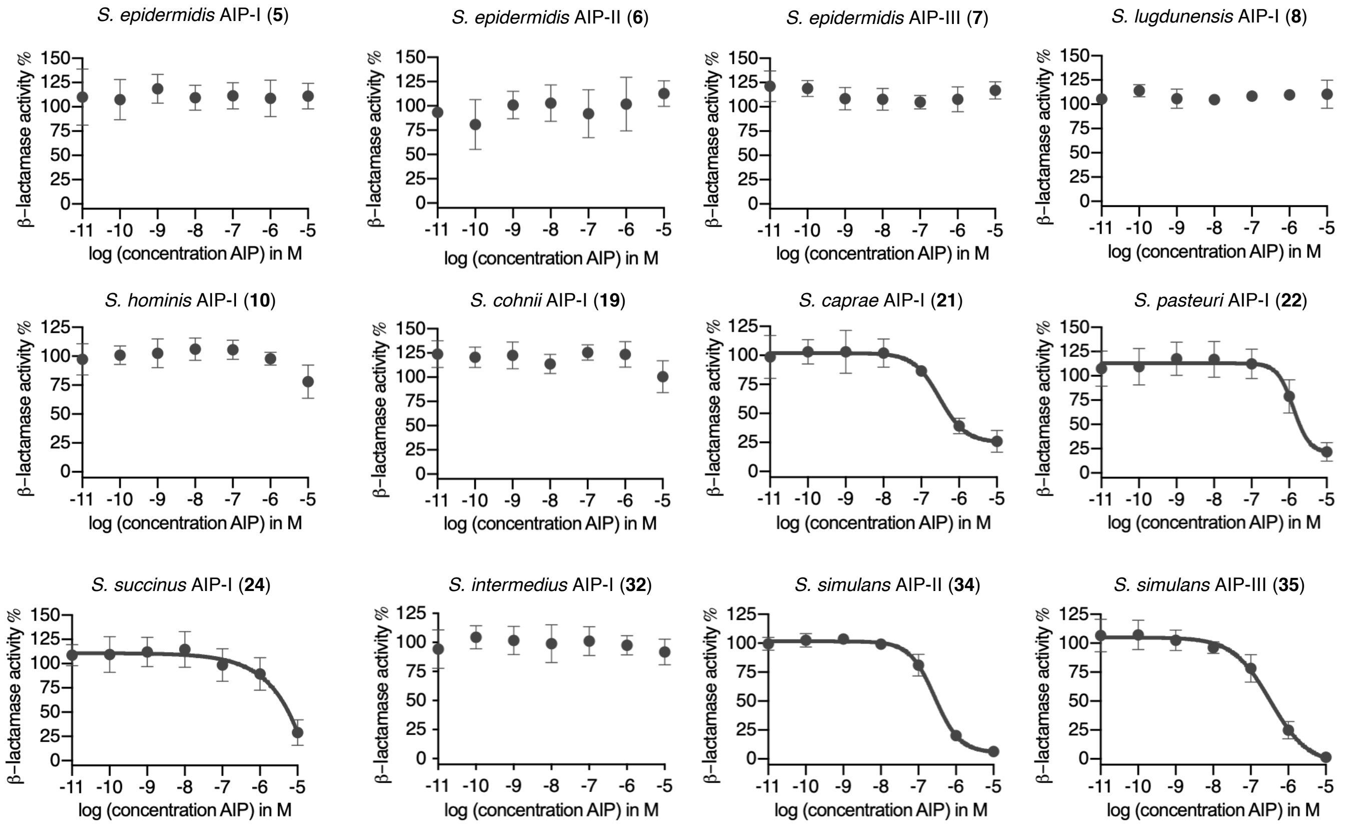

**Supplementary Figure S24. Dose-response curves of AgrC inhibition of *S. aureus agr*-IV.** Inhibition properties of synthetic AIPs (10 µM to 10 pM) were determined through *β*-lactamase activity using the *S. aureus agr*-IV reporter strain in the presence of 100 nM *S. aureus* AIP-IV (**4**). The curves were generated from three individual assays performed in technical duplicate and shown error bars are the standard deviation of the mean (SD).

#### Overnight growth and fluorescence curves in presence of synthetic AIPs

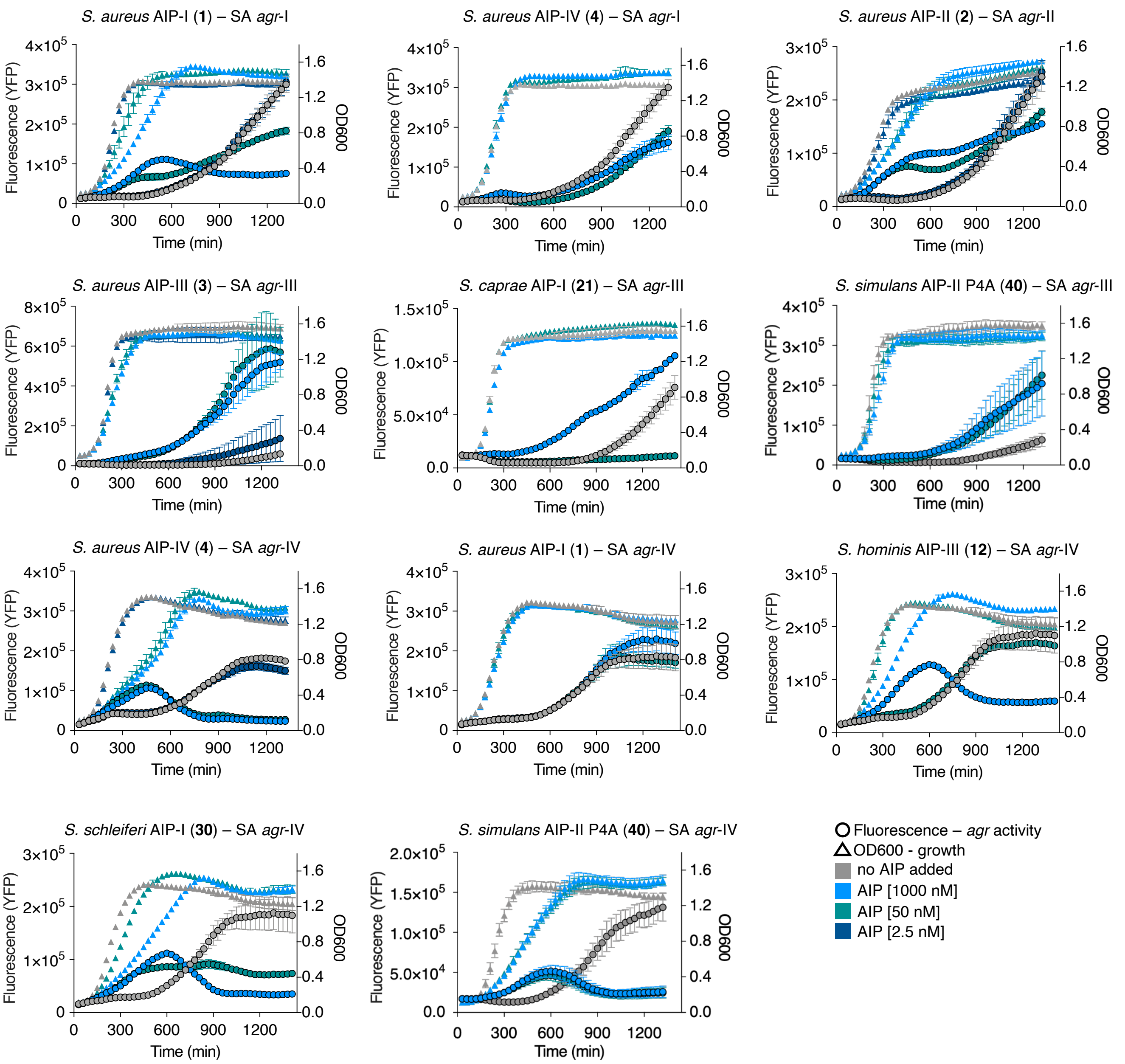

**Supplementary Figure S25. Fluorescence and $OD_{600}$ curves for peptides with *S. aureus* (SA) *agr*-I–IV.** YFP-producing reporter strains of SA were grown overnight in the absence or presence of AIPs (1000 nM, 50 nM and 2.5 nM) with continuous measurements of fluorescence (YFP) and $OD_{600}$. The curves represent single experiments performed in technical triplicate. Data points are the mean and error bars are the standard deviation of the mean (SD).

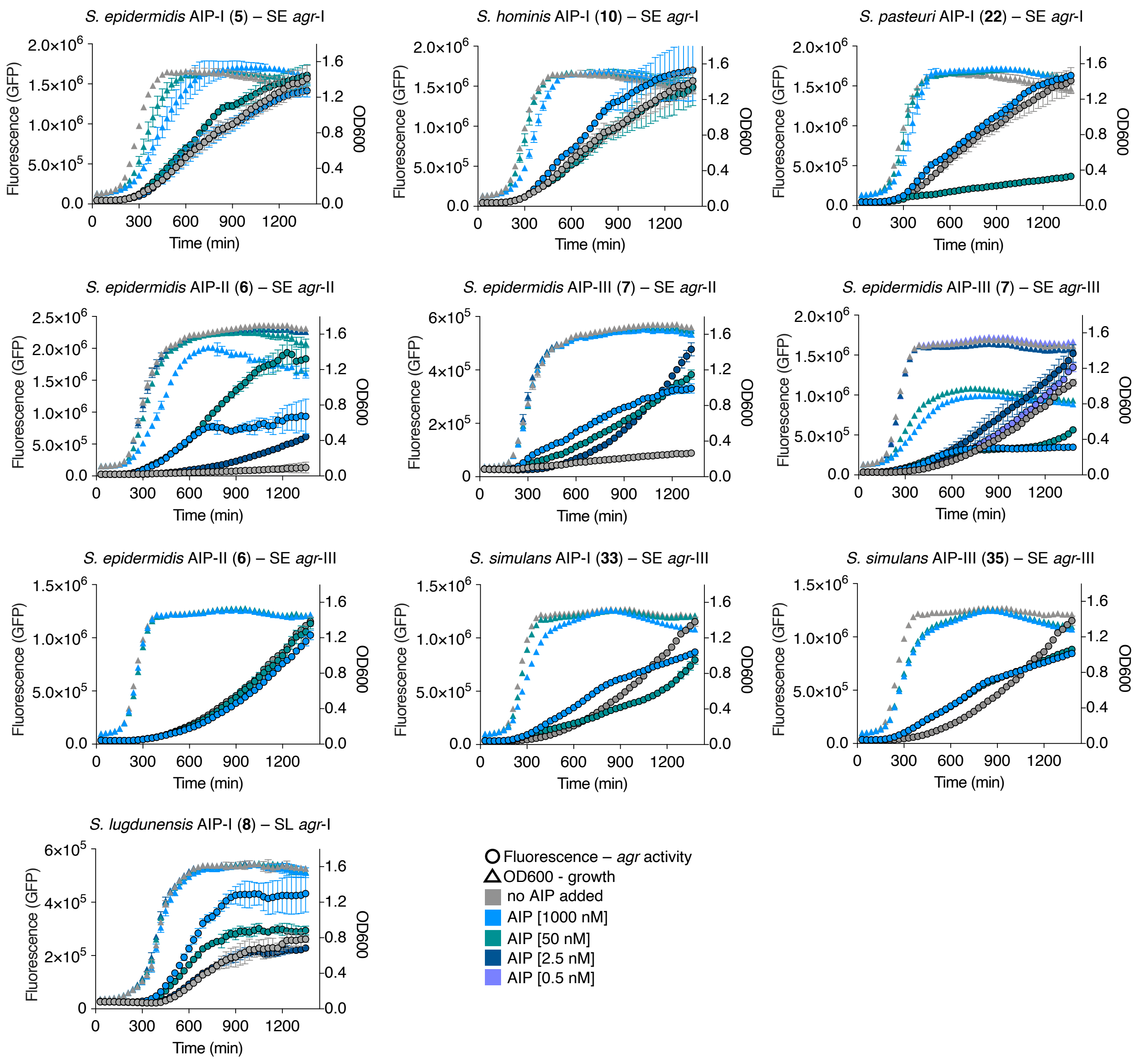

**Supplementary Figure S26. Fluorescence and OD$_{600}$ curves for peptides with *S. epidermidis* (SE) *agr*-I–III and *S. lugdunensis* (SL) *agr*-I.** sGFP-producing reporter strains of SE and SL were grown overnight in the absence or presence of AIPs (1000 nM, 50 nM, 2.5 nM and 0.5 nM) with continuous measurements of fluorescence (GFP) and OD$_{600}$. The curves represent single experiments performed in technical triplicate. Data points are the mean and error bars are the SD.

#### Dose-response curves for *agr* interference of *S. simulans* AIPs

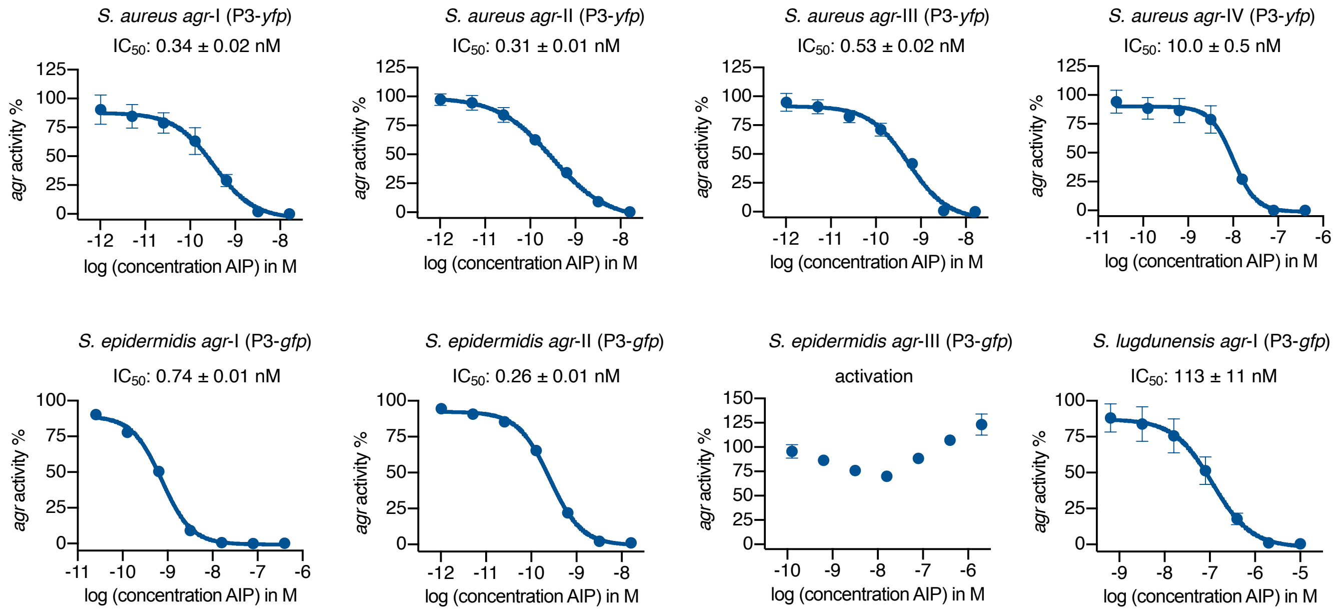

**Supplementary Figure S27. Dose-response ($IC_{50}$) curves for *agr* inhibition by *S. simulans* AIP-I (33).** Inhibition properties of *S. simulans* AIP-I (**33**) were determined through fluorescence readout as a measure of *agr* activity against reporter strains of of *S. aureus* (SA), *S. epidermidis* (SE), S. *lugdunensis* (SL). The curves were generated from three individual assays performed in technical duplicate and shown error bars are the standard deviation of the mean (SD).

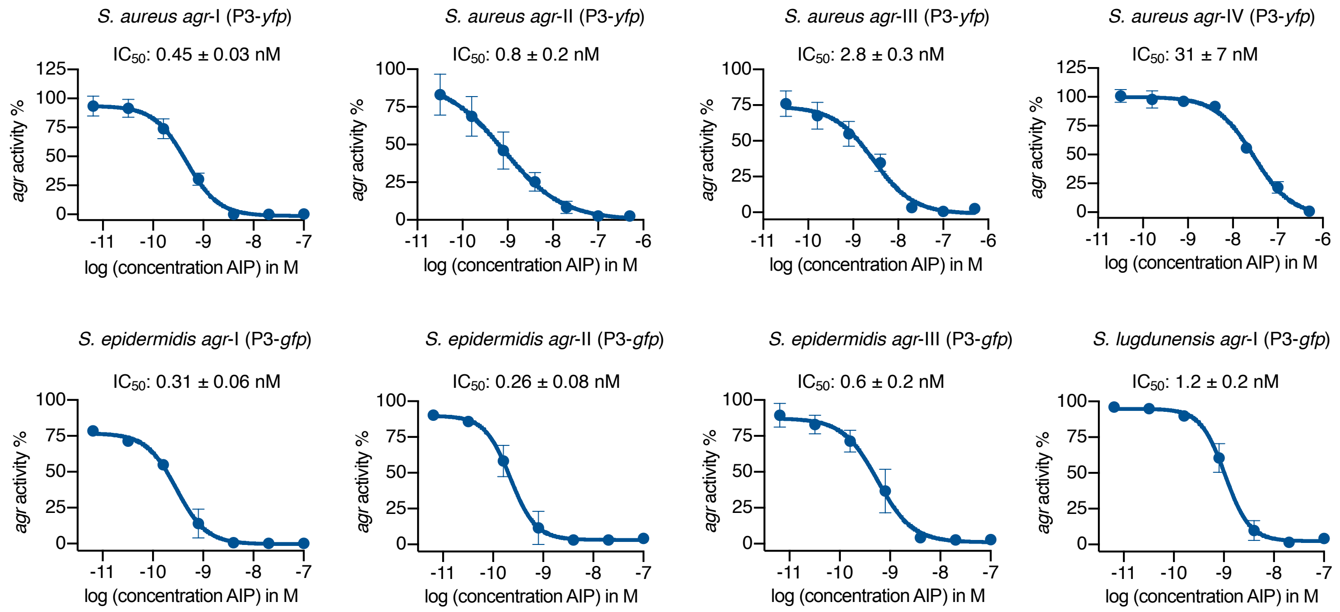

**Supplementary Figure S28. Dose-response ($IC_{50}$) curves for *agr* inhibition by *S. simulans* AIP-II (34).** Inhibition properties of *S. simulans* AIP-II (**34**) were determined through fluorescence readout as a measure of *agr* activity against reporter strains of of *S. aureus* (SA), *S. epidermidis* (SE), S. *lugdunensis* (SL). The curves were generated from three individual assays performed in technical duplicate and shown error bars are the standard deviation of the mean (SD).

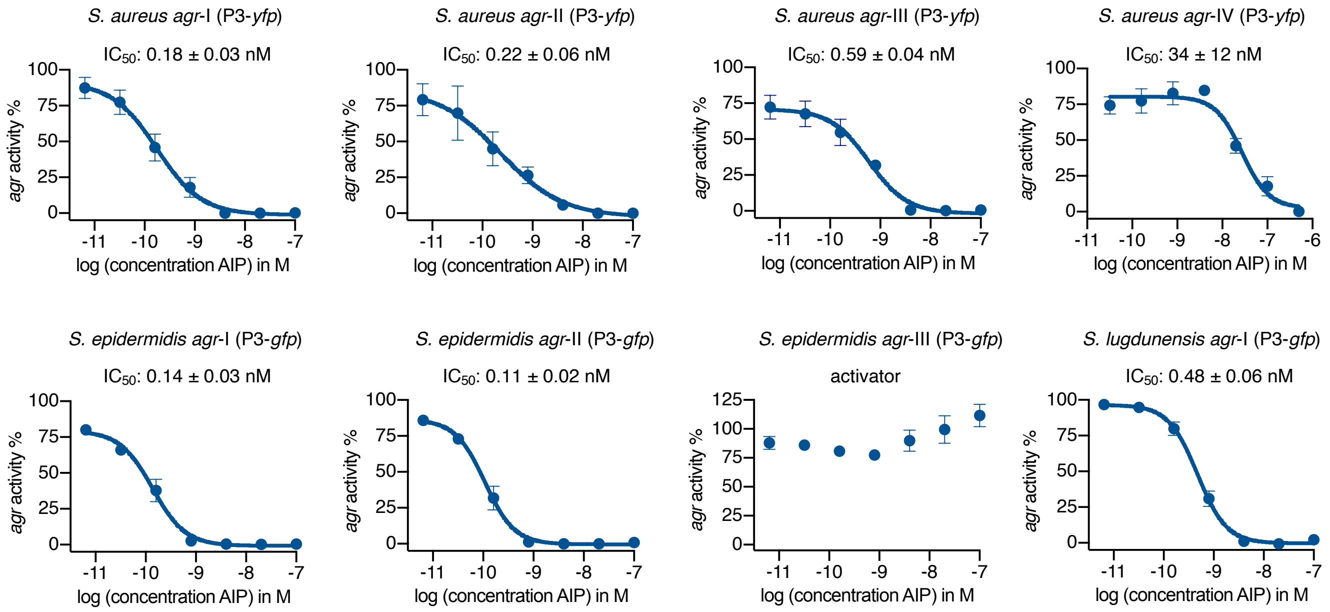

**Supplementary Figure S29. Dose-response ($IC_{50}$) curves for *agr* inhibition by *S. simulans* AIP-III (35).** Inhibition properties of *S. simulans* AIP-III (**35**) were determined through fluorescence readout as a measure of *agr* activity against reporter strains of of *S. aureus* (SA), *S. epidermidis* (SE), S. *lugdunensis* (SL). The curves were generated from three individual assays performed in technical duplicate and shown error bars are the standard deviation of the mean (SD).

#### Bar graphs for *agr* interference of *S. simulans* AIPs SAR study

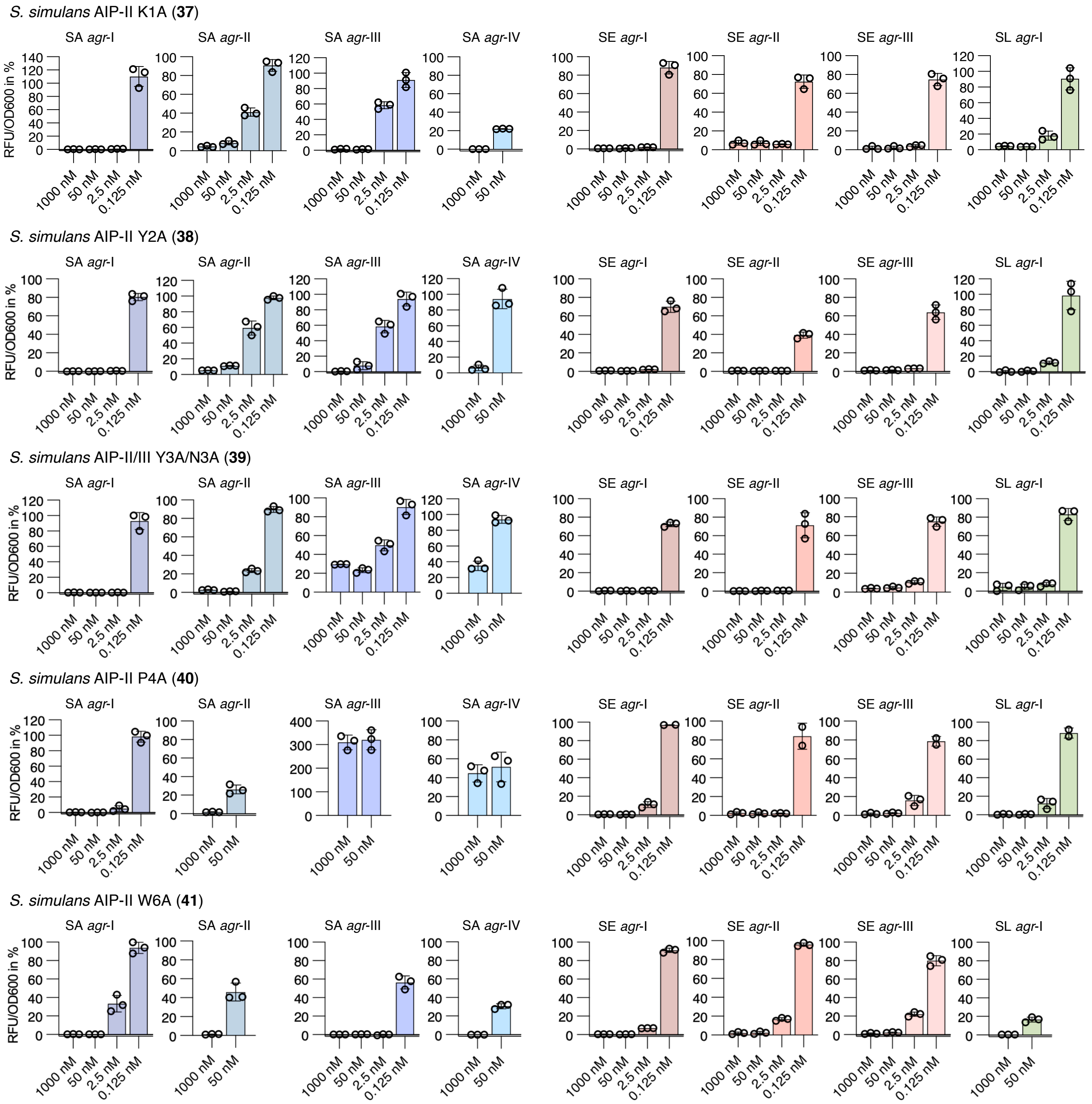

**Supplementary Figure S30. Fluorescence reporter strain assay for *agr* interference with SAR peptides.** Fluorescent reporter strains of of *S. aureus* (SA), *S. epidermidis* (SE), S. *lugdunensis* (SL) were treated with AIPs at 1000 nM and 50 nM. AIPs were further tested at 2.5 nM and 0.125 nM in case >75% inhibition was observed at higher concentrations. Error bars are the SEM of at least two individual biological assays performed in technical triplicate.

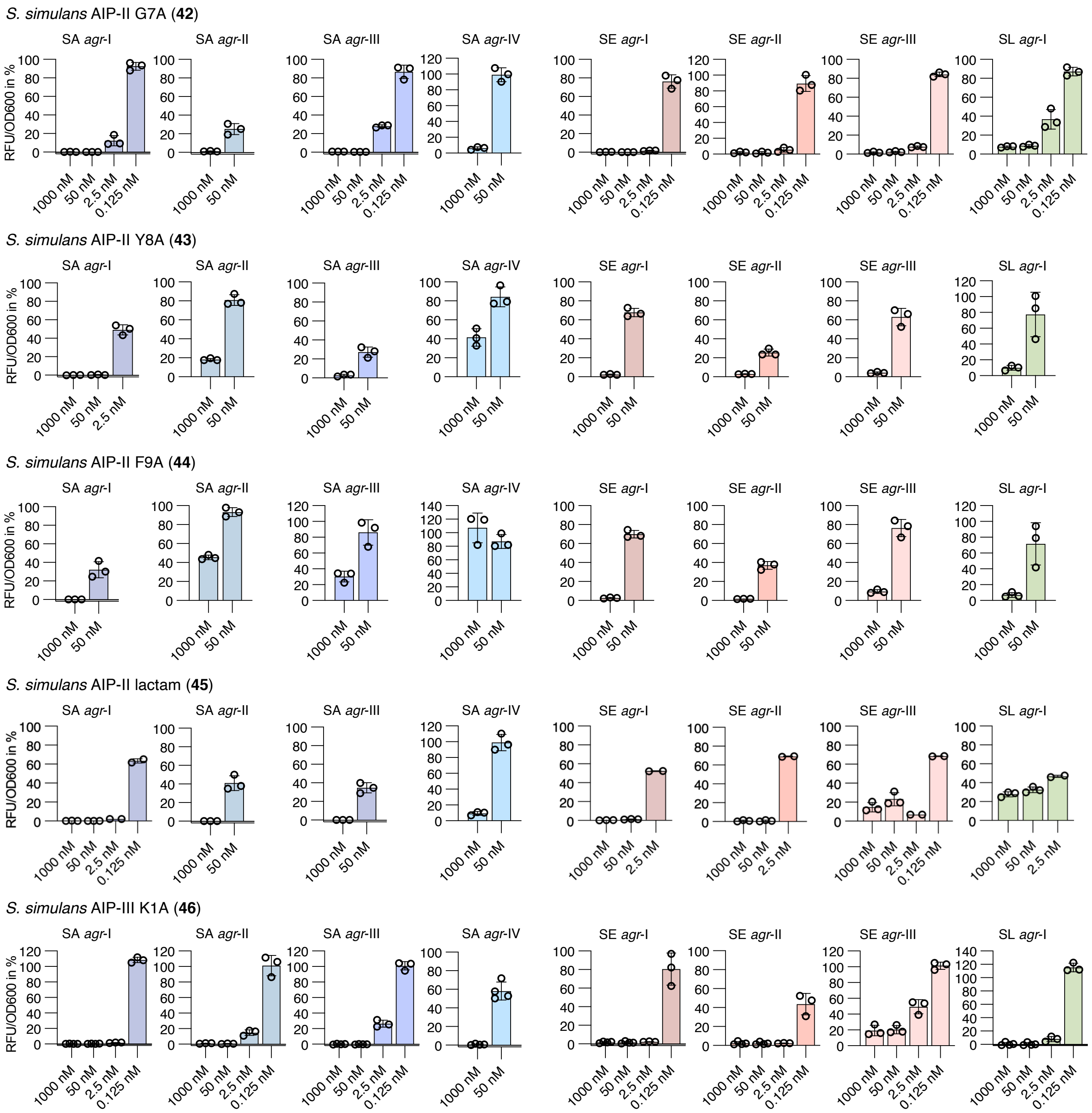

**Supplementary Figure S31. Fluorescence reporter strain assay for *agr* interference with SAR peptides.** Fluorescent reporter strains of of *S. aureus* (SA), *S. epidermidis* (SE), S. *lugdunensis* (SL) were treated with AIPs at 1000 nM and 50 nM. AIPs were further tested at 2.5 nM and 0.125 nM in case >75% inhibition was observed at higher concentrations. Error bars are the SEM of at least two individual biological assays performed in technical triplicate.

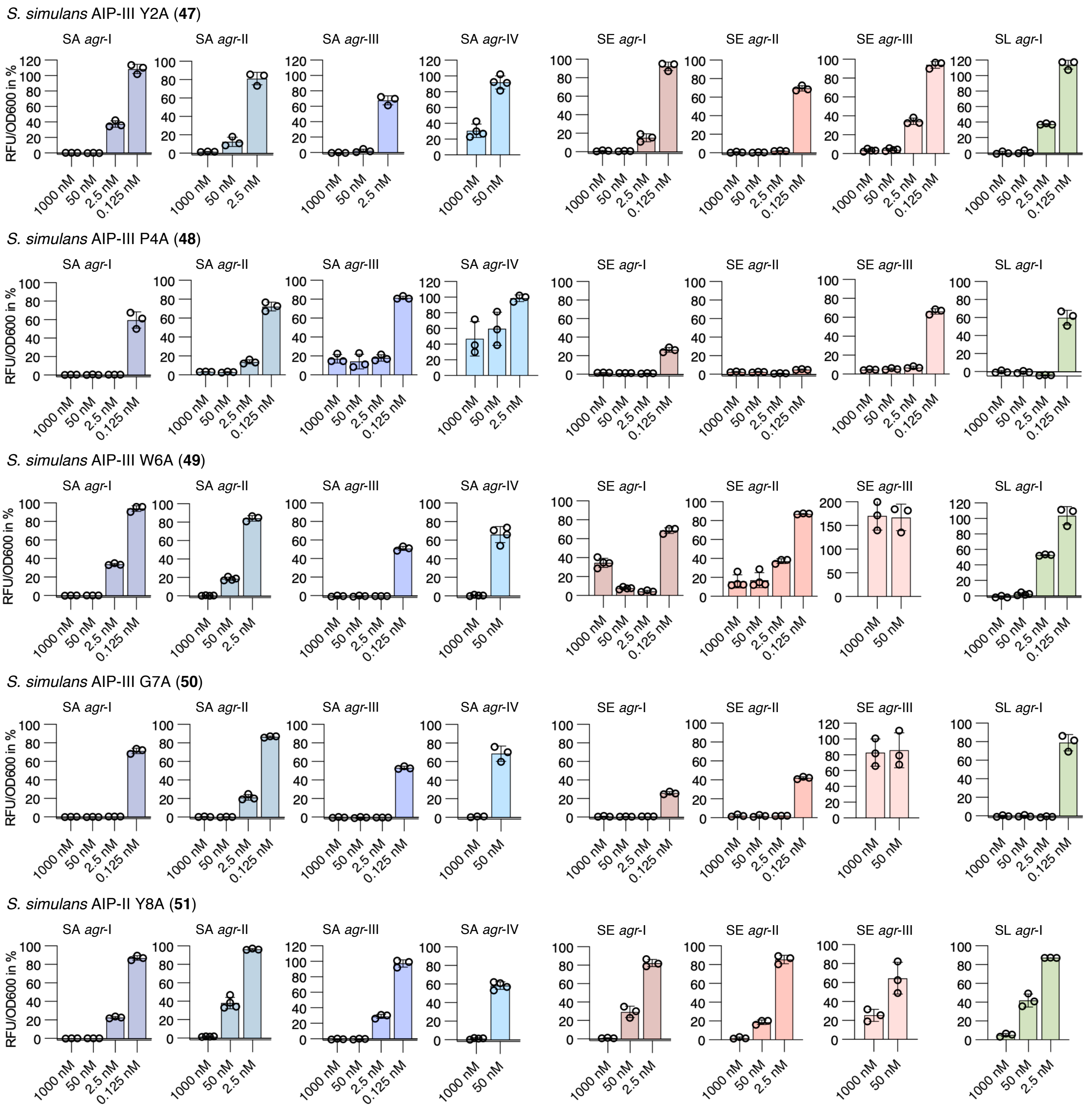

**Supplementary Figure S32. Fluorescence reporter strain assay for *agr* interference with SAR peptides.** Fluorescent reporter strains of of *S. aureus* (SA), *S. epidermidis* (SE), S. *lugdunensis* (SL) were treated with AIPs at 1000 nM and 50 nM. AIPs were further tested at 2.5 nM and 0.125 nM in case >75% inhibition was observed at higher concentrations. Error bars are the SEM of at least two individual biological assays performed in technical triplicate.

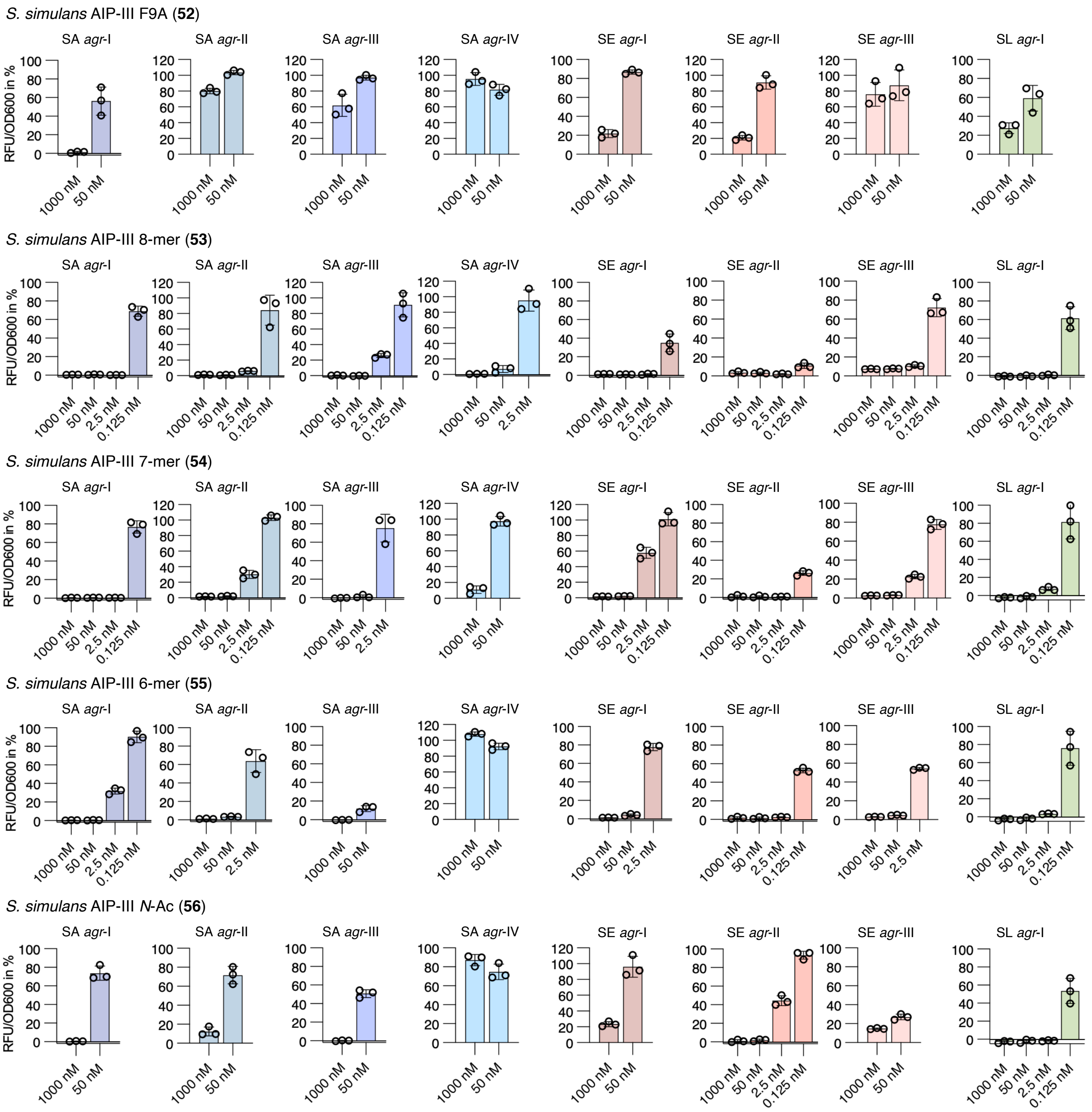

**Supplementary Figure S33. Fluorescence reporter strain assay for *agr* interference with SAR peptides.** Fluorescent reporter strains of of *S. aureus* (SA), *S. epidermidis* (SE), S. *lugdunensis* (SL) were treated with AIPs at 1000 nM and 50 nM. AIPs were further tested at 2.5 nM and 0.125 nM in case >75% inhibition was observed at higher concentrations. Error bars are the SEM of at least two individual biological assays performed in technical triplicate.

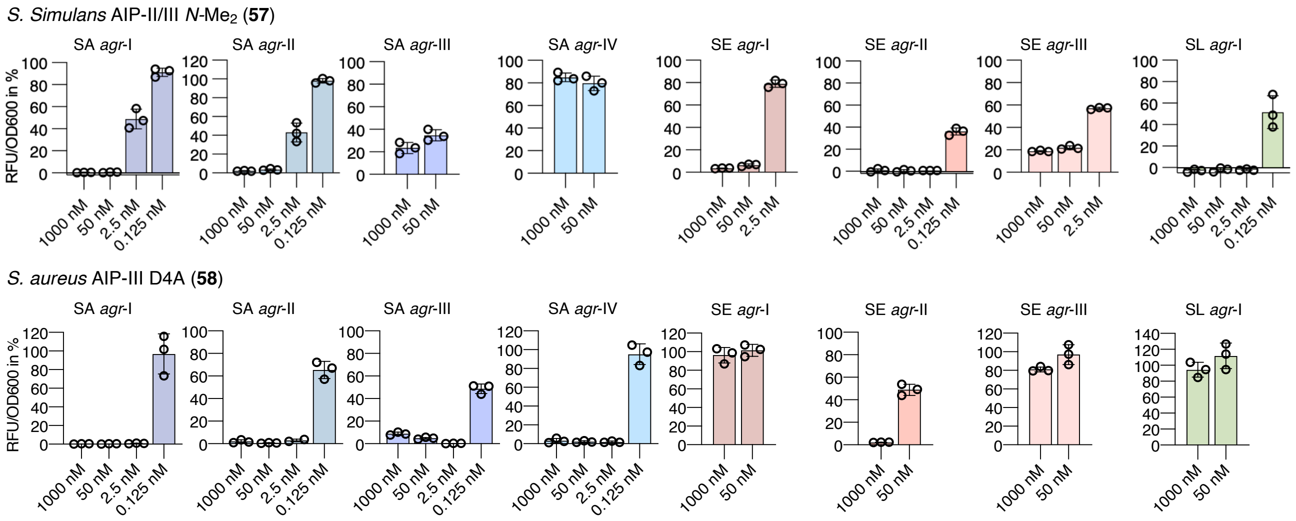

**Supplementary Figure S34. Fluorescence reporter strain assay for *agr* interference with SAR peptides.** Fluorescent reporter strains of of *S. aureus* (SA), *S. epidermidis* (SE), S. *lugdunensis* (SL) were treated with AIPs at 1000 nM and 50 nM. AIPs were further tested at 2.5 nM and 0.125 nM in case >75% inhibition was observed at higher concentrations. Error bars are the SEM of at least two individual biological assays performed in technical triplicate.

#### Overnight growth and fluorescence curves of *spa*-GFP agr deactivation assay

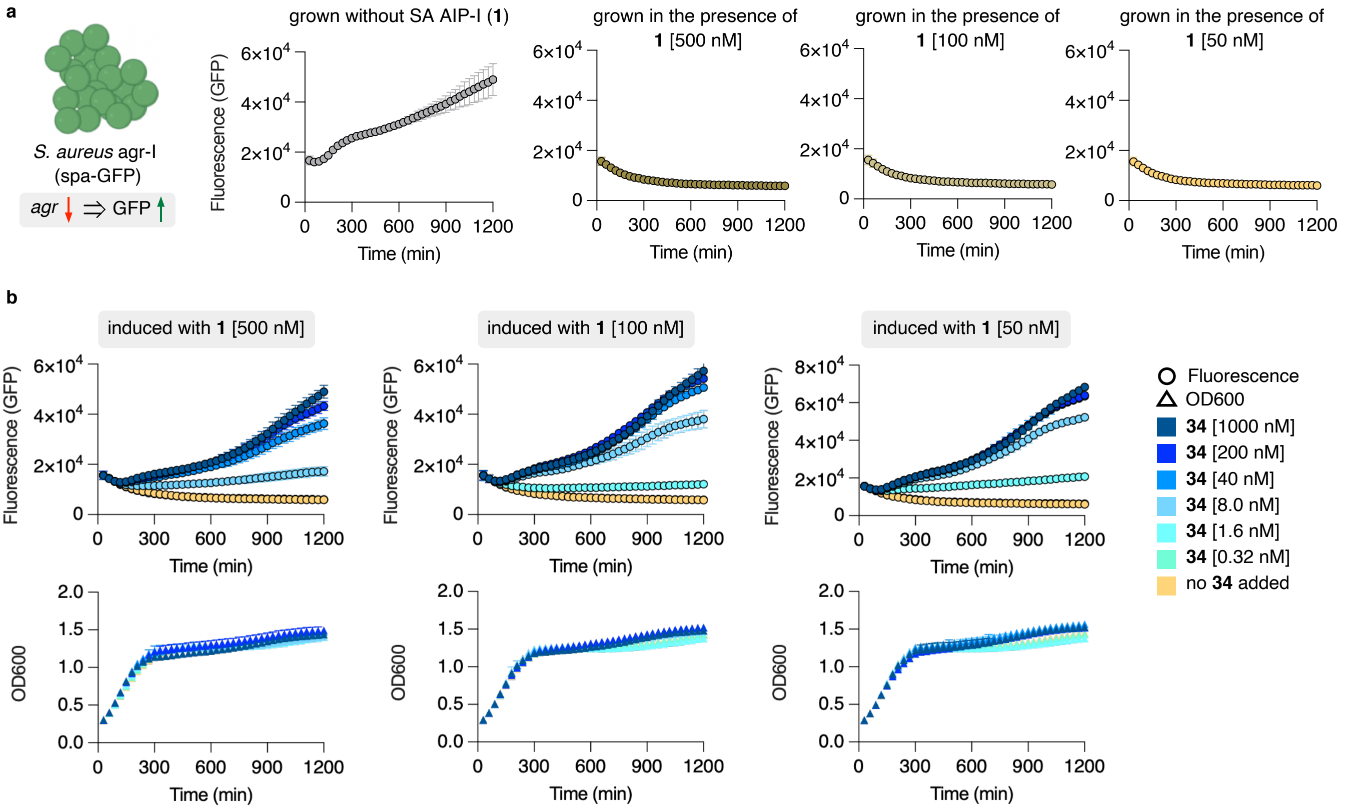

**Supplementary Figure S35. Fluorescence and $OD_{600}$ curves for *spa*-GFP *S. aureus agr*-I reporter strain assays.** **a**, Fluorescence monitored of *S. aureus agr*-I *spa*-GFP reporter strain grown in the presence and absence of SA AIP-I (**1**). **b**, Fluorescence and growth curves of *S. aureus agr*-I *spa*-GFP induced with different concentrations of **1** (50, 100 and 500 nM) for 2-3 h followed by the addition of *S. simulans* AIP-II (**34**) at different concentrations (1000–0.32 nM). The curves were generated from three individual assays performed in technical duplicate and shown error bars are the standard deviation of the mean (SD).

#### 2. Supplementary schemes

##### Synthesis of thiolactone-containing AIPs

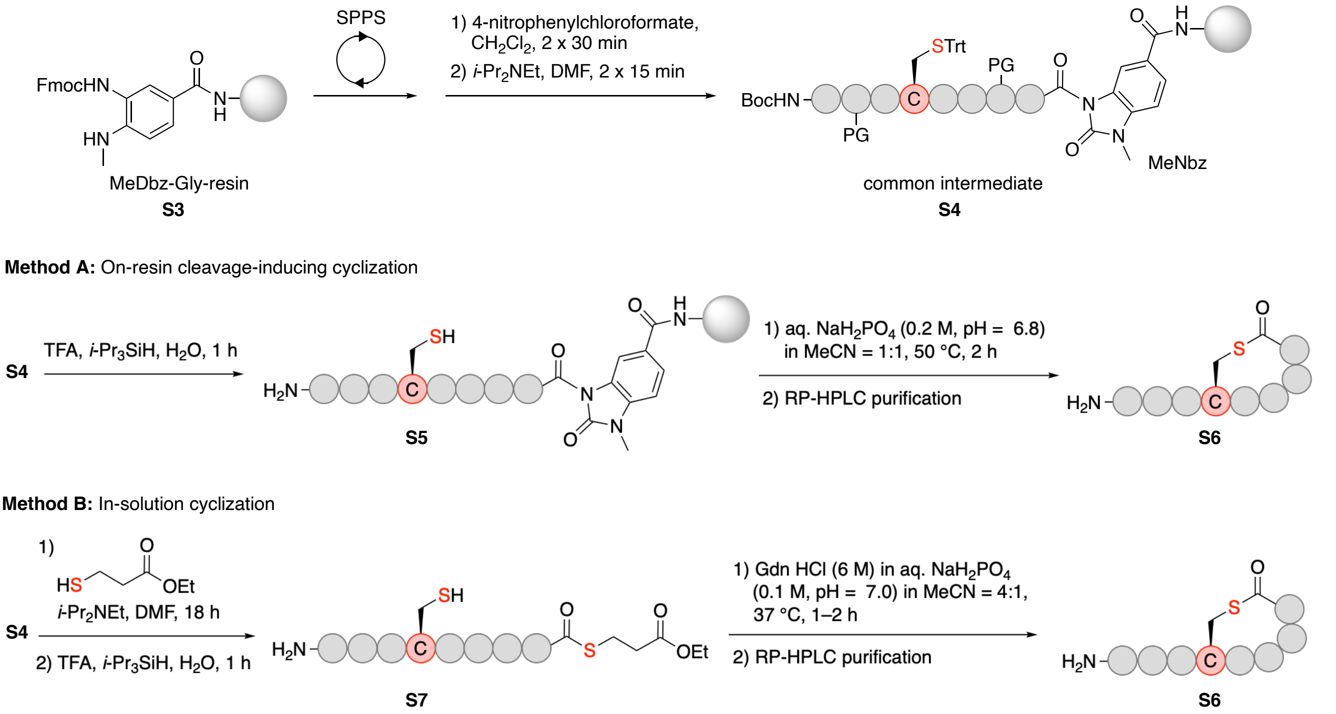

**Supplementary Scheme S1. Synthesis of thiolactone-containing AIPs using the MeDbz linker.** *N*-acyl-benzimidazolinone (Nbz) intermediates **S4** were synthesized on MeDbz-Gly resin and used for method A and B. In method A, **S4** was used for an on-resin cleavage-inducing cyclization protocol and in method B **S4** was treated with a thiol to induce a cleavage-inducing thioesterification followed by a chemoselective cyclization in solution to afford AIPs **S6**.

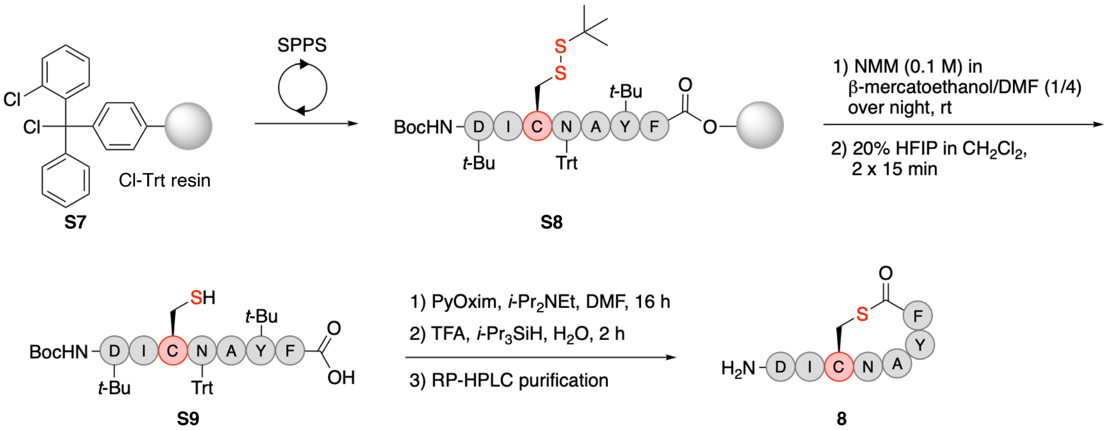

**Supplementary Scheme S2. Synthesis of *S. lugdunensis* AIP-I (8) via PyOxim-mediated thiolactonization.** The linear peptide **S8** was synthesized on Cl-Trt resin **S7** and the cysteine side chain protecting group removed on-resin. The partially-protected peptide **S9** was released from the resin and the peptide cyclized via PyOxim-mediated thiolactonization to give peptide **8**.

#### Synthesis of *S. intermedius* AIP

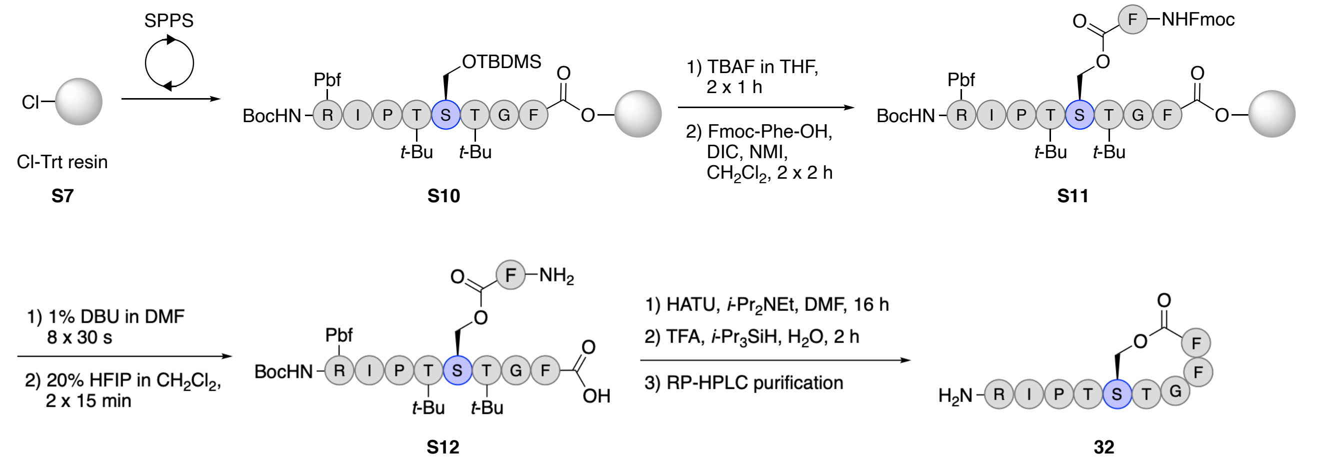

**Supplementary Scheme S3. Synthesis of *S. intermedius* AIP-I (32) using an on-resin esterification strategy.** The linear peptide **S10** was synthesized on Cl-Trt resin **S7** and the serine side chain protecting group was removed on-resin followed by esterification to give **S11**. The partially-protected peptide **S12** was released from the resin and the peptide cyclized via HATU-mediated amide coupling to give peptide **32**.

#### Synthesis of *S simulans* AIP-II lactam

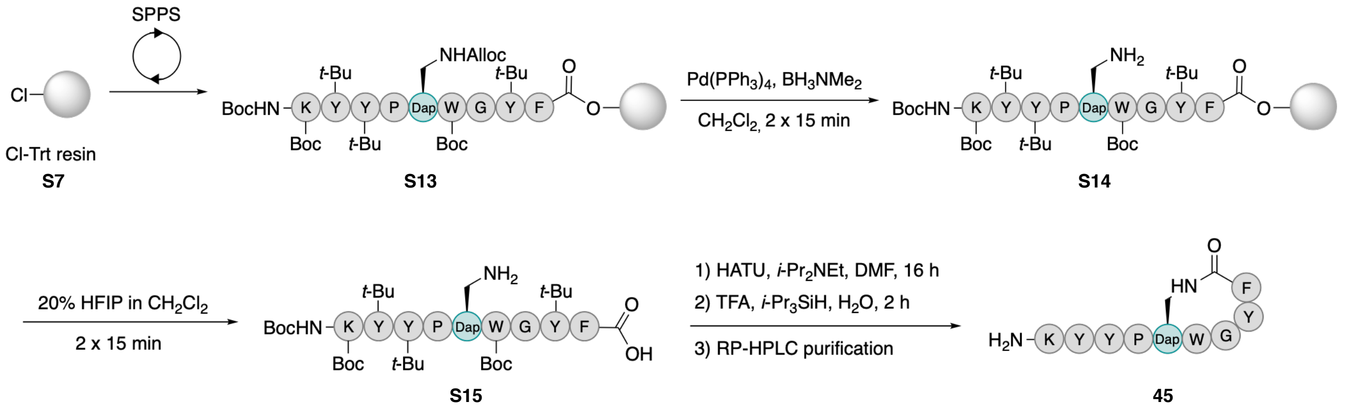

**Supplementary Scheme S4. Synthesis of *S. simulans* AIP-II lactam (45).** The linear peptide **S13** was synthesized on Cl-Trt resin **S7** and the 2,3-diaminopropionic acid (Dap) side chain protecting group was removed on-resin. The partially-protected peptide **S15** was released from the resin and the peptide cyclized via HATU-mediated amide coupling to give peptide **45**.

### 3. Supplementary tables

#### Staphylococci species and sequences of known AIPs.

##### Supplementary Table S1. Information about staphylococci species and sequences of known AIPs.

| Species AIP | Sequence | Associated Host | Phylogenetic species group |
|---|---|---|---|
| *S. aureus* AIP-I (**1**) [*S. argenteus* AIP-I] | YST-[CDFIM] | human/animal | Epidermidis-Aureus |
| *S. aureus* AIP-II (**2**) | GVNA-[CSSLF] | human/animal | Epidermidis-Aureus |
| *S. aureus* AIP-III (**3**) | IN-[CDFLL] | human/animal | Epidermidis-Aureus |
| *S. aureus* AIP-IV (**4**) [*S. schweitzeri* AIP-I] | YST-[CYFIM] | human/animal | Epidermidis-Aureus |
| *S. epidermidis* AIP-I (**5**) | DSV-[CASYF] | human/animal | Epidermidis-Aureus |
| *S. epidermidis* AIP-II (**6**) | NASKYNP-[CSNYL] | human/animal | Epidermidis-Aureus |
| *S. epidermidis* AIP-III (**7**) | NAAKYNP-[CASYL] | human/animal | Epidermidis-Aureus |
| *S. lugdunensis* AIP-I (**8**) | DI-[CNAYF] | human | Epidermidis-Aureus |
| *S. lugdunensis* AIP-II (**9**) | DM-[CNGYF] | human | Epidermidis-Aureus |
| *S. hominis* AIP-I (**10**) | SYNV-[CGGYF] | human | Epidermidis-Aureus |
| *S. hominis* AIP-II (**11**) | SYSP-[CATYF] | human | Epidermidis-Aureus |
| *S. hominis* AIP-III (**12**) | TYST-[CYGYF] | human | Epidermidis-Aureus |
| *S. hominis* AIP-IV (**13**) | TINT-[CGGYF] | human | Epidermidis-Aureus |
| *S. hominis* AIP-V (**14**) | SQTV-[CSGYF] | human | Epidermidis-Aureus |
| *S. capitis* AIP-I (**15**) | GANP-[CALYY] | human | Epidermidis-Aureus |
| *S. haemolyticus* AIP-I (**16**) | SFTP-[CTTYF] | human | Epidermidis-Aureus |
| *S. warneri* AIP-I (**17**) | YSP-[CTNFF] | human | Epidermidis-Aureus |
| *S. warneri* AIP-II (**18**) | ANP-[CAMFY] | human | Epidermidis-Aureus |
| *S. cohnii* AIP-I (**19**) | TISSVKP-[CTGFV] | human | Saprophyticus |
| *S. saprophyticus* AIP-I (**20**) | INP-[CFGYT] | human | Saprophyticus |
| *S. caprae* AIP-I (**21**) | YST-[CSYYF] | animal | Epidermidis-Aureus |
| *S. pasteuri* AIP-I (**22**) | ANP-[CAGYF] | animal | Epidermidis-Aureus |
| *S. devriesei* AIP-I (**23**) | YKP-[CFGYF] | animal | Epidermidis-Aureus |
| *S. succinus* AIP-I (**24**) | ATAGALP-[CGGFF] | animal | Saprophyticus |
| *S. equorum* AIP-I (**25**) | AVAGARP-[CYGYF] | animal | Saprophyticus |
| *S. equorum* AIP-II (**26**) | AVAGLSP-[CGGYF] | animal | Saprophyticus |
| *S. hyicus* AIP-I (**27**) | KINP-[CTVFF] | animal | Hyicus-Intermedius |
| *S. chromogenes* AIP-I (**28**) | SINP-[CTGFF] | animal | Hyicus-Intermedius |
| *S. chromogenes* AIP-II (**29**) | AMDP-[CTGFF] | animal | Hyicus-Intermedius |
| *S. chromogenes* AIP-III (**30**) | SINP-[CTAFF] | animal | Hyicus-Intermedius |
| *S. schleiferi* AIP-I (**31**) | KYPF-[CIGYF] | animal | Hyicus-Intermedius |
| *S. intermedius* AIP-I (**32**) | RIPT-[STGFF] | animal | Hyicus-Intermedius |
| *S. simulans* AIP-I (**33**) | KYNP-[CLGFL] | animal | Hyicus-Intermedius |
| *S. simulans* AIP-II (**34**) | KYYP-[CWGYF] | animal | Hyicus-Intermedius |
| *S. simulans* AIP-III (**35**) | KYNP-[CWGYF] | animal | Hyicus-Intermedius |
| *S. vitulinus* AIP-I (**36**) | VIRG-[CTAFL] | animal | Sciuri |

#### List of recorded and reported quorum sensing interactions of staphylococcal AIPs

**Supplementary Table S2. $IC_{50}$ and $EC_{50}$ values (nM) measured in $\beta$-lactamase assay using *S. aureus* (SA) AgrC-I–IV reporter strains.**

| Species AIP | SA AgrC-I | SA AgrC-II | SA AgrC-III | SA AgrC-IV |
|---|---|---|---|---|
| *S. aureus* AIP-I (**1**) | **EC50:** 5.4 ± 0.5 | 700 ± 165 | 80 ± 11 | **EC50:** 790 ± 66 |
| *S. aureus* AIP-II (**2**) | 110 ± 12 | **EC50:** 3.7 ± 0.5 | 40 ± 15 | 230 ± 19 |
| *S. aureus* AIP-III (**3**) | 120 ± 15 | 150 ± 63 | **EC50:** 11 ± 1 | >1000 |
| *S. aureus* AIP-IV (**4**) | **EC50:** 59 ± 2 | 47 ± 9 nM | 9.8 ± 0.2[a] | **EC50:** 2.6 ± 0.2 |
| *S. epidermidis* AIP-I (**5**) | >1000 | -[b] | >1000 | -[b] |
| *S. epidermidis* AIP-II (**6**) | - | -[b] | -[b] | -[b] |
| *S. epidermidis* AIP-III (**7**) | >10000 | -[b] | >10000 | -[b] |
| *S. lugdunensis* AIP-I (**8**) | >1000 | -[b] | >1000 | -[b] |
| *S. lugdunensis* AIP-II (**9**) | >1000 | -[b] | -[b] | -[b] |
| *S. hominis* AIP-I (**10**) | 224 ± 37 | >1000 | 338 ± 83 | -[b] |
| *S. hominis* AIP-III (**12**) | 134 ± 8 | 580 ± 61 | 220 ± 46[a] | **EC50:** 54 ± 5 |
| *S. haemolyticus* AIP-I (**16**) | 340 ± 25 | -[b] | 340 ± 91 | - |
| *S. warneri* AIP-I (**17**) | 100 ± 10 | 1000 ± 105 | 460 ± 59 | >1000 |
| *S. cohnii* AIP-I (**19**) | >10000 | -[b] | -[b] | -[b] |
| *S. saprophyticus* AIP-I (**20**) | 360 ± 30 | - | -[b] | -[b] |
| *S. caprae* AIP-I (**21**) | 5.7 ± 0.8 | 252 ± 44 | -[a] | 199 ± 19 |
| *S. pasteuri* AIP-I (**22**) | 155 ± 34 | -[b] | 191 ± 18 | >1000 |
| *S. succinus* AIP-I (**24**) | 117 ± 26 | 854 ± 65 | >1000 | >1000 |
| *S. hyicus* AIP-I (**27**) | 3.3 ± 0.5 | 350 ± 90 | 4.0 ± 0.8 | 180 ± 33 |
| *S. chromogenes* AIP-I (**28**) | 15 ± 1 | 200 ± 13 | 60 ± 13 | 350 ± 85 |
| *S. schleiferi* AIP-I (**31**) | 2.8 ± 0.8 | 86 ± 6 | 80 ± 16[a] | **EC50:** 31 ± 6 |
| *S. intermedius* AIP-I (**32**) | 87 ± 20 | >1000 | 104 ± 20 | -[b] |
| *S. simulans* AIP-I (**33**) | 8.6 ± 0.5 | 23 ± 2 | 50 ± 11 | 280 ± 66 |
| *S. simulans* AIP-II (**34**) | 1.9 ± 0.3 | 80 ± 7 | 198 ± 33 | 273 ± 26 |
| *S. simulans* AIP-III (**35**) | 1.6 ± 0.1 | 60 ± 7 | 103 ± 5 | 351 ± 60 |
| *S. vitulinus* AIP-I (**36**) | 190 ± 15 | 800 ± 116 | 690 ± 46 | -[b] |
| *S. aureus* AIP-III D4A (**58**) | 3.3 ± 0.7 | 34 ± 1 | 1.6 ± 0.2[a] | 10 ± 0.2 |

All inhibition assays were performed in the presence of 100 nM cognate AIP and the results represent means ± standard error of the mean (SEM) of at least duplicate determinations performed in biological triplicate. [a]Only partial inhibition observed. [b]No inhibition recorded at the highest AIP concentration tested. Blue shaded boxes represent $IC_{50}$ values determined in this study. White represent $IC_{50}$/$EC_{50}$ values determined previously.[1]

**Supplementary Table S3. Reported QS interactions of staphylococcal AIPs against *S. aureus* (SA) *agr*-I–IV.**

| Species | SA *agr*-I | SA *agr*-II | SA *agr*-III | SA *agr*-IV | Assay type | Ref. |
|---|---|---|---|---|---|---|
| *S. aureus* AIP-I (**1**) | n. t. | **$IC_{50}$:** 3.4 ± 1.2 nM | **$IC_{50}$:** 3.1± 1.3 nM | n. t. | *β*-lactamase | 2 |
| | **$EC_{50}$:** 40 ± 9 nM | **$IC_{50}$:** 26 ± 7 nM | n. t. | n. t. | *β*-lactamase | 3 |
| | **$EC_{50}$:** 23 ± 7 nM | **$IC_{50}$:** 135 ± 52 nM | n. t. | n. t. | *β*-lactamase | 3 |
| | **$EC_{50}$:** 28 nM (18–46 CI) | **$IC_{50}$:** 25 nM (14–45 CI) | **$IC_{50}$:** 3 nM (2–5 CI) | **$EC_{50}$:** 26 μM (23–29 CI) | *β*-lactamase | 4 |
| | n. v. | **$IC_{50}$:** 8.00 nM (3.7–17.5 CI) | **$IC_{50}$:** 0.53 nM (0.31–0.88 CI) | n. v. | fluorescence | 5 |
| | n. t. | **$IC_{50}$:** 3.34 nM (0.67–16.5 CI) | **$IC_{50}$:** 6.12 nM (5.20–7.21 CI) | n. t. | hemolysis | 5 |
| | **$EC_{50}$:** 11 nM (9–12 CI) | n. t. | n. t. | n. t. | *β*-lactamase | 6 |
| | **$EC_{50}$:** 3.21 nM (1.21–8.57 CI) | n. t. | n. t. | n. t. | *β*-lactamase | 7 |
| *S. aureus* AIP-II (**2**) | **$IC_{50}$:** 2.9 ± 1.2 nM | n. t. | **$IC_{50}$:** 3.2 ± 1.3 nM | n. t. | *β*-lactamase | 2 |
| | **$IC_{50}$:** 90 ± 30 nM | **$EC_{50}$:** 34 ± 6 nM | n. t. | n. t. | *β*-lactamase | 3 |
| | **$IC_{50}$:** 78 ± 12 nM | **$EC_{50}$:** 28 ± 14 nM | n. t. | n. t. | *β*-lactamase | 3 |
| | **$IC_{50}$:** 40 nM (12–140 CI) | **$EC_{50}$:** 30 nM (10–90 CI) | **$IC_{50}$:** 1 nM (0.7–2.6 CI) | **$IC_{50}$:** 86 nM (65–111 CI) | *β*-lactamase | 4 |
| | **$IC_{50}$:** 1.62 nM (0.93–2.82 CI) | n. v. | **$IC_{50}$:** 0.53 nM (0.24–1.19 CI) | **$IC_{50}$:** 0.40 nM (0.21–0.76 CI) | fluorescence | 5 |
| | **$IC_{50}$:** 0.89 nM (0.37–2.15 CI) | n. t. | **$IC_{50}$:** 3.59 nM (1.22–10.6 CI) | **$IC_{50}$:** 1.19 nM (0.55–2.60 CI) | hemolysis | 5 |
| | n. t. | **$EC_{50}$:** 40.9 nM (30.3–55.3 CI) | n. t. | n. t. | *β*-lactamase | 7 |
| | **$IC_{50}$:** 12 ± 2.9 nM | n. t. | n. t. | n. t. | *β*-lactamase | 8 |
| *S. aureus* AIP-III (**3**) | **$IC_{50}$:** 70 nM (30–150 CI) | **$IC_{50}$:** 6 nM (5–6.5 CI) | **$EC_{50}$:** 26 nM (22–31 CI) | **$IC_{50}$:** 150 nM (104–207 CI) | *β*-lactamase | 4 |
| | **$IC_{50}$:** 5.05 nM (2.46–10.4 CI) | **$IC_{50}$:** 5.63 nM (1.89–16.7 CI) | n. v. | **$IC_{50}$:** 8.53 nM (4.15–17.5 CI) | fluorescence | 5 |
| | **$IC_{50}$:** 8.07 nM (4.34–15.0 CI) | **$IC_{50}$:** 0.46 nM (0.27–0.77 CI) | n. t. | **$IC_{50}$:** 23.8 nM (13.7–41.3 CI) | hemolysis | 5 |
| | n. t. | n. t. | **$EC_{50}$:** 406 nM (281–586 CI) | n. t. | *β*-lactamase | 7 |
| | **$IC_{50}$:** 8 ± 1.1 nM | n. t. | n. t. | n. t. | *β*-lactamase | 8 |
| *S. aureus* AIP-IV (**4**) | **$EC_{50}$:** 62 nM (52–75 CI) | **$IC_{50}$:** 4 nM (3–5 CI) | **IC50:** 1 nM (0.5–3 CI) | **$EC_{50}$:** 13 nM (7–40 CI) | *β*-lactamase | 4 |
| | n. v. | **$IC_{50}$:** 0.37 nM (0.22–0.64 CI) | **IC50:** 0.46 nM (0.21–0.99 CI) | n. v. | fluorescence | 5 |

| | | | | | | |
|---|---|---|---|---|---|---|
| | n. t. | **IC$_{50}$:** 0.090 nM (0.078–0.103 CI) | **IC50:** 1.49 nM (0.70–3.14 CI) | n. t. | hemolysis | 5 |
| | n. t. | n. t. | n. t. | **EC$_{50}$:** 7.90 nM (4.93–12.7 CI) | *β*-lactamase | 7 |
| *S. epidermidis* AIP-I (**5**) | **IC$_{50}$:** ~250 nM | **IC$_{50}$:** ~30-40 nM | **IC$_{50}$:** ~10 nM | no inhibition at 1000 nM | HPLC δ-toxin | 9 |
| | **IC$_{50}$:** 166 nM (68.7–402 CI) | **IC$_{50}$:** >1000 nM | **IC$_{50}$:** 13.0 nM (6.41–26.5 CI) | **IC$_{50}$:** >1000 nM | fluorescence | 10 |
| *S. epidermidis* AIP-II (**6**) | no inhibition at 100 nM | no inhibition at 100 nM | no inhibition at 100 nM | no inhibition at 100 nM | fluorescence | 11 |
| *S. lugdunensis* AIP-I (**8**) | **IC$_{50}$:** 384 nM (353–418 CI) | **IC$_{50}$:** 419 nM (353–418 CI) | **IC$_{50}$:** 36.6 nM (353–418 CI) | **IC$_{50}$:** >1000 nM | fluorescence | 10 |
| *S. hominis* AIP-I (**10**) | **IC$_{50}$:** 0.6243 nM | ~70% inhibition by supernatant | ~80% inhibition by supernatant | no inhibition by supernatant | fluorescence | 12 |
| | **IC$_{50}$:** 13 nM (11.0–15.2 CI) | **IC$_{50}$:** 31 nM (25.4–39.0 CI) | **IC$_{50}$:** 5 nM (4.2–6.0 CI) | **IC$_{50}$:** 2910 nM (2155–4543 CI) | fluorescence | 13 |
| *S. hominis* AIP-II (**11**) | **IC$_{50}$:** 15 nM (13.5–16.8 CI) | **IC$_{50}$:** 2109 nM (1504–3473 CI) | **IC$_{50}$:** 3 nM (3.0–3.8 CI) | **IC$_{50}$:** 1130 nM (882.2–1474 CI) | fluorescence | 13 |
| *S. hominis* AIP-III (**12**) | **IC$_{50}$:** 11 nM (8.8–13.0 CI) | **IC$_{50}$:** 4 nM (2.7–6.8 CI) | **IC$_{50}$:** 6 nM (5.3–7.7 CI) | NA | fluorescence | 13 |
| *S. hominis* AIP-IV (**13**) | **IC$_{50}$:** 128 nM (103.5–157.4 CI) | **IC$_{50}$:** 140 nM (109.8–178.0 CI) | **IC$_{50}$:** 37 nM (30.0–46.4 CI) | NA | fluorescence | 13 |
| *S. hominis* AIP-V (**14**) | **IC$_{50}$:** 43 nM (36.8–51.3 CI) | **IC$_{50}$:** 59 nM (50.2–69.3 CI) | **IC$_{50}$:** 4 nM (3.3–5.0 CI) | **IC$_{50}$:** 3809 nM (3199–7372 CI) | fluorescence | 13 |
| *S. warneri* AIP-I (**17**) | **IC$_{50}$:** 10 nM (9.1–11.7 CI) | **IC$_{50}$:** 4 nM (3.9–4.4 CI) | **IC$_{50}$:** 13 nM (11.4–14.0 CI) | **IC$_{50}$:** 146 nM (122.0–188.0 CI) | fluorescence | 14 |
| *S. warneri* AIP-II (**18**) | **IC$_{50}$:** 2 nM (1.6–2.5 CI) | **IC$_{50}$:** 30 nM (22.4–39.1 CI) | **IC$_{50}$:** 2 nM (1.5–1.9 CI) | **IC$_{50}$:** 2 nM (1.4–2.5 CI) | fluorescence | 14 |
| *S. caprae* AIP-I (**21**) | **IC$_{50}$0:** 0.6 ± 0.02 nM | **IC$_{50}$:** 0.26 ± 0.08 nM | **IC$_{50}$:** 0.2 nM | **IC$_{50}$:** 8.98 ± 0.32 nM | fluorescence | 15 |
| *S. intermedius* AIP-I (**32**) | 80–100% inhibition from supernatant | 40–90% inhibition from supernatant | 60–100% inhibition from supernatant | 10–60% inhibition from supernatant | *β*-lactamase | 16 |
| *S. simulans* AIP-I (**33**) | **IC$_{50}$:** 2.2 nM | **IC$_{50}$:** 1.1 nM | **IC$_{50}$:** 3.5 nM | **IC$_{50}$:** 23 nM | fluorescence | 17 |
| *S. simulans* AIP-II (**34**) | **IC$_{50}$:** 1.6 nM | **IC$_{50}$:** 15 nM | **IC$_{50}$:** 11.5 nM | **IC$_{50}$:** 40 nM | fluorescence | 17 |
| *S. simulans* AIP-III (**35**) | **IC$_{50}$:** 1.7 nM | **IC$_{50}$:** 6.0 nM | **IC$_{50}$:** 3.2 nM | **IC$_{50}$:** 48 nM | fluorescence | 17 |
| *S. aureus* AIP-III D4A (**58**) | **IC$_{50}$:** 0.49 nM (0.29–0.81 CI) | **IC$_{50}$:** 0.43 nM (0.21–0.89 CI) | **IC$_{50}$:** 0.051 nM (0.023–0.113CI) | **IC$_{50}$:** 0.035 nM (0.012–0.099 CI) | fluorescence | 5 |
| | **IC$_{50}$:** 0.082 nM (0.047–0.142 CI) | **IC$_{50}$:** 0.060 nM (0.038–0.094 CI) | **IC$_{50}$:** 0.16 nM (0.069–0.39 CI) | **IC$_{50}$:** 0.11 nM (0.066–0.17 CI) | hemolysis | 5 |
| | **IC$_{50}$:** 0.16 ± 0.01 nM | n. t. | n. t. | n. t. | *β*-lactamase | 8 |

**Supplementary Table S4. Reported QS interactions of staphylococcal AIPs against *S. epidermidis* (SE) *agr*-I–III.**

| Species | SE *agr*-I | SE *agr*-II | SE *agr*-III | Assay type | Ref. |
|---|---|---|---|---|---|
| *S. aureus* AIP-I (**1**) | no inhibition at 1000 nM | n. t. | n. t. | HPLC δ-toxin | 9 |
| | no inhibition at 1000 nM | n. t. | n. t. | fluorescence | 11 |
| *S. aureus* AIP-II (**2**) | no inhibition at 1000 nM | n. t. | n. t. | HPLC δ-toxin | 9 |
| | **$IC_{50}$:** 62.9 nM (26.1–151 CI) | n. t. | n. t. | fluorescence | 11 |
| *S. aureus* AIP-III (**3**) | no inhibition at 1000 nM | n. t. | n. t. | HPLC δ-toxin | 9 |
| | no inhibition at 1000 nM | n. t. | n. t. | fluorescence | 11 |
| *S. aureus* AIP-IV (**4**) | 40% inhibition at 1000 nM | n. t. | n. t. | HPLC δ-toxin | 9 |
| | **$IC_{50}$:** >1000 nM | n. t. | n. t. | fluorescence | 11 |
| *S. epidermidis* AIP-I (**5**) | activation from supernatant | ~90% inhibition from supernatant | ~90% inhibition from supernatant | fluorescence | 18 |
| | **$EC_{50}$:** 196 nM (162–238 CI) | n. t. | n. t. | fluorescence | 11 |
| *S. epidermidis* AIP-II (**6**) | ~90% inhibition from supernatant | activation from supernatant | no effect from supernatant | fluorescence | 18 |
| | **$IC_{50}$:** 9.64 nM (7.99–11.6 CI) | activation at 10 µM | no effect at 10 µM | fluorescence | 11 |
| *S. epidermidis* AIP-III (**7**) | ~50% inhibition from supernatant | no effect from supernatant | activation from supernatant | fluorescence | 18 |
| | **$IC_{50}$:** 34.3 nM (31.4–37.4 CI) | n. t. | n. t. | fluorescence | 11 |
| *S. hominis* AIP-I (**10**) | NA | **$IC_{50}$:** 34 nM (27.3–41.6 CI) | **$IC_{50}$:** 16 nM (14.5–18.5 CI) | fluorescence | 13 |
| *S. hominis* AIP-II (**11**) | **$IC_{50}$:** 20 nM (18.6–21.5 CI) | **$IC_{50}$:** 19 nM (16.9–21.2 CI) | **$IC_{50}$:** 62 nM (51.3–74.4 CI) | fluorescence | 13 |
| *S. hominis* AIP-III (**12**) | **$IC_{50}$:** 4 nM (2.5–7.3 CI) | **$IC_{50}$:** 3 nM (2.6–4.5 CI) | **$IC_{50}$:** 3 nM (2.1–5.6 CI) | fluorescence | 13 |
| *S. hominis* AIP-IV (**13**) | **$IC_{50}$:** 237 nM (199.3–281.1 CI) | **$IC_{50}$:** 93 nM (84.8–102.3 CI) | **$IC_{50}$:** 28 nM (26.1–29.6 CI) | fluorescence | 13 |
| *S. hominis* AIP-V (**14**) | **$IC_{50}$:** 10 nM (8.8–10.7 CI) | **$IC_{50}$:** 22 nM (17.9–27.8 CI) | **$IC_{50}$:** 2 nM (1.5–2.2 CI) | fluorescence | 13 |
| *S. warneri* AIP-I (**17**) | **$IC_{50}$:** 3 nM (2.8–3.4 CI) | **$IC_{50}$:** 19 nM (16.8–21.2 CI) | n. t. | fluorescence | 14 |
| *S. warneri* AIP-II (**18**) | **$IC_{50}$:** 12 nM (8.6–17.3 CI) | **$IC_{50}$:** 4 nM (3.2–4.1 CI) | n. t. | fluorescence | 14 |
| *S. aureus* AIP-III D4A (**58**) | ~25% inhibition at 10 µM | ~70% inhibition at 100 nM | ~25% inhibition at 10 µM | fluorescence | 11 |

#### MRSA mouse skin infection model data

**Supplementary Table S5. Skin lesion size, colony-forming unit (CFU) values and body weight for MRSA mouse skin infection model.**

| Treatment | Mouse | Skin lesion in mm$^2$ | | | $log_{10}$ CFU | | | Body weight in g | | |
|---|---|---|---|---|---|---|---|---|---|---|
| | | Day 1 | Day 2 | Day 4 | Day 1 | Day 2 | Day 4 | Day 1 | Day 2 | Day 4 |
| vehicle | 1 | 71.0 | - | - | 7.68 | - | - | 18.6 | - | - |
| | 2 | 76.2 | - | - | 7.89 | - | - | 21.4 | - | - |
| | 3 | 62.3 | - | - | 7.98 | - | - | 20.8 | - | - |
| | 4 | 69.9 | - | - | 8.11 | - | - | 19.6 | - | - |
| | 5 | 88.1 | - | - | 8.06 | - | - | 19.1 | - | - |
| | 6 | 69.7 | - | - | 7.95 | - | - | 18.9 | - | - |
| | 7 | 74.3 | - | - | 7.74 | - | - | 18.0 | - | - |
| | 8 | 79.4 | - | - | 7.68 | - | - | 18.6 | - | - |
| | 9 | 63.4 | 98.3 | - | - | 7.38 | - | 18.6 | 20.6 | - |
| | 10 | 70.5 | 99.3 | - | - | 7.89 | - | 20.8 | 20.4 | - |
| | 11 | 61.8 | 70.3 | - | - | 7.49 | - | 18.7 | 20.4 | - |
| | 12 | 52.8 | 101.3 | - | - | 7.56 | - | 20.4 | 19.0 | - |
| | 13 | 82.2 | 103.7 | 64.4 | - | - | 7.11 | 19.3 | 19.9 | 19.6 |
| | 14 | 73.4 | 96.5 | 58.8 | - | - | 7.76 | 19.9 | 18.8 | 18.6 |
| | 15 | 60.4 | 96.3 | 74.7 | - | - | 7.24 | 20.5 | 18.6 | 18.6 |
| | 16 | 71.0 | 97.5 | 58.2 | - | - | 6.19 | 19.4 | 20.9 | 21.2 |
| 2% fusidic acid ointment | 1 | 64.3 | 76.7 | - | - | 7.27 | - | 18.8 | 18.3 | - |
| | 2 | 82.5 | 98.4 | - | - | 6.89 | - | 18.8 | 20.5 | - |
| | 3 | 69.1 | 69.7 | - | - | 7.18 | - | 18.9 | 20.7 | - |
| | 4 | 47.4 | 66.4 | - | - | 7.38 | - | 20.4 | 19.1 | - |
| | 5 | 55.2 | 80.5 | 38.2 | - | - | 4.12 | 19.2 | 18.9 | 19.5 |
| | 6 | 74.9 | 81.4 | 53.9 | - | - | 5.70 | 19.1 | 20.2 | 21.4 |
| | 7 | 69 | 88.1 | 41 | - | - | 6.12 | 19.9 | 18.8 | 19.0 |
| | 8 | 63.3 | 70.1 | 56.5 | - | - | 6.04 | 19.3 | 19.6 | 19.8 |
| *S. simulans* AIP-II (**34**) | 1 | 75.5 | 84.7 | - | - | 7.88 | - | 19.1 | 20.0 | - |
| | 2 | 77.6 | 91.5 | - | - | 7.95 | - | 20.7 | 19.3 | - |
| | 3 | 111.4 | 81.3 | - | - | 8.08 | - | 19.0 | 20.2 | - |
| | 4 | 79.6 | 81.5 | - | - | 7.51 | - | 18.5 | 20.7 | - |
| | 5 | 63.3 | 94.1 | 47.7 | - | - | 5.68 | 19.5 | 21.7 | 22.0 |
| | 6 | 64.6 | 77.4 | 41.9 | - | - | 5.19 | 20.3 | 19.3 | 19.8 |
| | 7 | 51 | 61 | 50 | - | - | 5.30 | 21.3 | 19.1 | 19.4 |
| | 8 | 64.5 | 69 | 52.6 | - | - | 5.04 | 19.7 | 18.6 | 18.8 |

#### Bacterial strains and *agrD* sequencing

**Supplementary Table S6. Bacterial strains used in this study.**

| Name | Characterisitics | Reference |
|---|---|---|
| P3-*blaZ*/*pagrC*-I | *S. aureus* RN10829 reporter expressing AgrC-I | 1 |
| P3-*blaZ*/*pagrC*-II | *S. aureus* RN10829 reporter expressing AgrC-II | 1 |
| P3-*blaZ*/*pagrC*-III | *S. aureus* RN10829 reporter expressing AgrC-III | 1 |
| P3-*blaZ*/*pagrC*-IV | *S. aureus* RN10829 reporter expressing AgrC-IV | 1 |
| AH3408 | *S. epidermidis* ATCC1228(AH1740) + pCM40 (ermR, arP3_sGFP, type-I) | 18 |
| AH3623 | *S. epidermidis* 1457(AH1738) ica::DHFR + pCM40 (ermR, arP3_sGFP, type-II) | 18 |
| AH3409 | *S. epidermidis* Fey 8247+ pCM40 (ermR, arP3_sGFP, type-III) | 18 |
| AH4031 | *S. lugdunensis* N920143+ pCM40 (ermR, arP3_sGFP, type-I) | 19 |
| AH1677 | *S. aureus* LAC/pDB59 (*agr*-I P3::YFP-reporter strain) | 20 |
| AH430 | *S. aureus* 502A/pDB59 (*agr*-II P3::YFP-reporter strain) | 20 |
| AH1747 | *S. aureus* MW2/pDB59 (*agr*-III P3::YFP-reporter strain) | 20 |
| AH1872 | *S. aureus* MNEV/pDB59 (*agr*-IV P3::YFP-reporter strain) | 20 |
| JE2/pALC1741 | *S. aureus* JE2/pALC1741 (*agr*-I spa::GFP-reporter strain) | This study |
| 5927201 | *S. pasteuri* human nose isolate | This study |
| 102 L88 | *S. succinus* horse isolate | This study |
| 2934 c-30869-41 | *S. cohnii* isolate from skim milk | This study |
| AW 1555 | *S. capitis* isolate from bovine milk | This study |
| AW 0837 | *S. devriesei* isolate from bovine milk | This study |
| KS 155 | *S. equorum* isolate from bovine milk | This study |
| DV 026 | *S. equorum* isolate from bovine milk | This study |

**Supplementary Table S7. Primer used for *agrD* sequencing.**

| Species | Forward primer | Reverse primer |
|---|---|---|
| *S. pasteuri* | ccattgttagataaaaacttacagcc | gaaaaggaaacgaatactctccattg |
| *S. cohnii* | ccgtctgaaataatgtataccttgtc | ttatgtctacaattagactacttaaccac |
| *S. succinus* | gtcaaagtagcaagccatgtag | cattagcaatcattggctttgttg |

**Supplementary Table S8. AgrD sequences used for AIP identification.**

| Species | AgrD sequence | |
|---|---|---|
| *S. pasteuri* | METLVNLFFKFFTSIMEFVGLVAGANPCAGYFDEPEVPDELTKLYE | sequenced in this study |
| *S. cohnii* | MNIFESILTIFAKFFTFIGTISSVKPCTGFVDEPEIPKELTDLYE | sequenced in this study |
| *S. succinus* | MTILESLLTLITKFFSVLGATAGALPCGGFFDEPEVPSEITKLHE | sequenced in this study |
| *S. equorum agr-I* | MHIFESIFSIIAKFFTVLGAVAGARPCYGYFDETEVPKEITELYE | WP_021339970.1 |
| *S. equorum agr-II* | MHIFESIFSLIAKFFSTLGAVAGLSPCGGYFDEPEVPKEITDLYE | WP_046466292.1 |
| *S. devriesei* | MMFITDLFFKFFAAILETLGNVAAYKPCFGYFDEAEVPEELTNLKR | WP_103165878.1 |
| *S. capitis* | MDALFNLVLKFFTIIFEFIGFVAGANPCALYYDEPEVPDELSKLYE | WP_049428106.1 |

### 4. Methods and protocols

#### 4.1 General information

##### Abbreviations of chemicals

2-(1*H*-Benzotriazol-1-yl)-1,1,3,3-tetramethyluronoium hexafluorophosphate (HBTU), 2-(1*H*-7-azabenzotriazol-1-yl)-1,1,3,3-tetramethyluronoium hexafluorphosphate (HATU), [ethyl cyano(hydroxyimino)acetato-$O^2$]tri-1-pyrrolidinylphosphonium hexafluorophosphate (PyOxim), trifluoroacetic acid (TFA), *N*,*N*-dimethylformamide (DMF), tetrahydrofuran (THF), *N*-methylmorpholine (NMM), hexafluoroisopropanol (HFIP), 1,4-dithiothreitol (DTT), tris(2-carboxyethyl)phosphine hydrochloride (TCEP·HCl), *N*-methyl-2- pyrrolidone (NMP), guanidine hydrochloride (Gdn·HCl), dimethyl sulfoxide (DMSO), 9-fluorenylmethyloxycarbonyl (Fmoc), 2,3-diaminopropionic acid (Dap), *N*,*N*'-diisopropylcarbodiimide (DIC), *N*-methylimidazole (NMI), *sec*-isoamyl mercaptan (SIT), *tert*-butyldimethylsilyl (TBDMS), 4-(methylamino)benzoic acid (MeDbz), *N*-acyl-benzimidazolinone (Nbz), 1,8-biazabicyclo[5.4.0]undec-7-ene (DBU), tetrabutylammonium fluoride (TBAF), chloramphenicol (CAM), erythromycin (ERM).

##### Reagents and materials

Bacteria were cultured in tryptic soy broth (TSB) medium and on tryptic soy agar (TSA) plates (both Oxoid) supplemented with 10 µg/ml CAM or ERM (Sigma-Aldrich) when appropriate. *N*α-Fmoc protected and α-*N*-Boc protected amino acids, HBTU and HATU were obtained from ChemImpex and PepChem. Aminomethyl ChemMatrix resin was obtained from PCAS BioMatrix. Aminomethyl PEGA resin, *N*,*N*-diisopropylethylamine, 4-nitrophenylchloroformate, maleic acid (qNMR grade) and nitrocefin were obtained from Sigma-Aldrich. All other chemicals used were obtained in the highest available purity from CombiBlocks or IrisBiotech. All solvents used were of analytical grade and purchased from Fisher Scientific. Manual solid-phase peptide synthesis was performed in polypropylene syringes equipped with fritted disks, purchased from Torviq.

##### Compound analysis and purification

Analytical ultra-performance liquid chromatography (UPLC) analyses were performed on a C18 Agilent InfinityLab Poroshell 120 column (2.7 µm, 100 × 3.0 mm) using an Agilent 1260 Infinity II series system equipped with a diode array UV detector. A gradient with eluent A (water–MeCN–TFA, 95:5:0.1, v/v/v) and eluent B (0.1% TFA in MeCN) rising linearly from 0 to 50% of B over 10.0 min at a flow rate of 1.2 mL $min^{-1}$ was applied to determine the purity of peptides ($\lambda = 215$ nm). UPLC-mass spectrometry (MS) analyses were performed on a Phenomenex Kinetex column (1.7 µm, 100 Å, 50 × 2.10 mm) using a Waters Acquity system. A gradient with eluent C (0.1% HCOOH in water) and eluent D (0.1% HCOOH in MeCN) rising linearly from 0 to 95% of D over 5.20 min at a flow rate of 0.6 mL $min^{-1}$ was applied to analyse reaction mixtures. Preparative high-performance liquid chromatography (HPLC) purification was performed on a C18 Phenomenex Luna column (5 µm, 100 Å, 250 × 20 mm) using an Agilent 1260 LC system equipped with a diode array ultraviolet detector. Various gradients with eluent A and eluent B at a flow rate of 20 mL $min^{-1}$ were applied for the purification. Fractions containing the purified target peptide were identified using UPLC-MS or matrix assisted laser desorption ionisation–time of flight mass spectrometry (MALDI-TOF MS). MALDI-TOF mass spectra were recorded with a Bruker microflex bench-top MALDI using a matrix of alpha-cyano-4-hydroxycinnamic acid or 2,5-dihydroxybenzoic acid in water/MeCN (1:1, v/v) containing 0.1% TFA. The observed *m/z* corresponded to the monoisotopic ions, unless otherwise stated. Nuclear magnetic resonance (NMR) spectra were recorded at 298 K using a Bruker Avance III HD ($^1$H NMR and $^{13}$C NMR recorded at 600 MHz and 150 MHz, respectively). Chemical shifts are reported in parts per million (ppm) relative to the deuterated solvent peak of DMSO-$d_6$ ($\delta_H$ = 2.50 ppm; $\delta_C$ = 39.52 ppm) or $CDCl_3$ ($\delta_H$ = 7.26 ppm; $\delta_C$ = 77.16 ppm) as internal standard.

**Concentration determination of DMSO stock solutions**

Concentrations of DMSO stock solutions for compounds were determined by quantitative NMR relative to the signal ($\delta_H$ = 6.24 ppm, 2H) of the internal standard maleic acid (qNMR grade).

**Plate reader for reporter strain assays**

For fluorescence-based reporter strain assays, fluorescence (for YFP: excitation 500 nm and emission 541 nm; for sGFP: excitation 479 nm and emission 520 nm; automatic gain adjustment) was measured along with optical density at $\lambda$ = 600 nm (OD600) using a BioTek Synergy H1 microplate reader with Gen5TM software.

**Statistical analysis**

All statistical analyses were performed using GraphPad Prism 10.4 software. *P* values were determined using one-way analysis of variance (ANOVA) and Dunnet's test. *P* values < 0.05 considered significant.

#### 4.2 NCL trapping of AIPs from bacterial supernatants

**Preparation of NCL trapping resins**

Amino PEGA resin (1.00 g, loading: 0.42 mmol/g, 0.42 mmol) was placed in a polypropylene syringe equipped with a fritted disk, swelled in DMF for 15 min and washed with DMF (5 × 1 min). Fmoc-Rink-amide linker (1.13 g, 2.10 mmol, 5.00 equiv), HATU (782 mg, 2.06 mmol, 4.90 equiv), and *i*-$Pr_2NEt$ (736 µL, 4.20 mmol, 10.0 equiv) were pre-incubated in DMF (10.0 mL) for 2 min and then added to the resin. After 2 h, the resin was washed with DMF (3 × 1 min), MeOH (3 × 1 min), and $CH_2Cl_2$ (3 × 1 min) and treated with a capping solution ($Ac_2O$–*i*-$Pr_2NEt$–$CH_2Cl_2$, 2:2:6, v/v/v, 10.0 mL). After 2 h, the resin was washed with DMF (3 × 1 min), MeOH (3 × 1 min), and $CH_2Cl_2$ (3 × 1 min). The resin was then treated with piperidine in DMF (1:4, v/v, 10.0 mL) (1 × 2 min, 1 × 20 min) and washed with DMF (3 × 1 min), MeOH (3 × 1 min), and $CH_2Cl_2$ (3 × 1 min). Fmoc-Cys(S*t*-Bu)-OH (272 mg, 0.63 mmol, 1.50 equiv) or Fmoc-Cys(SIT)-OH[21] (335 mg, 0.63 mmol, 1.50 equiv), HATU (240 mg, 0.63 mmol, 1.50 equiv), and *i*-$Pr_2NEt$ (220 µL, 1.26 mmol, 3.00 equiv) were pre-incubated in DMF (10.0 mL) for 2 min and then added to the resin. After 2 h, the resin was washed with DMF (3 × 1 min), MeOH (3 × 1 min), and $CH_2Cl_2$ (3 × 1 min) and dried under high vacuum for 16 h.

**NCL trapping of AIPs from bacterial supernatants**

Bacterial isolates were streaked on agar plates and grown over night at 37 °C. Single colonies were then inoculated in 50 mL TSB media overnight at 37 °C in an incubator at 200 rpm shaking. Overnight cultures were centrifuged at 8000 rpm at 4 °C and supernatants filtered through a sterile filter (0.22 µm) and stored at 4 °C for direct use or frozen and stored at -20 °C until use. *Resin preparation:* Fmoc-Cys(S*t*-Bu)-Rink-PEGA resin (50 mg) or Fmoc-Cys(SIT)-Rink-PEGA resin (50 mg) was placed in a 2.0 mL polypropylene syringe equipped with a fritted disk, swelled in DMF for 15 min and washed with DMF (5 × 1 min). The resin was treated with piperidine in DMF (1:4, v/v, 2.0 mL) (1 × 2 min, 1 × 20 min) and washed with DMF (5 × 1 min). The resin was then treated with a solution of *β*-mercaptoethanol in DMF (1:4, v/v, 2.0 mL) containing *N*-methyl morpholine (NMM) (0.1 M) or DL-dithiothreitol (DTT) in DMF (0.05:0.95, w/v, 2.0 mL) containing NMM (0.1 M) (3 × 10 min) and subsequently washed with DMF (3 × 1 min), MeOH (3 × 1 min) and $H_2O$ (3 × 1 min). *NCL trapping:* The sterile and filtered bacterial supernatant (~50 mL) was added to 50 mL centrifugal tube and the pH adjusted to pH = ~7.0 using aqueous NaOH (1.0 M). An aqueous tris(2-carboxyethyl)phosphine hydrochloride (TCEP) solution (1.0 mL, 0.5 M, pH = 7.0; final conc. = 10.0 mM) was added to the supernatant followed by the deprotected Cys-Rink-PEGA-resin and the centrifugal tube containing the trapping mixture was agitated at 37 °C overnight. The next day, the resin was separated from the supernatant through filtration using a 10 mL polypropylene syringe equipped with a fritted disk under suction and washed with DMF (3 × 1 min), $H_2O$ (3 × 1 min), and DMF (3 × 1 min). A solution of DTT in DMF (0.05:0.95, w/v, 2.0 mL) containing NMM (0.1 M) was added to the resin and the resin was agitated at 37 °C. After 30 min, the resin was washed with DMF (3 × 1 min), MeOH (3 × 1 min), and $CH_2Cl_2$ (3 × 1 min) and dried under suction for 15 min. The dried resin was treated with a cleavage cocktail (2.0 mL, TFA–MilliQ water, 97:3, v/v) for 2 h at room temperature. The peptide containing cleavage solution was removed from the resin, collected and the resin rinsed with neat TFA (1.0 mL). The combined TFA fractions were evaporated under $N_2$ stream to near dryness, redissolved in a solution of MeCN in $H_2O$ (100 µL, 1:1, v/v) and filtered (0.22 µm). *LC-MS analysis*: The filtered TFA cleavage solution was analysed using an Waters Acquity system equipped with a Phenomenex Kinetex column (1.7 µm, 100 Å, 50 × 2.10 mm) applying a gradient with eluent C (0.1% HCOOH in water) and eluent D (0.1% HCOOH in MeCN) rising linearly from 0 to 50% of D over 10.0 min at a flow rate of 0.6 mL min$^{1}$ and an injection volume of 40 µL. The total ion chromatograms (TIC) were analyzed by displaying extracted ion chromatograms (EIC) of *m/z* $[M+H]^+$ values of the possible linear peptides with an additional *C*-terminal cysteine and amide functionality based on the AgrD sequence.

#### 4.3 Assay procedures using bacterial reporter strains

**Fluorescence reporter assay for screening *agr*-interference of peptides**

Peptides were evaluated for the ability to interfere with *agr*-mediated quorum sensing in *S. aureus* (AH1677, AH430, AH1747 and AH1872 for *agr*-I–IV, respectively)[20] using reporter strains expressing yellow fluorescent protein (YFP) upon *agr* activation. Interference with *agr*-mediated quorum sensing in *S. epidermidis* (AH3408, AH3623 and AH3409 for *agr*-I–III, respectively)[18] and *S. lugdunensis* (AH4031 for *agr*-I)[19] was evaluated using reporter strains expressing superfolder green fluorescent protein (sGFP) upon *agr* activation. Overnight cultures of the reporter strains were grown in TSB medium containing chloramphenicol (CAM, 10 μg/mL for *S. aureus*) or erythromycin (ERM, 10 μg/mL of *S. epidermidis* and *S. lugdunensis*) and diluted 1:100 in fresh TSB medium containing the same antibiotic. Assays were performed in sterile black 96-well plates with clear bottom. All peptides were screened at concentrations of 1.0 μM and 50 nM and peptides showing at least 75% inhibition at 50 nM were screened further at 2.5 nM against the respective reporter strain, and similarly at 0.125 nM in the case of at least 75% inhibition at 2.5 nM. DMSO stock solutions of peptides (1 mM) were diluted in TSB media and added (15 μL) in technical triplicate to the 96-well plate followed by diluted bacterial overnight cultures (135 μL). Control wells for 100% *agr* activity were wells replacing the peptide solution with TSB medium (15 μL). Wells containing 150 μL TSB medium were used to measure background fluorescence. The 96-well plates were incubated in a humidified incubator at 37°C shaking at 1000 rpm for 22–24 h and fluorescence (for GFP: excitation 479 nm, emission 520 nm; for YFP: excitation 500 nm, emission 541 nm; automatic gain) and $OD_{600}$ values were subsequently measured using a plate-reader. Background fluorescence was subtracted from all wells and further normalized to the corresponding $OD_{600}$ value of the respective wells. Average fluorescence of control wells was used as relative measure for 100% activation of the *agr*-circuit and bar graphs were generated using GraphPad Prism 10.4 software. All assays were performed in at least biological duplicate.

**Fluorescence reporter assay for $IC_{50}$ determination**

Overnight cultures of the reporter strains were grown in TSB medium containing chloramphenicol (CAM, 10 μg/mL for *S. aureus*) or erythromycin (ERM, 10 μg/mL of *S. epidermidis* and *S. lugdunensis*) and diluted 1:100 in fresh TSB medium containing the same antibiotic. Assays were performed in sterile black 96-well plates with clear bottom. Peptide solutions (15 μL) in 1:5 serial dilutions from DMSO stock solutions (1 mM) in TSB medium were added to the 96-well plate in technical duplicate followed by diluted bacterial overnight cultures (135 μL). Control wells for 100% *agr* activity were wells replacing the peptide solution with TSB medium (15 μL). Wells containing 150 μL TSB medium were used to measure background fluorescence. The 96-well plates were incubated in a humidified incubator at 37°C shaking at 1000 rpm for 22–24 h and fluorescence (for GFP: excitation 479 nm, emission 520 nm; for YFP: excitation 500 nm, emission 541 nm; automatic gain) and $OD_{600}$ values were subsequently measured using a plate-reader. Background fluorescence was subtracted from all wells and further normalized to the corresponding $OD_{600}$ value of the respective wells. Average fluorescence of control wells were used as relative measure for 100% activation of the *agr*-circuit. Relative *agr* activity was plotted to obtain $IC_{50}$

values by non-linear regression with variable slope using GraphPad Prism 10.4 software. All assays were performed in biological triplicate.

***β*-Lactamase assay for $IC_{50}$ determination against *S. aureus agr*-I–IV**

The *β*-lactamase reporter strains RN10829 [(P2-*agrA*:P3-*blaZ*)],[22] with p*agrC*-I,[23] p*agrC*-II, p*agrC*-III or p*agrC*-IV[1] substituting the native *agr* locus with a chromosomal integration of P2-*agrA* and P3-*blaZ* and a plasmid from which a wild-type variant of the corresponding AgrC is expressed, were used to asses inhibition and activation of the AgrC receptor via *β*-lactamase activity in response to varying concentrations of the QS modulating peptides. Overnight cultures of the reporter strains in TSB medium were diluted 1:250 in fresh TSB medium and grown to $OD_{600}$ = 0.35–0.40 (early exponential phase) at 37 °C. Peptide solutions (10 µL) in 1:10 serial dilutions from DMSO stock solutions (1 mM) in TSB medium (final concentrations = 10 µM–10 pM) were added to each well of a clear 96-well plate as well as solutions (10 µL) of cognate AIP (final concentration = 100 nM) in TSB medium followed by 80 µL of bacterial cells. Control wells for 100% *β*-lactamase activity were wells replacing peptide solution with TSB media (10 µL) and control wells for 0% *β*-lactamase activity were wells replacing both peptide and cognate AIP solutions with TSB medium (20 µL). The 96-well plates were incubated at 37 °C shaking at 200 rpm for 1 h and immediately frozen down at -80 °C to minimize growth during nitrocefin treatment. Next, the 96-well plates were thawed and $OD_{600}$ values were determined using a plate-reader followed by addition of 50 µL of nitrocefin solution to the wells (final concentration = 33.3 µg/mL). *β*-Lactamase activity was monitored at $OD_{486}$ every 20 s for 10 min at 37 °C using a plate-reader. Linear nitrocefin conversion rates were plotted to obtain $IC_{50}$ values by non-linear regression with variable slope using GraphPad Prism 10.4 software. Assays were performed at least as duplicate determinations in biological triplicate.

**Fluorescence reporter assay for *agr* deactivation of virulent *S. aureus***

A fluorescent reporter strain of *S. aureus* JE2 containing a plasmid with a *spa::GFP* promotor was constructed by transforming the strain with pALC1741.[24] Cultures of the reporter strain were grown from single colonies in TSB medium containing chloramphenicol (CAM, 10 µg/mL) and *S. aureus* AIP-I (**1**) (concentrations = 500 nM, 100 nM or 50 nM) to $OD_{600}$ = 0.2–0.3 (early exponential phase) at 37 °C. Assays were performed in sterile black 96-well plates with clear bottom. Peptide solutions (15 µL) in 1:5 serial dilutions from DMSO stock solutions (1 mM) in TSB medium were added to the 96-well plate in technical triplicate followed by the induced bacterial cultures (135 µL). Control wells were uninduced cultures containing no *S. aureus* AIP-I (**1**). The 96-well plates were incubated in a plate reader at 37°C shaking at 283 rpm and fluorescence (GFP: excitation 479 nm, emission 520 nm, automatic gain) and $OD_{600}$ values were continuously measured. For $IC_{50}$ values, final fluorescence measurements were normalized to the corresponding $OD_{600}$ value of the respective wells and average fluorescence of control wells were used as relative measure for 100% GFP expression. Relative GFP expression was plotted to obtain $IC_{50}$ values by non-linear regression with variable slope using GraphPad Prism 10.4 software. All assays were performed in biological triplicate.

**Quorum sensing inhibition resistance development experiment**

Bacterial cultures (n = 3 per treatment) of a fluorescent reporter strain of *S. aureus* (AH1677) with a *P3::YFP* promoter plasmid were cultured in 15 mL centrifugal tubes in 2.0 mL of TSB medium containing chloramphenicol (CAM, 10 µg/mL) and *S. simulans* AIP-II (**34**) (concentrations = 100 nM, or 2.0 nM) or no peptide overnight at 37 °C in a shaking incubator. The next day, YFP expression levels were assessed by flow cytometry (YFP: excitation 500 nm, emission 541 nm) and the overnight cultures were diluted 1:250 in fresh TSB medium containing CAM and **34** or no peptide (final volume 2.0 mL) and incubated overnight at 37 °C in a shaking incubator. The procedure was repeated in total for 15 days and the passaged cultures were subsequently frozen at -80 °C. The passaged cultures were inoculated from frozen stocks in 2.0 mL TSB medium containing CAM in 15 mL centrifugal tubes and incubated overnight at 37 °C in a shaking incubator. Overnight cultures were diluted 1:250 in fresh TSB medium containing CAM and 1:5 serial dilutions of *S. simulans* AIP-II (**34**) (concentrations = 10 nM–0.08 nM) were added and the cultures incubated overnight at 37 °C in a shaking incubator. The next day, YFP expression levels were assessed by flow cytometry (YFP: excitation 500 nm, emission 541 nm) to determine the susceptibility towards *agr* inhibition by *S. simulans* AIP-II (**34**).

#### 4.4 *In vivo* MRSA skin infection model

The mouse model was performed under contract at Statens Serum Institut (DK) essentially as previously described. In brief, eight to ten-week-old Balb/c female mice (Taconic Denmark) mice were used for all experiments [n = 16 for vehicle, n = 8 for Fucidin® (2% fusidic acid ointment) treatment, and n = 8 for *S. simulans* AIP-II (**34**) treatment]. All animal experiments were approved by the National Committee of Animal Ethics, Denmark. Mice were anaesthetized and the hair was removed on a 2 $cm^2$ skin area on the back and thereafter was the outer most layer of the skin scraped off with a dermal curette to obtain a 1 $cm^2$ superficial skin lesion. For vehicle control and Fucidin® treatment, 10 µL inoculum containing approximately $10^7$ CFU of methilicin-resistant *S. aureus* (MRSA43484) were spread on the skin lesions. For *S. simulans* AIP-II (**34**) treatment, 10 µL of the same inoculum containing **34** (100 µM prepared from a 10 mM DMSO stock solution of **34**•HCl immediately before application) were spread on the skin lesions. After the applied inoculum had dried, the mice were placed in a cage and kept in a warming cabinet until fully awake. The topical skin treatment with Fucidin® was initiated one day after inoculation (day 2, 3, 4) by spreading 50 µL of Fucidin® on the inoculated skin area once a day. Skin lesion size and body weight was measured on day 1, 2, and 4 of all mice. Mice were sacrificed on day 1 (n = 8 vehicle), day 2 (n = 4 vehicle, Fucidin®, **34**) and day 4 (n = 4 vehicle, Fucidin®, **34**) and the infected skin area was cut out and homogenized to determine the CFU count in the skin lesions.

#### 4.5 Synthetic procedures

**General protocol for automated peptide synthesis**

Automated peptide synthesis was carried out on a Biotage SyroWave$^{TM}$ synthesizer using standard Fmoc SPPS chemistry. The following Fmoc-protected amino acids with side chain protecting groups were used: Fmoc-Ala-OH, Fmoc-Asn(Trt)-OH, Fmoc-Cys(Trt)-OH, Fmoc-Cys(*S*t-Bu)-OH, Fmoc-Dap(Alloc)-OH, Fmoc-Gly-OH, Fmoc-Ile-OH, Fmoc-Leu-OH, Fmoc-Phe-OH, Fmoc-Pro-OH, Fmoc-Ser(*t*-Bu)-OH, Fmoc-Ser(TBDMS)-OH, Fmoc-Thr(*t*-Bu)-OH, Fmoc-Trp(Boc)-OH, Fmoc-Tyr(*t*-Bu)-OH, and Fmoc-Val-OH. The following amino acids with side chain protecting groups were used as *N*-terminal amino acids: Ac-OH, Boc-Ala-OH, Boc-Asn(Trt)-OH, Boc-Asp(O*t*-Bu)-OH, Boc-Arg(Pbf)-OH, $Me_2N$-Cys(Trt)-OH,[25] Boc-Lys(Boc)-OH, Boc-Ser(*t*-Bu)-OH, Boc-Thr(*t*-Bu)-OH and Boc-Tyr(*t*-Bu)-OH.

SPPS was performed on 0.02 mmol (method A) or 0.04 mmol (method B) scale using MeDbz-Gly-ChemMatrix resin[8] or on 0.04 mmol scale using preloaded chlorotrityl (Cl-Trt) resin. Fmoc deprotection was performed in two stages: 1) piperidine in DMF (2:3, v/v) for 3 min and 2) piperidine in DMF (1:4, v/v) for 12 min. The deprotection was followed by washing with DMF (2 × 45 s), $CH_2Cl_2$ (1 × 45 s), and DMF (2 × 45 s). The first coupling reaction on MeDbz-Gly-ChemMatrix resin was performed as double coupling using Fmoc-AA-OH (5.00 equiv to the resin loading), HATU (4.90 equiv) and *i*-$Pr_2$NEt in NMP (10.0 equiv, 2.0 M) in DMF (final concentration = 0.2 M for Fmoc-AA-OH) for 90 min. Standard coupling reactions were performed as double couplings with Fmoc-AA-OH (5.00 equiv to the resin loading), HBTU (4.90 equiv) and *i*-$Pr_2$NEt in NMP (10.0 equiv, 2.0 M) in DMF (final concentration = 0.2 M for Fmoc-AA-OH) for 40 min for each coupling. The last amino acid was incorporated as Boc-AA-OH.

**General procedure for *N*-acyl-benzimidazolinone (Nbz) formation**

After automated peptide elongation, the peptidyl-MeDbz-Gly-ChemMatrix resin (1.00 equiv) was transferred into a polypropylene syringe equipped with a fritted disk using $CH_2Cl_2$ and the resin was then washed with $CH_2Cl_2$ (5 × 1 min). A solution of 4-nitrophenyl-chloroformate (5.00 equiv) in $CH_2Cl_2$ (concentration = 0.1 M) was added to the resin and the suspension was agitated for 30 min. The resin was then washed with $CH_2Cl_2$ (2 × 1 min) and the procedure was repeated. The resin was then washed with $CH_2Cl_2$ (3 × 1 min) and DMF (3 × 1 min) and a solution of *i*-$Pr_2$NEt (25.0 equiv) in DMF (0.5 M) was added to the resin. After 15 min, the resin was washed with DMF (3 × 1 min) and the procedure was repeated. The resin was then washed with DMF (3 × 1 min), *i*-$Pr_2$NEt in DMF (5%, v/v) (3 × 1 min), DMF (3 × 1 min), MeOH (3 × 1 min), and $CH_2Cl_2$ (3 × 1 min) and dried under high vacuum.

**Method A: General procedure for on-resin cleavage-inducing cyclization**

Dried peptidyl-MeNbz-Gly-ChemMatrix resin **S4** (0.02 mmol, 1.00 equiv) was placed in a polypropylene syringe equipped with a fritted disk and treated with a deprotection/cleavage cocktail (2.0 mL, TFA–*i*-$Pr_3SiH$–water, 94:3:3, v/v/v) for 1 h and the TFA cocktail removed from the resin. The resin **S5** was washed with $CH_2Cl_2$ (3 × 1 min), DMF (3 × 1 min), and $CH_2Cl_2$ (3 × 1 min) and dried under suction for several minutes. The cyclization buffer (5.0 mL, phosphate buffer (0.2 M, pH = 6.8)–MeCN, 1:1, v/v) was added to the resin (final concentration = 4.0 mM) and the suspension was agitated at 50 °C for 2 h. The solution was removed from the resin, collected and the resin was rinsed with fresh cyclisation buffer. The combined peptide-containing solution was purified by preparative RP-HPLC. Fractions containing pure peptide were lyophilized to afford the desired cyclized peptide.

**Method B: General procedure for solution cyclization**

The procedure was adapted from a previously published protocol.[26] Dried peptidyl-MeNbz-Gly-ChemMatrix resin **S4** (0.04 mmol, 1.00 equiv) was placed in a polypropylene syringe equipped with a fritted disk and treated with a solution of 3-mercaptopropionic acid ethyl ester (50.6 µL, 0.40 mmol, 10.0 equiv) and *i*-$Pr_2NEt$ (69.8 µL, 0.40 mmol, 10.0 equiv) in DMF (3.0 mL). After overnight incubation, the peptide containing solution was removed from the resin and the resin washed twice with DMF (2.0 mL). The combined organic phase was concentrated to dryness under pressure and the remaining residue was treated with a deprotection cocktail (2.0 mL, TFA–*i*-$Pr_3SiH$–water, 94:3:3, v/v/v) for 2 h. The reaction mixture was concentrated under a stream of nitrogen and precipitated by addition of ice-cold diethyl ether. The crude peptide thioester **S6** was lyophilized and used in the next step without further purification. Crude **S6** (10.0 µmol) was dissolved in cyclization buffer (10.0 mL, guanidinium hydrochloride (6 M) in phosphate buffer (0.1 M, pH = 7.0)–MeCN, 4:1, v/v) with a final concentration of 1.0 mM and the reaction mixture was agitated at 37 °C. After for 2 h, the reaction mixture was purified by preparative RP-HPLC. Fractions containing pure peptide were lyophilized to afford the desired cyclized peptide.

### 5. Synthetic peptides

#### 5.1 Native AIPs

The following peptides have been characterized elsewhere: *S. aureus* AIP-I–IV (**1**–**4**),[8] *S. lugdunensis* AIP-II (**8**), *S. hominis* AIP-III (**12**), *S. haemolyticus* AIP-I (**16**), *S. warneri* AIP-I (**17**), *S. sparophyticus* AIP-I (**20**), *S. hyicus* AIP-I (**27**), *S. chromogenes* AIP-I–III (**28**–**30**), *S. schleiferi* AIP-I (**31**), *S. simulans* AIP-I (**33**) and *S. vitulinus* AIP-I (**36**).[1]

***S. epidermidis*** **AIP-I (5)**

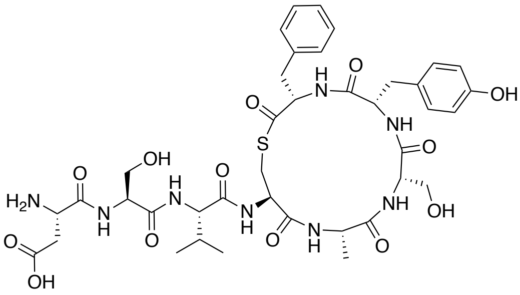

$C_{39}H_{52}N_8O_{13}S$
MW: 872.95

The peptide was synthesized according to general method B for solution cyclization (section 4.5). Purification by preparative RP-HPLC afforded *S. epidermidis* AIP-I (**5**) as a fluffy white solid after lyophilization. Purity >95% determined by UPLC ($\lambda$ = 215 nm). **UPLC-MS** (ESI) *m/z* calcd for $[M+H]^+$ $C_{39}H_{53}N_8O_{13}S^+$: 873.35, found 873.32. **$^1$H NMR** (600 MHz, DMSO-$d_6$) $\delta$ = 0.83 (t, $J$ = 7.1 Hz, 6H), 1.21 (d, $J$ = 7.3 Hz, 3H), 1.95–2.04 (m, 1H), 2.58–2.67 (m, 2H), 2.67–2.72 (m, 1H), 2.73–2.82 (m, 2H), 2.90–2.95 (m, 1H), 3.16 (dd, $J$ = 12.8, 3.8 Hz, 1H), 3.28 (dd, $J$ = 14.0, 3.9 Hz, 1H), 3.54–3.66 (m, 4H), 4.05–4.07 (m, 2H), 4.15–4.23 (m, 3H), 4.24–4.30 (m, 2H), 4.41–4.46 (m, 1H), 5.04–5.11 (m, 1H), 5.13–5.20 (m, 1H), 6.62 (d, $J$ = 8.5 Hz, 2H), 6.85 (d, $J$ = 8.5 Hz, 2H), 7.01–7.04 (m, 2H), 7.16–7.20 (m, 1H), 7.21–7.24 (m, 2H), 7.83 (d, $J$ = 8.7 Hz, 1H), 7.91 (d, $J$ = 8.8 Hz, 1H), 8.04 (d, $J$ = 7.5 Hz, 1H), 8.21 (d, $J$ = 8.3 Hz, 1H), 8.45 (d, $J$ = 7.8 Hz, 1H), 8.58 (d, $J$ = 7.6 Hz, 1H), 8.88 (d, $J$ = 8.2 Hz, 1H), 9.22 (s, 1H).

***S. epidermidis*** **AIP-II (6)**

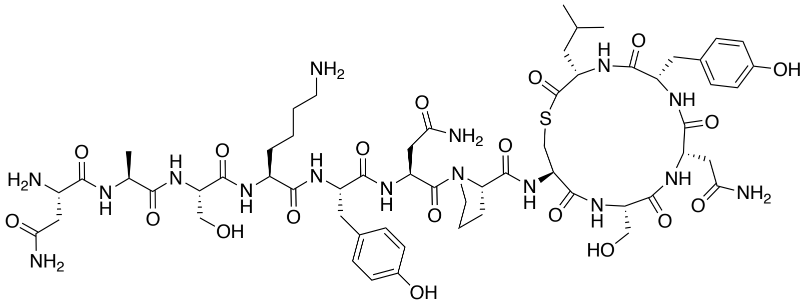

$C_{59}H_{86}N_{16}O_{19}S$
MW: 1355.49

The peptide was synthesized according to general method A for on-resin cleavage-inducing cyclization (section 4.5). Purification by preparative RP-HPLC afforded *S. epidermidis* AIP-II (**6**) as a fluffy white solid after lyophilization. Purity >95% determined by UPLC ($\lambda$ = 215 nm). **UPLC-MS** (ESI) *m/z* calcd for $[M+2H]^{2+}$ $C_{59}H_{88}N_{16}O_{19}S^{2+}$: 678.31, found 678.61; $[M+H]^+$ $C_{59}H_{87}N_{16}O_{19}S^+$: 1355.61, found 1355.73. **$^1$H NMR** (600 MHz, DMSO-$d_6$) $\delta$ = 0.75 (d, $J$ = 6.5 Hz, 3H), 0.83 (d, $J$ = 6.6 Hz, 3H), 1.18–1.27 (m, 5H), 1.29–1.37 (m, 1H), 1.41–1.53 (m, 3H), 1.54–1.64 (m, 2H), 1.64–1.71 (m, 1H), 1.80–1.89 (m, 3H), 1.93–2.01 (m, 1H), 2.37 (dd, $J$ = 15.6, 6.3 Hz, 1H), 2.51–2.65 (m, 5H), 2.66–2.76 (m, 3H), 2.78–2.90 (m, 3H), 2.94 (dd, $J$ = 13.7, 7.6 Hz, 1H), 3.13 (dd, $J$ = 12.9,

4.1 Hz, 1H), 3.49 (dd, $J$ = 11.2, 4.4 Hz, 1H), 3.52–3.66 (m, 5H), 4.05–4.15 (m, 2H), 4.14–4.23 (m, 3H), 4.27–4.35 (m, 3H), 4.36–4.46 (m, 3H), 4.77 (q, $J$ = 7.2 Hz, 1H), 4.92 (s, 1H), 5.16 (s, 1H), 6.62 (d, $J$ = 8.3 Hz, 2H), 6.65 (d, $J$ = 8.3 Hz, 2H), 6.92 (s, 1H), 6.93–6.97 (m, 4H), 7.01 (s, 1H), 7.26 (s, 1H), 7.47 (s, 1H), 7.51 (s, 1H), 7.62–7.75 (m, 4H), 7.84 (d, $J$ = 8.0 Hz, 1H), 7.90–7.96 (m, 2H), 8.04 (d, $J$ = 8.9 Hz, 1H), 8.06–8.12 (m, 4H), 8.15 (d, $J$ = 7.7 Hz, 1H), 8.19 (d, $J$ = 8.0 Hz, 1H), 8.27 (d, $J$ = 7.7 Hz, 1H), 8.56 (d, $J$ = 8.4 Hz, 1H), 8.60 (d, $J$ = 7.4 Hz, 1H), 9.20 (s, 2H).

***S. epidermidis*** **AIP-III (7)**

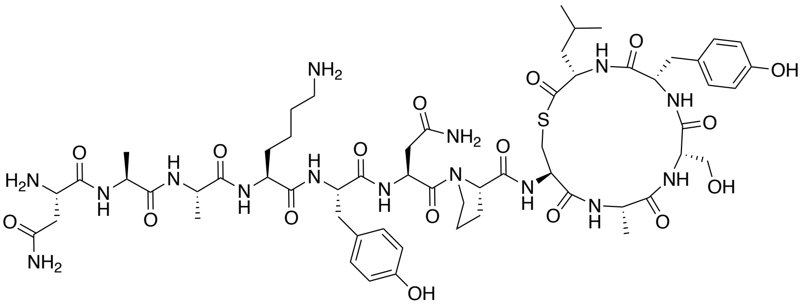

$C_{58}H_{85}N_{15}O_{17}S$
MW: 1296.47

The peptide was synthesized according to general method A for on-resin cleavage-inducing cyclization (section 4.5). Purification by preparative RP-HPLC afforded *S. epidermidis* AIP-III (**7**) as a fluffy white solid after lyophilization. Purity 95% determined by UPLC ($\lambda$ = 215 nm). **UPLC-MS** (ESI) *m/z* calcd for $[M+2H]^{2+}$ $C_{58}H_{87}N_{15}O_{17}S^{2+}$: 648.81, found 649.15; $[M+H]^{+}$ $C_{58}H_{86}N_{15}O_{17}S^{+}$: 1296.60, found 1296.72. **$^1$H NMR** (600 MHz, DMSO-$d_6$) $\delta$ = 0.75 (d, $J$ = 6.5 Hz, 3H), 0.82 (d, $J$ = 6.6 Hz, 3H), 1.16–1.26 (m, 11H), 1.26–1.33 (m, 1H), 1.42–1.53 (m, 3H), 1.53–1.59 (m, 3H), 1.80–1.88 (m, 3H), 1.96–2.04 (m, 1H), 2.37 (dd, $J$ = 15.5, 6.2 Hz, 1H), 2.54–2.71 (m, 4H), 2.71–2.78 (m, 2H), 2.80–2.89 (m, 3H), 2.90–2.95 (m, 1H), 3.11 (dd, $J$ = 12.9, 3.8 Hz, 1H), 3.53–3.64 (m, 4H), 4.04–4.10 (m, 2H), 4.13–4.21 (m, 4H), 4.24 (t, $J$ = 7.2 Hz, 1H), 4.25–4.30 (m, 1H), 4.30–4.36 (m, 2H), 4.44 (td, $J$ = 7.9, 5.3 Hz, 1H), 4.77 (q, $J$ = 7.3 Hz, 1H), 5.17 (s, 1H), 6.60 (d, $J$ = 8.5 Hz, 2H), 6.64 (d, $J$ = 8.5 Hz, 2H), 6.91–6.95 (m, 3H), 6.97 (d, $J$ = 8.5 Hz, 2H), 7.26 (s, 1H), 7.49 (s, 1H), 7.66–7.75 (m, 5H), 7.77 (d, $J$ = 8.7 Hz, 1H), 7.82 (d, $J$ = 8.0 Hz, 1H), 7.96 (d, $J$ = 7.6 Hz, 1H), 8.04 (d, $J$ = 7.9 Hz, 1H), 8.08–8.11 (m, 3H), 8.13 (d, $J$ = 7.3 Hz, 1H), 8.18 (d, $J$ = 7.9 Hz, 1H), 8.38 (d, $J$ = 7.7 Hz, 1H), 8.59 (d, $J$ = 7.4 Hz, 1H), 8.68 (d, $J$ = 8.3 Hz, 1H), 9.14–9.27 (m, 2H).

***S. lugdunensis*** **AIP-I (8)**

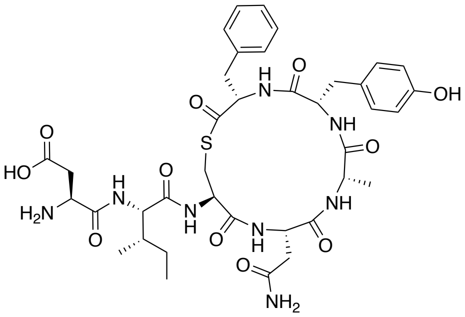

$C_{38}H_{50}N_8O_{11}S$
MW: 826.92

The peptide was synthesized as followed (Supporting Scheme S2). The fully protected linear peptide **S8** was synthesized in 40.0 μmol scale on Cl-Trt polystyrene resin **S7** preloaded with Fmoc-Phe-OH (0.69 mmol/g) using the general procedures for automated SPPS. After completed peptide elongation, the peptidyl resin **S8** was transferred into a polypropylene syringe equipped with a fritted disk using $CH_2Cl_2$ and washed with DMF (3 × 1 min). A solution of NMM (22.1 µL, 0.20 mmol, final concentration = 0.1 M) in a mixture of *β*-mercaptoethanol–DMF (2.0 mL, 1:4, v/v) was added to the peptidyl resin **S8** and the resin was agitated overnight at room temperature. The next day, the thiol-containing solution was removed by suction and the resin was washed with DMF (3 × 1 min), MeOH (3 × 1 min) and $CH_2Cl_2$ (3 × 1 min) and dried under suction for 15 min. The dried resin was treated with a solution of HFIP (0.4 mL) in $CH_2Cl_2$ (1.6 mL) for 15 min at room temperature to cleave the partially-protected peptide **S9** from the resin. The cleavage solution was removed from the resin, collected and a fresh HFIP–$CH_2Cl_2$ solution (2.0 mL, 1:4, v/v) was added to the resin. After 15 min, the cleavage solution was removed from the resin, collected and the resin rinsed with $CH_2Cl_2$ (2.0 mL). The combined cleavage solution was concentrated to dryness under reduced pressure to yield the partially protected peptide **S9**, which was used without further purification.

The crude peptide **S9** (0.04 mmol based on the resin loading) was dissolved in anhydrous DMF (5.0 mL) under nitrogen atmosphere and added dropwise to a solution of PyOxim (42.2 mg, 0.08 mmol, 2.00 equiv) and *i*-$Pr_2NEt$ (48.6 µL, 0.28 mmol, 7.00 equiv) in anhydrous DMF (35.0 mL). The reaction mixture was stirred overnight at room temperature and after full consumption of **S9** was confirmed by UPLC-MS, the reaction was reduced to dryness under reduced pressure. The remaining residue was treated with a deprotection cocktail (3.0 mL, TFA–*i*-$Pr_3SiH$–water, 94:3:3, v/v/v) for 2 h and subsequently concentrated under a stream of nitrogen followed by precipitation in ice-cold diethyl ether. Purification by preparative RP-HPLC afforded *S. lugdunensis* AIP-I (**8**) as a fluffy white solid after lyophilization. Purity >95% determined by UPLC ($\lambda$ = 215 nm). **UPLC-MS** (ESI) *m/z* calcd for $[M+H]^+$ $C_{38}H_{51}N_8O_{11}S^+$: 827.34, found 827.40. **$^1$H NMR** (600 MHz, DMSO-$d_6$) $\delta$ = 0.79–0.83 (m, 6H), 1.03–1.11 (m, 1H), 1.17 (d, *J* = 6.9 Hz, 3H), 1.37–1.47 (m, 1H), 1.65–1.73 (m, 1H), 2.30 (dd, *J* = 15.7, 5.2 Hz, 1H), 2.50–2.59 (m, 2H), 2.64 (dd, *J* = 13.6, 7.0 Hz, 1H), 2.70 (dd, *J* = 17.3, 3.8 Hz, 1H), 2.76–2.86 (m, 2H), 2.92 (t, *J* = 12.0 Hz, 1H), 3.15 (dd, *J* = 12.7, 4.0 Hz, 1H), 3.27 (1H) signal partially overlapping with HDO peak, 3.98–4.04 (m, 1H), 4.06 (q, *J* = 7.6 Hz, 1H), 4.12–4.20 (m, 1H), 4.21 (t, *J* = 7.2 Hz, 1H), 4.31 (td, *J* = 9.2, 4.1 Hz, 1H), 4.43 (dt, *J* = 8.4, 5.8 Hz, 1H), 4.48 (ddd, *J* = 11.8, 8.1, 4.0 Hz, 1H), 6.64 (d, *J* = 8.2 Hz, 2H), 6.87 (d, *J* = 8.2 Hz, 2H), 6.91 (s, 1H), 7.03 (d, *J* = 7.4 Hz, 2H), 7.15–7.19 (m, 1H), 7.19–7.25 (m, 2H), 7.40 (s, 1H), 7.86 (d, *J* = 8.7 Hz, 1H), 8.13 (d, *J* = 8.5 Hz, 1H), 8.25 (d, *J* = 7.7 Hz, 1H), 8.45 (d, *J* = 8.5 Hz, 1H), 8.59 (d, *J* = 8.1 Hz, 1H), 8.80 (d, *J* = 8.4 Hz, 1H), 9.23 (s, 1H).

***S. hominis* AIP-I (10)**

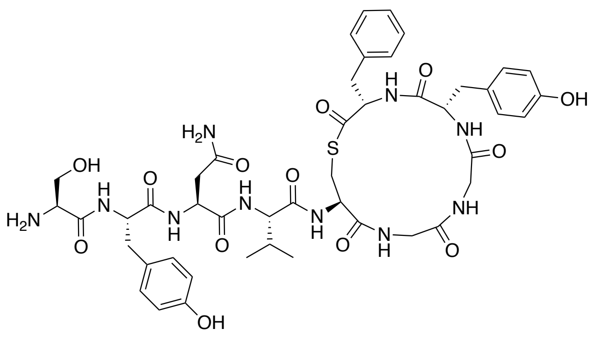

$C_{46}H_{58}N_{10}O_{13}S$
MW: 991.09

The peptide was synthesized according to general method A for on-resin cleavage-inducing cyclization (section 4.5). Purification by preparative RP-HPLC afforded *S. hominis* AIP-I (**10**) as a fluffy white solid after lyophilization. Purity >95% determined by UPLC ($\lambda$ = 215 nm). **UPLC-MS** (ESI) *m/z* calcd for $[M+H]^+$ $C_{46}H_{59}N_{10}O_{13}S^+$: 991.40, found 991.57. **$^1$H NMR** (600 MHz, DMSO-$d_6$) $\delta$ = 0.81 (d, *J* = 6.8 Hz, 3H), 0.83 (d, *J* = 6.8 Hz, 3H), 1.95–2.07 (m, 1H), 2.32 (dd, *J* = 14.3, 10.6 Hz, 1H), 2.42 (dd, *J* = 15.5, 7.4 Hz, 1H), 2.57 (dd, *J* = 15.6, 6.2 Hz, 1H), 2.62–2.70 (m, 2H), 2.76 (dd, *J* = 13.0, 10.5 Hz, 1H), 2.88–2.98 (m, 2H), 3.22 (dd, *J* = 13.0, 4.8 Hz, 1H), 3.34 (2H) signal partially overlapping with HDO peak, 3.43–3.49 (m, 1H), 3.60–3.67 (m, 1H), 3.73–3.82 (m, 3H), 4.12–4.19 (m, 2H), 4.22 (dd, *J* = 8.8, 5.9 Hz, 1H), 4.41 (ddd, *J* = 10.4, 7.7, 5.0 Hz, 1H), 4.53 (td, *J* = 8.9, 3.9 Hz, 1H), 4.58–4.65 (m, 2H), 5.43–5.49 (m, 1H), 6.60–6.65 (m, 4H), 6.89 (d, *J* = 8.5 Hz, 2H), 6.96 (s, 1H), 7.05 (d, *J* = 8.5 Hz, 2H), 7.20–7.26 (m, 3H), 7.27–7.32 (m, 2H), 7.42 (s, 1H), 7.76 (d, *J* = 8.9 Hz, 1H), 7.91–8.05 (m, 3H), 8.26–8.29 (m, 2H), 8.30–8.33 (m, 1H), 8.35 (dd, *J* = 8.2, 4.4 Hz, 1H), 8.39–8.44 (m, 2H), 8.54 (d, *J* = 8.3 Hz, 1H), 9.18 (s, 1H), 9.19 (s, 1H).

***S. hominis* AIP-II (11)**

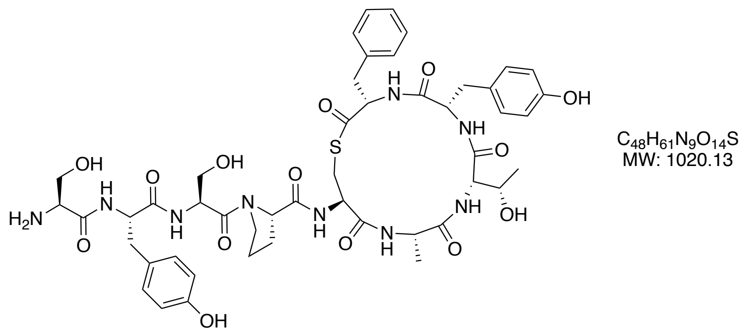

The peptide was synthesized according to general method A for on-resin cleavage-inducing cyclization (section 4.5). Purification by preparative RP-HPLC afforded *S. hominis* AIP-II (**11**) as a fluffy white solid after lyophilization. Purity >95% determined by UPLC ($\lambda$ = 215 nm). **HRMS** (MALDI) *m/z* calcd for $[M+H]^+$ $C_{48}H_{62}N_9O_{14}S^+$: 1020.4130, found 1020.4133. **$^1$H NMR** (600 MHz, DMSO-$d_6$) $\delta$ = 0.97 (d, *J* = 6.3 Hz, 3H), 1.27 (d, *J* = 7.3 Hz, 3H), 1.77–1.93 (m, 3H), 2.03–2.11 (m, 1H), 2.64–2.78 (m, 4H), 2.87–2.98 (m, 2H), 3.15–3.24 (m, 2H), 3.58–3.72 (m, 4H), 3.73–3.82 (m, 2H), 3.96–4.01 (m, 1H), 4.01–4.06 (m, 1H), 4.12–4.17 (m, 1H), 4.18–4.25 (m, 1H), 4.26–4.32 (m, 2H), 4.38 (dd, *J* = 8.4, 4.2 Hz, 1H), 4.53–4.64 (m, 2H), 5.02 (t, *J* = 5.8 Hz, 1H), 5.10 (d, *J* = 5.1 Hz, 1H), 5.46 (t, *J* = 4.8 Hz, 1H), 6.61–6.69 (m, 4H), 6.82–6.86 (m, 2H), 6.92 (d, *J* = 8.4 Hz, 2H), 7.03 (d, *J* = 8.5 Hz, 1H), 7.13–7.17 (m, 3H), 7.75 (d, *J* = 8.0 Hz, 1H), 7.86 (d, *J* = 9.8 Hz, 1H), 8.04 (s, 4H), 8.13 (d, *J* = 8.4 Hz, 1H), 8.30–8.36 (m, 1H), 8.50 (d, *J* = 8.1 Hz, 1H), 8.95 (d, *J* = 7.6 Hz, 1H), 9.21 (s, 1H), 9.24 (s, 1H).

***S. hominis* AIP-IV (13)**

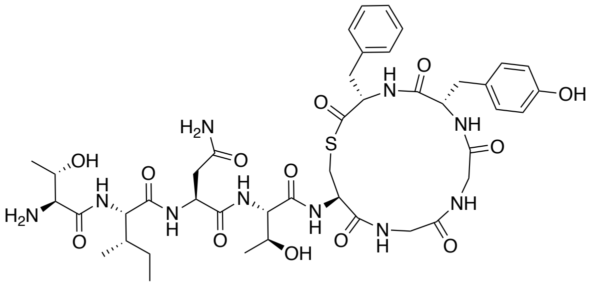

$C_{43}H_{60}N_{10}O_{13}S$
MW: 957.07

The peptide was synthesized according to general method A for on-resin cleavage-inducing cyclization (section 4.5). Purification by preparative RP-HPLC afforded *S. hominis* AIP-IV (**13**) as a fluffy white solid after lyophilization. Purity >95% determined by UPLC ($\lambda = 215$ nm). **UPLC-MS** (ESI) *m/z* calcd for $[M+H]^+$ $C_{43}H_{61}N_{10}O_{13}S^+$: 957.41, found 957.73. **$^1$H NMR** (600 MHz, DMSO-$d_6$) $\delta = 0.82$ (t, $J = 7.4$ Hz, 3H), 0.86 (d, $J = 6.7$ Hz, 3H), 1.00 (d, $J = 6.3$ Hz, 3H), 1.03–1.12 (m, 0H), 1.14 (d, $J = 6.3$ Hz, 3H), 1.47 (ddt, $J = 10.8$, 7.4, 3.3 Hz, 1H), 1.68–1.77 (m, 1H), 2.33 (dd, $J = 14.3$, 10.4 Hz, 1H), 2.43 (dd, $J = 15.6$, 6.8 Hz, 1H), 2.58 (dd, $J = 15.6$, 7.0 Hz, 1H), 2.64 (dd, $J = 14.2$, 4.3 Hz, 1H), 2.81 (dd, $J = 13.1$, 10.2 Hz, 1H), 2.91 (dd, $J = 14.1$, 11.0 Hz, 1H), 3.23 (dd, $J = 13.0$, 4.8 Hz, 1H), 3.33–3.40 (m, 1H), 3.46 (dd, $J = 14.3$, 6.3 Hz, 1H), 3.65–3.69 (m, 2H), 3.77–3.84 (m, 2H), 4.05–4.19 (m, 4H), 4.30 (dd, $J = 8.7$, 7.0 Hz, 1H), 4.44 (ddd, $J = 10.1$, 7.8, 4.9 Hz, 1H), 4.56–4.66 (m, 2H), 4.87 (d, $J = 5.2$ Hz, 1H), 5.54 (d, $J = 5.0$ Hz, 1H), 6.62 (dd, $J = 8.4$, 2.0 Hz, 2H), 6.88 (d, $J = 8.5$ Hz, 2H), 6.96 (s, 1H), 7.18–7.26 (m, 3H), 7.27–7.32 (m, 2H), 7.45 (s, 1H), 7.65 (d, $J = 8.5$ Hz, 1H), 7.94–8.09 (m, 5H), 8.26 (t, $J = 5.6$ Hz, 1H), 8.29 (d, $J = 9.2$ Hz, 1H), 8.35 (d, $J = 7.4$ Hz, 2H), 8.40 (d, $J = 7.9$ Hz, 1H), 8.46 (d, $J = 8.7$ Hz, 1H), 9.19 (s, 1H).

***S. hominis* AIP-V (14)**

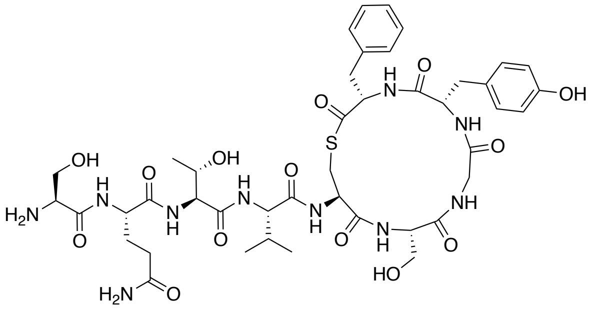

$C_{43}H_{60}N_{10}O_{14}S$
MW: 973.07

The peptide was synthesized according to general method A for on-resin cleavage-inducing cyclization (section 4.5). Purification by preparative RP-HPLC afforded *S. hominis* AIP-V (**14**) as a fluffy white solid after lyophilization. Purity >95% determined by UPLC ($\lambda = 215$ nm). **HRMS** (MALDI) *m/z* calcd for $[M+H]^+$ $C_{43}H_{61}N_{10}O_{14}S^+$: 973.4083, found 973.4096. **$^1$H NMR** (600 MHz, DMSO-$d_6$) $\delta = 0.81$–0.85 (m, 6H), 1.04 (d, $J = 6.3$ Hz, 3H), 1.77–1.85 (m, 1H), 1.88–2.01 (m, 2H), 2.09–2.21 (m, 2H), 2.31 (dd, $J = 14.4$, 10.8 Hz, 1H), 2.62–2.71 (m, 2H), 2.92 (dd, $J = 14.2$, 11.1 Hz, 1H), 3.17 (dd, $J = 12.9$, 4.7 Hz, 1H), 3.33 (1H) signal overlapping with HDO peak, 3.47 (dd, $J = 14.2$, 7.4 Hz, 1H), 3.52–3.57 (m, 1H), 3.60–3.66 (m, 1H), 3.72–3.81 (m, 2H), 3.85–3.90 (m, 1H), 3.97–4.03 (m, 1H), 4.12–4.18 (m, 1H), 4.25 (dd, $J = 8.8$, 6.3 Hz, 1H), 4.30 (dd, $J = 8.3$, 4.3 Hz, 1H), 4.35–4.48 (m, 2H), 4.58–4.65 (m, 1H), 4.95 (t, $J = 5.4$ Hz, 1H), 4.97 (d, $J = 4.8$ Hz, 1H), 5.46 (t, $J = 4.9$ Hz, 1H), 6.62 (d, $J = 8.4$ Hz, 2H), 6.80 (s, 1H), 6.90 (d, $J = 8.5$ Hz, 2H), 7.18–7.28 (m, 4H), 7.27–7.33 (m, 2H), 7.76 (d, $J = 8.8$ Hz, 1H),

7.93–7.98 (m, 1H), 8.09 (s, 3H), 8.16 (d, $J$ = 9.3 Hz, 1H), 8.34 (d, $J$ = 7.7 Hz, 1H), 8.45–8.51 (m, 2H), 8.64 (d, $J$ = 7.9 Hz, 1H), 9.20 (s, 1H).

***S. capitis* AIP-I (15)**

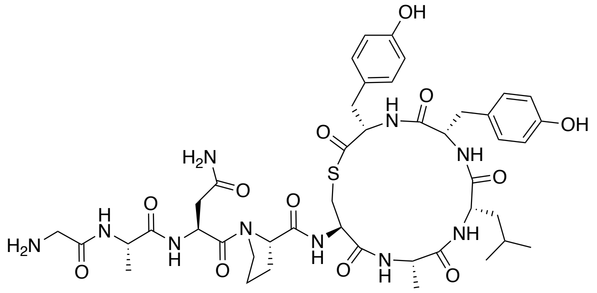

$C_{44}H_{60}N_{10}O_{12}S$
MW: 953.08

The peptide was synthesized according to general method A for on-resin cleavage-inducing cyclization (section 4.5). Purification by preparative RP-HPLC afforded *S. capitis* AIP-I (**15**) as a fluffy white solid after lyophilization. Purity >95% determined by UPLC ($\lambda$ = 215 nm). **HRMS** (MALDI) *m/z* calcd for [M+H $C_{44}H_{61}N_{10}O_{12}S^{+}$: 953.4185, found 953.4182. **$^{1}$H NMR** (600 MHz, DMSO-$d_6$) $\delta$= 0.79 (d, $J$ = 6.5 Hz, 3H), 0.82 (d, $J$ = 6.5 Hz, 3H), 1.13–1.23 (m, 6H), 1.33–1.41 (m, 1H), 1.40–1.51 (m, 3H), 1.78–1.83 (m, 1H), 1.84–1.93 (m, 2H), 1.95–2.10 (m, 4H), 2.35–2.44 (m, 1H), 2.52–2.59 (m, 1H), 2.65–2.73 (m, 2H), 2.72–2.81 (m, 1H), 2.90 (t, $J$ = 12.1 Hz, 1H), 3.11–3.22 (m, 2H), 3.60–3.71 (m, 2H), 4.03–4.13 (m, 2H), 4.12–4.19 (m, 1H), 4.18–4.27 (m, 2H), 4.29–4.34 (m, 1H), 4.34–4.43 (m, 1H), 4.71–4.85 (m, 1H), 6.58–6.69 (m, 4H), 6.81 (d, $J$ = 8.4 Hz, 2H), 6.88 (d, $J$ = 8.2 Hz, 2H), 6.93–7.00 (m, 1H), 7.19 (s, 1H), 7.46 (s, 1H), 7.78 (d, $J$ = 8.9 Hz, 1H), 7.92 (d, $J$ = 7.8 Hz, 1H), 8.07 (s, 1H), 8.31 (d, $J$ = 8.0 Hz, 1H), 8.36 (d, $J$ = 7.3 Hz, 1H), 8.46 (d, $J$ = 7.8 Hz, 1H), 8.93 (d, $J$ = 8.2 Hz, 1H), 9.17–9.23 (m, 2H).

***S. warneri* AIP-II (18)**

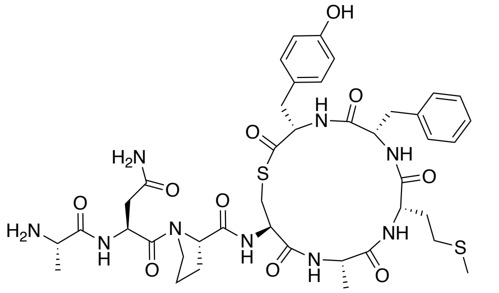

$C_{41}H_{55}N_9O_{10}S_2$
MW: 898.06

The peptide was synthesized according to general method A for on-resin cleavage-inducing cyclization (section 4.5). Purification by preparative RP-HPLC afforded *S. warneri* AIP-II (**18**) as a fluffy white solid after lyophilization. Purity >95% determined by UPLC ($\lambda$ = 215 nm). **UPLC-MS** (ESI) *m/z* calcd for [M+H]$^{+}$ $C_{41}H_{56}N_9O_{10}S_2^{+}$: 898.36, found 898.14. Major conformer (conformers were observed in a ration of 1:0.1): **$^{1}$H NMR** (600 MHz, DMSO-$d_6$) $\delta$= 1.23 (d, $J$ = 7.4 Hz, 3H), 1.31 (d, $J$ = 7.0 Hz, 3H), 1.75–1.84 (m, 3H), 1.86–1.93 (m, 2H), 1.99 (s, 3H), 2.00–2.06 (m, 1H), 2.13–2.27 (m, 2H), 2.42 (dd, $J$ = 15.8, 7.6 Hz, 1H), 2.53–2.66 (m, 1H), 2.73 (dd, $J$ = 14.2, 10.3 Hz, 1H), 2.79–2.97 (m, 3H), 3.10–3.23 (m, 2H), 3.66 (t, $J$ = 6.9 Hz, 2H), 3.77–3.83 (m, 1H), 4.08–4.21 (m, 3H), 4.22–4.35 (m, 3H), 4.85 (q, $J$ = 7.1 Hz, 1H), 6.64 (d, $J$ = 8.0 Hz, 2H), 6.91 (d, $J$ = 8.1 Hz, 2H), 6.99

(s, 1H), 7.06 (d, $J$ = 7.4 Hz, 2H), 7.16–7.33 (m, 3H), 7.50 (s, 1H), 7.88 (d, $J$ = 8.6 Hz, 1H), 7.97 (d, $J$ = 7.6 Hz, 1H), 7.98–8.07 (m, 3H), 8.22 (d, $J$ = 7.7 Hz, 1H), 8.30 (d, $J$ = 8.0 Hz, 1H), 8.70 (d, $J$ = 7.2 Hz, 1H), 8.83 (d, $J$ = 8.4 Hz, 1H), 9.19 (s, 1H).

***S. cohnii* AIP-I (19)**

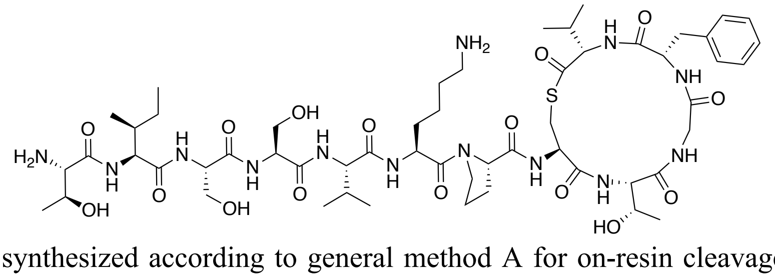

$C_{55}H_{89}N_{13}O_{16}S$
MW: 1220.45

The peptide was synthesized according to general method A for on-resin cleavage-inducing cyclization (section 4.5). Purification by preparative RP-HPLC afforded *S. cohnii* AIP-I (**19**) as a fluffy white solid after lyophilization. Purity 95% determined by UPLC ($\lambda$ = 215 nm). **UPLC-MS** (ESI) *m/z* calcd for $[M+2H]^{2+}$ $C_{55}H_{91}N_{13}O_{16}S^{2+}$: 610.82, found 610.93; $[M+H]^{+}$ $C_{55}H_{90}N_{13}O_{16}S^{+}$: 1220.63, found 1220.69. **$^{1}$H NMR** (600 MHz, DMSO-$d_6$) $\delta$ = 0.80 (d, $J$ = 6.7 Hz, 3H), 0.81–0.85 (m, 6H), 0.86 (d, $J$ = 6.8 Hz, 3H), 0.90 (d, $J$ = 6.8 Hz, 3H), 0.95 (d, $J$ = 6.9 Hz, 3H), 0.99 (d, $J$ = 6.2 Hz, 3H), 1.05–1.12 (m, 1H), 1.14 (d, $J$ = 6.3 Hz, 3H), 1.34 –1.40 (m 2H), 1.44–1.59 (m, 4H), 1.62–1.69 (m, 1H), 1.69–1.77 (m, 2H), 1.80–1.92 (m, 2H), 1.93–2.04 (m, 2H), 2.37–2.45 (m, 1H), 2.63 (dd, $J$ = 13.1, 10.8 Hz, 1H), 2.75–2.81 (m, 2H), 2.85 (dd, $J$ = 14.2, 11.4 Hz, 1H), 3.21 (dd, $J$ = 13.1, 4.8 Hz, 1H), 3.25 (dd, $J$ = 14.2, 3.5 Hz, 1H), 3.41–3.46 (m, 1H), 3.50–3.61 (m, 4H), 3.61–3.71 (m, 4H), 3.76–3.83 (m, 1H), 3.86–3.91 (m, 1H), 4.19 (dd, $J$ = 8.8, 6.2 Hz, 1H), 4.24 (dd, $J$ = 9.1, 4.4 Hz, 1H), 4.32 (dd, $J$ = 8.6, 7.1 Hz, 1H), 4.34–4.46 (m, 6H), 4.56 (dd, $J$ = 9.9, 4.6 Hz, 1H), 4.76-4.80 (m, 1H), 4.95–4.99 (m, 1H), 5.05–5.09 (m, 1H), 5.52–5.58 (m, 1H), 7.19–7.24 (m, 1H), 7.24–7.34 (m, 5H), 7.65–7.75 (m, 4H), 7.89–7.96 (m, 2H), 8.03–8.10 (m, 3H), 8.11–8.16 (m, 2H), 8.19 (d, $J$ = 7.7 Hz, 1H), 8.44 (d, $J$ = 8.6 Hz, 1H), 8.53 (dd, $J$ = 6.8, 4.6 Hz, 1H), 8.77 (d, $J$ = 8.4 Hz, 1H).

***S. caprae* AIP-I (21)**

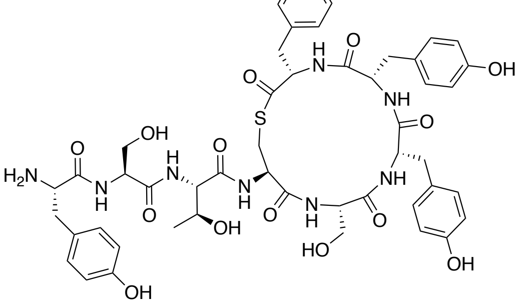

$C_{49}H_{58}N_8O_{14}S$
MW: 1015.11

The peptide was synthesized according to general method A for on-resin cleavage-inducing cyclization (section 3). Purification by preparative RP-HPLC afforded *S. caprae* AIP-I (**21**) as a fluffy white solid after lyophilization. Purity >95% determined by UPLC ($\lambda$ = 215 nm). **UPLC-MS** (ESI) *m/z* calcd for $[M+H]^{+}$ $C_{49}H_{59}N_8O_{14}S^{+}$: 1015.39, found 1015.53. **$^{1}$H NMR** (600 MHz, DMSO-$d_6$) $\delta$ = 1.05 (d, $J$ = 6.3 Hz, 3H), 2.62–2.72 (m, 2H), 2.73–2.86 (m,

4H), 2.90 (dd, $J$ = 12.7, 11.0 Hz, 1H), 3.03 (dd, $J$ = 14.3, 4.7 Hz, 1H), 3.20–3.28 (m, 2H), 3.35–3.39 (m, 1H), 3.44–3.50 (m, 1H), 3.54–3.61 (m, 1H), 3.65–3.72 (m, 1H), 4.01 (dd, $J$ = 8.3, 4.8 Hz, 1H), 4.06–4.14 (m, 4H), 4.26 (dd, $J$ = 8.5, 3.4 Hz, 1H), 4.36 (dt, $J$ = 9.2, 7.0 Hz, 1H), 4.45 (ddd, $J$ = 11.7, 8.1, 4.0 Hz, 1H), 4.55 (dt, $J$ = 7.7, 6.2 Hz, 1H), 4.92 (d, $J$ = 5.1 Hz, 1H), 5.03 (t, $J$ = 5.3 Hz, 1H), 5.25 (t, $J$ = 5.2 Hz, 1H), 6.60–6.66 (m, 4H), 6.69 (d, $J$ = 8.5 Hz, 1H), 6.78 (d, $J$ = 8.5 Hz, 2H), 6.89–6.96 (m, 4H), 7.07 (d, $J$ = 8.5 Hz, 2H), 7.14–7.22 (m, 3H), 7.85 (d, $J$ = 9.1 Hz, 1H), 7.86–7.99 (m, 3H), 7.97 (d, $J$ = 8.5 Hz, 1H), 8.02 (d, $J$ = 8.1 Hz, 1H), 8.18–8.23 (m, 2H), 8.72 (d, $J$ = 7.7 Hz, 1H), 8.94 (d, $J$ = 7.9 Hz, 1H), 9.22 (s, 1H), 9.24 (s, 1H), 9.34 (s, 1H).

***S. pasteuri* AIP-I (22)**

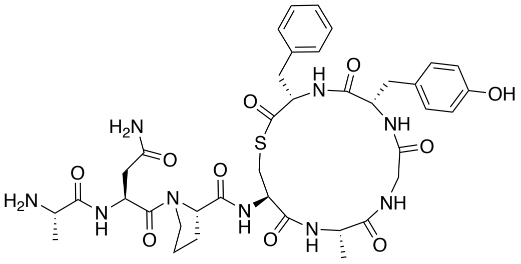

$C_{38}H_{49}N_9O_{10}S$
MW: 823.92

The peptide was synthesized according to general method A for on-resin cleavage-inducing cyclization (section 4.5). Purification by preparative RP-HPLC afforded *S. pasteuri* AIP-I (**22**) as a fluffy white solid after lyophilisation. Purity >95% determined by UPLC ($\lambda$ = 215 nm). **UPLC-MS** (ESI) *m/z* calcd for $[M+H]^+$ $C_{38}H_{50}N_9O_{10}S^+$: 824.34, found 824.26. Major conformer (conformers were observed in a ration of 1:0.15): **$^1$H NMR** (600 MHz, DMSO-$d_6$) $\delta$ = 1.19 (d, $J$ = 7.1 Hz, 3H), 1.31 (d, $J$ = 7.0 Hz, 3H), 1.79–1.94 (m, 3H), 1.95–2.03 (m, 1H), 2.31 (dd, $J$ = 14.3, 10.8 Hz, 1H), 2.42 (dd, $J$ = 15.7, 7.9 Hz, 1H), 2.59 (dd, $J$ = 15.7, 5.5 Hz, 1H), 2.63–2.72 (m, 2H), 2.97 (dd, $J$ = 14.2, 11.2 Hz, 1H), 3.18 (dd, $J$ = 13.0, 5.0 Hz, 1H), 3.30–3.39 (m, 2H), 3.62–3.67 (m, 2H), 3.78–3.88 (m, 2H), 4.14 (ddd, $J$ = 11.5, 7.9, 4.0 Hz, 1H), 4.34–4.42 (m, 2H), 4.43–4.49 (m, 1H), 4.60 (ddd, $J$ = 11.2, 9.3, 3.8 Hz, 1H), 4.81–4.88 (m, 1H), 6.62 (d, $J$ = 8.4 Hz, 2H), 6.90 (d, $J$ = 8.4 Hz, 2H), 6.99 (s, 1H), 7.20–7.24 (m, 1H), 7.25–7.33 (m, 4H), 7.48 (s, 1H), 8.03–8.10 (m, 4H), 8.21 (d, $J$ = 9.4 Hz, 1H), 8.25 (d, $J$ = 8.7 Hz, 1H), 8.48 (d, $J$ = 8.0 Hz, 1H), 8.53 (t, $J$ = 5.5 Hz, 1H), 8.70 (d, $J$ = 7.3 Hz, 1H), 9.20 (br s, 1H).

***S. devriesei* AIP-I (23)**

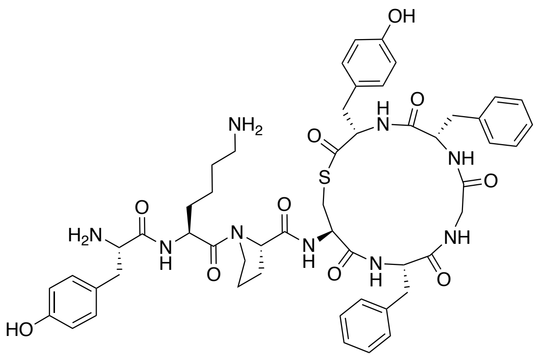

$C_{52}H_{63}N_9O_{10}S$
MW: 1006.19

The peptide was synthesized according to general method A for on-resin cleavage-inducing cyclization (section 4.5). Purification by preparative RP-HPLC afforded *S. devriesei* AIP-I (**23**) as a fluffy white solid after lyophilization. Purity >95% determined by UPLC ($\lambda$ = 215 nm). **UPLC-MS** (ESI) *m/z* calcd for $[M+H]^+$

$C_{52}H_{64}N_9O_{10}S^+$: 1006.45, found 1006.71. Major conformer (conformers were observed in a ration of 1:0.14): **$^1$H NMR** (600 MHz, DMSO-$d_6$) $\delta$ = 1.34–1.47 (m, 2H), 1.48–1.66 (m, 4H), 1.68–1.76 (m, 1H), 1.77–1.89 (m, 2H), 1.89–2.00 (m, 1H), 2.31 (dd, $J$ = 14.3, 10.9 Hz, 1H), 2.61 (dd, $J$ = 12.9, 10.5 Hz, 1H), 2.68 (dd, $J$ = 14.3, 3.8 Hz, 1H), 2.72–2.87 (m, 4H), 2.88–2.96 (m, 2H), 3.03 (dd, $J$ = 14.4, 11.4 Hz, 1H), 3.19 (dd, $J$ = 12.9, 4.9 Hz, 1H), 3.29–3.42 (m, 2H), 3.54 (t, $J$ = 6.8 Hz, 2H), 3.86 (dd, $J$ = 14.2, 4.7 Hz, 1H), 3.94–4.06 (m, 1H), 4.14 (ddd, $J$ = 11.3, 8.0, 3.8 Hz, 1H), 4.38 (dd, $J$ = 8.3, 4.3 Hz, 1H), 4.42 (ddd, $J$ = 10.4, 8.2, 5.1 Hz, 1H), 4.51 (q, $J$ = 7.4 Hz, 1H), 4.59–4.68 (m, 2H), 6.62 (d, $J$ = 8.5 Hz, 2H), 6.69 (d, $J$ = 8.4 Hz, 2H), 6.91 (d, $J$ = 8.5 Hz, 2H), 6.99 (d, $J$ = 8.5 Hz, 2H), 7.15–7.38 (m, 10H), 7.73–7.86 (m, 3H), 8.04–8.14 (m, 4H), 8.18 (d, $J$ = 9.3 Hz, 1H), 8.39 (d, $J$ = 9.1 Hz, 1H), 8.59 (d, $J$ = 7.9 Hz, 1H), 8.67 (t, $J$ = 5.5 Hz, 1H), 8.75 (d, $J$ = 7.9 Hz, 1H), 8.96–9.62 (m, 2H).

***S. succinus* AIP-I (24)**

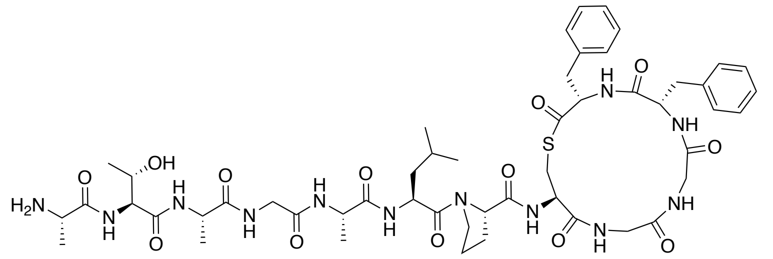

The peptide was synthesized according to general method A for on-resin cleavage-inducing cyclization (section 4.5). Purification by preparative RP-HPLC afforded *S. succinus* AIP-I (**24**) as a fluffy white solid after lyophilization. Purity >95% determined by UPLC ($\lambda$ = 215 nm). **UPLC-MS** (ESI) *m/z* calcd for $[M+2H]^{2+}$ $C_{51}H_{74}N_{12}O_{13}S^{2+}$: 547.26, found 547.39; $[M+H]^+$ $C_{51}H_{73}N_{12}O_{13}S^+$: 1093.51, found 1093.48. Major conformer (conformers were observed in a ration of 1:0.07): **$^1$H NMR** (600 MHz, DMSO-$d_6$) $\delta$ = 0.87–0.92 (m, 6H), 1.08 (d, $J$ = 6.3 Hz, 3H), 1.18 (d, $J$ = 7.0 Hz, 3H), 1.24 (d, $J$ = 7.0 Hz, 3H), 1.35 (d, $J$ = 6.9 Hz, 3H), 1.40–1.52 (m, 2H), 1.60–1.69 (m, 1H), 1.71–1.77 (m, 1H), 1.82–1.88 (m, 1H), 1.88–1.95 (m, 1H) 1.98–2.07 (m, 1H), 2.44 (dd, $J$ = 14.3, 10.7 Hz, 1H), 2.69 (dd, $J$ = 13.0, 10.2 Hz, 1H), 2.78 (dd, $J$ = 14.3, 4.1 Hz, 1H), 2.93 (dd, $J$ = 14.1, 11.0 Hz, 1H), 3.23 (dd, $J$ = 13.0, 4.8 Hz, 1H), 3.30–3.37 (m, 2H), 3.42–3.51 (m, 2H), 3.62–3.75 (m, 3H), 3.79 (dd, $J$ = 14.2, 4.4 Hz, 1H), 3.96–4.06 (m, 2H), 4.17 (dd, $J$ = 15.6, 8.4 Hz, 1H), 4.22–4.33 (m, 4H), 4.34–4.40 (m, 2H), 4.50 (ddd, $J$ = 9.6, 8.0, 4.8 Hz, 1H), 4.64 (ddd, $J$ = 11.0, 9.2, 4.0 Hz, 1H), 7.08–7.12 (m, 2H), 7.16–7.20 (m, 1H), 7.20–7.28 (m, 5H), 7.29–7.34 (m, 2H), 7.85 (d, $J$ = 7.5 Hz, 1H), 8.01 (d, $J$ = 7.0 Hz, 1H), 8.03–8.07 (m, 4H), 8.17 (t, $J$ = 5.8 Hz, 1H), 8.20 (d, $J$ = 7.7 Hz, 1H), 8.31 (d, $J$ = 9.3 Hz, 1H), 8.33–8.39 (m, 2H), 8.41 (d, $J$ = 8.4 Hz, 1H), 8.51 (d, $J$ = 8.0 Hz, 1H).

***S. equorum* AIP-I (25)**

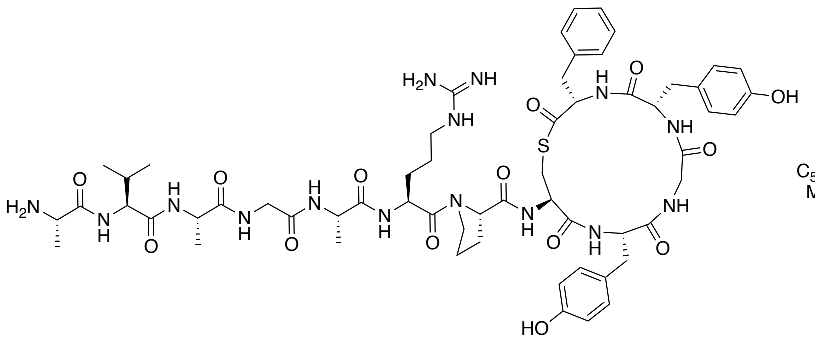

The peptide was synthesized according to general method A for on-resin cleavage-inducing cyclization (section 4.5). Purification by preparative RP-HPLC afforded *S. equorum* AIP-I (**25**) as a fluffy white solid after lyophilization. Purity >95% determined by UPLC ($\lambda$ = 215 nm). **UPLC-MS** (ESI) *m/z* calcd for $[M+H]^+$ $C_{59}H_{82}N_{15}O_{14}S^+$: 1256.59, found 1256.11. Major conformer (conformers were observed in a ration of 1:0.12): **$^1$H NMR** (600 MHz, DMSO-$d_6$) $\delta$ = 0.85 (d, *J* = 6.7 Hz, 3H), 0.89 (d, *J* = 6.7 Hz, 3H), 1.19 (d, *J* = 7.0 Hz, 3H), 1.19–1.25 (m, 3H), 1.32 (d, *J* = 6.9 Hz, 3H), 1.51–1.65 (m, 4H), 1.68–1.76 (m, 1H), 1.80–2.04 (m, 4H), 2.30 (dd, *J* = 14.1, 10.9 Hz, 1H), 2.57 (dd, *J* = 13.0, 10.6 Hz, 1H), 2.66–2.75 (m, 2H), 2.78 (dd, *J* = 14.0, 5.5 Hz, 1H), 2.98–3.07 (m, 1H), 3.07–3.21 (m, 3H), 3.25–3.32 (m, 1H), 3.32–3.38 (m, 1H), 3.49–3.59 (m, 1H), 3.60–3.70 (m, 2H), 3.74 (dd, *J* = 16.8, 5.8 Hz, 1H), 3.83 (dd, *J* = 14.1, 4.7 Hz, 1H), 3.89–3.99 (m, 1H), 4.10–4.18 (m, 1H), 4.18–4.23 (m, 1H), 4.22–4.29 (m, 1H), 4.27–4.33 (m, 1H), 4.32–4.37 (m, 1H), 4.37–4.44 (m, 1H), 4.44–4.52 (m, 1H), 4.51–4.58 (m, 1H), 4.57–4.64 (m, 1H), 6.58–6.63 (m, 4H), 6.70–7.41 (broad signal, 4H), 6.91 (d, *J* = 8.3 Hz, 2H), 7.01 (d, *J* = 8.4 Hz, 2H), 7.20–7.26 (m, 1H), 7.27–7.39 (m, 4H), 7.48–7.64 (m, 1H), 7.87 (d, *J* = 7.2 Hz, 1H), 8.02–8.07f (m, 3H), 8.08–8.21 (m, 5H), 8.27 (d, *J* = 9.1 Hz, 1H), 8.32 (d, *J* = 8.6 Hz, 1H), 8.51 (d, *J* = 7.9 Hz, 1H), 8.66 (d, *J* = 5.6 Hz, 1H), 9.05–9.28 (m, 2H).

***S. equorum* AIP-II (26)**

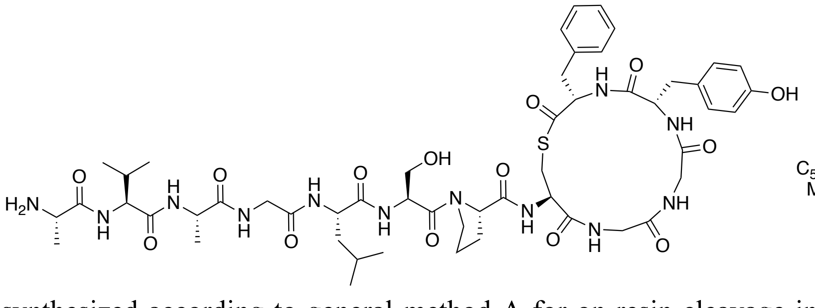

The peptide was synthesized according to general method A for on-resin cleavage-inducing cyclization (section 4.5). Purification by preparative RP-HPLC afforded *S. equorum* AIP-II (**26**) as a fluffy white solid after lyophilization. Purity >95% determined by UPLC ($\lambda$ = 215 nm). **UPLC-MS** (ESI) *m/z* calcd for $[M+H]^+$ $C_{52}H_{75}N_{12}O_{14}S^+$: 1123.52, found 1123.27. Major conformer (conformers were observed in a ration of 1:0.45) **$^1$H NMR** (600 MHz, DMSO-$d_6$) $\delta$ = 0.79–0.83 (m, 3H), 0.83–0.88 (m, 6H), 0.86–0.91 (m, 3H), 1.21 (d, *J* = 7.1 Hz, 3H), 1.22–1.31 (m, 1H), 1.32 (d, *J* = 6.9 Hz, 3H), 1.33–1.42 (m, 1H), 1.40–1.49 (m, 1H), 1.49–1.64 (m, 1H), 1.78–2.06 (m, 4H), 2.08–2.22 (m, 1H), 2.27 (t, *J* = 8.8 Hz, 1H), 2.54–2.66 (m, 1H), 2.68–2.79 (m, 1H), 2.81–3.09 (m, 3H), 3.34 (t, *J* = 10.6 Hz, 1H), 3.50–3.59 (m, 3H), 3.60–3.73 (m, 2H), 3.74–4.02 (m, 4H), 4.04–4.17 (m, 1H), 4.17–

4.53 (m, 6H), 4.53–4.72 (m, 2H), 4.99–5.13 (m, 1H), 6.66 (d, $J$ = 8.4 Hz, 2H), 7.09–7.18 (m, 2H), 7.18–7.42 (m, 5H), 7.61 (d, $J$ = 9.4 Hz, 1H), 7.85–8.02 (m, 2H), 8.02–8.18 (m, 5H), 8.32–8.41 (m, 2H), 8.55 (t, $J$ = 6.3 Hz, 1H), 9.11–9.21 (m, 1H), 9.28–9.40 (m, 1H).

***S. intermedius*** **AIP-I (32)**

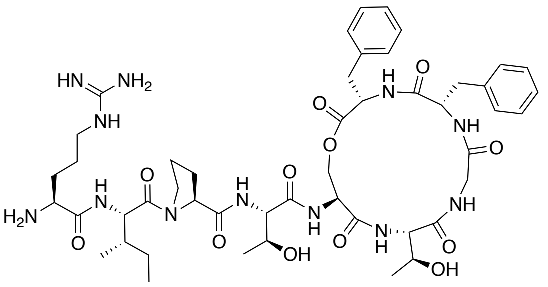

The peptide was synthesized as followed (Supporting Scheme S3). The fully protected linear peptide **S10** was synthesized in 40.0 μmol scale on Cl-Trt polystyrene resin **S7** preloaded with Fmoc-Phe-OH (0.69 mmol/g) using the general procedures for automated SPPS. After completed peptide elongation, the peptidyl-resin **S10** was transferred into a polypropylene syringe equipped with a fritted disk using $CH_2Cl_2$, washed with $CH_2Cl_2$ (3 × 1 min) and dried overnight under vacuum. The peptidyl-resin **S10** was swelled in anhydrous THF (1.5 mL) for 15 min and subsequently a solution of tetrabutylammonium fluoride (TBAF) in THF (1.0 M) (0.40 mL, 0.40 mmol, 10.0 equiv) in anhydrous THF (4.6 mL) was added to the resin and the resin was agitated at room temperature. After 1 h, the TBAF solution was removed by suction and resin treated with a fresh TBAF–THF solution. After 1 h, the TBAF solution was removed by suction and the resin washed with DMF (3 × 1 min), MeOH (3 × 1 min) and $CH_2Cl_2$ (3 × 1 min) and dried overnight under vacuum. On-resin esterification was performed according to a previously published protocol.[27] The resin was swelled in anhydrous $CH_2Cl_2$ (1.5 mL) for 15 min and subsequently a solution of Fmoc-Phe-OH (93.0 mg, 0.24 mmol, 6.00 equiv), *N*,*N*'-diisopropylcarbodiimide (DIC) (37.6 μL, 0.24 mmol, 6.00 equiv), *N*-methylimidazole (NMI) (17.2 μL, 0.22 mmol, 5.40 equiv) in anhydrous $CH_2Cl_2$ (1.5 mL) was added. The resin was agitated at room temperature for 2 h and subsequently washed with anhydrous $CH_2Cl_2$ (3 × 1 min). A fresh Fmoc-Phe-OH–DIC–NMI solution was added to resin and after 2 h of incubation, the resin was washed with DMF (3 × 1 min), $CH_2Cl_2$ (3 × 1 min) and DMF (5 × 1 min). Fmoc-removal of the *O*-acylated peptidyl resin **S11** was performed by treatment of the resin with a solution of 1,8-biazabicyclo[5.4.0]undec-7-ene (DBU) in DMF (1.5 mL, 1:99, v/v) (8 × 30 s). The resin was subsequently washed with DMF (3 × 1 min), MeOH (3 × 1 min) and $CH_2Cl_2$ (3 × 1 min) and dried under suction for 15 min. The dried resin was treated with a solution of HFIP (0.4 mL) in $CH_2Cl_2$ (1.6 mL) for 15 min at room temperature to cleave the partially protected peptide **S12** from the resin. The cleavage solution was removed from the resin and collected and a fresh HFIP–$CH_2Cl_2$ solution was added to the resin. After 15 min the cleavage solution was removed from the resin and collected and the resin rinsed with $CH_2Cl_2$ (2.0 mL). The combined cleavage solutions and the rinsing solution were evaporated to dryness under reduced pressure to yield the partially protected peptide **S12,** which was used without further purification. Crude peptide **S12** (0.04 mmol based on the resin loading) was dissolved in anhydrous DMF (5.0 mL) under nitrogen atmosphere and added dropwise to a solution of HATU (15.2 mg, 0.04 mmol, 1.00 equiv) and *i*-$Pr_2NEt$ (20.8 μL, 0.12 mmol,

3.00 equiv) in anhydrous DMF (35.0 mL). The reaction mixture was stirred overnight at room temperature and after full consumption of **S12** was confirmed by UPLC-MS, the reaction was reduced to dryness under reduced pressure. The remaining residue was treated with a deprotection cocktail (3.0 mL, TFA–*i*-$Pr_3SiH$–water, 94:3:3, v/v/v) for 2 h and subsequently concentrated under a stream of nitrogen followed by precipitation in ice-cold diethyl ether. Purification by preparative RP-HPLC afforded *S. intermedius* AIP-I (**32**) as a fluffy white solid after lyophilization. Purity >95% determined by UPLC ($\lambda$ = 215 nm). **UPLC-MS** (ESI) *m/z* calcd for $[M+2H]^{2+}$ $C_{48}H_{72}N_{12}O_{12}^{2+}$: 504.27, found 504.53; $[M+H]^{+}$ $C_{48}H_{71}N_{12}O_{12}^{+}$: 1007.53, found 1007.68. Major conformer (conformers were observed in a ration of 1:0.12): **$^1$H NMR** (600 MHz, DMSO-$d_6$) $\delta$ = 0.81–0.86 (t, *J* = 7.4 Hz, 3H), 0.92 (d, *J* = 6.7 Hz, 3H), 1.03 (d, *J* = 6.3 Hz, 3H), 1.07 (d, *J* = 6.3 Hz, 3H), 1.08–1.13 (m, 1H), 1.41–1.51 (m, 2H), 1.51–1.59 (m, 1H), 1.60–1.71 (m, 1H), 1.72–1.88 (m, 3H), 1.88–1.96 (m, 1H), 1.96–2.05 (m, 1H), 2.82 (dd, *J* = 14.1, 11.5 Hz, 1H), 3.03 (dd, *J* = 13.8, 9.3 Hz, 1H), 3.06–3.16 (m, 3H), 3.26–3.32 (m, 1H), 3.60 (dt, *J* = 9.7, 6.6 Hz, 1H), 3.70–3.79 (m, 2H), 3.82–3.92 (m, 3H), 4.01–4.06 (m, 1H), 4.12–4.20 (m, 2H), 4.19–4.27 (m, 2H), 4.38 (t, *J* = 8.3 Hz, 1H), 4.43 (dd, *J* = 8.2, 4.6 Hz, 1H), 4.62 (td, *J* = 9.2, 6.2 Hz, 1H), 4.75–4.83 (m, 1H), 4.90–5.07 (m, 2H), 6.85–7.48 (4H) broad signal of guanidium group, 7.16–7.36 (m, 10H), 7.64 (d, *J* = 7.9 Hz, 1H), 7.70 (t, *J* = 6.0 Hz, 1H), 7.86 (d, *J* = 8.5 Hz, 1H), 7.93 (d, *J* = 9.1 Hz, 1H), 8.12–8.21 (m, 3H), 8.31 (d, *J* = 8.0 Hz, 1H), 8.39 (d, *J* = 7.2 Hz, 1H), 8.59 (d, *J* = 8.0 Hz, 1H), 9.02 (t, *J* = 5.6 Hz, 1H).

***S. simulans* AIP-II (34)**

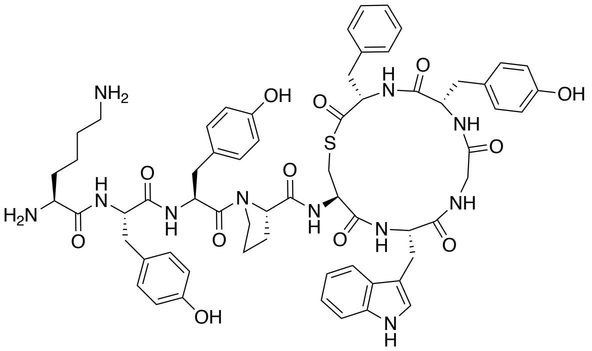

$C_{63}H_{73}N_{11}O_{12}S$
MW: 1208.40

The peptide was synthesized according to general method A for on-resin cleavage-inducing cyclization (section 4.5). Purification by preparative RP-HPLC afforded *S. simulans* AIP-II (**34**) as a fluffy white solid after lyophilization. Purity >95% determined by UPLC ($\lambda$ = 215 nm). **UPLC-MS** (ESI) *m/z* calcd for $[M+2H]^{2+}$ $C_{63}H_{75}N_{11}O_{12}S^{2+}$: 604.77, found 605.07; $[M+H]^{+}$ $C_{63}H_{74}N_{11}O_{12}S^{+}$: 1208.52, found 1208.70. Major conformer (conformers were observed in a ration of 1:0.21): **$^1$H NMR** (600 MHz, DMSO-$d_6$) $\delta$ = 1.24–1.34 (m, 2H), 1.47–1.55 (m, 2H), 1.63–1.74 (m, 3H), 1.75–1.82 (m, 1H), 1.83–1.95 (m, 2H), 2.32 (dd, *J* = 14.2, 10.9 Hz, 1H), 2.57–2.78 (m, 6H), 2.86–2.99 (m, 3H), 3.00–3.09 (m, 2H), 3.27 (dd, *J* = 12.9, 5.0 Hz, 1H), 3.31–3.41 (3H) signals overlapping with HDO peak, 3.48–3.54 (m, 1H), 3.69–3.76 (m, 1H), 3.84 (dd, *J* = 14.0, 4.7 Hz, 1H), 4.11–4.19 (m, 1H), 4.33–4.38 (m, 1H), 4.44–4.53 (m, 2H), 4.52–4.60 (m, 1H), 4.59–4.66 (m, 1H), 4.65–4.72 (m, 1H), 6.59–6.69 (m, 6H), 6.89–6.93 (m, 2H), 6.94–6.99 (m, 1H), 7.01 (d, *J* = 8.5 Hz, 2H), 7.03–7.07 (m, 1H), 7.08–7.13 (m, 2H), 7.15 (d, *J* = 2.4 Hz, 1H), 7.20–7.26 (m, 1H), 7.26–7.37 (m, 5H), 7.59 (d, *J* = 7.9 Hz, 1H), 7.65–7.76 (m, 3H), 7.99–8.09 (m, 5H), 8.19 (d, *J* = 9.3 Hz, 1H), 8.34–8.40 (m, 2H), 8.46 (d, *J* = 8.2 Hz, 1H), 8.51 (d, *J* = 8.0 Hz, 1H), 8.73 (t, *J* = 5.6 Hz, 1H), 9.16–9.21 (m, 3H), 10.80 (d, *J* = 2.4 Hz, 1H).

***S. simulans* AIP-III (35)**

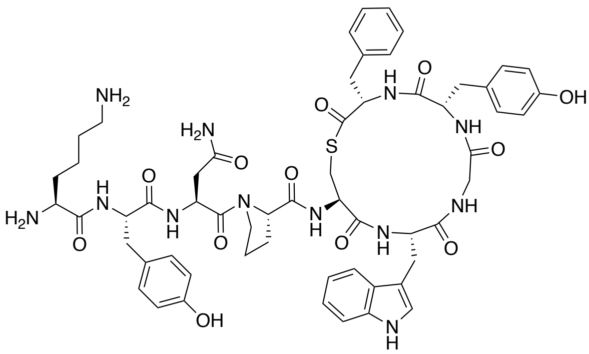

$C_{58}H_{70}N_{12}O_{12}S$
MW: 1159.33

The peptide was synthesized according to general method A for on-resin cleavage-inducing cyclization (section 4.5). Purification by preparative RP-HPLC afforded *S. simulans* AIP-III (**35**) as a fluffy white solid after lyophilization. Purity >95% determined by UPLC ($\lambda$ = 215 nm). **UPLC-MS** (ESI) *m/z* calcd for $[M+2H]^{2+}$ $C_{58}H_{72}N_{12}O_{12}S^{2+}$: 580.25, found 580.46; $[M+H]^{+}$ $C_{58}H_{71}N_{12}O_{12}S^{+}$: 1159.50, found 1159.68. Major conformer (conformers were observed in a ration of 1:0.10): $^{1}$**H NMR** (600 MHz, DMSO-$d_6$) $\delta$ = 1.27–1.36 (m, 2H), 1.48–1.57 (m, 2H), 1.66–1.74 (m, 2H), 1.73–1.80 (m, 1H), 1.78–1.88 (m, 2H), 1.86–1.95 (m, 1H), 2.31 (dd, *J* = 14.2, 10.7 Hz, 1H), 2.38 (dd, *J* = 15.6, 7.1 Hz, 1H), 2.58–2.64 (m, 1H), 2.64–2.71 (m, 3H), 2.72–2.81 (m, 2H), 2.89 (dd, *J* = 14.1, 4.7 Hz, 1H), 2.94–3.08 (m, 3H), 3.16 (dd, *J* = 13.0, 4.9 Hz, 1H), 3.31–3.41 (2H) signals overlapping with HDO peak, 3.50–3.62 (m, 2H), 3.72–3.81 (m, 2H), 4.11–4.17 (m, 1H), 4.32–4.37 (m, 1H), 4.36–4.43 (m, 1H), 4.48–4.55 (m, 1H), 4.58–4.68 (m, 2H), 4.82 (q, *J* = 7.1 Hz, 1H), 6.60–6.67 (m, 4H), 6.90 (d, *J* = 8.4 Hz, 2H), 6.93–6.99 (m, 2H), 7.02 (d, *J* = 8.5 Hz, 2H), 7.03–7.07 (m, 1H), 7.16 (d, *J* = 2.3 Hz, 1H), 7.20–7.26 (m, 1H), 7.26–7.34 (m, 5H), 7.47 (s, 1H), 7.56 (d, *J* = 7.9 Hz, 1H), 7.68–7.76 (m, 3H), 8.03 (d, *J* = 8.0 Hz, 1H), 8.06–8.09 (m, 3H), 8.18–8.26 (m, 2H), 8.48–8.54 (m, 3H), 8.64 (t, *J* = 5.6 Hz, 1H), 9.13–9.35 (m, 2H), 10.81 (d, *J* = 2.5 Hz, 1H).

#### 5.2 *S. simulans* AIP SAR study

The following peptide have been characterized elsewhere: *S. aureus* AIP-III D4A (**58**).[8]

***S. simulans* AIP-II K1A (37)**

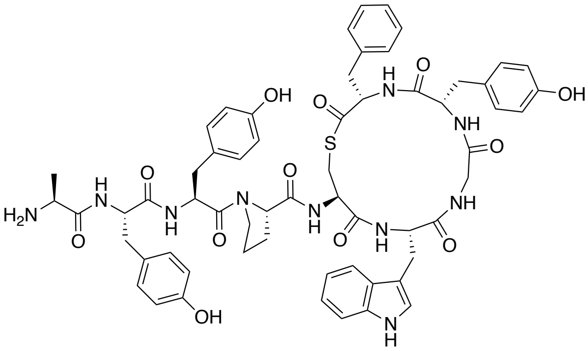

$C_{60}H_{66}N_{10}O_{12}S$
MW: 1151.31

The peptide was synthesized according to general method A for on-resin cleavage-inducing cyclization (section 4.5). Purification by preparative RP-HPLC afforded *S. simulans* AIP-II K1A (**37**) as a fluffy white solid after lyophilization. Purity >95% determined by UPLC ($\lambda = 215$ nm). **UPLC-MS** (ESI) *m/z* calcd for $[M+H]^+$ $C_{60}H_{67}N_{10}O_{12}S^+$: 1151.47, found 1151.45.

***S. simulans* AIP-II Y2A (38)**

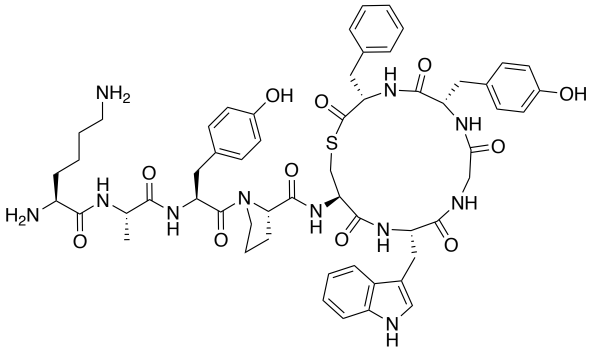

$C_{57}H_{69}N_{11}O_{11}S$
MW: 1116.31

The peptide was synthesized according to general method A for on-resin cleavage-inducing cyclization (section 4.5). Purification by preparative RP-HPLC afforded *S. simulans* AIP-II Y2A (**38**) as a fluffy white solid after lyophilization. Purity >95% determined by UPLC ($\lambda = 215$ nm). **UPLC-MS** (ESI) *m/z* calcd for $[M+H]^+$ $C_{57}H_{70}N_{11}O_{11}S^+$: 1116.50, found 1116.71.

***S. simulans* AIP-II/III Y3A/N3A (39)**

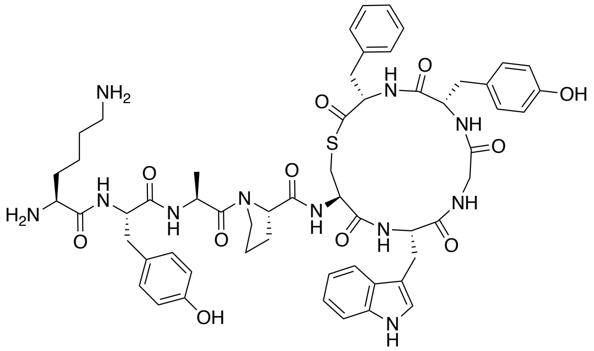

$C_{57}H_{69}N_{11}O_{11}S$
MW: 1116.31

The peptide was synthesized according to general method A for on-resin cleavage-inducing cyclization (section 4.5). Purification by preparative RP-HPLC afforded *S. simulans* AIP-II/III Y3A/N3A (**39**) as a fluffy white solid after lyophilization. Purity >95% determined by UPLC ($\lambda = 215$ nm). **UPLC-MS** (ESI) *m/z* calcd for $[M+H]^+$ $C_{57}H_{70}N_{11}O_{11}S^+$: 1116.50, found 1116.65.

***S. simulans* AIP-II P4A (40)**

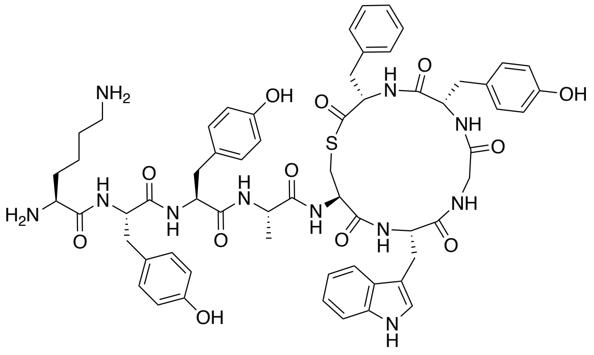

$C_{61}H_{71}N_{11}O_{12}S$
MW: 1182.36

The peptide was synthesized according to general method A for on-resin cleavage-inducing cyclization (section 4.5). Purification by preparative RP-HPLC afforded *S. simulans* AIP-II P4A (**40**) as a fluffy white solid after lyophilization. Purity >95% determined by UPLC ($\lambda = 215$ nm). **UPLC-MS** (ESI) *m/z* calcd for $[M+H]^+$ $C_{61}H_{72}N_{11}O_{12}S^+$: 1182.51, found 1182.76.

***S. simulans* AIP-II W6A (41)**

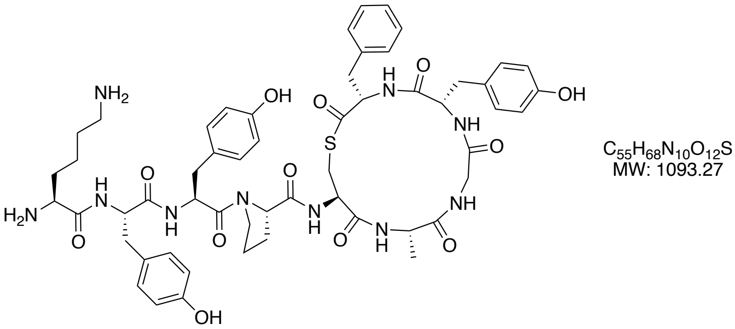

The peptide was synthesized according to general method A for on-resin cleavage-inducing cyclization (section 4.5). Purification by preparative RP-HPLC afforded *S. simulans* AIP-II W6A (**41**) as a fluffy white solid after lyophilization. Purity >95% determined by UPLC ($\lambda = 215$ nm). **UPLC-MS** (ESI) *m/z* calcd for $[M+H]^+$ $C_{55}H_{69}N_{10}O_{12}S^+$: 1093.48, found 1093.55.

***S. simulans* AIP-II G7A (42)**

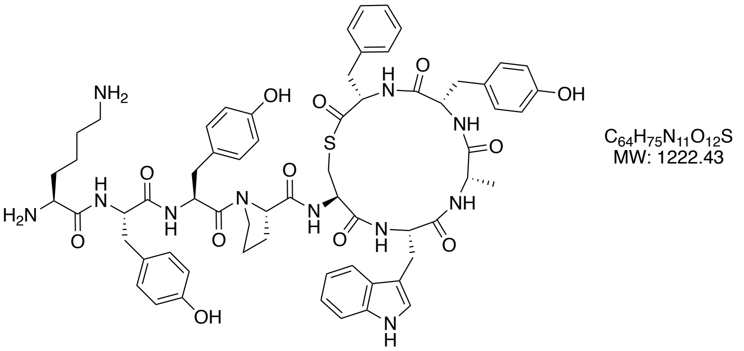

The peptide was synthesized according to general method A for on-resin cleavage-inducing cyclization (section 4.5). Purification by preparative RP-HPLC afforded *S. simulans* AIP-II G7A (**42**) as a fluffy white solid after lyophilization. Purity >95% determined by UPLC ($\lambda = 215$ nm). **UPLC-MS** (ESI) *m/z* calcd for $[M+H]^+$ $C_{64}H_{76}N_{11}O_{12}S^+$: 1222.54, found 1222.60.

***S. simulans* AIP-II Y8A (43)**

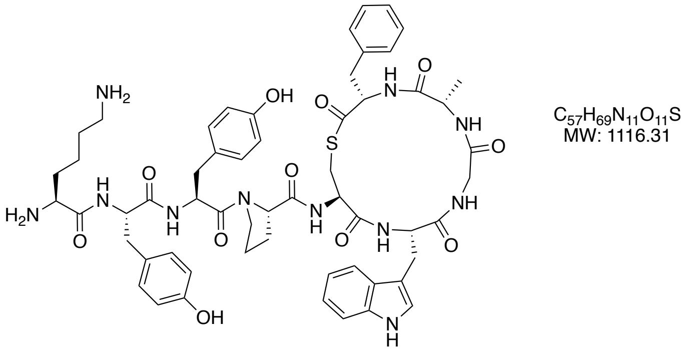

The peptide was synthesized according to general method A for on-resin cleavage-inducing cyclization (section 4.5). Purification by preparative RP-HPLC afforded *S. simulans* AIP-II Y8A (**43**) as a fluffy white solid after lyophilization. Purity >95% determined by UPLC ($\lambda = 215$ nm). **UPLC-MS** (ESI) *m/z* calcd for $[M+H]^+$ $C_{57}H_{70}N_{11}O_{11}S^+$: 1116.50, found 1116.61.

***S. simulans*** **AIP-II F9A (44)**

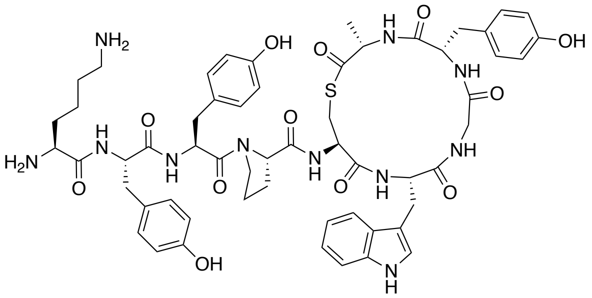

$C_{57}H_{69}N_{11}O_{12}S$
MW: 1132.30

The peptide was synthesized according to general method A for on-resin cleavage-inducing cyclization (section 4.5). Purification by preparative RP-HPLC afforded *S. simulans* AIP-II F9A (**44**) as a fluffy white solid after lyophilization. Purity >95% determined by UPLC ($\lambda = 215$ nm). **UPLC-MS** (ESI) *m/z* calcd for $[M+H]^+$ $C_{57}H_{70}N_{11}O_{12}S^+$: 1132.49, found 1132.64.

***S. simulans*** **AIP-II lactam (45)**

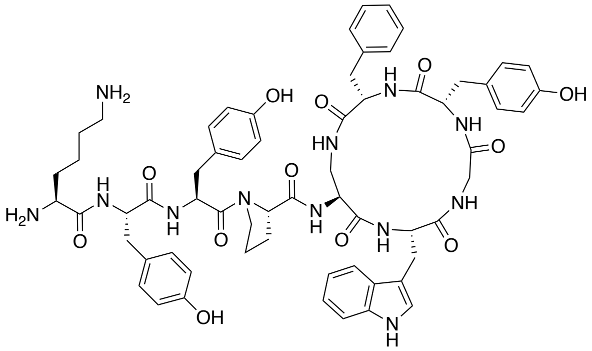

$C_{63}H_{74}N_{12}O_{12}$
MW: 1191.36

The peptide was synthesized as followed (Supporting Scheme S4). The fully protected linear peptide **S13** was synthesized in 40.0 μmol scale on Cl-Trt polystyrene resin **S7** preloaded with Fmoc-Phe-OH (0.69 mmol/g) using the general procedures for automated SPPS. After completed peptide elongation, the peptidyl-resin **S13** was transferred into a polypropylene syringe equipped with a fritted disk using $CH_2Cl_2$, washed with $CH_2Cl_2$ (3 × 1 min) and dried overnight under vacuum. The peptidyl-resin **S13** was treated twice with a solution of $Pd(PPh_3)_4$ (4.60 mg, 4.00 μmol, 0.1 equiv) and dimethylborane (12.0 mg, 0.20 mmol, 5.00 equiv) in anhydrous $CH_2Cl_2$ (3.0 mL) for 15 min at room temperature. The resin was then washed with $CH_2Cl_2$ (3 × 1 min), DMF (3 × 1 min) and $CH_2Cl_2$ (3 × 1 min) and dried under suction for 15 min. The dried resin **S14** was treated with a solution of HFIP (0.4 mL) in $CH_2Cl_2$ (1.6 mL) for 15 min at room temperature to cleave the partially protected peptide **S15** from the resin. The cleavage solution was removed from the resin and collected and a fresh HFIP–$CH_2Cl_2$ solution was added to the resin. After 15 min the cleavage solution was removed from the resin and collected and the resin rinsed with $CH_2Cl_2$ (2.0 mL). The combined cleavage solutions and the rinsing solution were evaporated to dryness under reduced pressure to yield the partially protected peptide **S15,** which was used without further purification. Crude peptide

**S15** (0.04 mmol based on the resin loading) was dissolved in anhydrous DMF (5.0 mL) under nitrogen atmosphere and added dropwise to a solution of HATU (15.2 mg, 0.04 mmol, 1.00 equiv) and *i*-$Pr_2NEt$ (20.8 µL, 0.12 mmol, 3.00 equiv) in anhydrous DMF (35.0 mL). The reaction mixture was stirred overnight at room temperature and after full consumption of **S15** was confirmed by UPLC-MS, the reaction was reduced to dryness under reduced pressure. The remaining residue was treated with a deprotection cocktail (3.0 mL, TFA–*i*-$Pr_3SiH$–water, 94:3:3, v/v/v) for 2 h and subsequently concentrated under a stream of nitrogen followed by precipitation in ice-cold diethyl ether. Purification by preparative RP-HPLC afforded *S. simulans* AIP-II lactam (**45**) as a fluffy white solid after lyophilization. Purity >95% determined by UPLC ($\lambda = 215$ nm). **UPLC-MS** (ESI) *m/z* calcd for $[M+H]^+$ $C_{63}H_{75}N_{12}O_{12}^+$: 1191.56, found 1191.72.

***S. simulans*** **AIP-III K1A (46)**

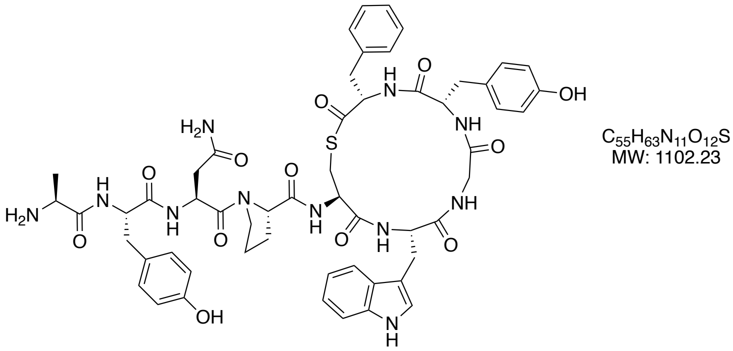

The peptide was synthesized according to general method A for on-resin cleavage-inducing cyclization (section 4.5). Purification by preparative RP-HPLC afforded *S. simulans* AIP-III K1A (**46**) as a fluffy white solid after lyophilization. Purity >95% determined by UPLC ($\lambda = 215$ nm). **UPLC-MS** (ESI) *m/z* calcd for $[M+H]^+$ $C_{55}H_{64}N_{11}O_{12}S^+$: 1102.45, found 1102.42.

***S. simulans*** **AIP-III Y2A (47)**

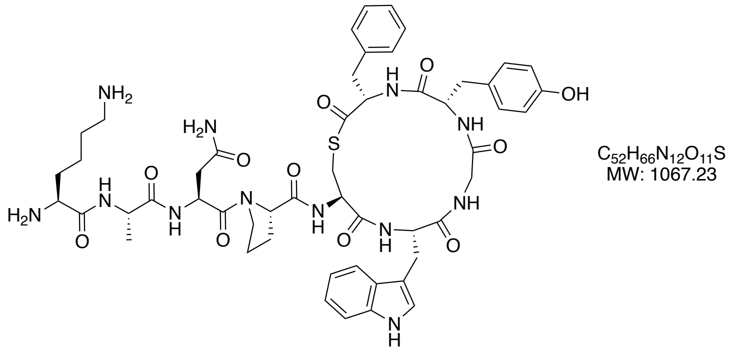

The peptide was synthesized according to general method A for on-resin cleavage-inducing cyclization (section 4.5). Purification by preparative RP-HPLC afforded *S. simulans* AIP-III Y2A (**47**) as a fluffy white solid after lyophilization. Purity >95% determined by UPLC ($\lambda = 215$ nm). **UPLC-MS** (ESI) *m/z* calcd for $[M+H]^+$ $C_{52}H_{67}N_{12}O_{11}S^+$: 1067.48, found 1067.50.

***S. simulans*** **AIP-III P4A (48)**

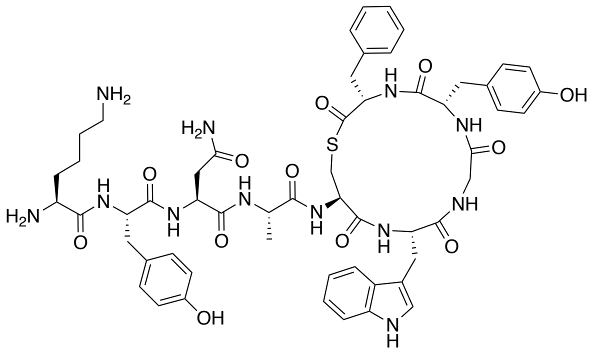

$C_{56}H_{68}N_{12}O_{12}S$
MW: 1133.29

The peptide was synthesized according to general method A for on-resin cleavage-inducing cyclization (section 4.5). Purification by preparative RP-HPLC afforded *S. simulans* AIP-III P4A (**48**) as a fluffy white solid after lyophilization. Purity >95% determined by UPLC ($\lambda$ = 215 nm). **UPLC-MS** (ESI) *m/z* calcd for $[M+H]^+$ $C_{56}H_{69}N_{12}O_{12}S^+$: 1133.49, found 1133.67.

***S. simulans* AIP-III W6A (49)**

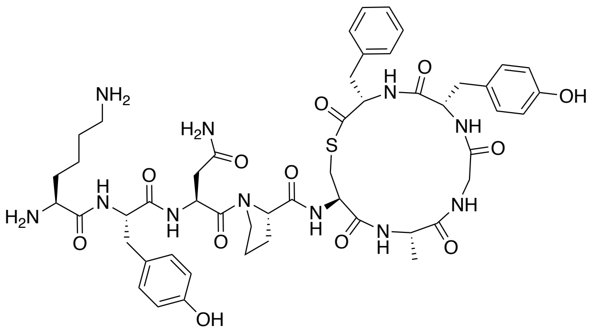

$C_{50}H_{65}N_{11}O_{12}S$
MW: 1044.20

The peptide was synthesized according to general method A for on-resin cleavage-inducing cyclization (section 4.5). Purification by preparative RP-HPLC afforded *S. simulans* AIP-III W6A (**49**) as a fluffy white solid after lyophilization. Purity >95% determined by UPLC ($\lambda$ = 215 nm). **UPLC-MS** (ESI) *m/z* calcd for $[M+H]^+$ $C_{50}H_{66}N_{11}O_{12}S^+$: 1044.46, found 1044.48.

***S. simulans* AIP-III G7A (50)**

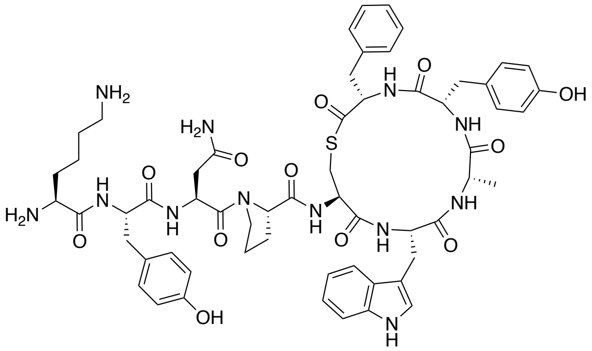

$C_{59}H_{72}N_{12}O_{12}S$
MW: 1173.36

The peptide was synthesized according to general method A for on-resin cleavage-inducing cyclization (section 4.5). Purification by preparative RP-HPLC afforded *S. simulans* AIP-III G7A (**50**) as a fluffy white solid after lyophilization. Purity >95% determined by UPLC ($\lambda$ = 215 nm). **UPLC-MS** (ESI) *m/z* calcd for $[M+H]^+$ $C_{59}H_{73}N_{12}O_{12}S^+$: 1173.52, found 1173.51.

***S. simulans* AIP-III Y8A (51)**

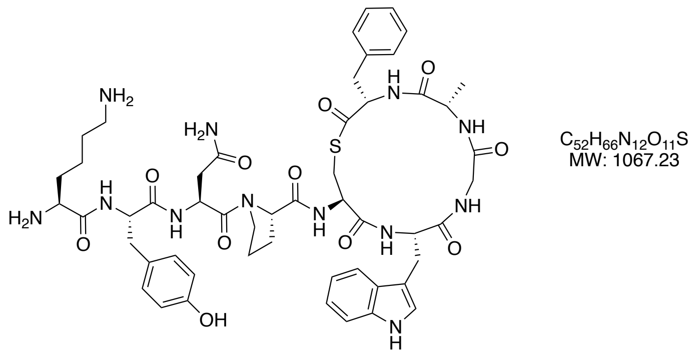

The peptide was synthesized according to general method A for on-resin cleavage-inducing cyclization (section 4.5). Purification by preparative RP-HPLC afforded *S. simulans* AIP-III Y8A (**51**) as a fluffy white solid after lyophilization. Purity >95% determined by UPLC ($\lambda$ = 215 nm). **UPLC-MS** (ESI) *m/z* calcd for $[M+H]^+$ $C_{52}H_{67}N_{12}O_{11}S^+$: 1067.48, found 1067.61.

***S. simulans*** **AIP-III F9A (52)**

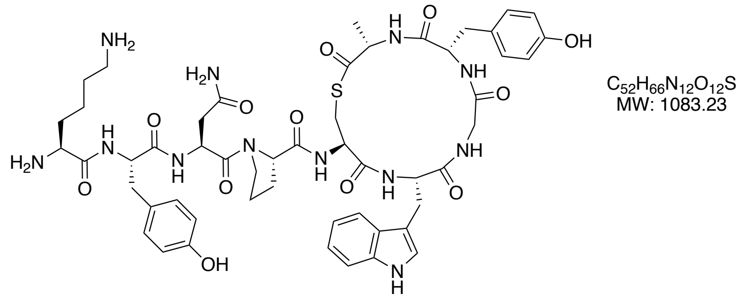

The peptide was synthesized according to general method A for on-resin cleavage-inducing cyclization (section 4.5). Purification by preparative RP-HPLC afforded *S. simulans* AIP-III F9A (**52**) as a fluffy white solid after lyophilization. Purity 93% determined by UPLC ($\lambda$ = 215 nm). **UPLC-MS** (ESI) *m/z* calcd for $[M+H]^+$ $C_{52}H_{67}N_{12}O_{12}S^+$: 1083.47, found 1083.52.

***S. simulans*** **AIP-III 8-mer (53)**

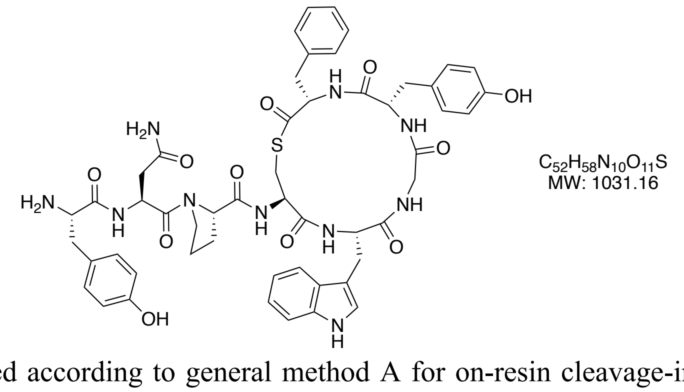

The peptide was synthesized according to general method A for on-resin cleavage-inducing cyclization (section 4.5). Purification by preparative RP-HPLC afforded *S. simulans* AIP-III 8-mer (**53**) as a fluffy white solid after lyophilization. Purity >95% determined by UPLC ($\lambda$ = 215 nm). **UPLC-MS** (ESI) *m/z* calcd for $[M+H]^+$ $C_{52}H_{59}N_{10}O_{11}S^+$: 1031.41, found 1031.67.

***S. simulans*** **AIP-III 7-mer (54)**

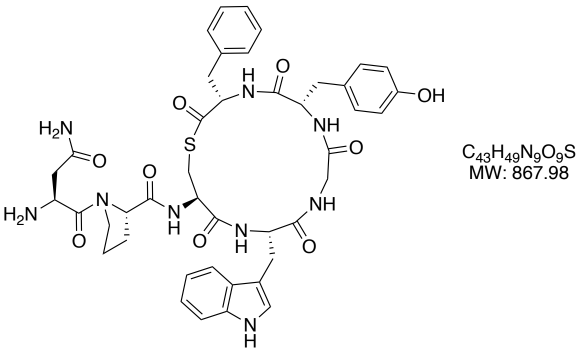

The peptide was synthesized according to general method A for on-resin cleavage-inducing cyclization (section 4.5). Purification by preparative RP-HPLC afforded *S. simulans* AIP-III 7-mer (**54**) as a fluffy white solid after lyophilization. Purity >95% determined by UPLC ($\lambda = 215$ nm). **UPLC-MS** (ESI) *m/z* calcd for $[M+H]^+$ $C_{43}H_{50}N_9O_9S^+$: 868.34, found 868.53.

***S. simulans*** **AIP-II/III 6-mer (55)**

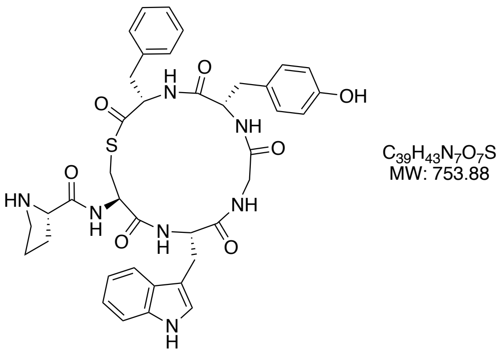

The peptide was synthesized according to general method A for on-resin cleavage-inducing cyclization (section 4.5). Purification by preparative RP-HPLC afforded *S. simulans* AIP-III 6-mer (**55**) as a fluffy white solid after lyophilization. Purity >95% determined by UPLC ($\lambda = 215$ nm). **UPLC-MS** (ESI) *m/z* calcd for $[M+H]^+$ $C_{39}H_{44}N_7O_7S^+$: 754.30, found 754.48.

***S. simulans*** **AIP-II/III 5-mer *N*-Ac (56)**

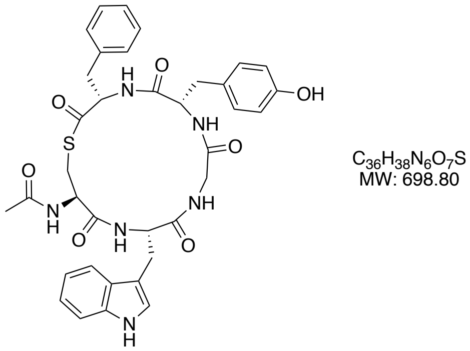

The peptide was synthesized according to general method A for on-resin cleavage-inducing cyclization (section 4.5). Purification by preparative RP-HPLC afforded *S. simulans* AIP-II/III 5-mer *N*-Ac (**56**) as a fluffy white solid after lyophilization. Purity >95% determined by UPLC ($\lambda = 215$ nm). **UPLC-MS** (ESI) *m/z* calcd for $[M+H]^+$ $C_{36}H_{39}N_6O_7S^+$: 699.26, found 699.41.

***S. simulans*** **AIP-II/III 5-mer *N*-Me₂ (57)**

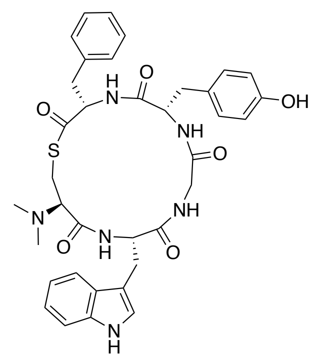

$C_{36}H_{40}N_6O_6S$
MW: 684.81

The peptide was synthesized according to general method A for on-resin cleavage-inducing cyclization (section 4.5). The amino acid $Me_2N$-Cys(Trt)-OH[25] was used for the synthesis. Purification by preparative RP-HPLC afforded *S. simulans* AIP-II/III 5-mer *N*-$Me_2$ (**57**) as a fluffy white solid after lyophilization. Purity >95% determined by UPLC ($\lambda$ = 215 nm). **UPLC-MS** (ESI) *m/z* calcd for $[M+H]^+$ $C_{36}H_{41}N_6O_6S^+$: 685.28, found 685.42.

### 6. UPLC traces of synthetic peptides

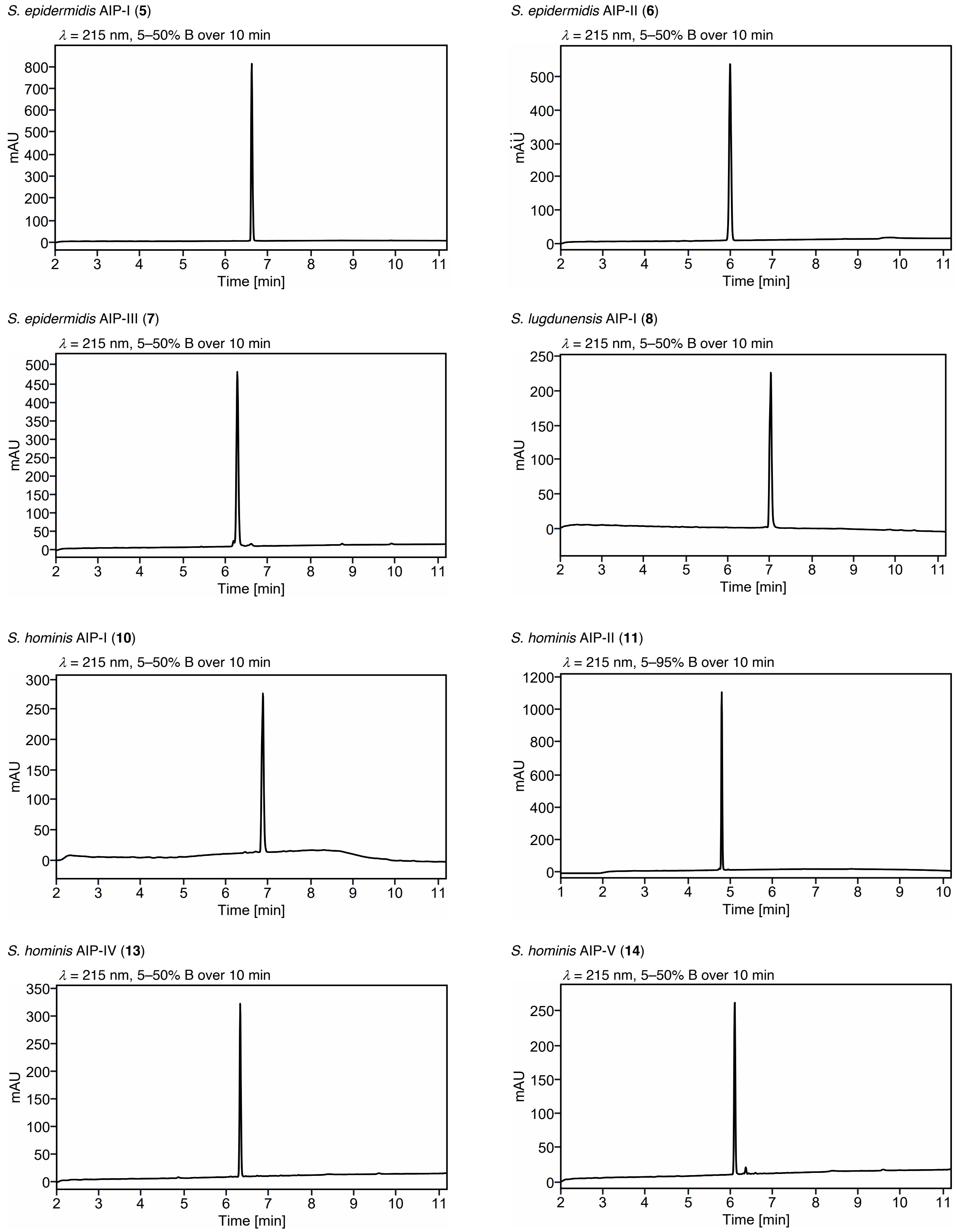

**Supplementary Figure S36.** UPLC traces of purified peptides.

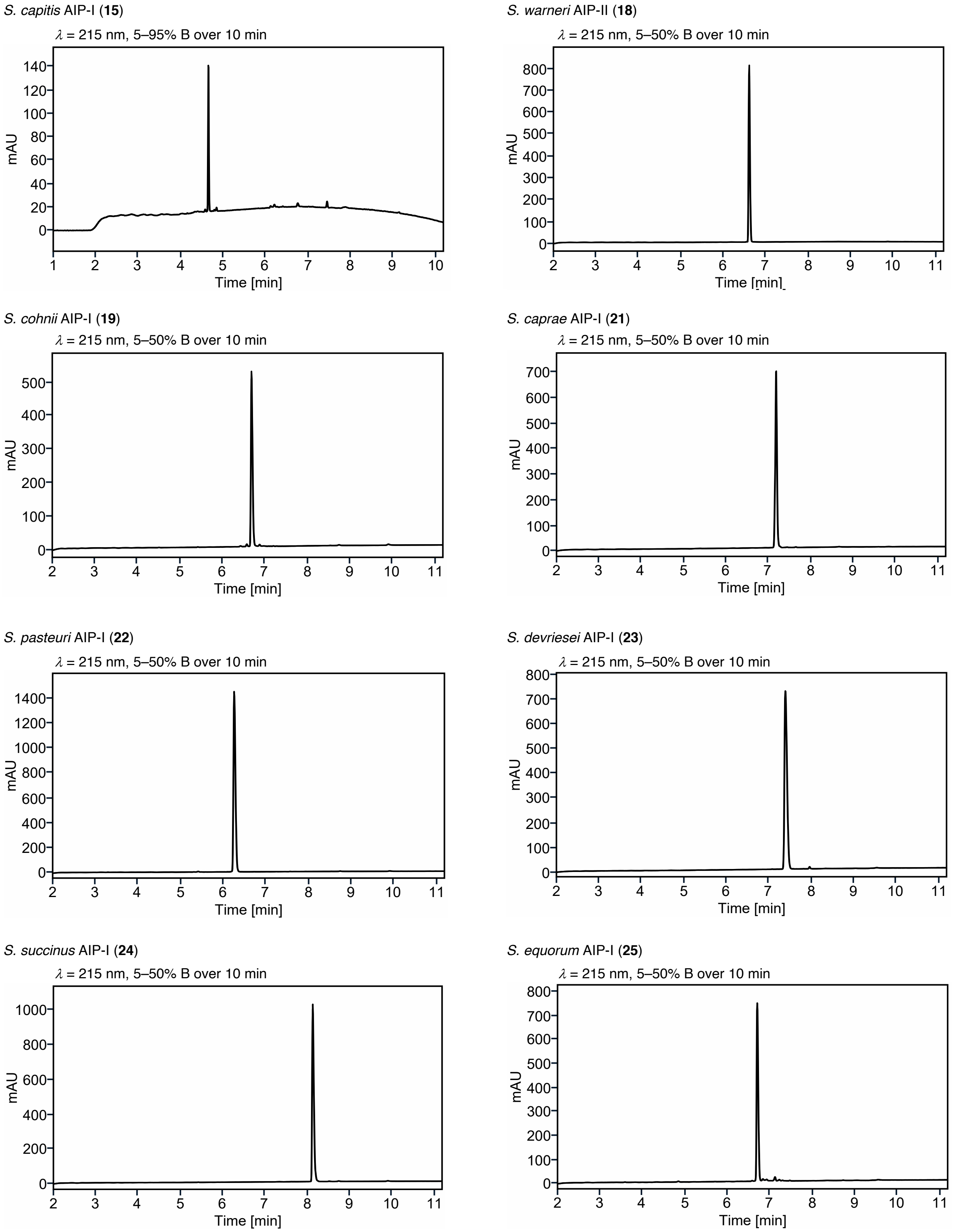

**Supplementary Figure S37.** UPLC traces of purified peptides.

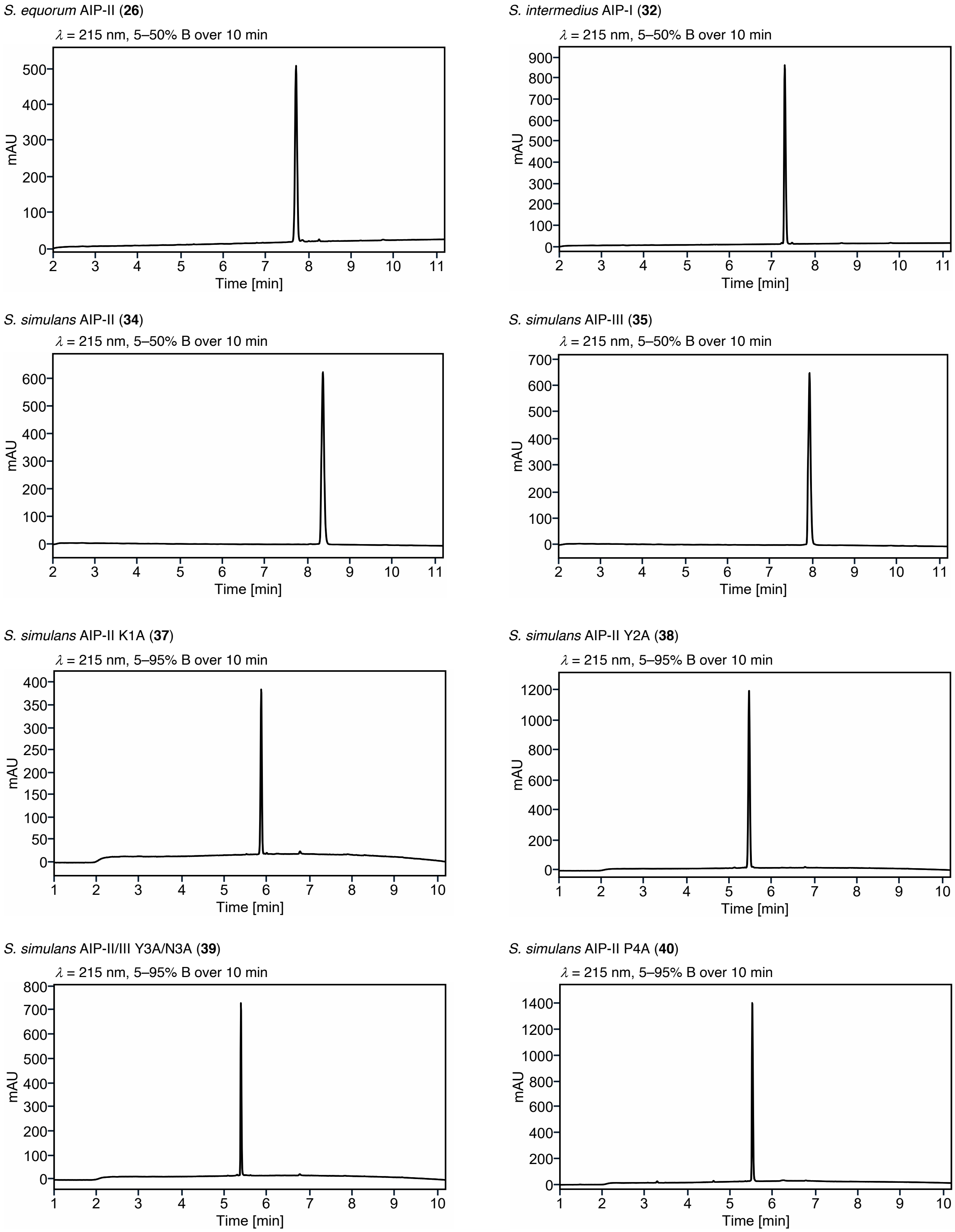

**Supplementary Figure S38.** UPLC traces of purified peptides.

*S. simulans* AIP-II W6A (**41**)

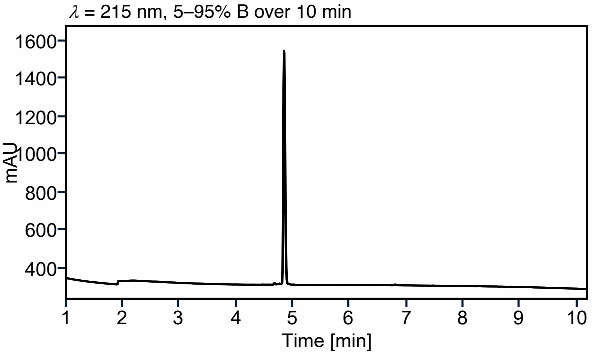

*S. simulans* AIP-II G7A (**42**)

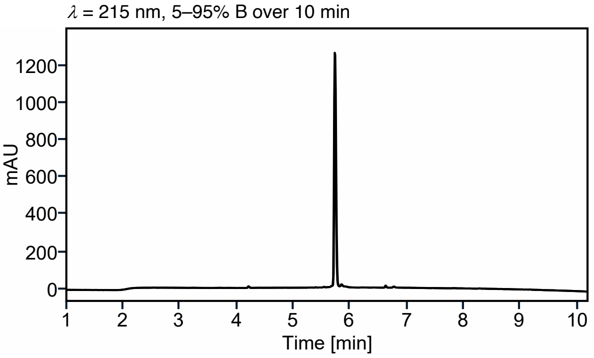

*S. simulans* AIP-II Y8A (**43**)

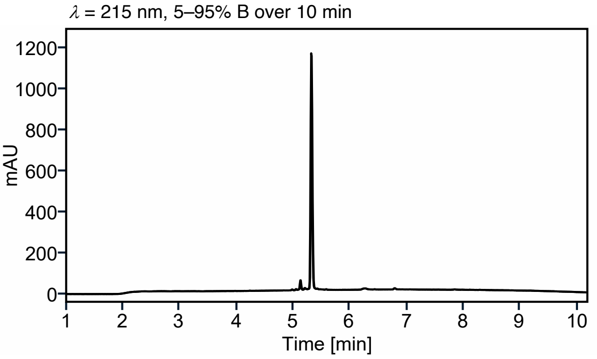

*S. simulans* AIP-II F9A (**44**)

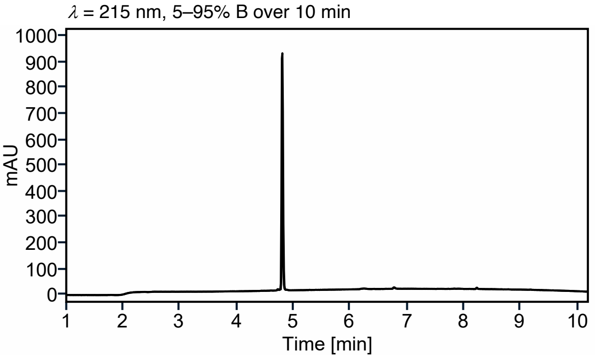

*S. simulans* AIP-II lactam (**45**)

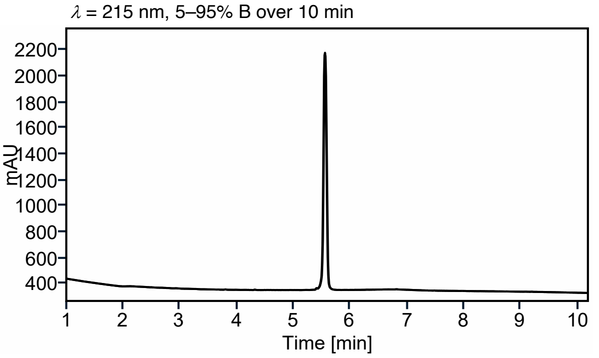

*S. simulans* AIP-III K1A (**46**)

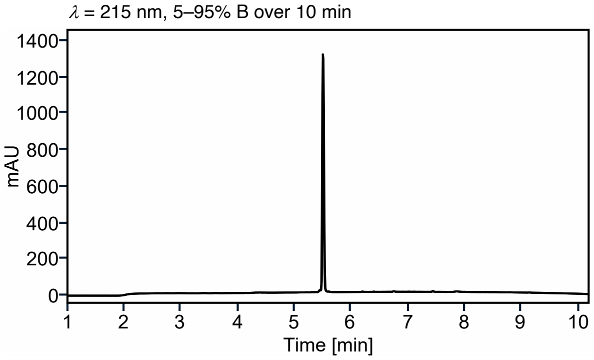

*S. simulans* AIP-III Y2A (**47**)

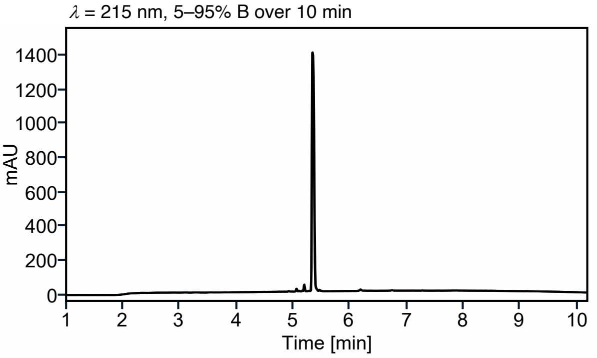

*S. simulans* AIP-III P4A (**48**)

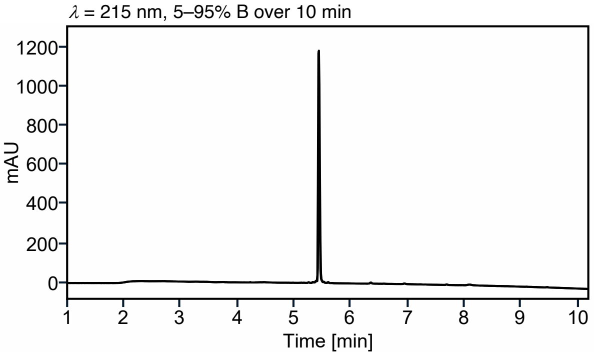

**Supplementary Figure S39.** UPLC traces of purified peptides.

*S. simulans* AIP-III W6A (**49**)

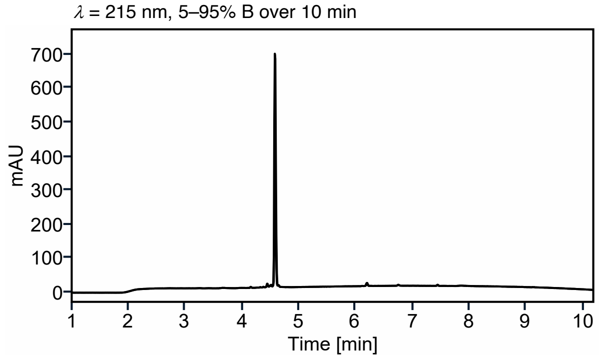

*S. simulans* AIP-III G7A (**50**)

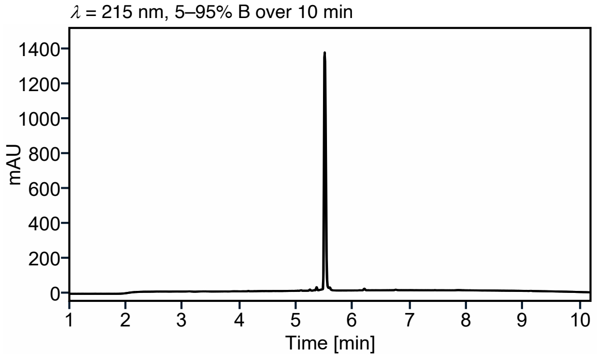

*S. simulans* AIP-III Y8A (**51**)

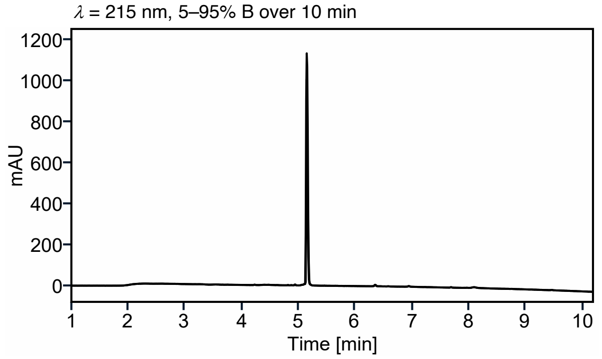

*S. simulans* AIP-III F9A (**52**)

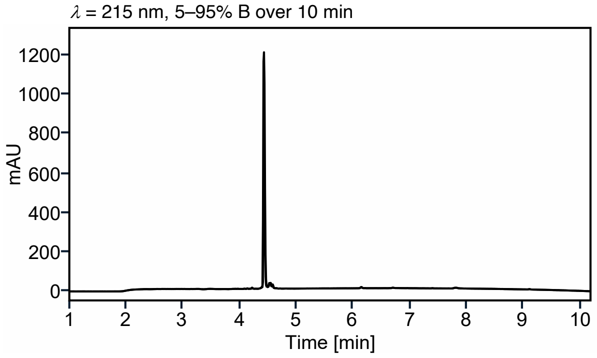

*S. simulans* AIP-III 8-mer (**53**)

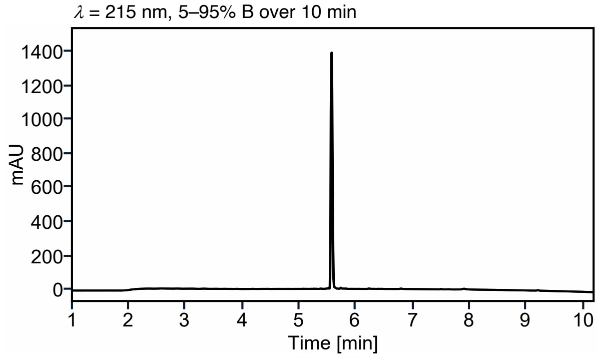

*S. simulans* AIP-III 7-mer (**54**)

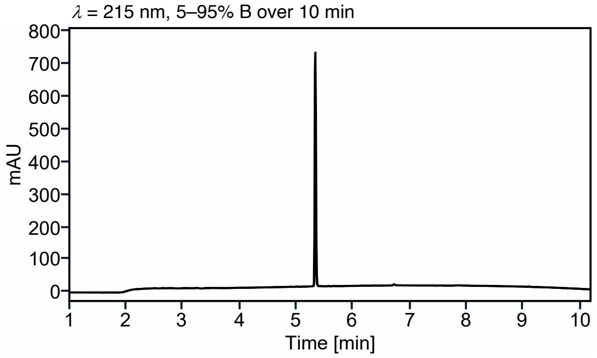

*S. simulans* AIP-II/III 6-mer (**55**)

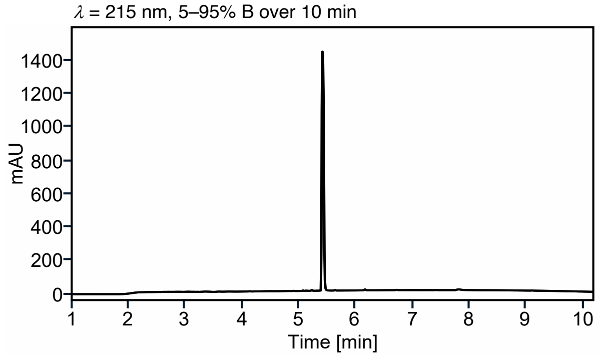

*S. simulans* AIP-II/III 5-mer *N*-Ac (**56**)

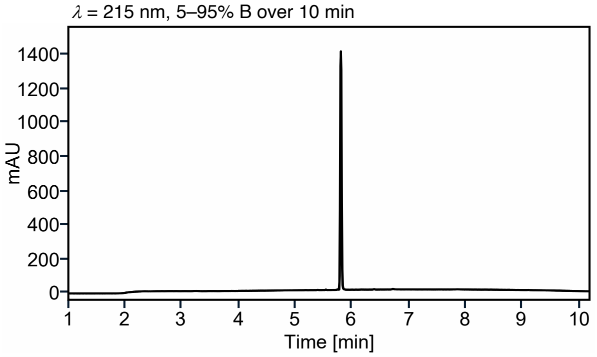

**Supplementary Figure S40.** UPLC traces of purified peptides.

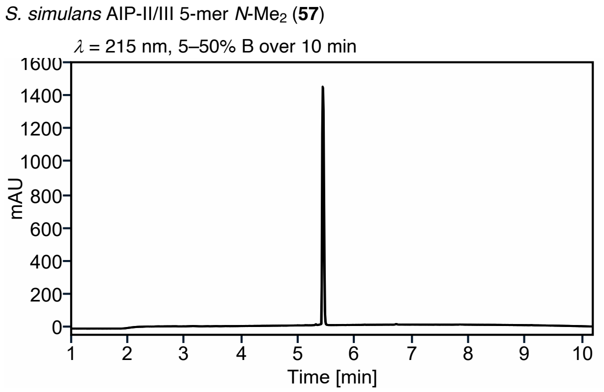

**Supplementary Figure S41.** UPLC traces of purified peptides.

### 7. Copies of NMR spectra

***S. epidermidis* AIP-I (5)**

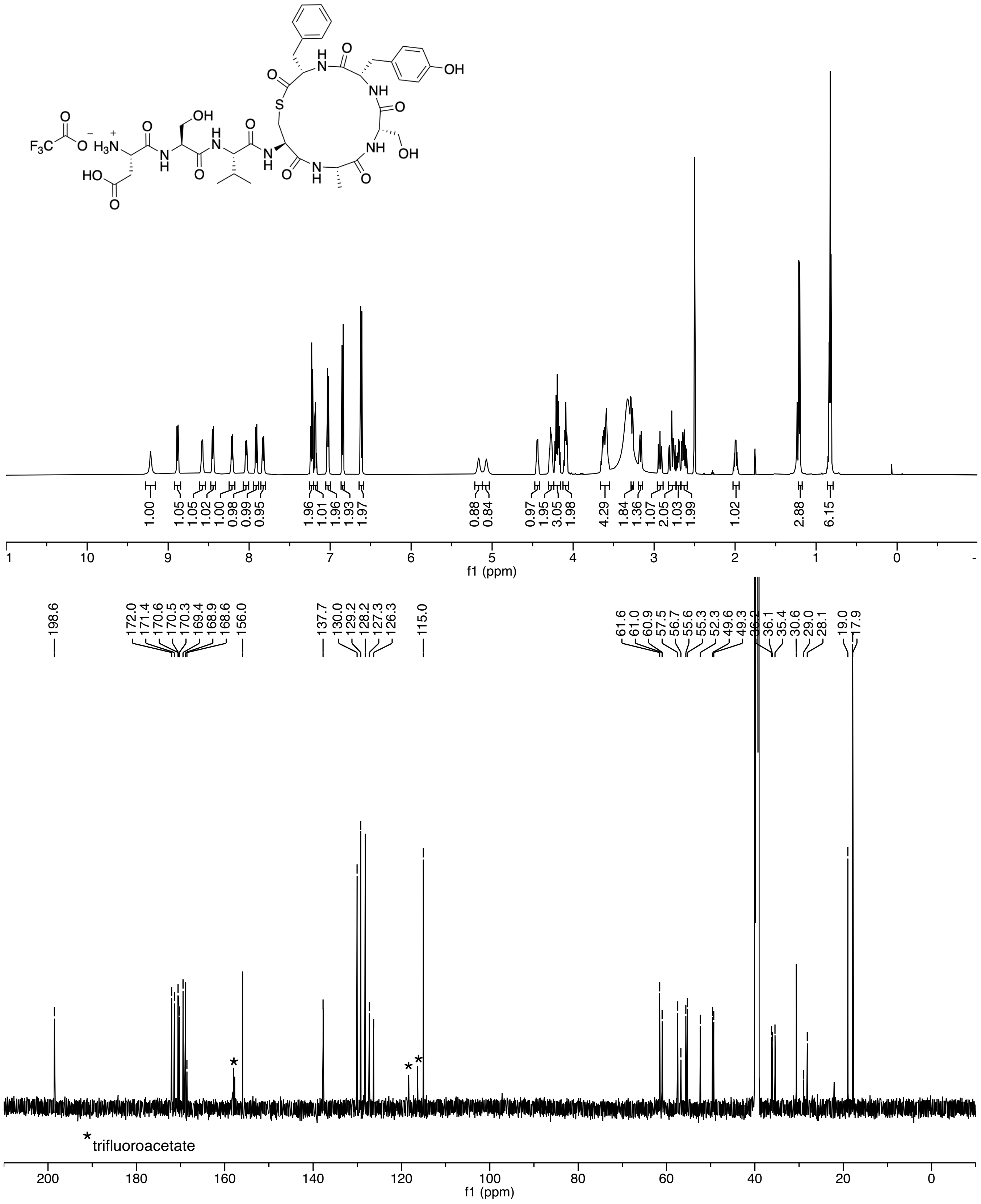

***S. epidermidis* AIP-II (6)**

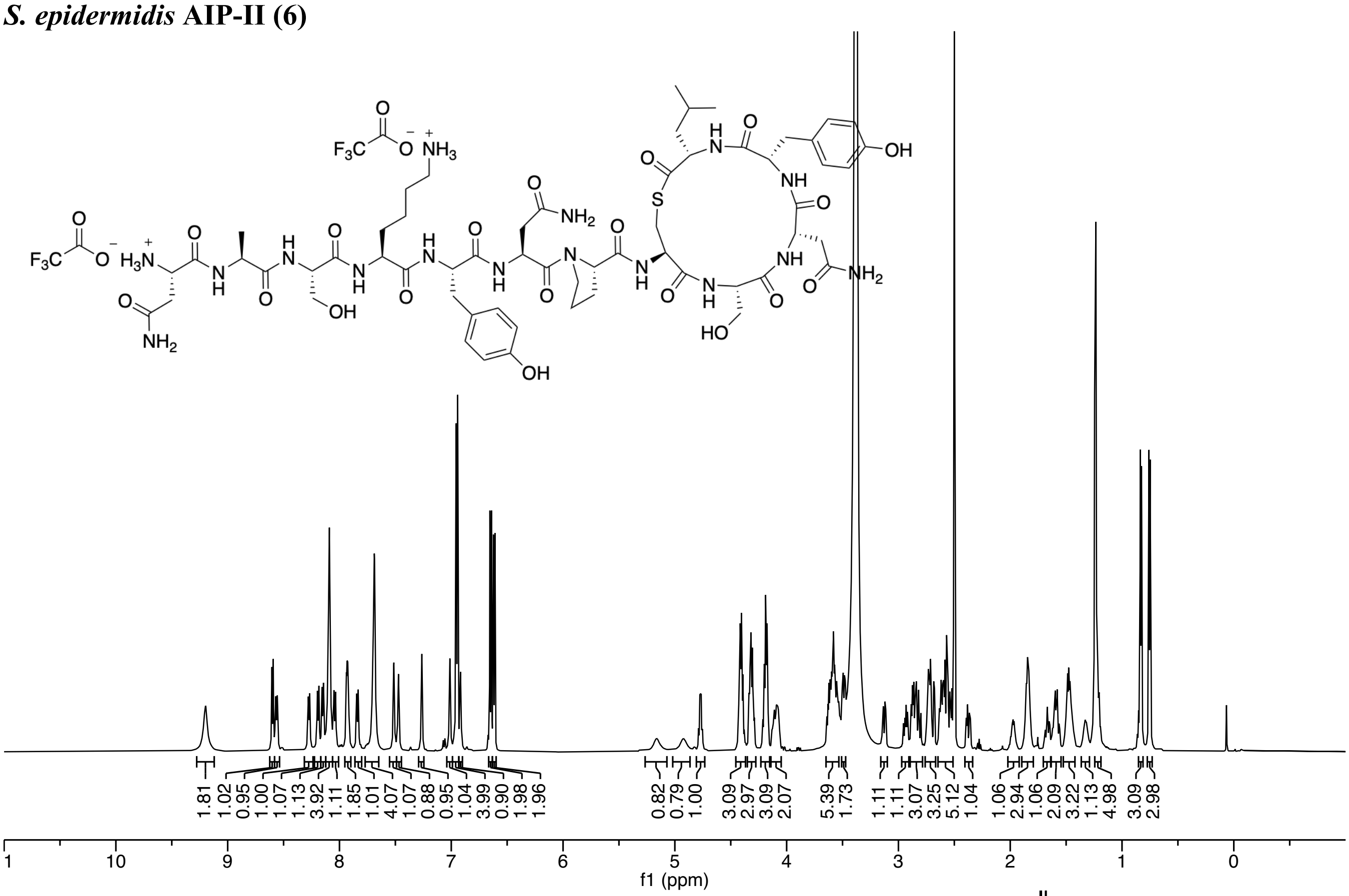

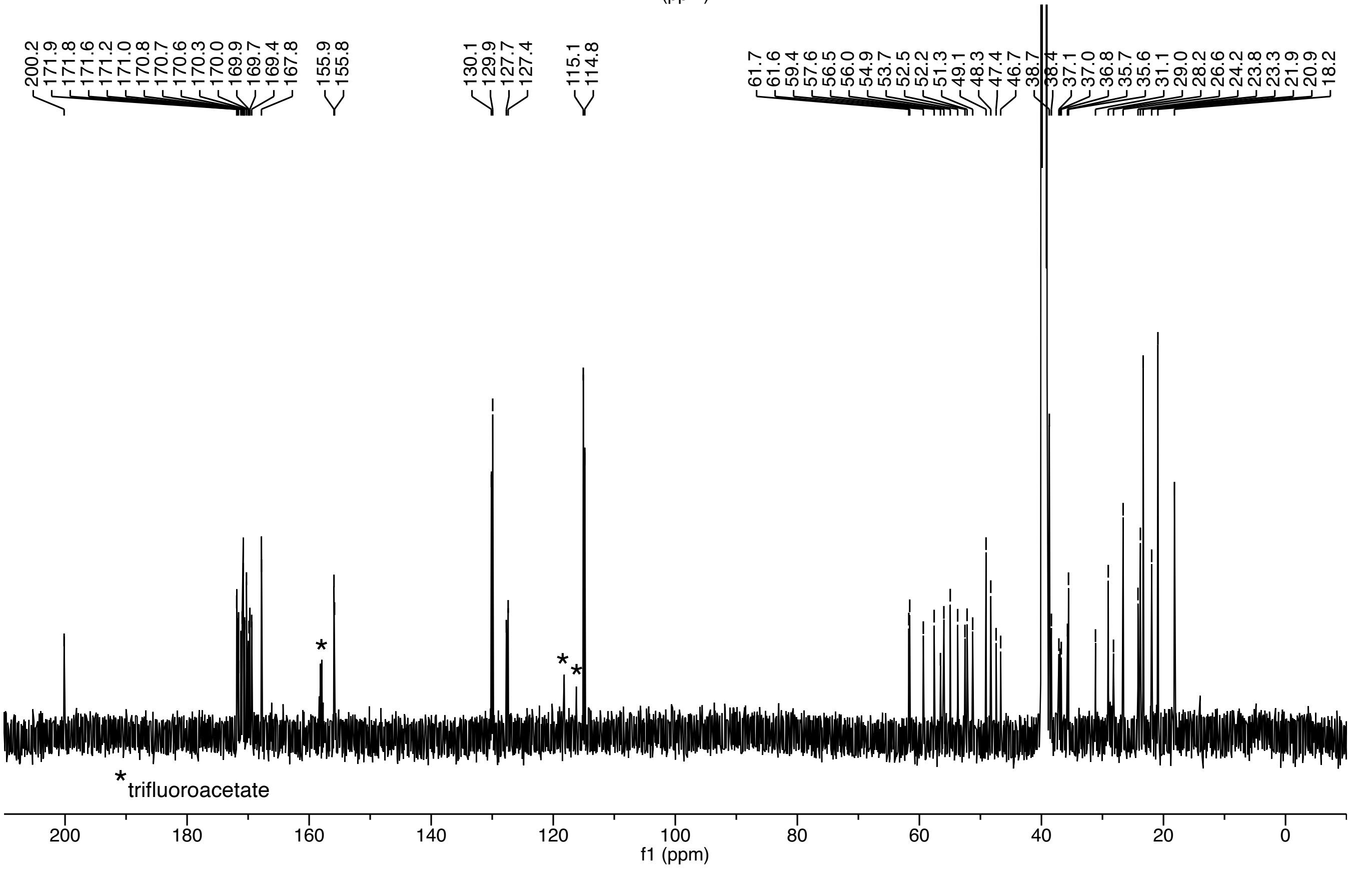

***S. epidermidis* AIP-III (7)**

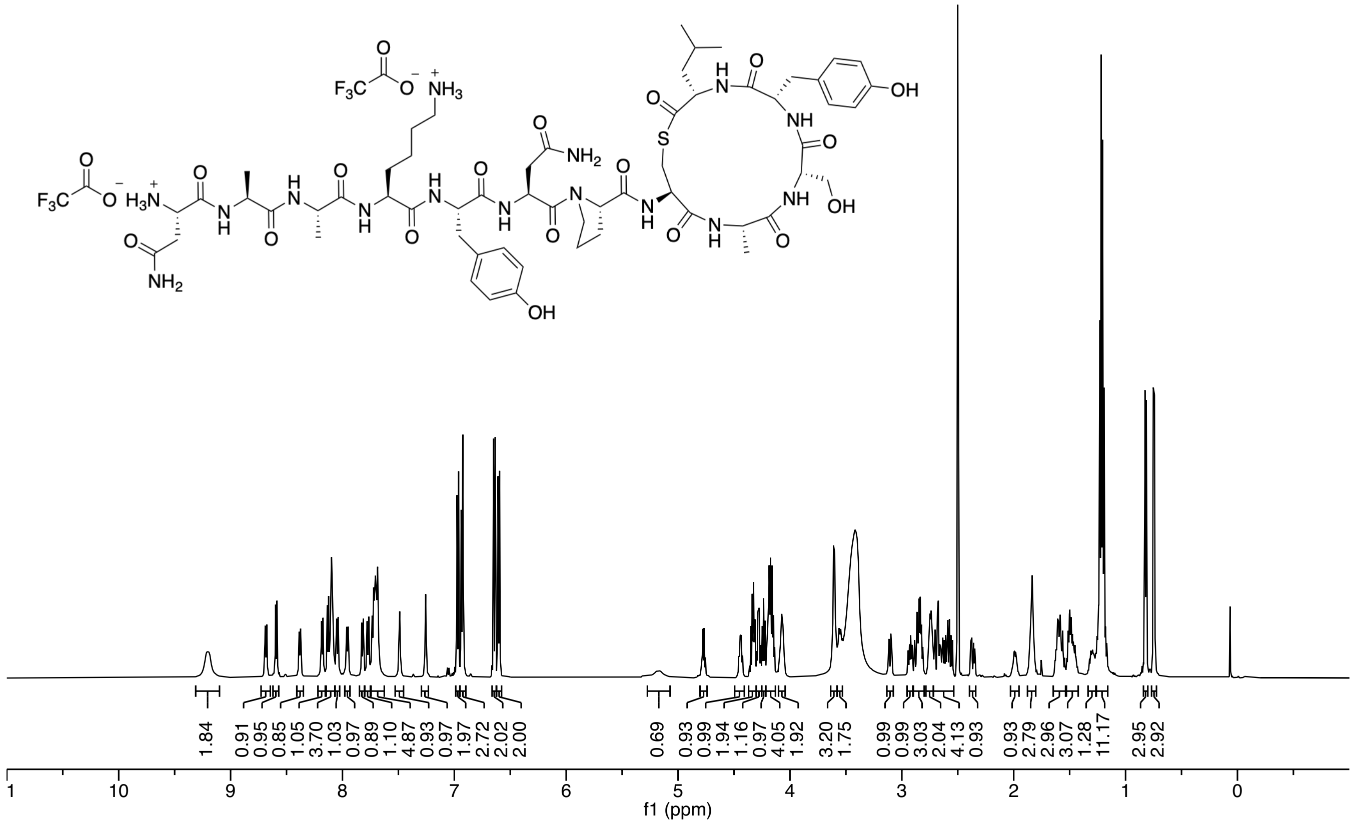

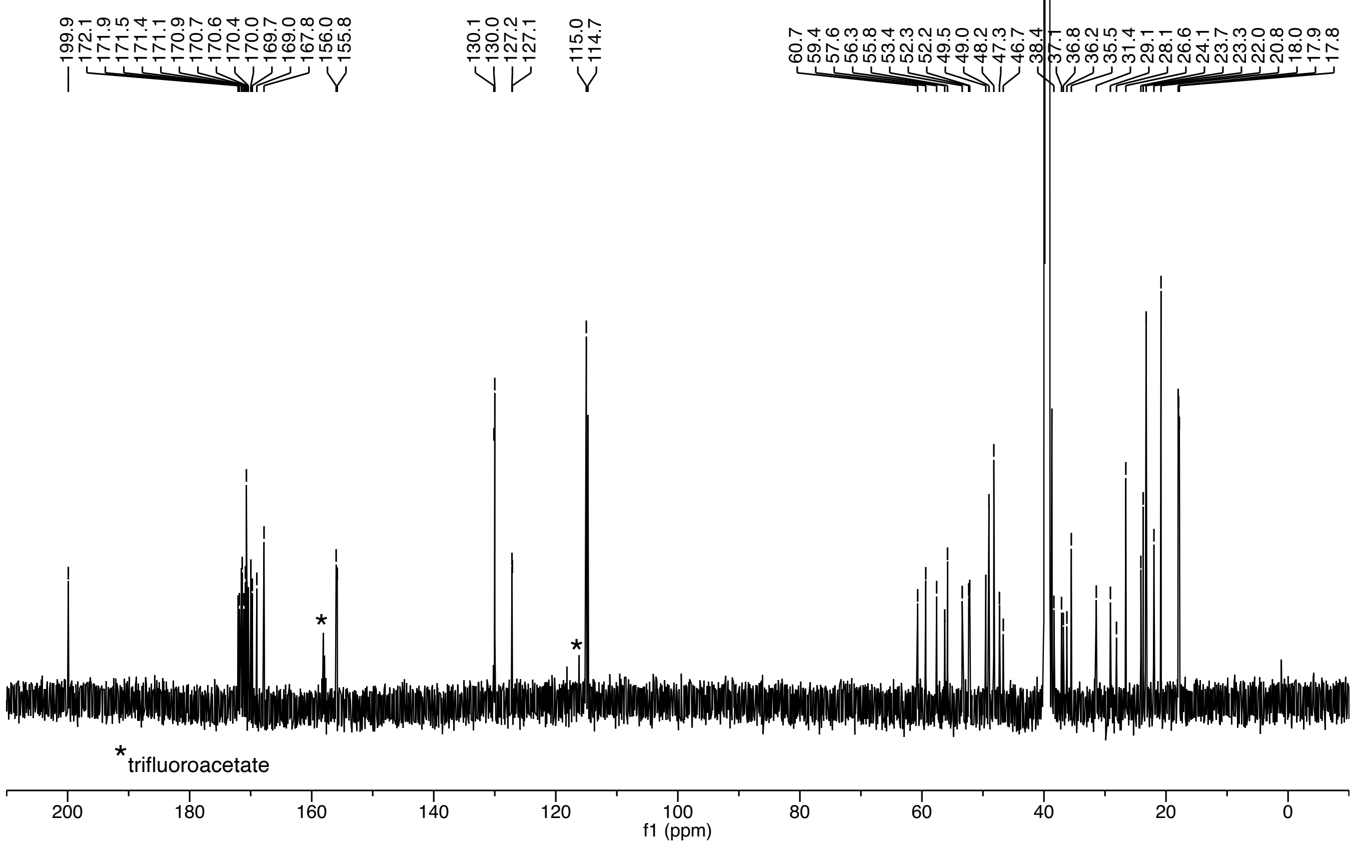

***S. lugdunensis* AIP-I (8)**

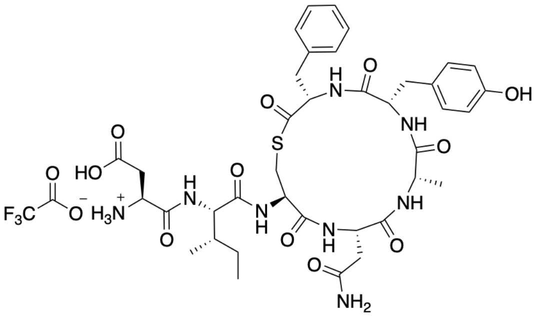

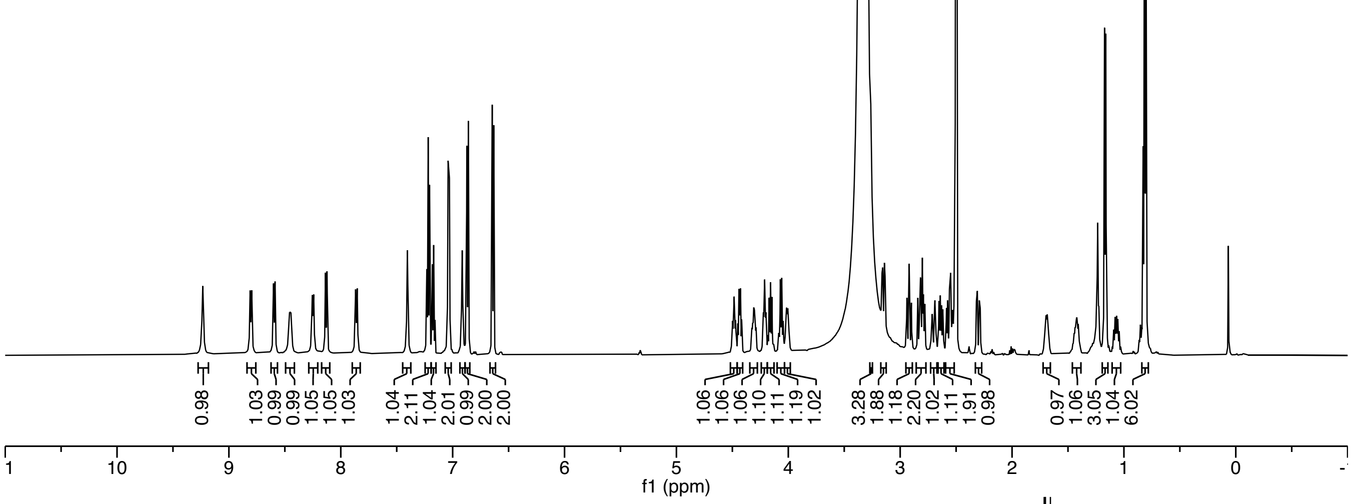

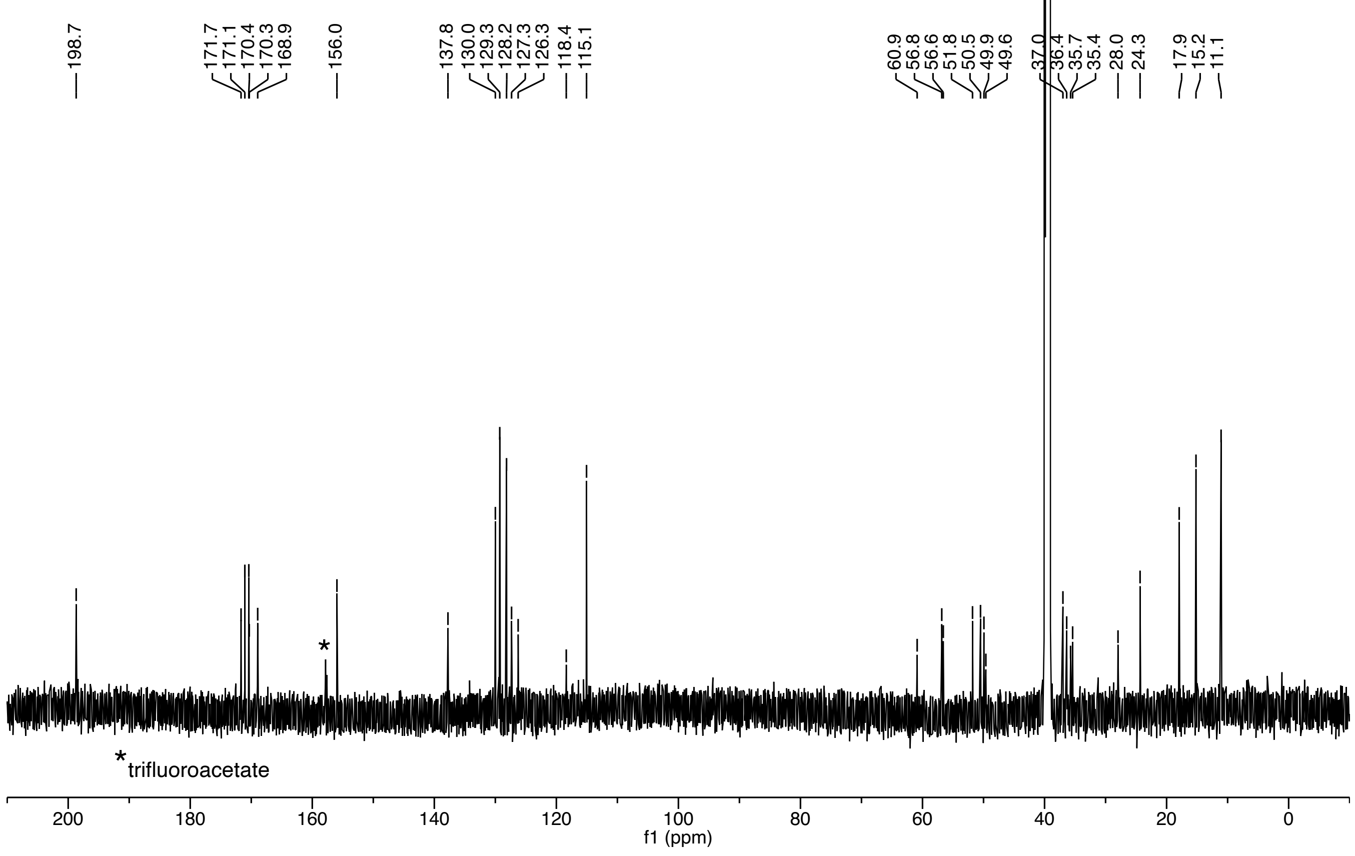

***S. hominis* AIP-I (10)**

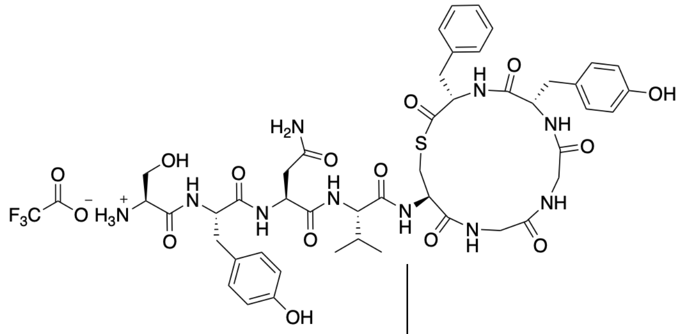

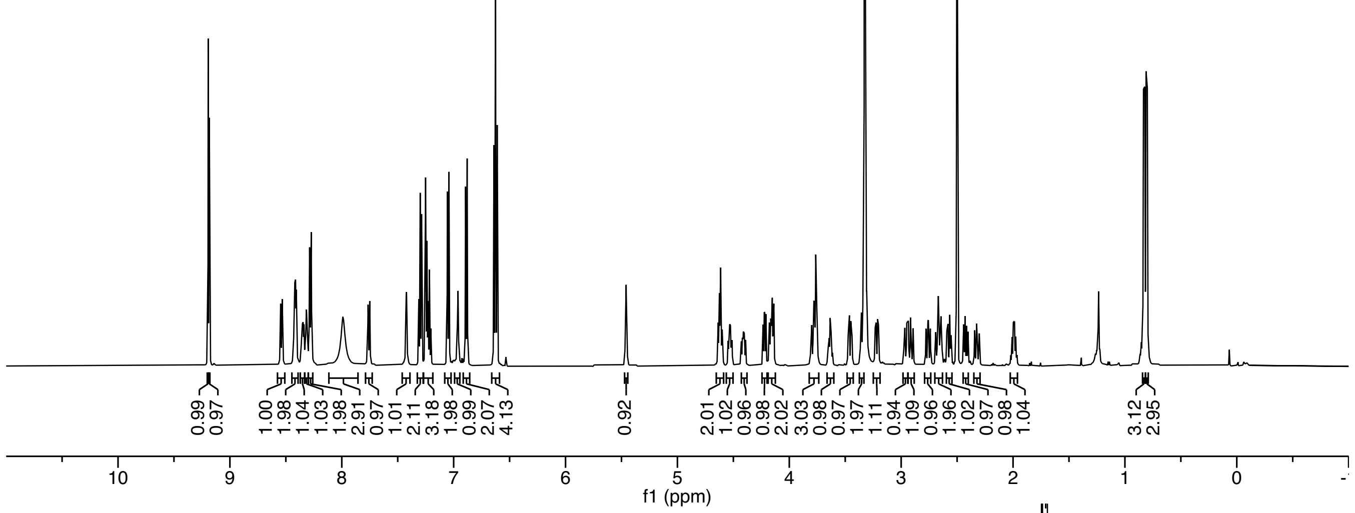

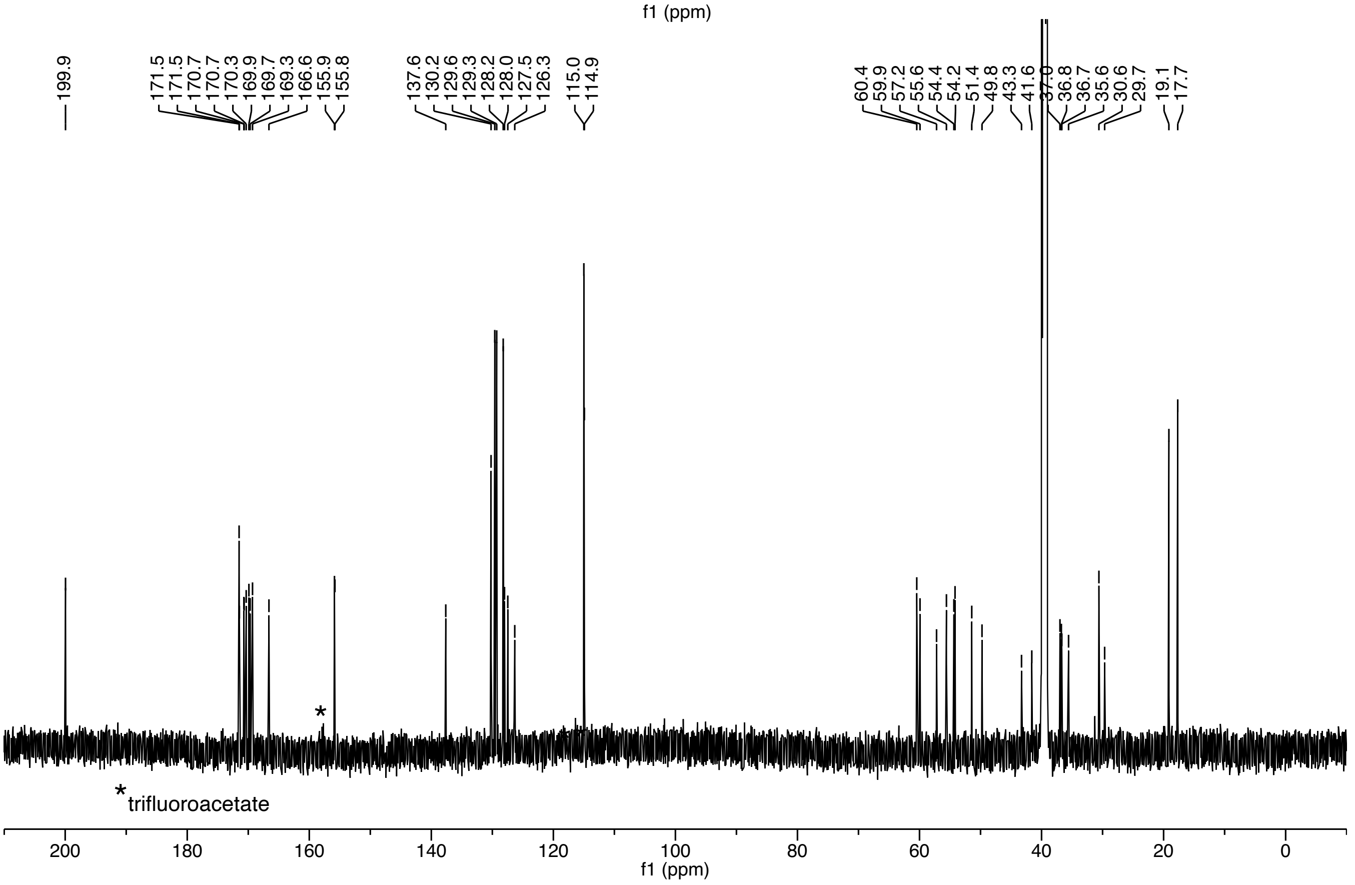

*S. hominis* **AIP-II (11)**

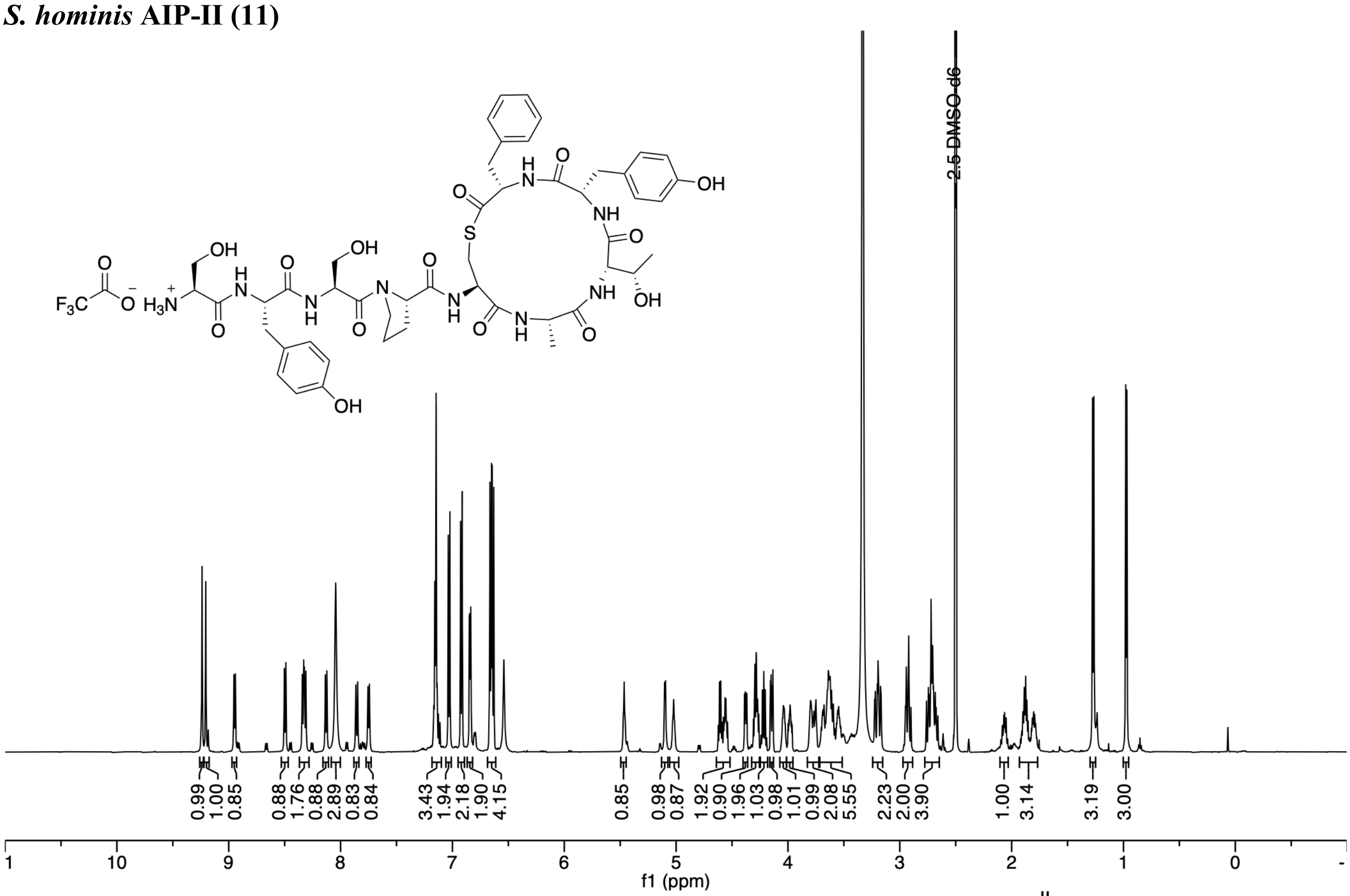

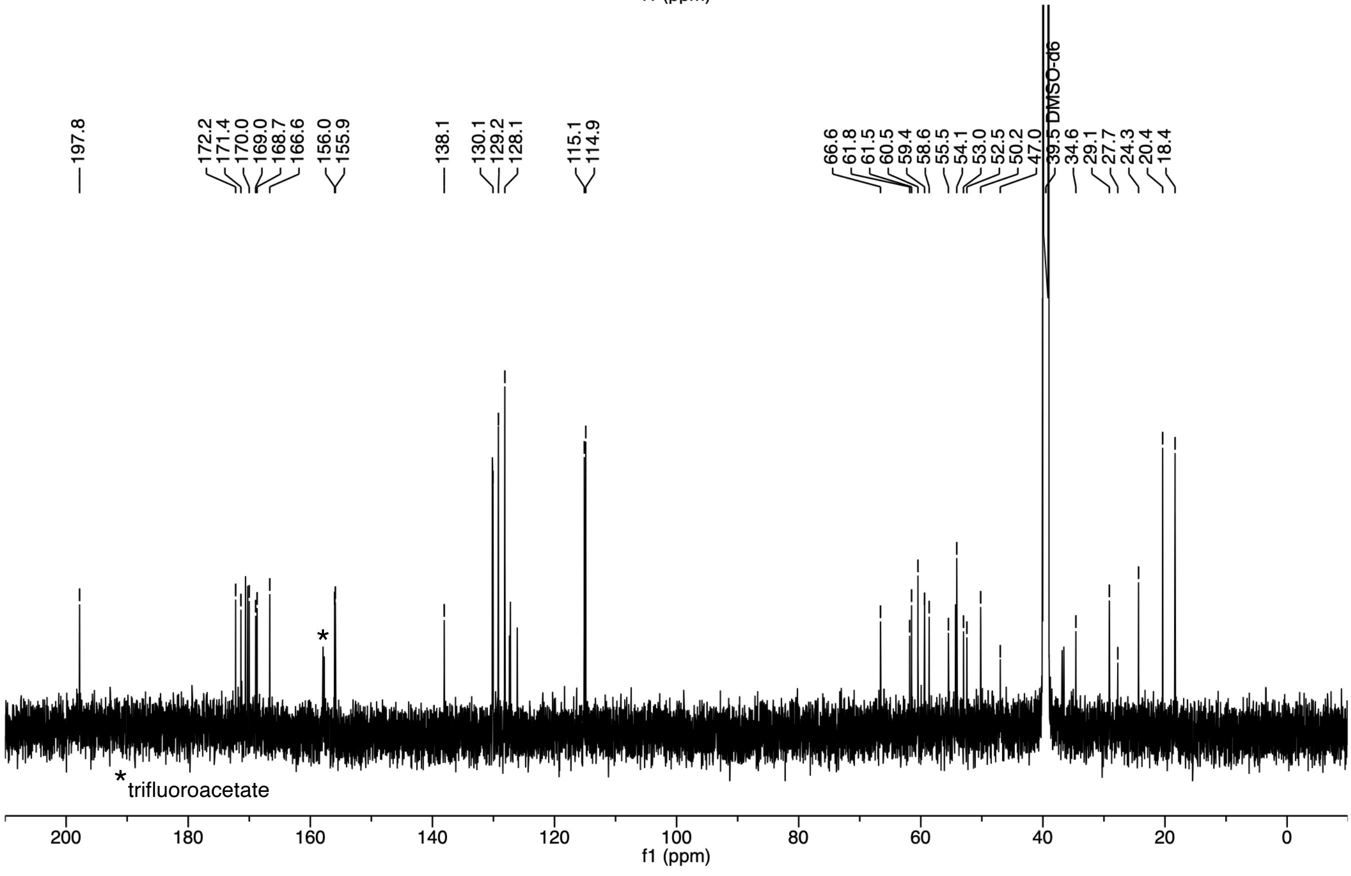

***S. hominis* AIP-IV (13)**

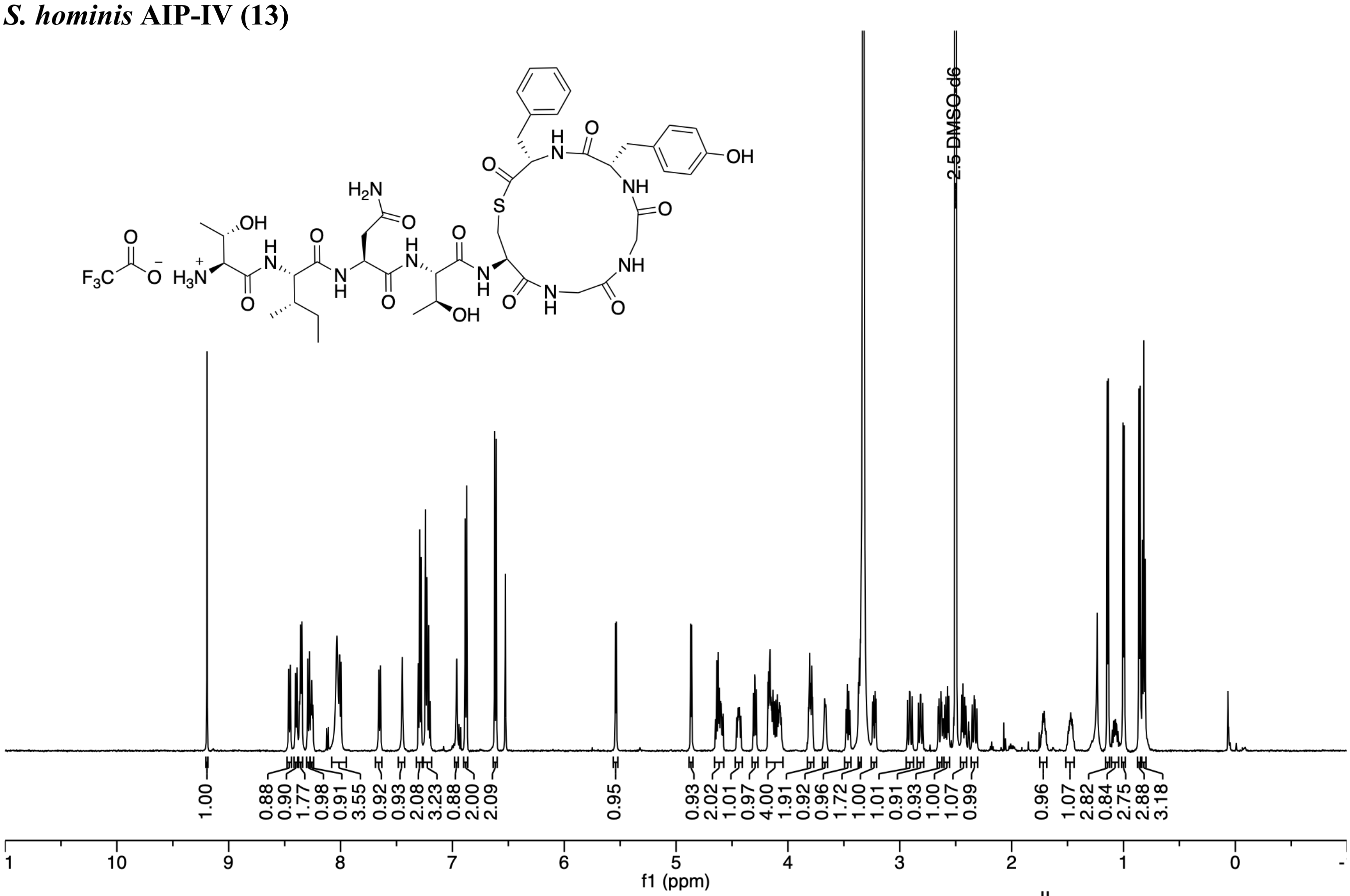

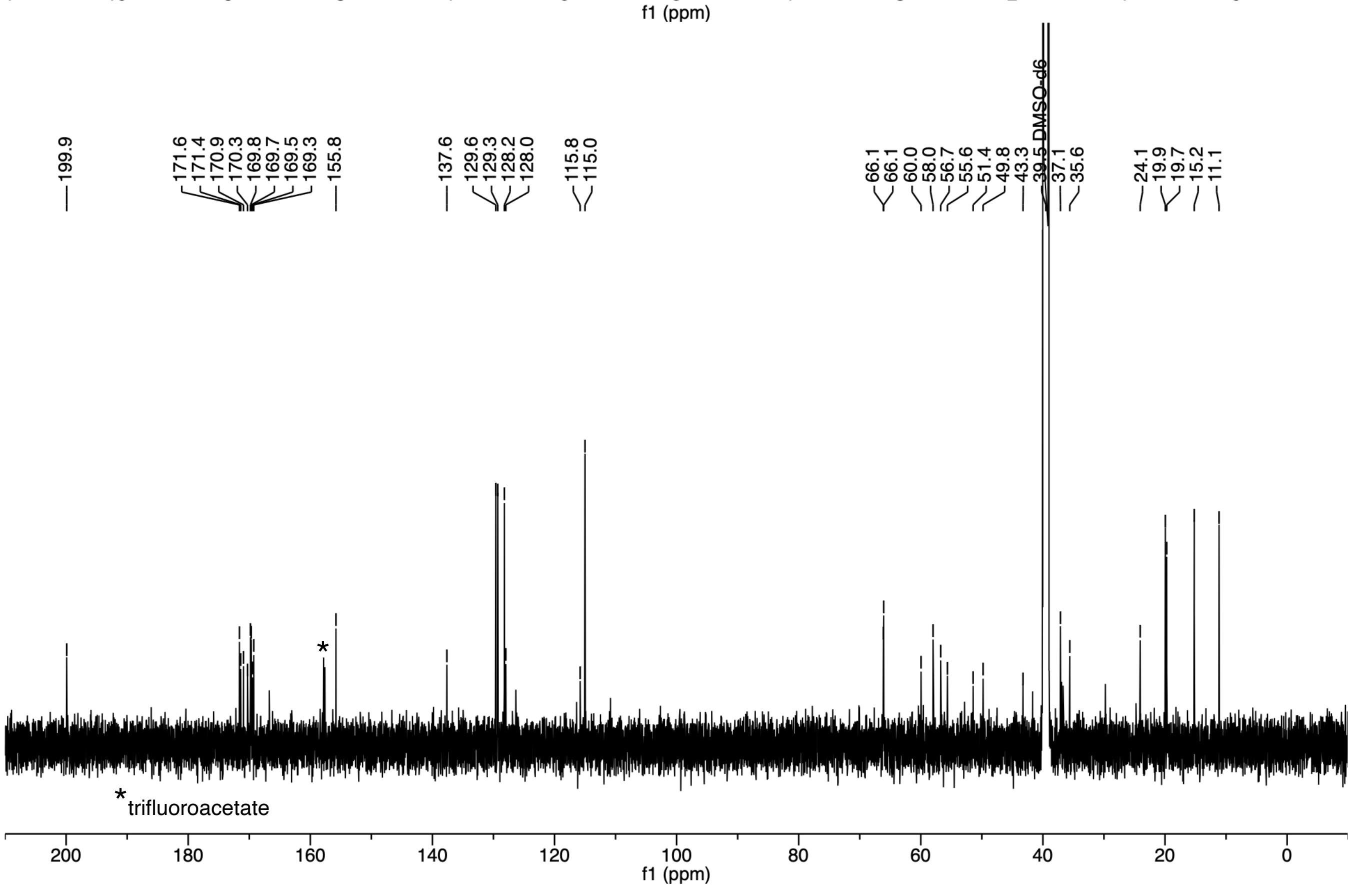

***S. hominis* AIP-V (14)**

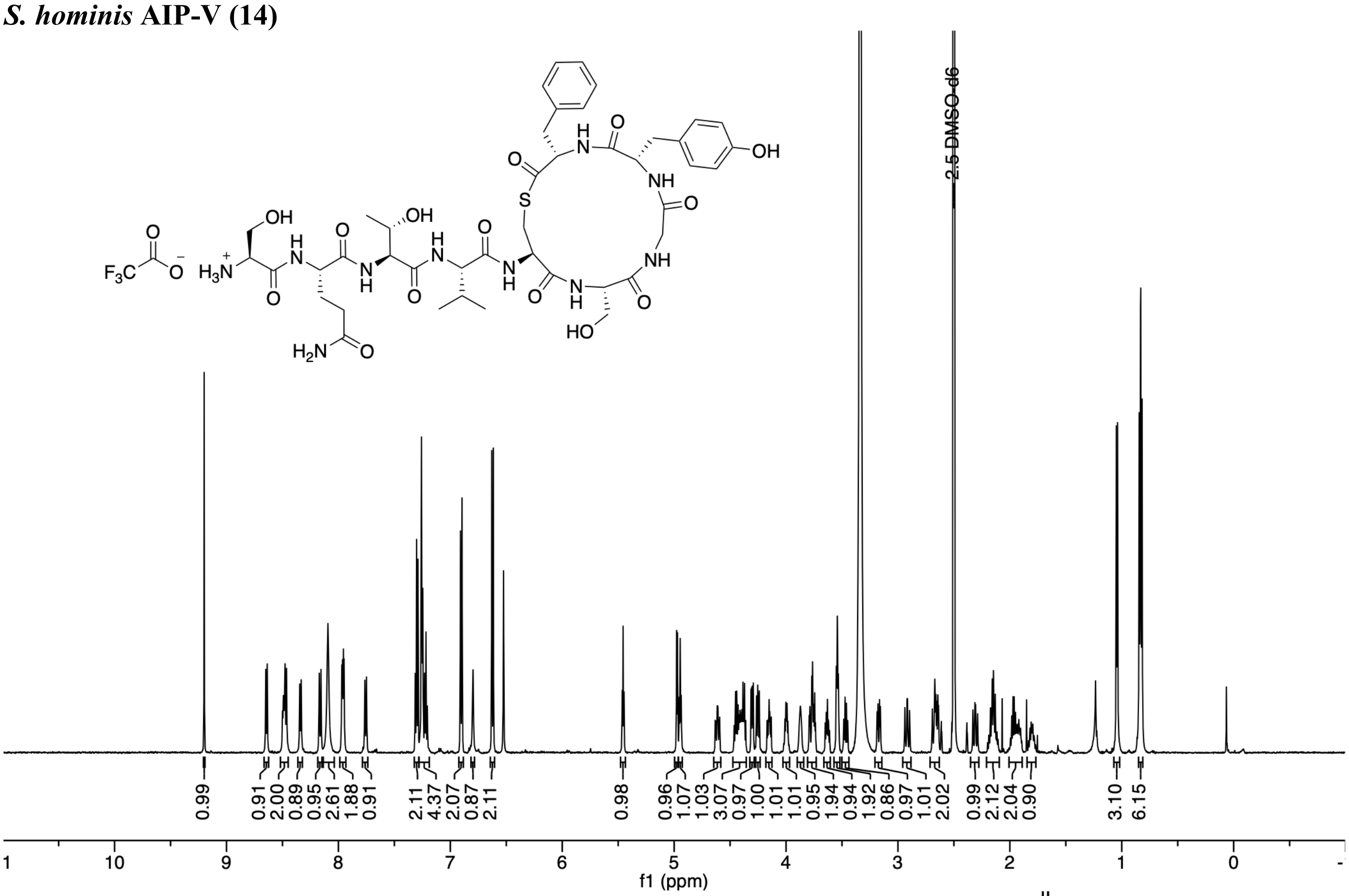

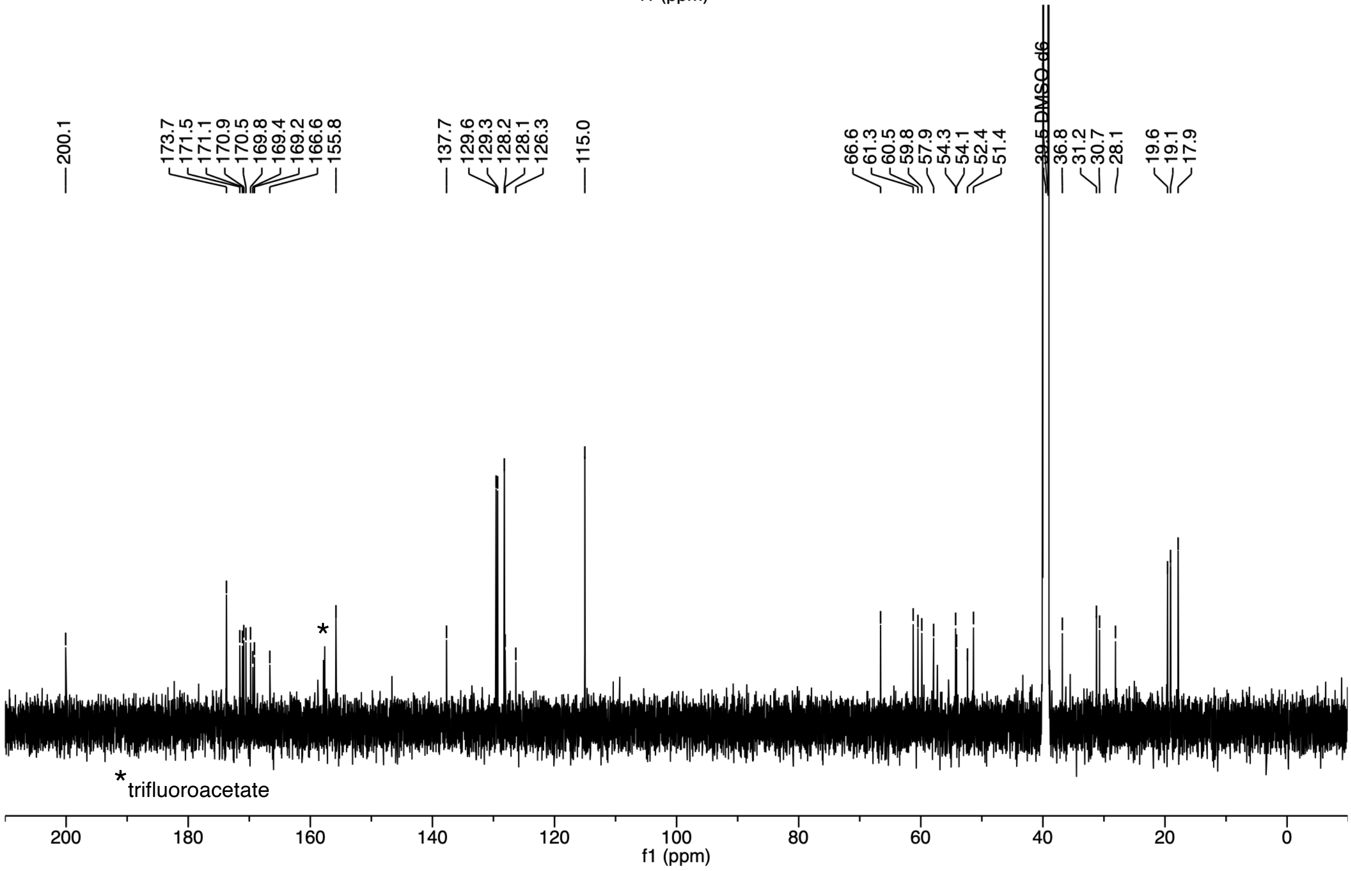

***S. capitis* AIP-I (15)**

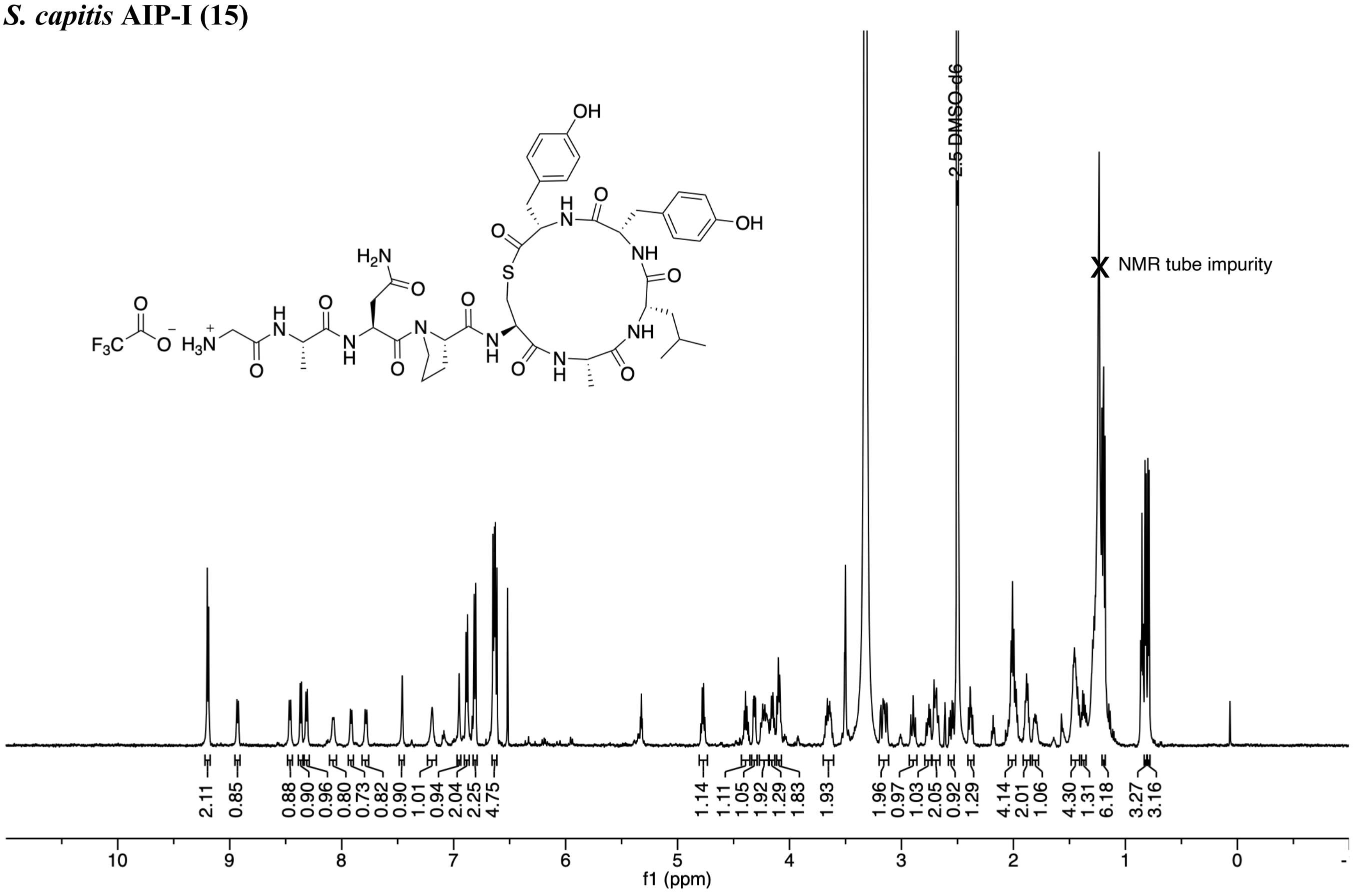

***S. warneri* AIP-II (18)**

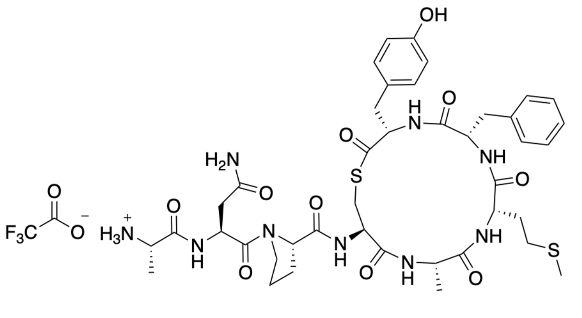

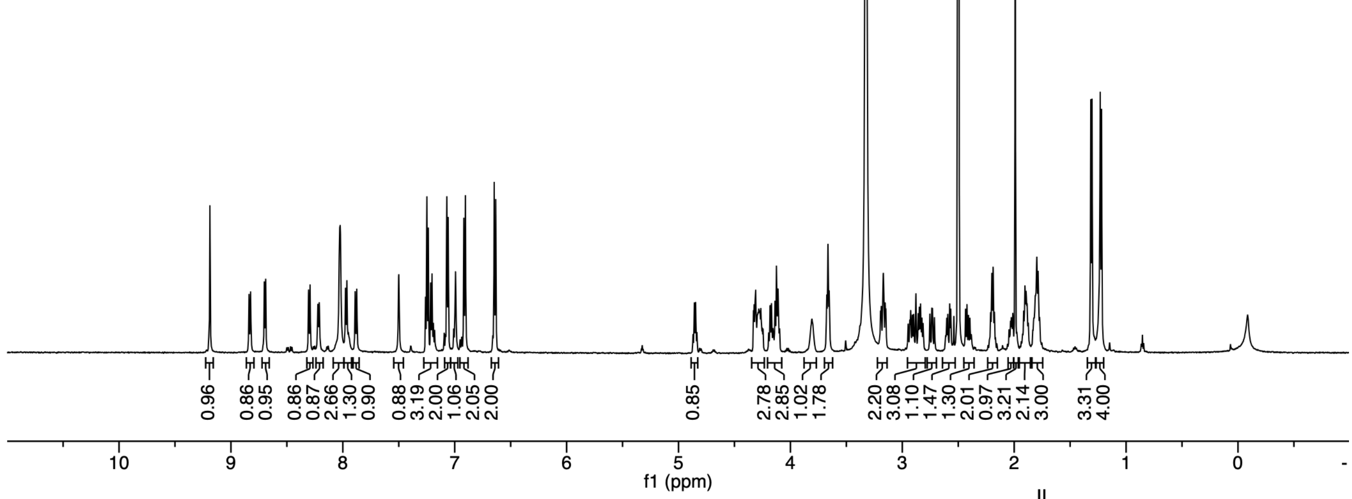

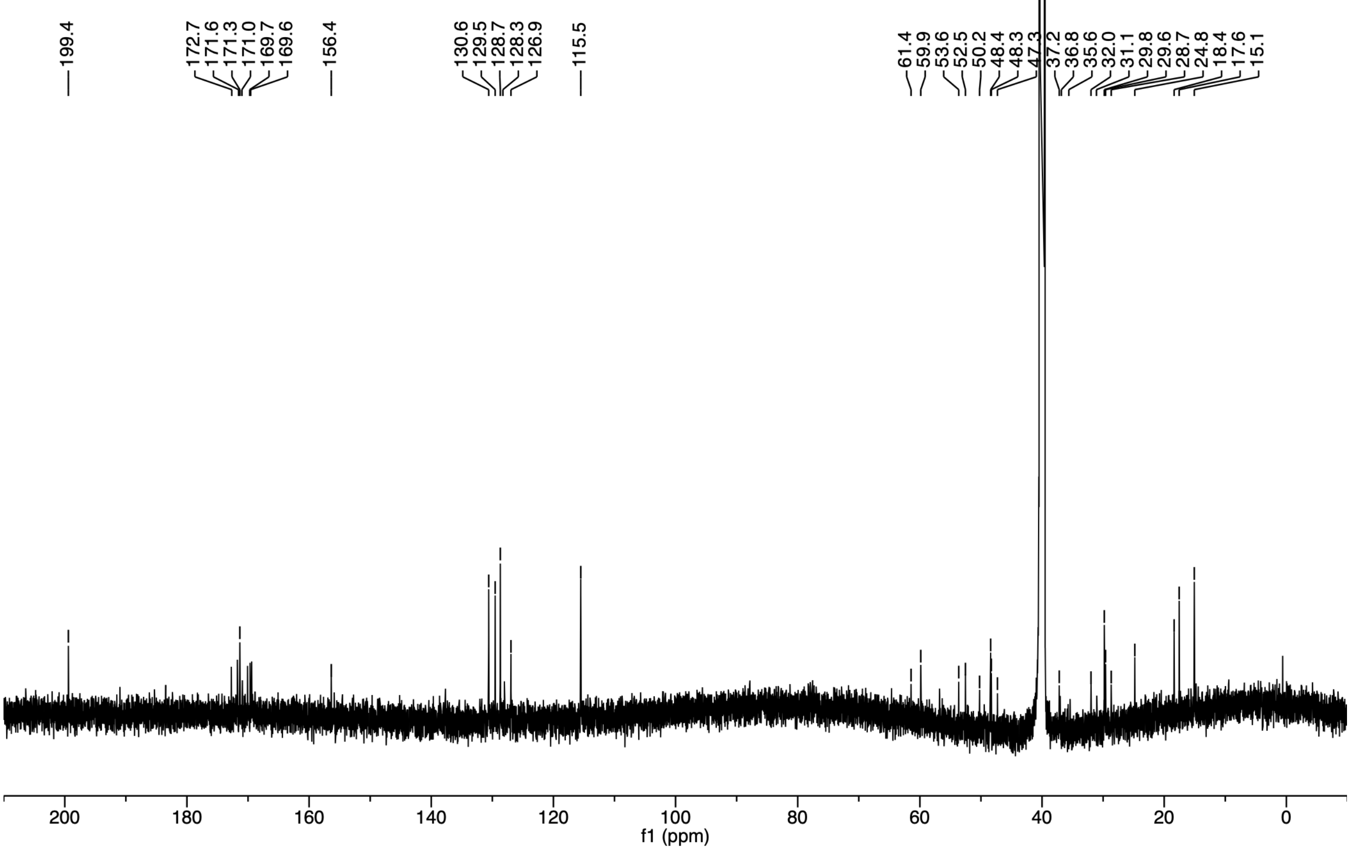

***S. cohnii* AIP-I (19)**

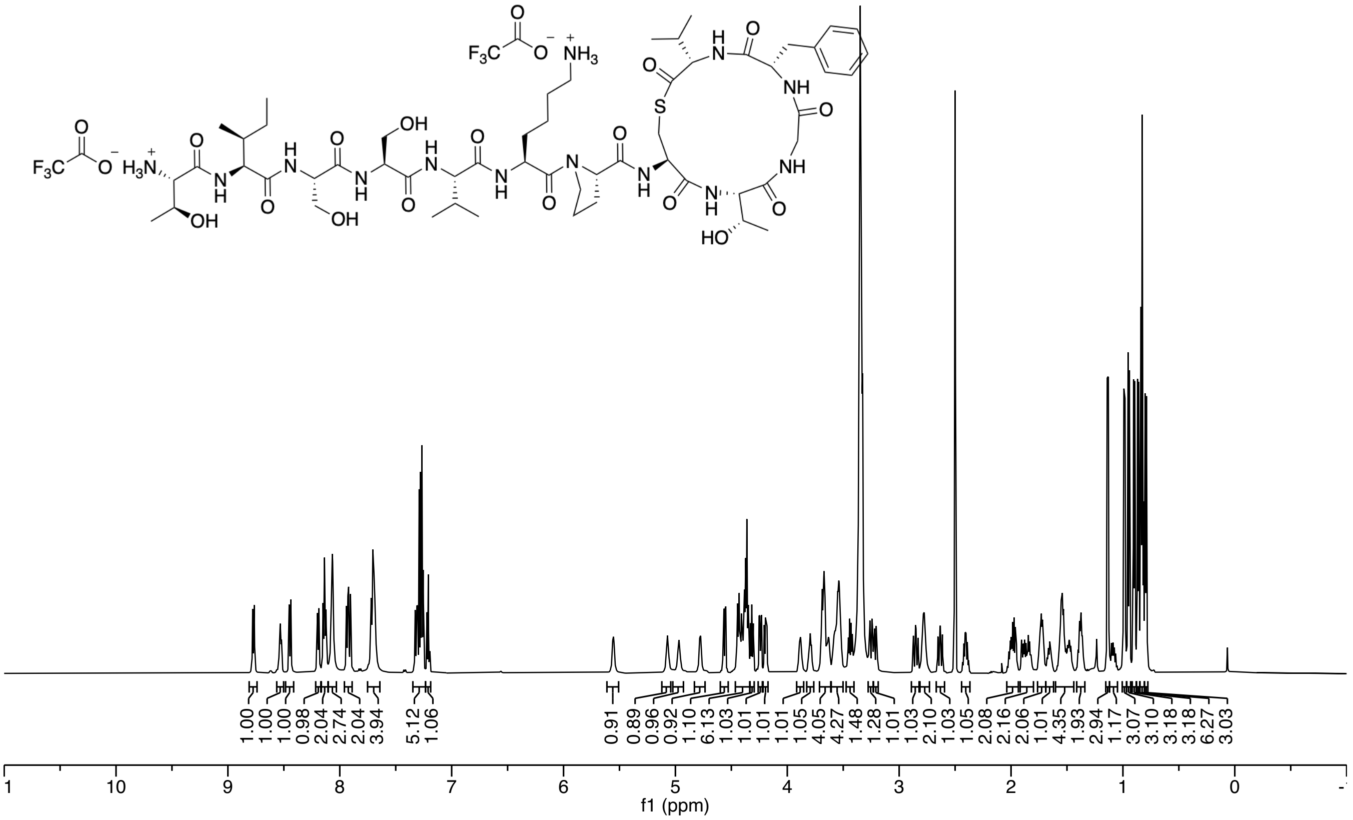

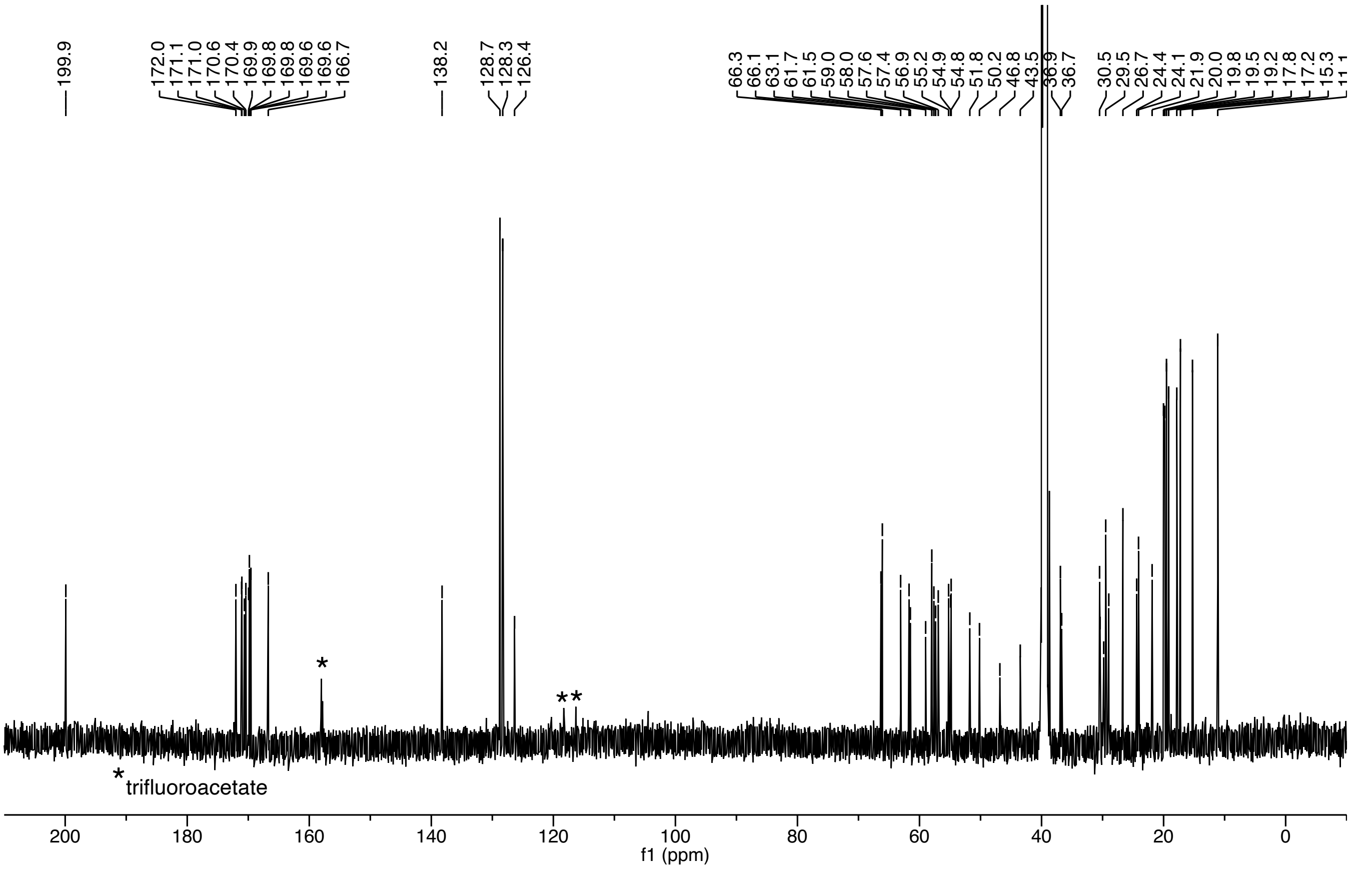

***S. caprae* AIP-I (21)**

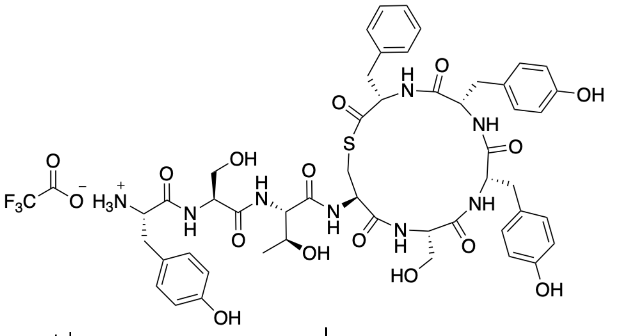

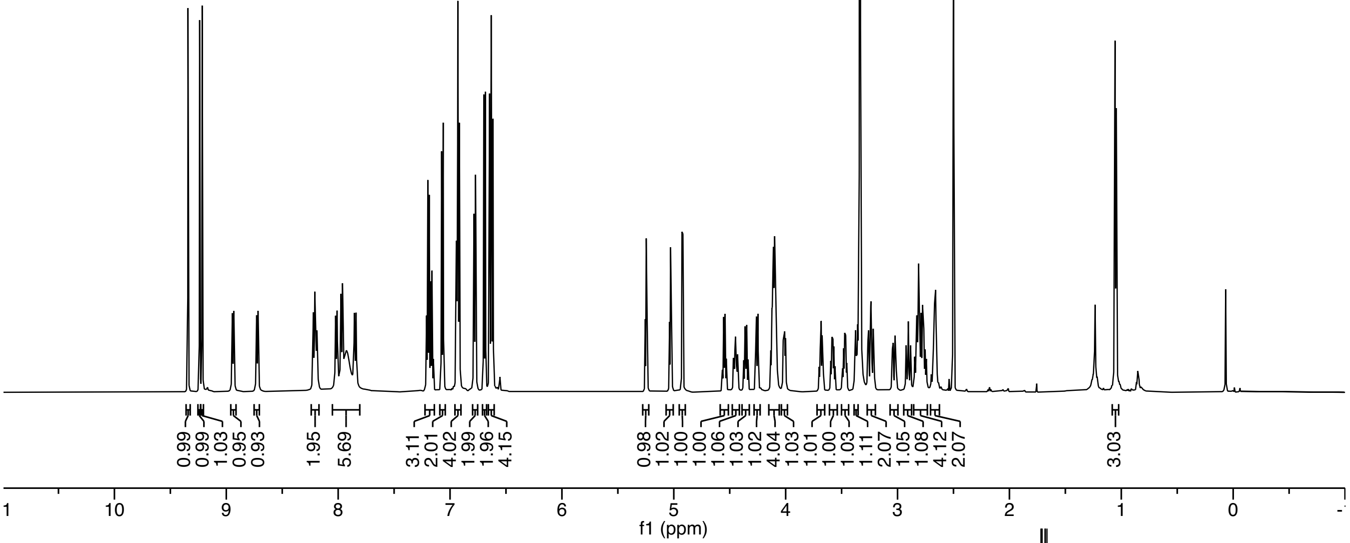

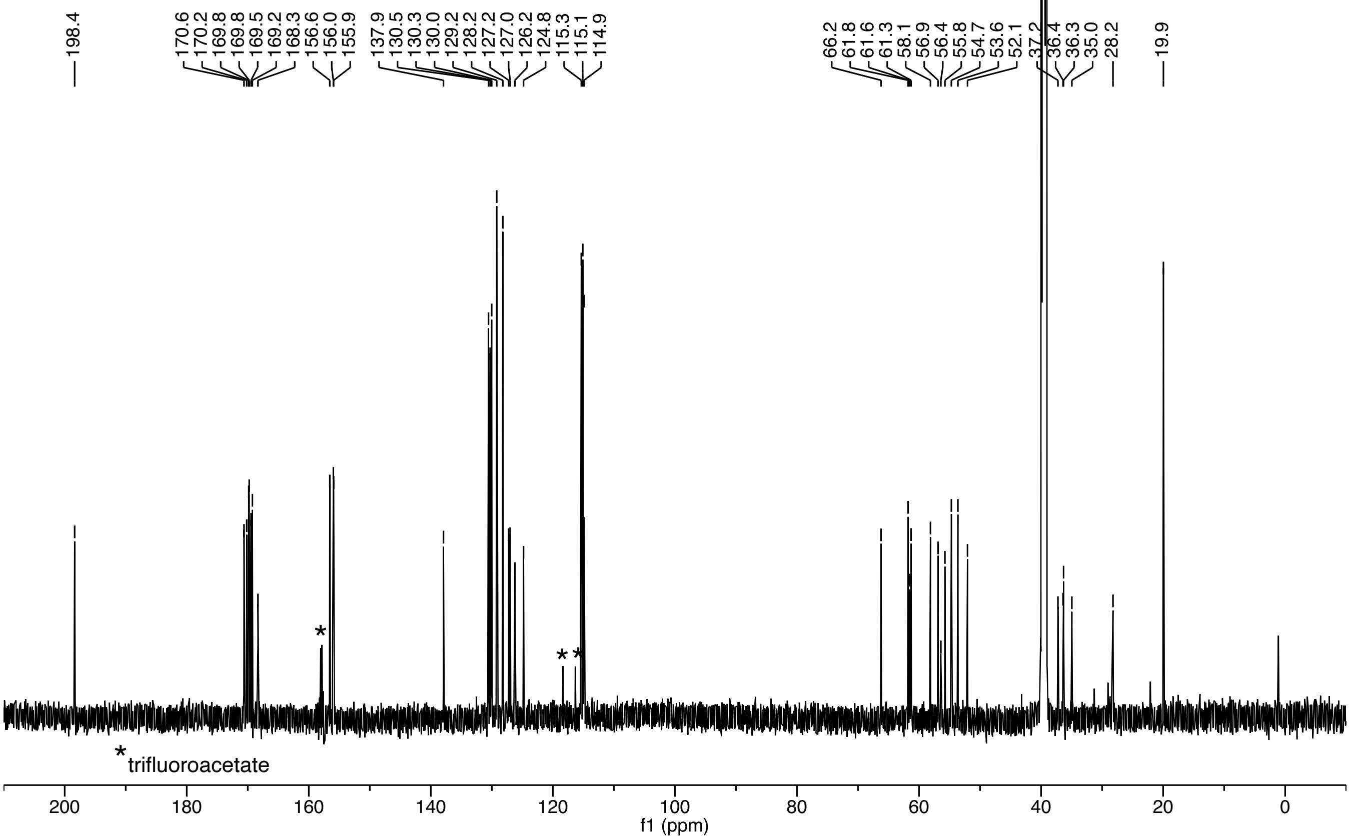

***S. pasteuri* AIP-I (22)**

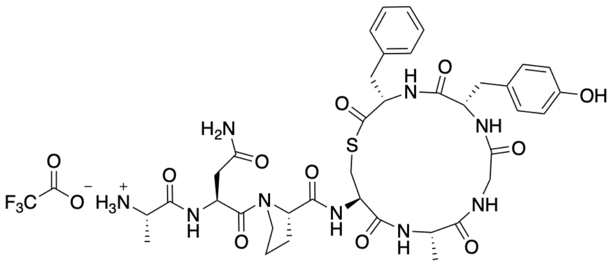

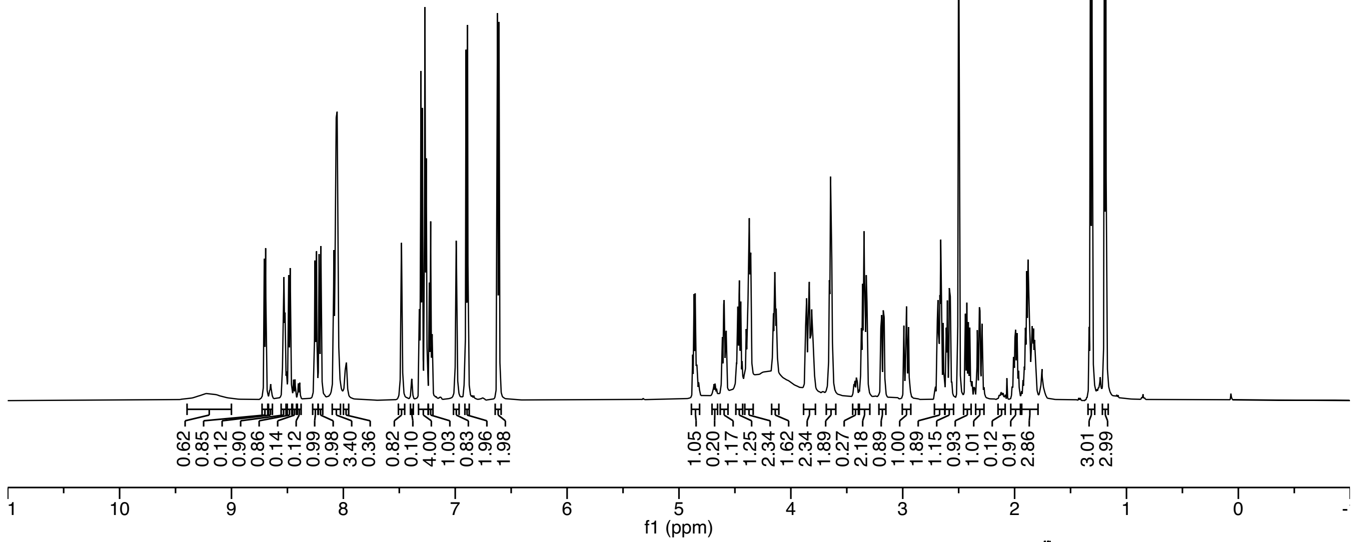

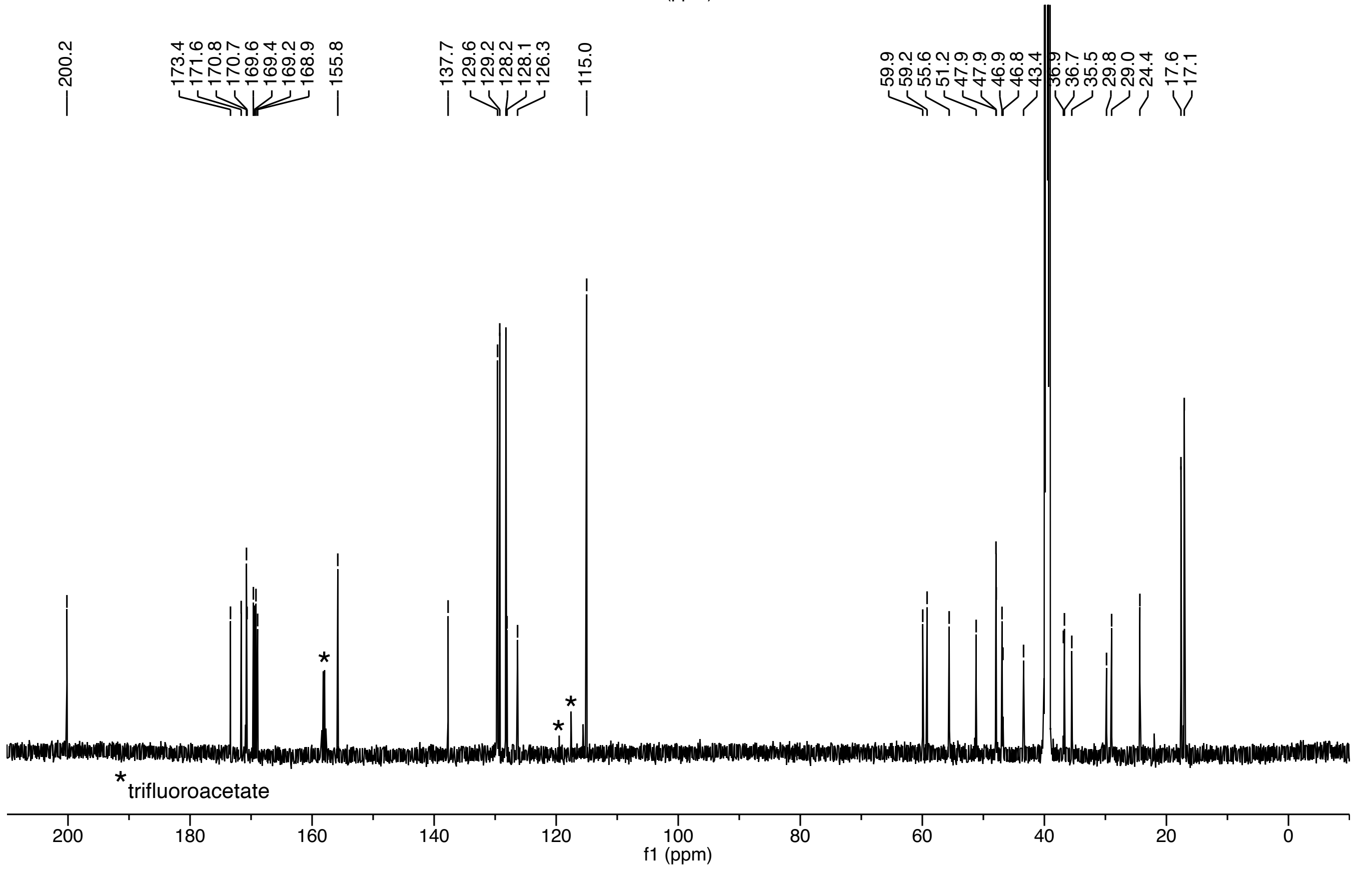

***S. devriesei* AIP-II (23)**

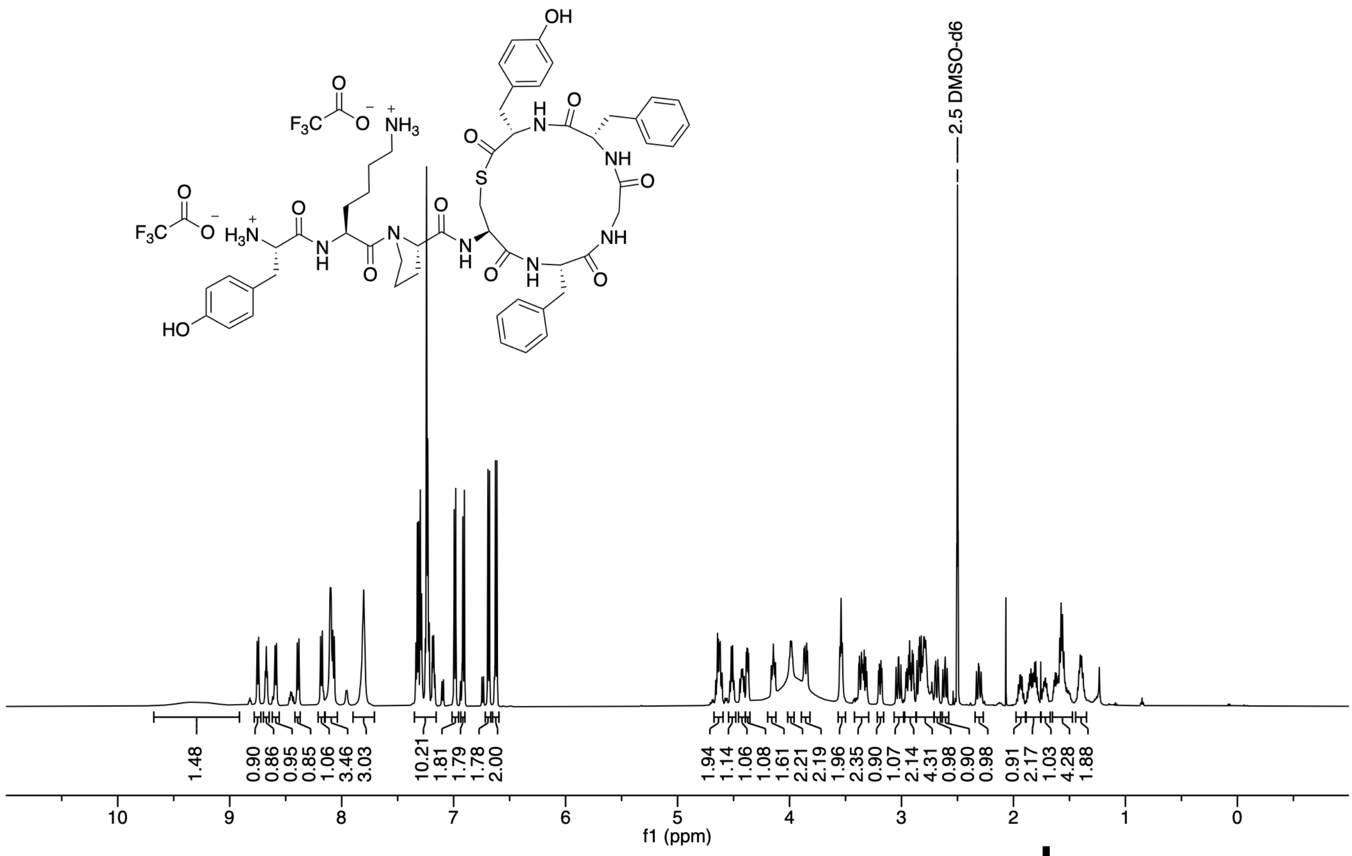

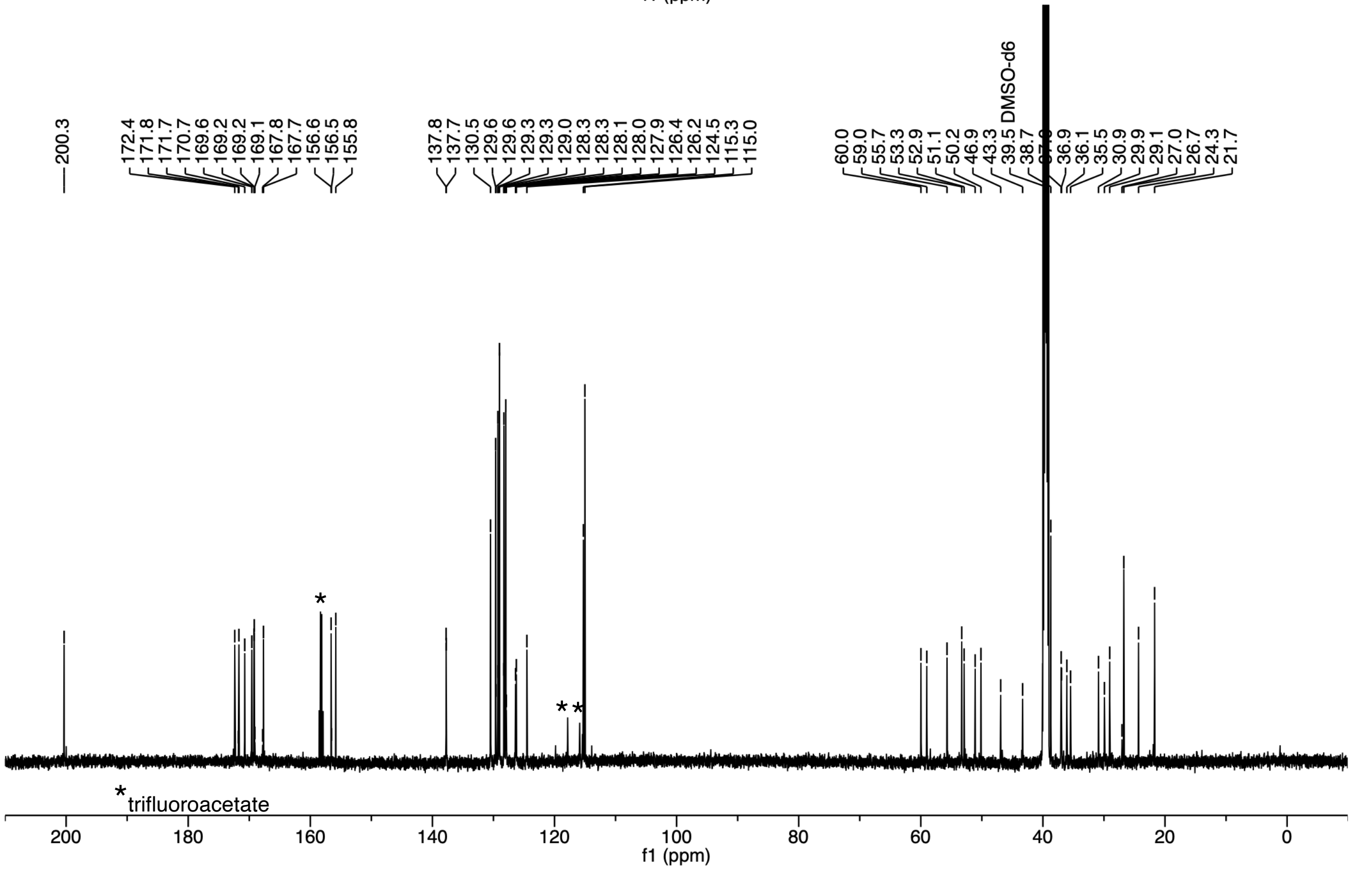

***S. succinus* AIP-I (24)**

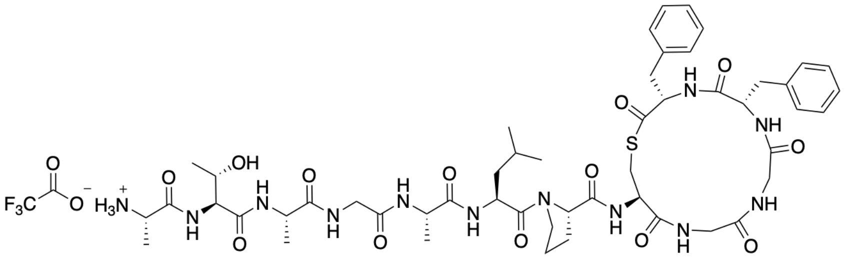

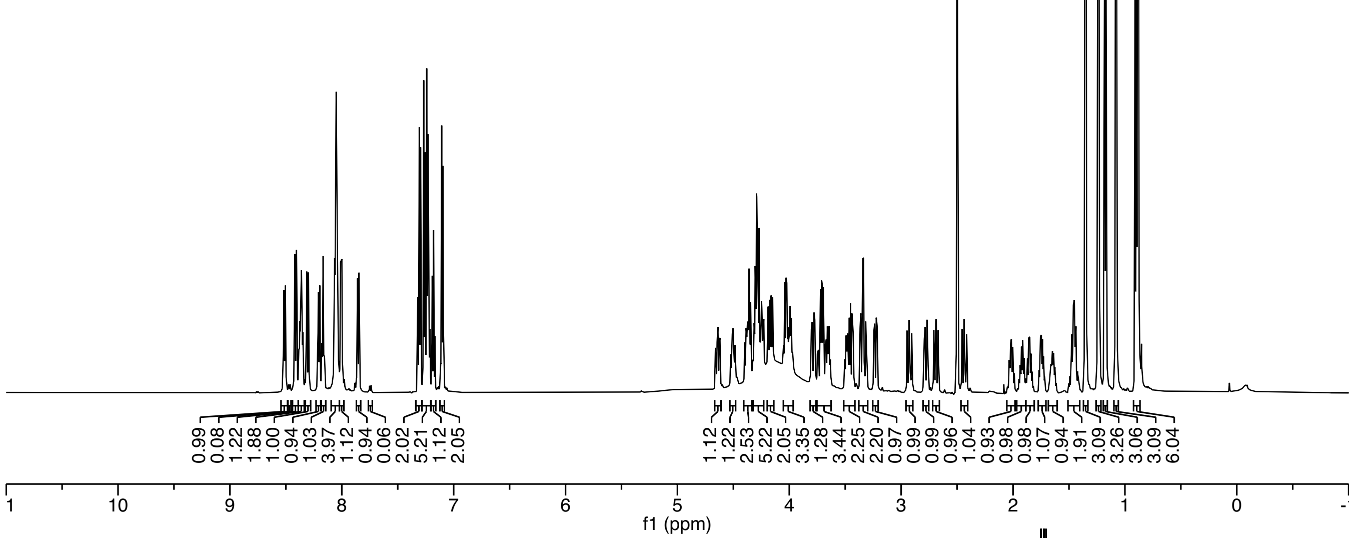

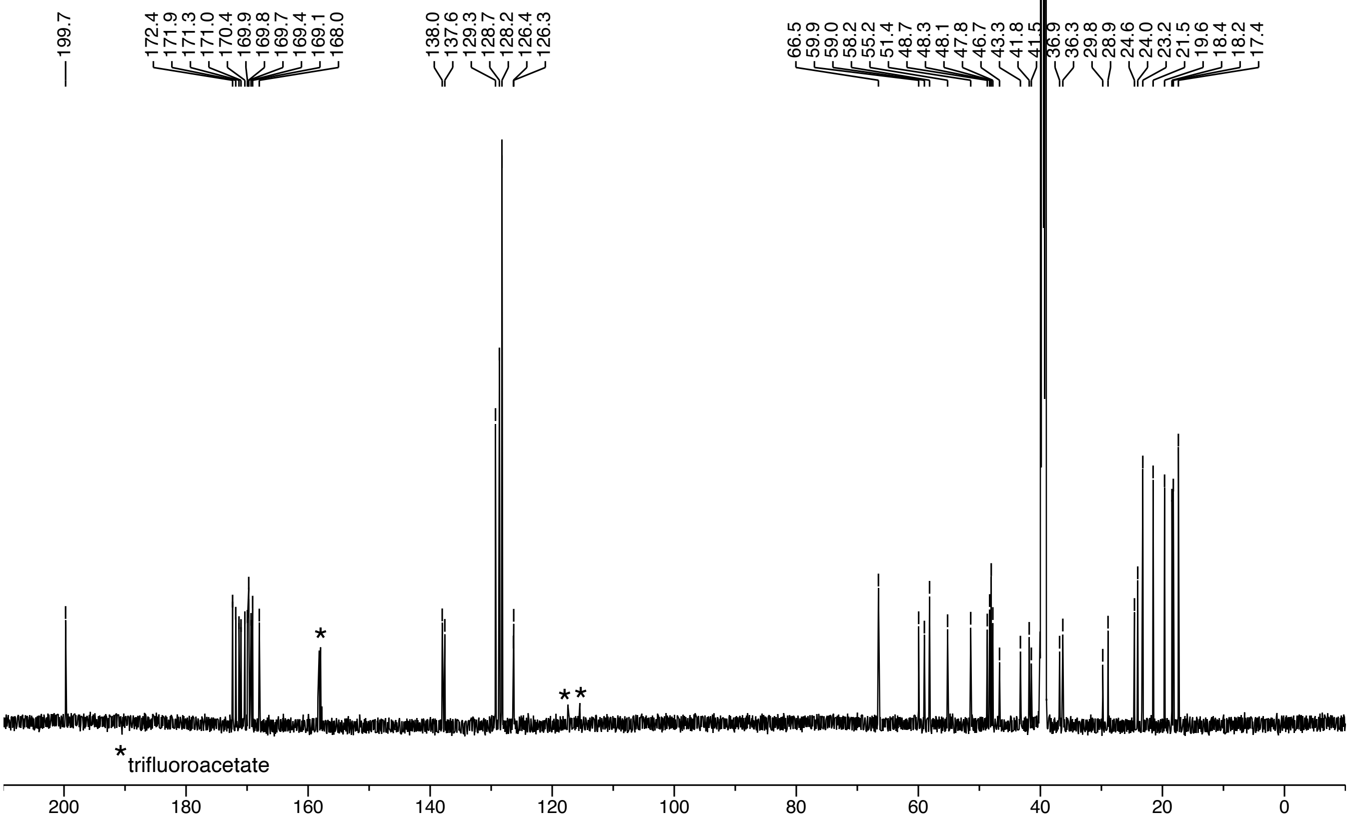

***S. equorum* AIP-I (25)**

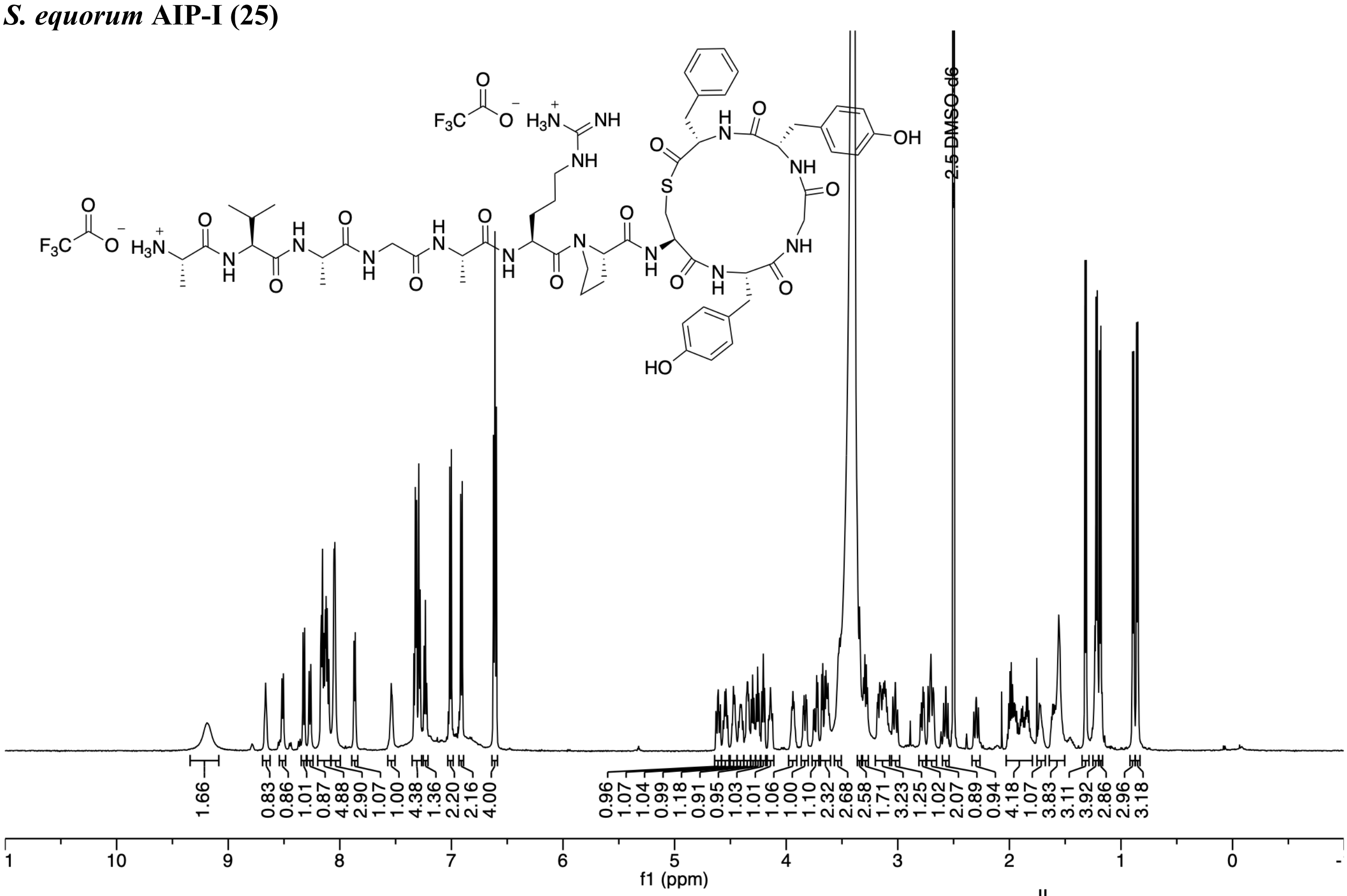

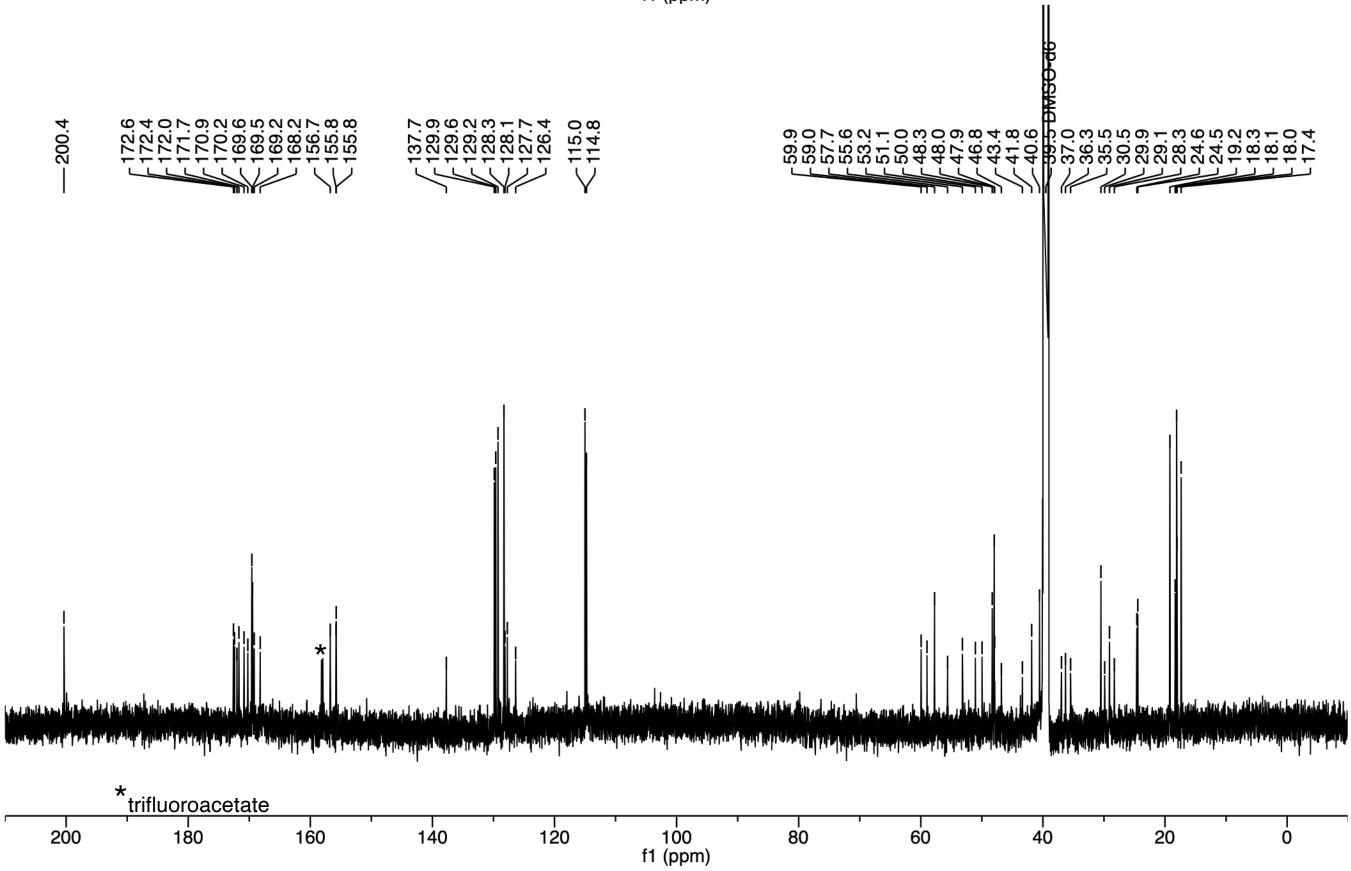

***S. equorum* AIP-II (26)**

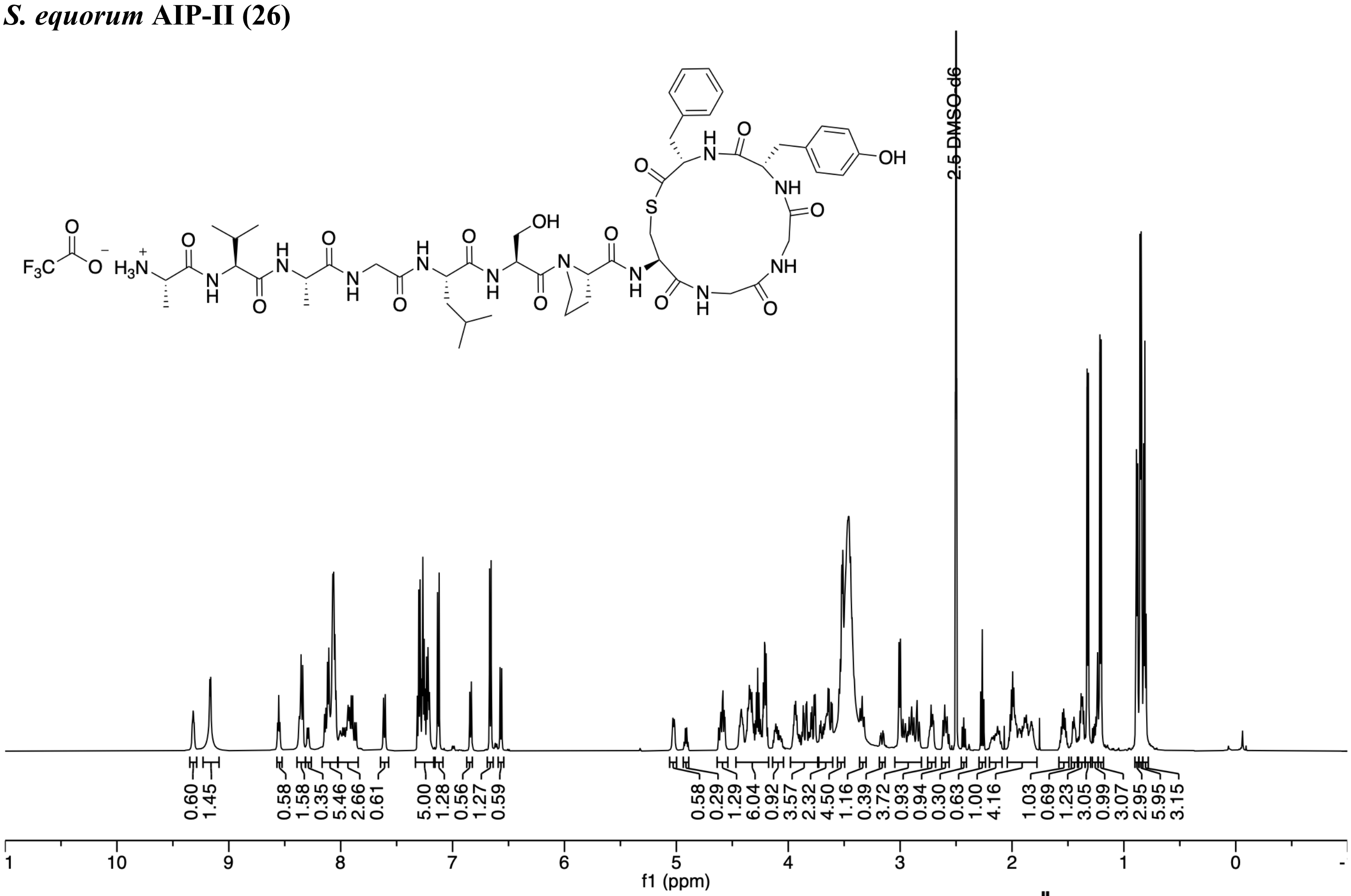

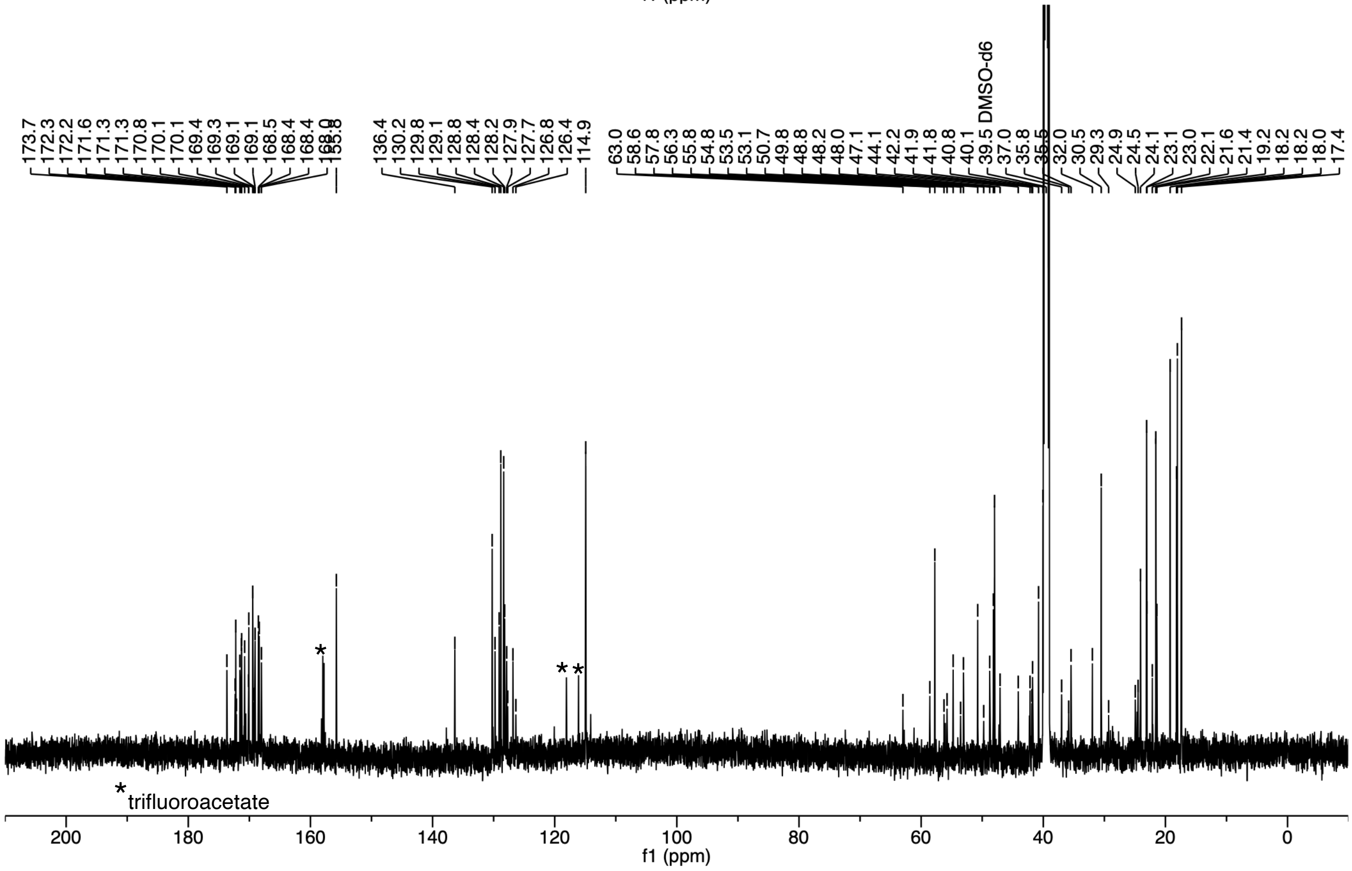

***S. intermedius* AIP-I (32)**

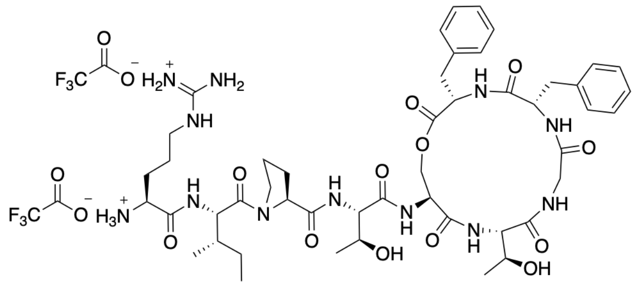

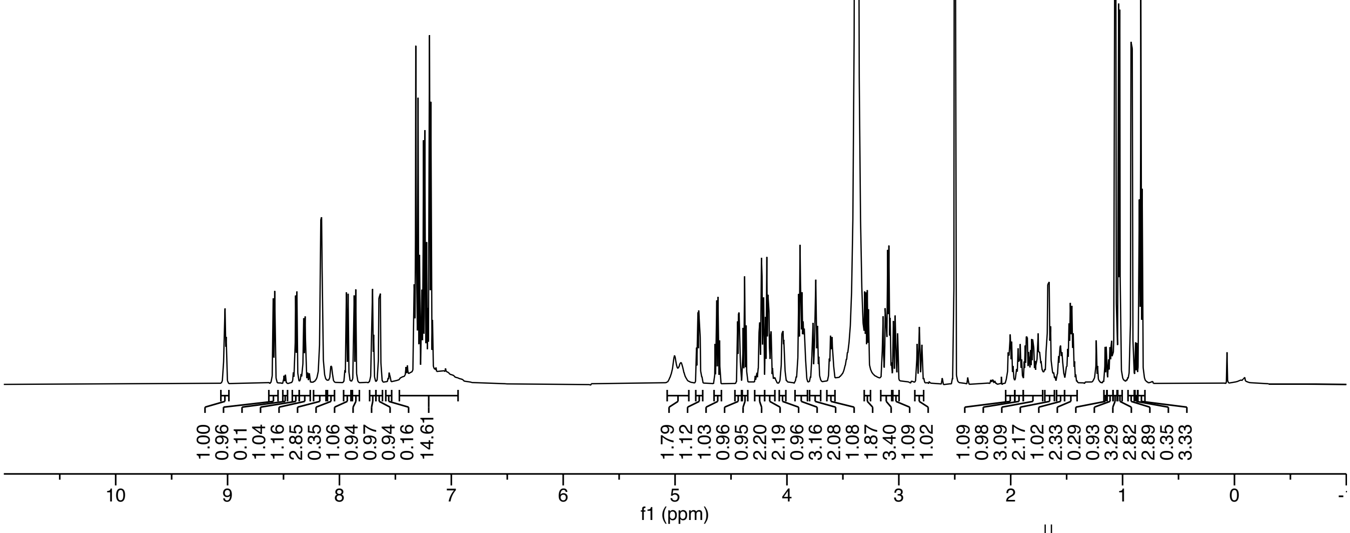

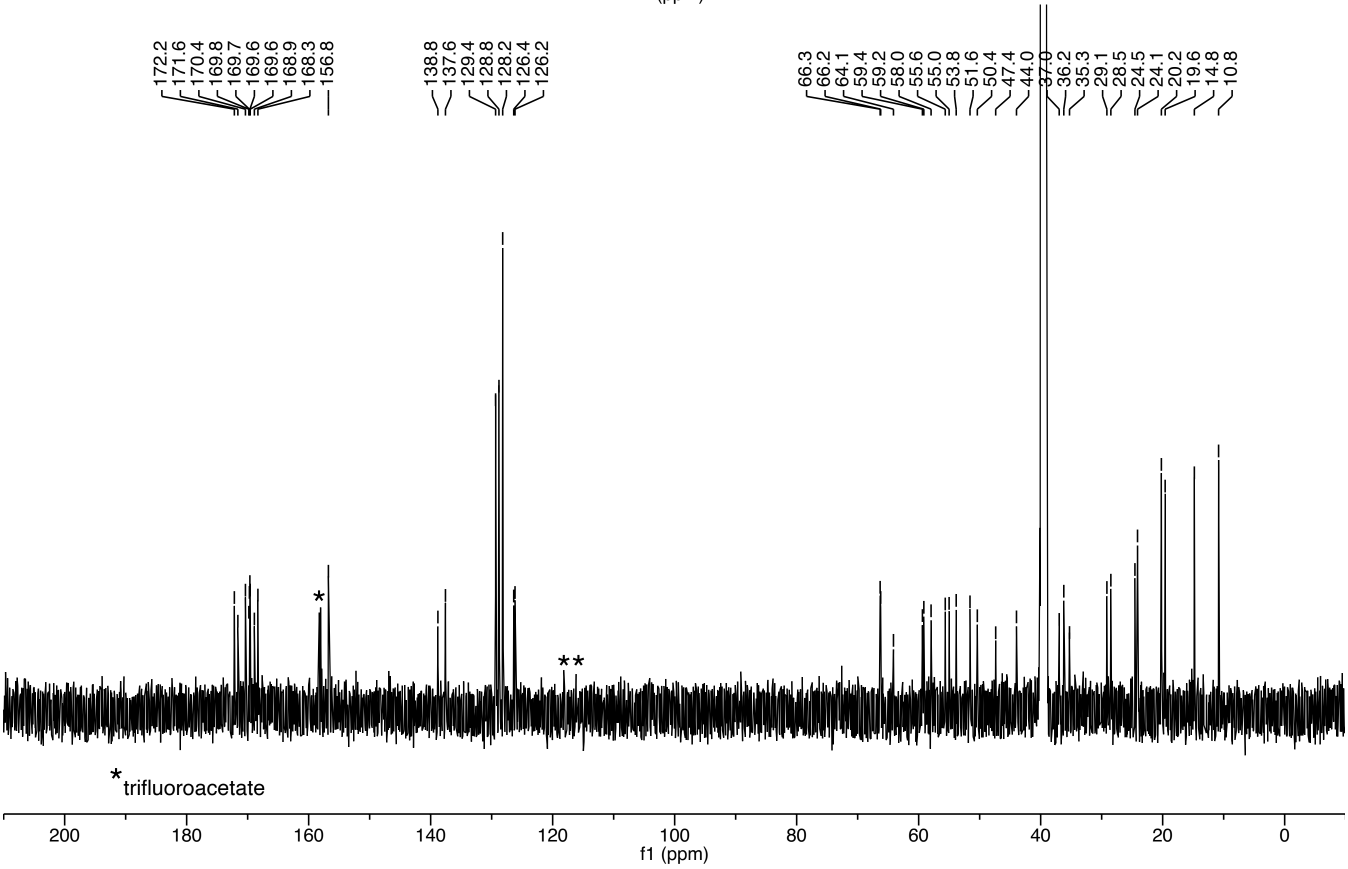

***S. simulans* AIP-II (34)**

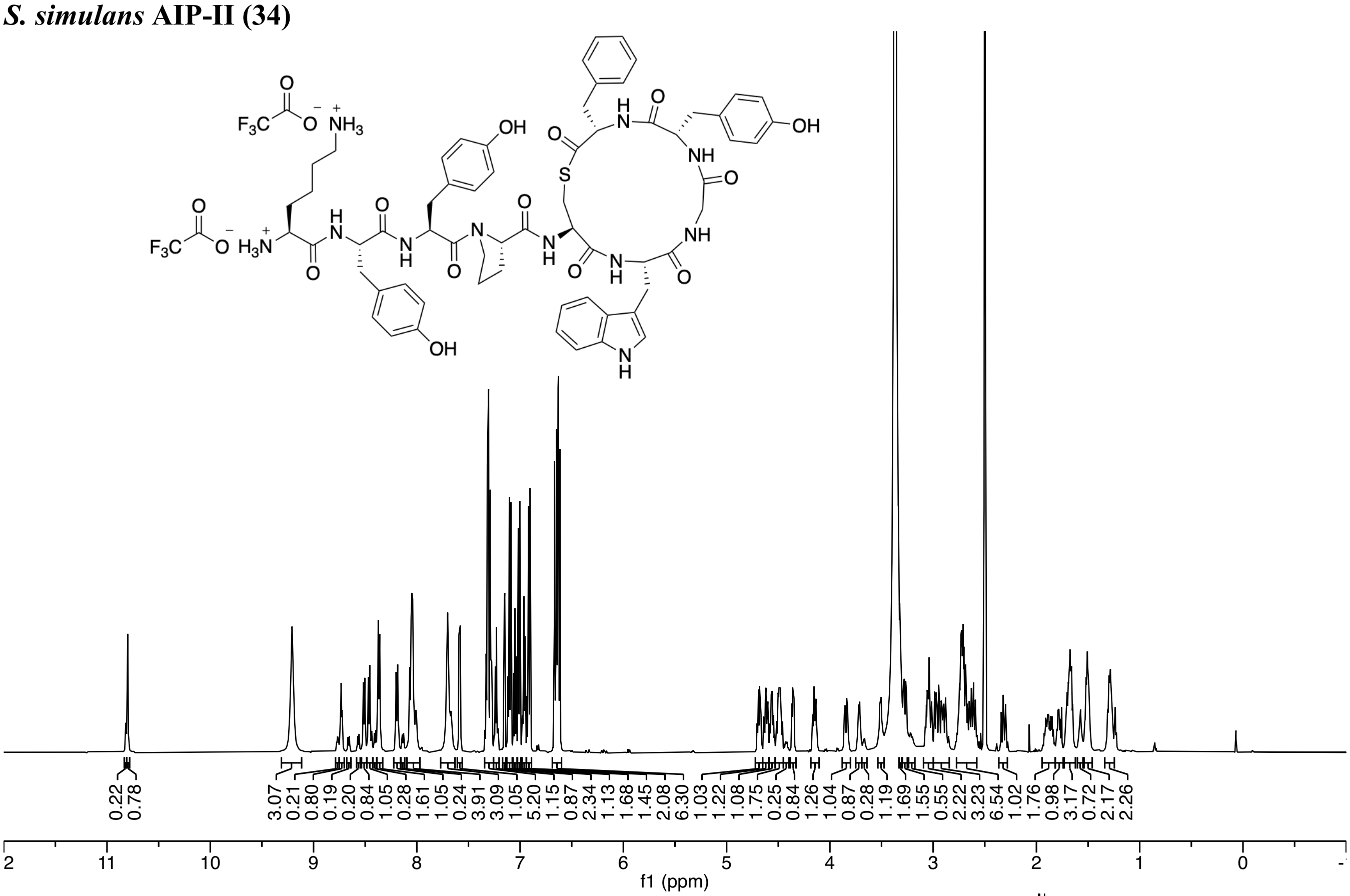

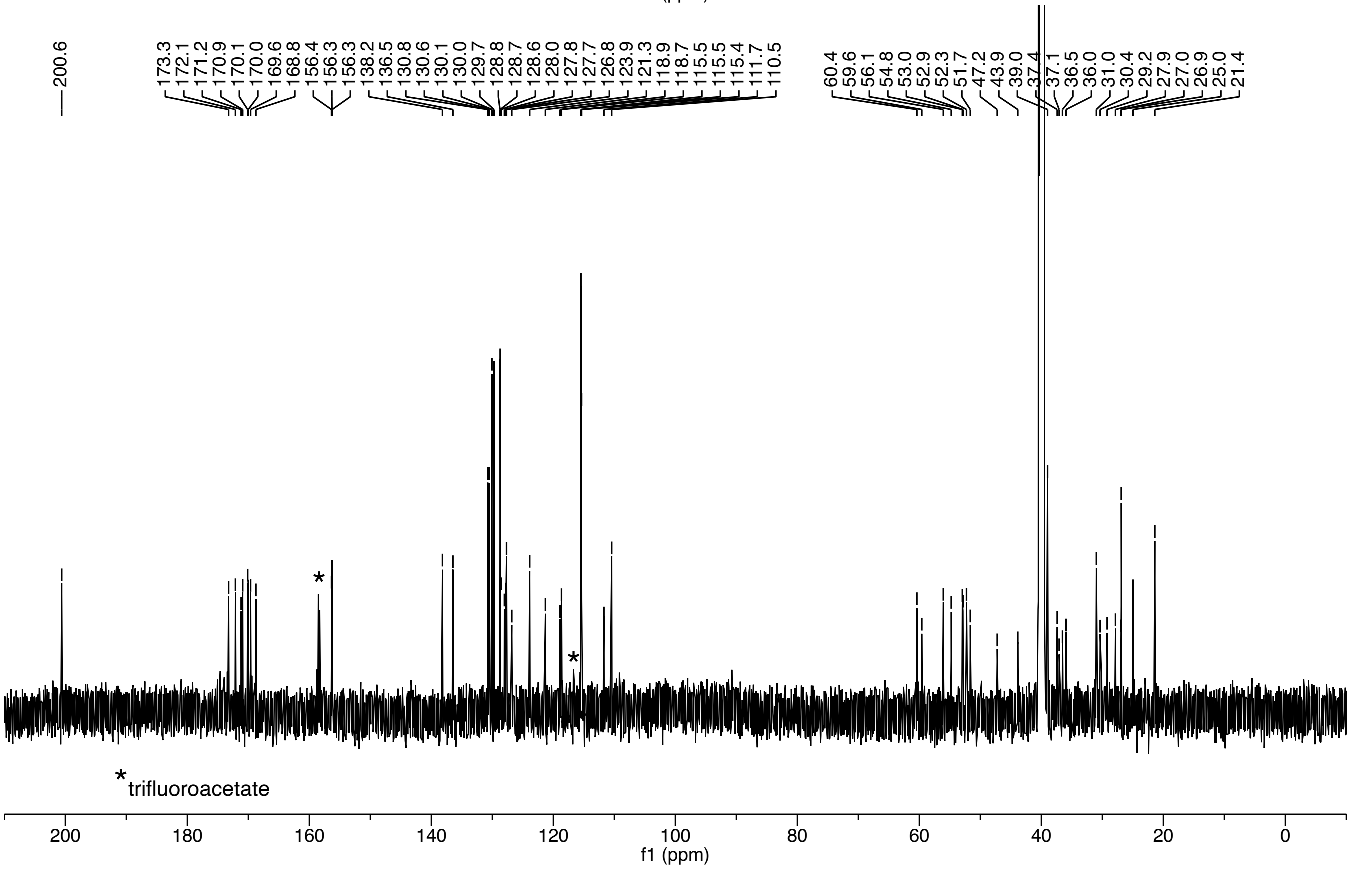

***S. simulans* AIP-III (35)**

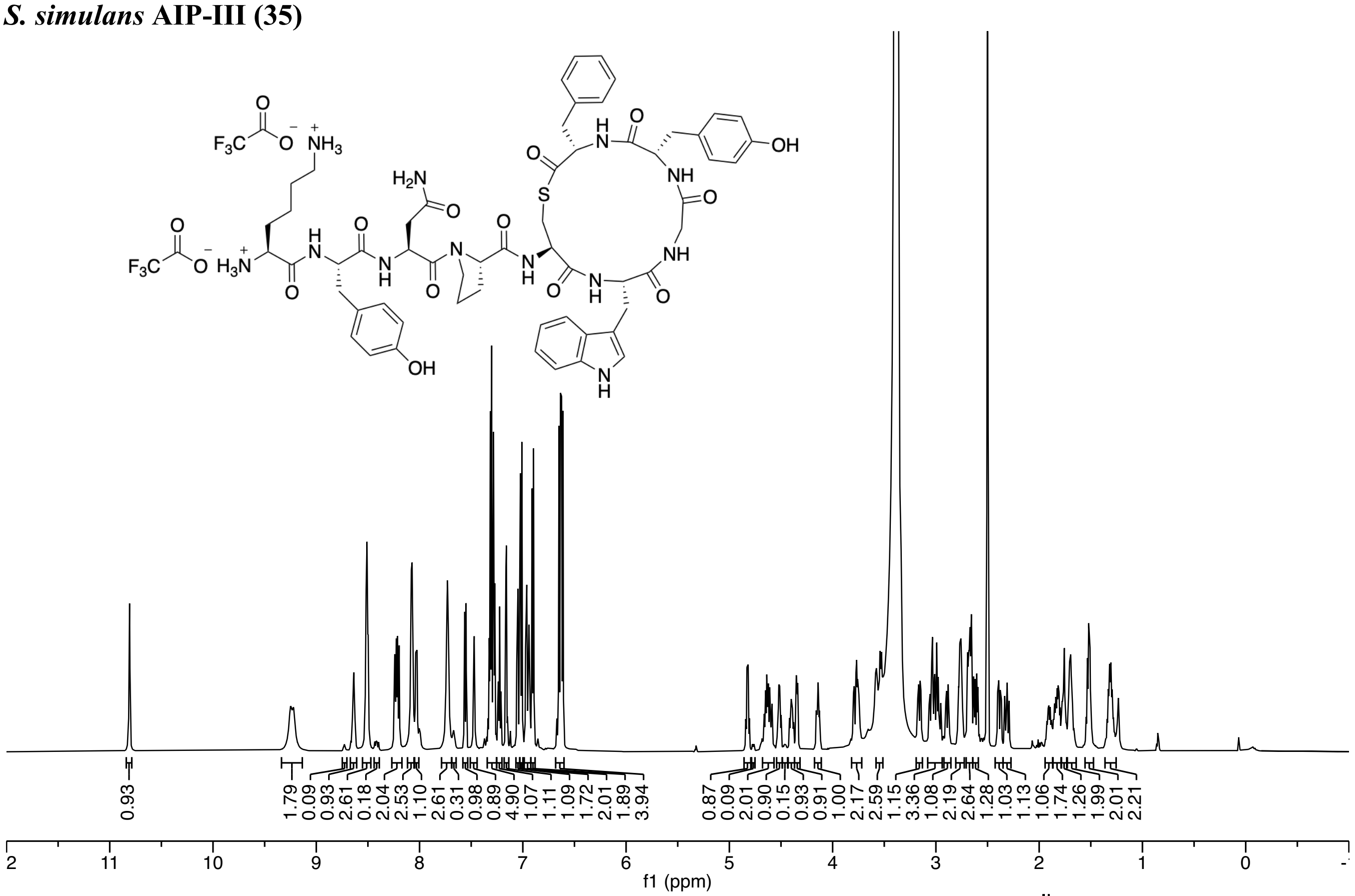

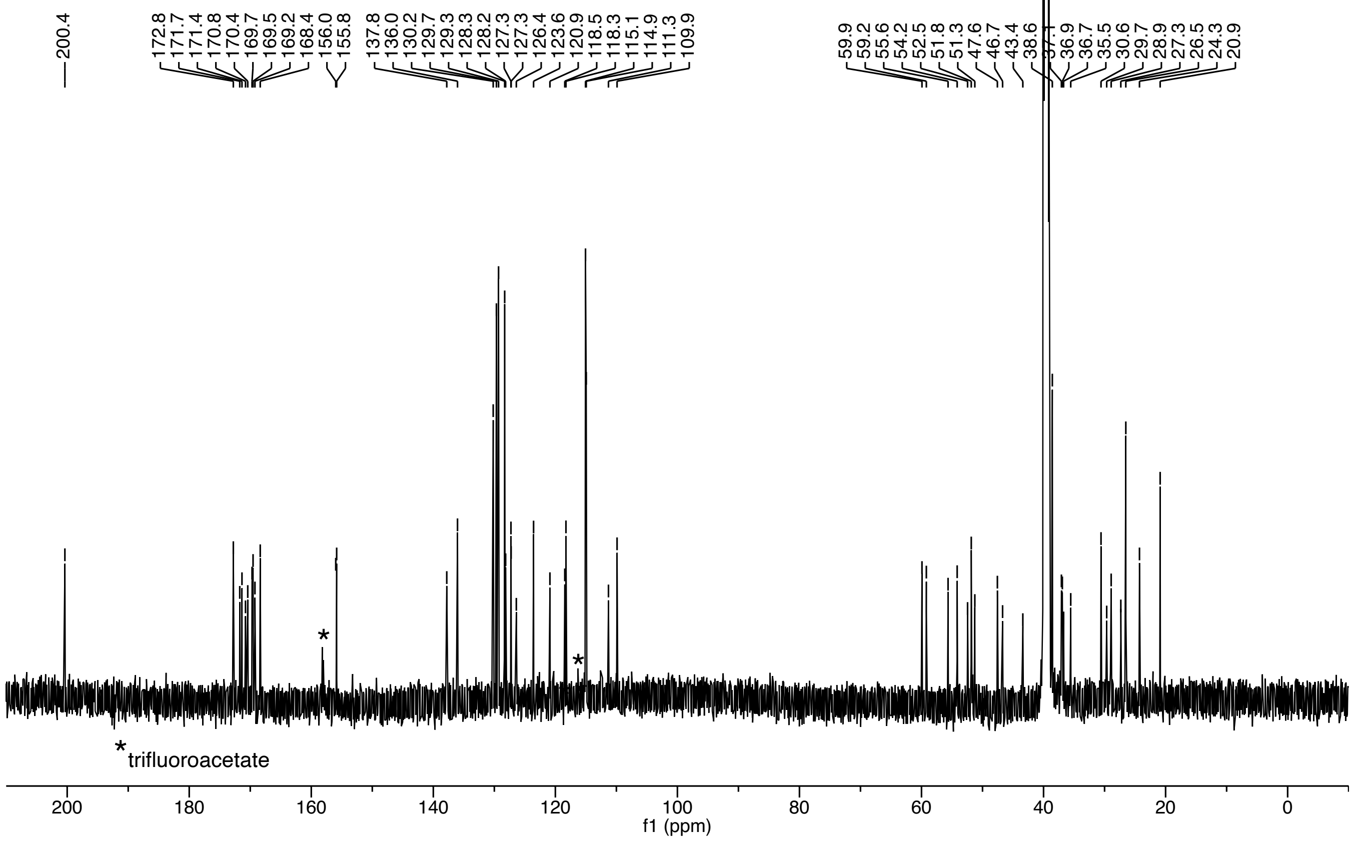

#### 8. Supplementary references

(1) Gless, B. H.; Bojer, M. S.; Peng, P.; Baldry, M.; Ingmer, H.; Olsen, C. A., Identification of autoinducing thiodepsipeptides from staphylococci enabled by native chemical ligation. *Nat. Chem.* **2019**, *11*, 463-469.

(2) Mayville, P.; Ji, G.; Beavis, R.; Yang, H.; Goger, M.; Novick, R. P.; Muir, T. W., Structure-activity analysis of synthetic autoinducing thiolactone peptides from *Staphylococcus aureus* responsible for virulence. *Proc. Natl. Acad. Sci. U. S. A.* **1999**, *96*, 1218-1223.

(3) Lyon, G. J.; Mayville, P.; Muir, T. W.; Novick, R. P., Rational design of a global inhibitor of the virulence response in *Staphylococcus aureus*, based in part on localization of the site of inhibition to the receptor-histidine kinase, AgrC. *Proc. Natl. Acad. Sci. U. S. A.* **2000**, *97*, 13330-13335.

(4) Lyon, G. J.; Wright, J. S.; Muir, T. W.; Novick, R. P., Key determinants of receptor activation in the *agr* autoinducing peptides of *Staphylococcus aureus*. *Biochemistry* **2002**, *41*, 10095-10104.

(5) Tal-Gan, Y.; Stacy, D. M.; Foegen, M. K.; Koenig, D. W.; Blackwell, H. E., Highly potent inhibitors of quorum sensing in *Staphylococcus aureus* revealed through a systematic synthetic study of the group-III autoinducing peptide. *J. Am. Chem. Soc.* **2013**, *135*, 7869-7882.

(6) Johnson, J. G.; Wang, B.; Debelouchina, G. T.; Novick, R. P.; Muir, T. W., Increasing AIP macrocycle size reveals key features of *agr* activation in *Staphylococcus aureus*. *ChemBioChem* **2015**, *16*, 1093-1100.

(7) Tal-Gan, Y.; Stacy, D. M.; Blackwell, H. E., N-Methyl and peptoid scans of an autoinducing peptide reveal new structural features required for inhibition and activation of AgrC quorum sensing receptors in *Staphylococcus aureus*. *Chem. Commun.* **2014**, *50*, 3000-3003.

(8) Gless, B. H.; Peng, P.; Pedersen, K. D.; Gotfredsen, C. H.; Ingmer, H.; Olsen, C. A., Structure-activity relationship study based on autoinducing peptide (AIP) from dog pathogen *S. schleiferi*. *Org. Lett.* **2017**, *19*, 5276–5279.

(9) Otto, M.; Echner, H.; Voelter, W.; Götz, F., Pheromone Cross-Inhibition between *Staphylococcus aureus* and *Staphylococcus epidermidis*. *Infect. Immun.* **2001**, *69*, 1957-1960.

(10) Tal-Gan, Y.; Ivancic, M.; Cornilescu, G.; Blackwell, H. E., Characterization of structural elements in native autoinducing peptides and non-native analogues that permit the differential modulation of AgrC-type quorum sensing receptors in Staphylococcus aureus. *Org. Biomol. Chem.* **2016**, *14*, 113-21.

(11) Yang, T.; Tal-Gan, Y.; Paharik, A. E.; Horswill, A. R.; Blackwell, H. E., Structure-function analyses of a *Staphylococcus epidermidis* autoinducing peptide reveals motifs critical for AgrC-type receptor modulation. *ACS Chem. Biol.* **2016**, *11*, 1982-91.

(12) Williams, M. R.; Costa, S. K.; Zaramela, L. S.; Khalil, S.; Todd, D. A.; Winter, H. L.; Sanford, J. A.; O'Neill, A. M.; Liggins, M. C.; Nakatsuji, T.; Cech, N. B.; Cheung, A. L.; Zengler, K.; Horswill, A. R.; Gallo, R. L., Quorum sensing between bacterial species on the skin protects against epidermal injury in atopic dermatitis. *Sci. Transl. Med.* **2019**, *11*, eaat8329.

(13) Severn Morgan, M.; Williams Michael, R.; Shahbandi, A.; Bunch Zoie, L.; Lyon Laurie, M.; Nguyen, A.; Zaramela Livia, S.; Todd Daniel, A.; Zengler, K.; Cech Nadja, B.; Gallo Richard, L.; Horswill Alexander, R., The Ubiquitous Human Skin Commensal Staphylococcus hominis Protects against Opportunistic Pathogens. *mBio* **2022**, *13*, e00930-22.

(14) Severn, M. M.; Cho, Y.-S. K.; Manzer, H. S.; Bunch, Z. L.; Shahbandi, A.; Todd, D. A.; Cech, N. B.; Horswill, A. R., The Commensal Staphylococcus warneri Makes Peptide Inhibitors of MRSA Quorum Sensing that Protect Skin from Atopic or Necrotic Damage. *J. Invest. Dermatol.* **2022**, *142*, 3349-3352.e5.

(15) Paharik, A. E.; Parlet, C. P.; Chung, N.; Todd, D. A.; Rodriguez, E. I.; Van Dyke, M. J.; Cech, N. B.; Horswill, A. R., Coagulase-negative staphylococcal strain prevents *Staphylococcus aureus* colonization and skin infection by blocking quorum sensing. *Cell Host Microbe* **2017**, *22*, 1-11.

(16) Ji, G.; Pei, W.; Zhang, L.; Qiu, R.; Lin, J.; Benito, Y.; Lina, G.; Novick, R. P., Staphylococcus intermedius Produces a Functional agr Autoinducing Peptide Containing a Cyclic Lactone. *J. Bacteriol.* **2005**, *187*, 3139-3150.

(17) Brown, M. M.; Kwiecinski, J. M.; Cruz, L. M.; Shahbandi, A.; Todd, D. A.; Cech, N. B.; Horswill, A. R., Novel Peptide from Commensal *Staphylococcus simulans* Blocks Methicillin-Resistant *Staphylococcus aureus* Quorum Sensing and Protects Host Skin from Damage. *Antimicrob. Agents Chemother.* **2020**, *64*, e00172-20.
(18) Olson, M. E.; Todd, D. A.; Schaeffer, C. R.; Paharik, A. E.; Van Dyke, M. J.; Büttner, H.; Dunman, P. M.; Rohde, H.; Cech, N. B.; Fey, P. D.; Horswill, A. R., *Staphylococcus epidermidis agr* quorum-sensing system: signal identification, cross talk, and importance in colonization. *J. Bacteriol.* **2014**, *196*, 3482-93.
(19) Gordon, C. P.; Olson, S. D.; Lister, J. L.; Kavanaugh, J. S.; Horswill, A. R., Truncated Autoinducing Peptides as Antagonists of *Staphylococcus lugdunensis* Quorum Sensing. *J. Med. Chem.* **2016**, *59*, 8879-8888.
(20) Malone, C. L.; Boles, B. R.; Lauderdale, K. J.; Thoendel, M.; Kavanaugh, J. S.; Horswill, A. R., Fluorescent reporters for Staphylococcus aureus. *J. Microbiol. Methods* **2009**, *77*, 251-260.
(21) Chakraborty, A.; Sharma, A.; Albericio, F.; de la Torre, B. G., Disulfide-Based Protecting Groups for the Cysteine Side Chain. *Org. Lett.* **2020**, *22*, 9644-9647.
(22) Geisinger, E.; Muir, T. W.; Novick, R. P., agr receptor mutants reveal distinct modes of inhibition by staphylococcal autoinducing peptides. *Proc. Natl. Acad. Sci. U. S. A.* **2009**, *106*, 1216-21.
(23) Nielsen, A.; Månsson, M.; Bojer, M. S.; Gram, L.; Larsen, T. O.; Novick, R. P.; Frees, D.; Frøkiær, H.; Ingmer, H., Solonamide B inhibits quorum sensing and reduces Staphylococcus aureus mediated killing of human neutrophils. *PLoS One* **2014**, *9*, e84992.
(24) Kupferwasser, L. I.; Yeaman, M. R.; Nast, C. C.; Kupferwasser, D.; Xiong, Y.-Q.; Palma, M.; Cheung, A. L.; Bayer, A. S., Salicylic acid attenuates virulence in endovascular infections by targeting global regulatory pathways in Staphylococcus aureus. *J. Clin. Investig.* **2003**, *112*, 222-233.
(25) Bejder, B. S.; Monda, F.; Gless, B. H.; Bojer, M. S.; Ingmer, H.; Olsen, C. A., A short-lived peptide signal regulates cell-to-cell communication in Listeria monocytogenes. *Commun. Biol.* **2024**, *7*, 942.
(26) Khoo, K. K.; Galleano, I.; Gasparri, F.; Wieneke, R.; Harms, H.; Poulsen, M. H.; Chua, H. C.; Wulf, M.; Tampé, R.; Pless, S. A., Chemical modification of proteins by insertion of synthetic peptides using tandem protein trans-splicing. *Nat. Commun.* **2020**, *11*, 2284.
(27) Coin, I.; Beyermann, M.; Bienert, M., Solid-phase peptide synthesis: from standard procedures to the synthesis of difficult sequences. *Nat. Protoc.* **2007**, *2*, 3247-3256.
